# SPIRO – the automated Petri plate imaging platform designed by biologists, for biologists

**DOI:** 10.1101/2021.03.15.435343

**Authors:** Jonas A Ohlsson, Jia Xuan Leong, Pernilla H Elander, Florentine Ballhaus, Sanjana Holla, Adrian N Dauphinee, Johan Johansson, Mark Lommel, Gero Hofmann, Staffan Betnér, Mats Sandgren, Karin Schumacher, Peter V Bozhkov, Elena A Minina

**Author notes:** **Corresponding authors:** Alyona Minina; Jonas Ohlsson; Jia Xuan Leong.

## Abstract

Phenotyping of model organisms grown on Petri plates is often carried out manually, despite the procedures being time-consuming and laborious. The main reason for this is the limited availability of automated phenotyping facilities, whereas constructing a custom automated solution can be a daunting task for biologists.

Here, we describe SPIRO, the Smart Plate Imaging Robot, an automated platform that acquires time-lapse photos of up to four vertically oriented Petri plates in a single experiment, corresponding to 192 seedlings for a typical root growth assay and up to 2500 seeds for a germination assay. SPIRO is catered specifically to biologists’ needs, requiring no engineering or programming expertise for assembly and operation. Its small footprint is optimized for standard incubators, the inbuilt green LED enables imaging under dark conditions, and remote control provides access to the data without interfering with sample growth. SPIRO’s excellent image quality is suitable for automated image processing, which we demonstrate on the example of seed germination and root growth assays. Furthermore, the robot can be easily customized for specific uses, as all information about SPIRO is released under open-source licenses.

Importantly, uninterrupted imaging allows considerably more precise assessment of seed germination parameters and root growth rates compared to manual assays. Moreover, SPIRO enables previously technically challenging assays such as phenotyping in the dark. We illustrate the benefits of SPIRO in proof-of-concept experiments which yielded a novel insight on the interplay between autophagy, nitrogen sensing and photoblastic response.

## Introduction

Manual imaging of Petri plates using cameras or scanners is a common practice in biology experiments that require phenotyping of plant seedlings or microbial colonies. However, manual imaging necessitates removing the plates from the growth conditions and increases the risk of introducing unwanted variables into the experiment, e.g., changes in temperature, humidity, illumination, vector of gravity, and mechanical stress. Such fluctuations, especially if triggered repeatedly by time-lapse imaging, might significantly impact phenotypes of interest, including plant seed germination, root growth, or microbial colony growth (De Ligne et al., 2019; Quint et al., 2016; Topham et al., 2017). Furthermore, manual imaging has limited time resolution, causes inconsistencies in the time of imaging, and impedes data acquisition during nights.

Automating labor- and time-intensive procedures is crucial to improving research quality and throughput, and open science hardware and software (tools which are freely available and modifiable) can help further this goal (Maia Chagas, 2018; Pearce, 2016, 2020). Moreover, opensource hardware improves the transparency and reproducibility of science while delivering radical cost savings (Pearce, 2020), enabling less well-funded labs (including those in low-income countries) to afford high-quality equipment (Maia Chagas, 2018; Wenzel, 2023).

A plethora of commercial and custom-made automated systems for imaging biological samples on Petri plates are already available (Fernandez et al., 2022; Lube et al., 2022; Nagel et al., 2020; Satbhai et al., 2017; Slovak et al., 2014a; Subramanian et al., 2013; Tovar et al., 2018; Yazdanbakhsh & Fisahn, 2009). However, we struggled to find an affordable platform that would be suitable for imaging Petri plates in standard plant growth incubators. Performing automated imaging under the growth conditions used for other experiments is crucial for direct comparison of the results, therefore we endeavored to develop a custom-made small footprint solution.

The result of our efforts is SPIRO, the compact **S**mart **P**late **I**maging **Ro**bot for timelapse imaging of vertical Petri plates, which fits into standard plant/microbe incubators. SPIRO was designed for biologists by biologists, introducing end-user insight into its development. We ensured that no prior knowledge of mechanical engineering, electronics, or computer science is necessary for its assembly and operation. SPIRO comprises the absolute minimum of components that warrants robust and reliable high-throughput time-lapse imaging and applicability for a broad range of experimental layouts. Owing to its minimalistic design, building the robot costs less than €300 (as of 2023), and it is not only easy to assemble but also to maintain, making it optimal for everyday use.

To further promote SPIRO’s applicability, we have developed two designated assays for high-throughput analysis of images produced by the robot: SPIRO Seed Germination and SPIRO Root Growth Assays. The assays are designed for analysis of phenotypic traits commonly used in plant biology: seed size, germination time, primary root length and growth rate. SPIRO assays encompass complete start-to-finish procedures comprising the preparation of Petri plates, automated imaging under user-defined conditions, semi-automated image processing, and statistical analysis of the quantitative data. The proof-of-concept experiments carried out for this article illustrate the benefits of using SPIRO.

SPIRO is powered by the open-source computer platform Raspberry Pi^1^ and comprises mostly 3D-printed hardware components, making it particularly suitable for customization. For the benefit of the scientific community, we are publishing SPIRO as an open-source project with all information about its structural design, electronics, software, and designated assays available under permissive licenses allowing unlimited use and modifications in the presence of correct attribution.

## Results

### General description/overview

SPIRO takes 8 megapixel (MP) timelapse images of up to four Petri plates positioned on a rotating cube-shaped stage. It is equipped with green LEDs for illuminating plates while imaging in the dark, and is controlled via a web-based user interface (**Fig. 1A-D, Movie S1**). The latter feature enables setting up imaging conditions remotely via Ethernet, Wi-Fi, or the built-in Wi-Fi hotspot, while the robot is inside a growth cabinet. SPIRO’s dimensions are approximately 50 cm × 30 cm × 30 cm (length × width × height), it weighs less than 3 kg, and can easily be transported (**Fig. 1A**).

**Figure 1.**
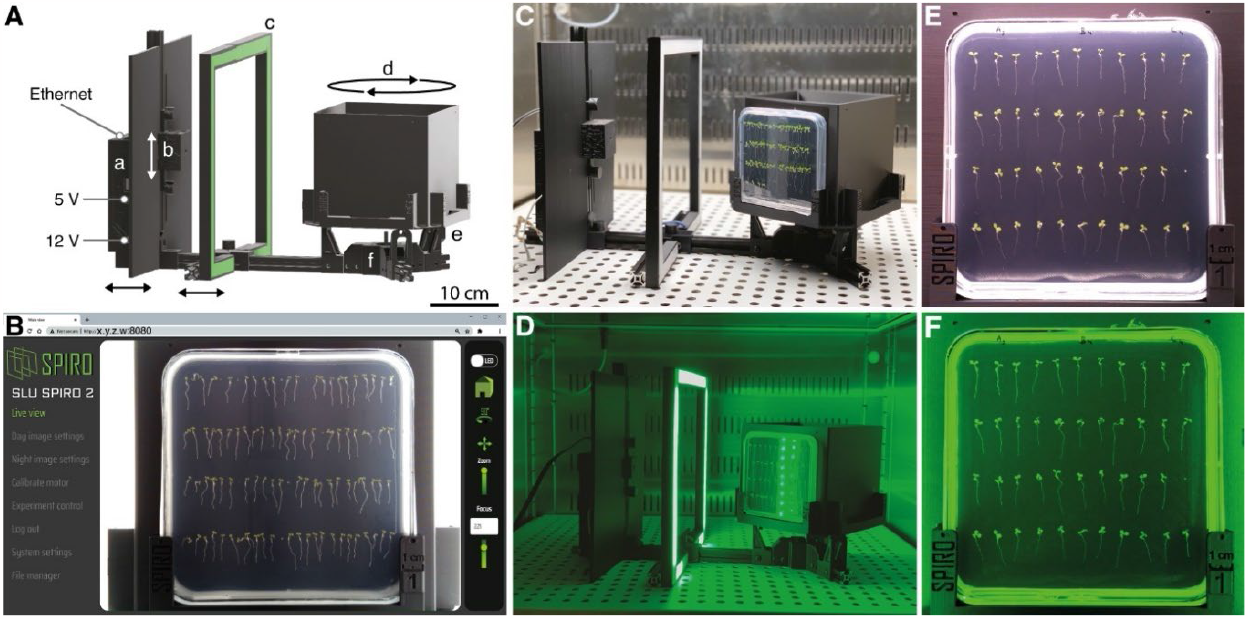
SPIRO, the Smart Plate Imaging Robot. **A**. 3D rendering image of SPIRO. The robot is controlled by a Raspberry Pi computer placed within the electronics housing (a). The camera house is mounted on a vertical axis with an anti-reflection screen (b). The white arrow indicates the possibility of adjusting the camera position along the vertical axis, the black arrows indicate the possibility to tune the distance between the camera and the stage. A green LED frame (c) provides illumination during imaging under night conditions. The cube-shaped stage can accommodate up to four Petri plates (d). At the beginning of each imaging cycle, the home position of the stage (where the first plate holder is facing the camera) is verified with the help of the positional sensor (e). The stage is rotated by a stepper motor (f). **B**. SPIRO is controlled by the designated software via a web-based graphical user interface, which allows users to adjust the settings for an experiment and to access imaging data. **C**. SPIRO in a plant growth cabinet under day conditions. **D**. SPIRO in a plant growth cabinet under night conditions. SPIRO automatically detects insufficient illumination and turns on the LED for a fraction of a second while the image is taken. **E, F**. Examples of images acquired by SPIRO under day and night conditions, respectively.

SPIRO performs imaging in cycles, where one cycle comprises: (i) finding the “home” position, at which the first plate holder is facing the camera; (ii) measuring the average light intensity to determine if it is sufficient for image acquisition without green LED illumination, or otherwise switching the LEDs on; (iii) taking an image and saving the file; (iv) rotating the stage by 90° to place the next plate holder in front of the camera, and repeating steps ii and iii until all plates are imaged (**Movie S1**). The duration of each cycle is less than two minutes, enabling high temporal resolution time-lapse imaging.

### Hardware design

We formulated the following three primary goals of the SPIRO hardware design: it should be affordable, customizable, and that a person with no previous experience could build it easily. For this reason, we opted to use 3D-printed parts and standard aluminium profiles for the structural components (**Figure 1, Tables S1 and S2, Files S1** and **S2**, and the SPIRO Hardware Repository^2^, and relatively cheap and readily available electronic components. 3D printing is inexpensive, allows for reproducible fabrication, rapid prototyping and modification, and is easily accessible. For example, printing can be done at publicly available facilities such as Makerspaces^3^ or ordered from online services (3dhubs^4^ or similar). Printed parts can be easily replicated if they get broken or customized for a specific application. We successfully printed and tested six SPIRO prototypes in four independent laboratories using black matte PLA filament (for detailed information about printing, see **Table S2** and the SPIRO Hardware Repository. The hardware proved to be easy to reproduce, robust and durable.

To facilitate use of SPIRO for a broad range of experiments, we designed plate holders compatible with the most commonly used plate formats: a 12 cm square (Greiner Bio-One International, Item: 688102) and a 9 cm round Petri plate (Sarstedt, 82.1473.001), and enabled adjusting the distance between the camera and the plate holders by moving the camera along the vertical and horizontal aluminum profiles (**Fig. 1A**).

#### Camera

SPIRO is equipped with a single 8 MP (3280×2464 pixels) color camera, saving images as RGB PNG files. Image files are stored in a user-defined experiment folder and are automatically sorted into four sub-folders corresponding to each plate holder. Metadata useful for further analysis is included in the file names, i.e., the names contain the plate holder number, date and time of acquisition, and information about day or night mode (for detailed information, please refer to **File S4**). SPIRO acquires excellent quality images regardless of ambient illumination conditions, which is crucial for downstream automated data analysis (**Fig. 1E and F, Movie S2**).

We provide assembly instructions for two possible configurations of SPIRO (GitHub SPIRO Hardware, **File S2**): the first is based on a Raspberry Pi v2 camera and requires manual adjustment of the focus before starting an experiment; the second one implements an Arducam camera module with motorized focus, which enables remote focusing via the SPIRO software. Both cameras are very compact and allow imaging of complete Petri plates from a short distance without fisheye distortion effects, a feature crucial for quantitative comparison of seed and seedling measurements. In our experience, both configurations deliver the same image quality, and while the first configuration is somewhat cheaper, the second one is more convenient. Notably, the first configuration can be relatively easily upgraded into the second. In our experience, current 8 MP cameras provide excellent image quality and manageable file size, which is especially important for analysis of long-term high-temporal-resolution experiments. As new cameras are being continuously developed, we strongly recommend checking the SPIRO Hardware repository for potential upgrades.

#### Computer

SPIRO’s electronics layout (**File S2**) is optimized to enable all essential features for robust high-throughput imaging while minimizing costs and complexity of assembly. SPIRO is powered by the cheap and readily available Raspberry Pi 3B+ single-board computer that controls the other four components: a camera, stepper motor, positional sensor (mini micro-switch) and LED light source. SPIRO’s software and acquired images are hosted on a microSD card mounted on the Raspberry Pi computer and can be remotely accessed via Ethernet or Wi-Fi connection. Notably, Raspberry Pi is an open-source computer platform designed for development of engineering solutions and is supported by a vivid community. Raspberry Pi is compatible with a multitude of electronics modules, sensors and components, often supplemented with suitable software packages^5^. It is thus optimal for further customizing the current SPIRO layout for specific uses.

#### Stepper motor and positional sensor

The cube-shaped stage of SPIRO is rotated by a stepper motor (i) during imaging, to position each of the four plate holders in front of the camera and (ii) between the imaging cycles, to ensure the plates are evenly exposed to the ambient conditions and that there is no shading effect from SPIRO’s light screen (**Fig. 1**).

The 12 V unipolar stepper motor we recommend provides sufficient force to reproducibly move the weight corresponding to the stage with 4 square Petri plates with agar growth medium and a holding torque that stably holds the stage position during imaging. Importantly, the motor movement is smooth and has no discernable impact on samples growth as was verified by monitoring Arabidopsis root growth under normal conditions (**Movie S2**).

The current layout of SPIRO requires two power supply units: a 5 V supply for the Raspberry Pi computer and a 12 V supply for the stepper motor and LED illuminator (**Fig. 1A, Table S1, File S2**). We decided for such configuration, as it drastically simplifies the assembly and maintenance of the robot, in comparison to implementing a single power supply unit.

At the beginning of each imaging cycle, the motor rotates the stage until the pin attached to the bottom of the stage presses the lever of a positional microswitch (**Fig. 1A, Movie S1)**. After the signal from the microswitch is detected, the stepper motor makes a predefined number of steps to place the first plate holder in front of the camera. If needed, the number of steps can be adjusted by the user in the “Calibrate motor” settings tab of the SPIRO software. After the image of the first plate is taken, the motor rotates the stage by 90° three more times, pausing to acquire images of the other three plate holders of SPIRO.

During prototyping, we considered implementation of either magnetic or infrared (IR) switches. However, a mechanical sensor provides the most robust system that is least susceptible to the presence of magnetic fields or stray light in the environment, and thus is applicable to a broader range of growth cabinets.

#### LEDs

SPIRO’s built-in light source enables imaging of Petri plates in the dark, providing another crucial benefit over manual imaging. The light source comprises a green LED strip mounted on a 3D-printed square frame with a diffuser (**Fig. 1A and D, Movie S1**). The LED frame can slide along the horizontal axis to finetune illumination of the Petri plates for individual conditions (**Fig. 1A**).

SPIRO does not require any instructions from the user about the day/night cycles in the growth cabinet. The intensity of ambient illumination is automatically assessed by SPIRO’s camera immediately prior to acquiring each image. If sufficient illumination is detected, SPIRO takes an image in the “day” mode (**Fig. 1C, E**), otherwise the robot turns on the LED light source and acquires a “night” image (**Fig. 1D, F**). ISO and shutter speed for image acquisition can be adjusted individually for day and night modes in the web-based interface of the SPIRO software (**Fig. 1B**).

During prototyping, we tested night imaging using IR LEDs and the IR-sensitive Raspberry Pi NoIR camera. However, this increased the cost of the robot, while significantly complicating its electronics layout and focusing procedure, and did not provide satisfactory quality of images suitable for automated image analysis. Typically, color camera detectors are most sensitive to green light, as they contain double the number of green pixels compared to red or blue. Hence, we speculated that using a green light source for illumination would be most efficient while using the color Raspberry Pi camera for imaging in the dark. Additionally, we took into consideration the use of SPIRO for plant imaging. Plants are known to be dependent on the light of blue and red wavelengths for photosynthesis and regulation of the circadian cycle(Eriksson & Millar, 2003; Wientjes et al., 2017). Although a number of studies have showed that green light wavelengths also have important regulatory effect during plant growth and development, the reported effects were observed after prolonged irradiation(Folta & Maruhnich, 2007). Thus, we speculated that illuminating Petri plates with green LEDs for only a fraction of a second during image acquisition should have the weakest impact on the growth and circadian cycle of imaged seedlings. To verify whether this was the case, we compared germination rates and root growth of *Arabidopsis thaliana* seeds and seedlings, respectively, imaged using two SPIRO systems, with and without green LEDs. Our analysis confirmed that green light indeed had no effect on germination and root growth (**Tables 1 and 2**).

**Table 1.** The results of an ANOVA test with one dependent variable (germinated before *t*=50 h), and three independent variables (genotype, system, LED), using four genotypes and 1154 seeds of *Arabidopsis thaliana*, indicate that SPIRO systems do not differ in performance and that LED illumination does not influence germination rates.

|  | *D.F.* | *Sum Sq* | *Mean Sq* | *F-value* | *p-value* |
| --- | --- | --- | --- | --- | --- |
| *Genotype* | 3 | 17.15 | 5.716 | 24.723 | $1.67 \times 10^{-15}$ |
| *System* | 1 | 0.09 | 0.094 | 0.407 | 0.524 |
| *LED* | 1 | 0.55 | 0.545 | 2.357 | 0.125 |
| *Residuals* | 1148 | 265.44 | 0.231 |  |  |

**Table 2.** Results of the comparison of Arabidopsis root growth models with and without effect of LED, using 316 seedlings on two different SPIRO systems. The likelihood ratio test revealed that the models are not significantly different, indicating that LED illumination has no effect on root growth

| *Model* | *Log likelihood* | *Deviance* | $\chi^2$ | $\chi^2$ *D.F.* | *p-value* |
| --- | --- | --- | --- | --- | --- |
| *Without LED* | 65440 | −130880 |  |  |  |
| *With LED* | 65281 | −130563 | 0 | 30 | 1.00 |

#### SPIRO Accessories

To aid the use of SPIRO we designed a set of 3D-printable accessories (**Table S2** and the SPIRO Hardware GitHub Repository): seed plating guides and anti-reflection lids.

Seed plating guides help the user to place Arabidopsis seeds on plates at regular distances from each other and from the edges of the plate. The first ensures optimal distance between individual seeds or seedlings for later recognition as distinct objects during image analysis. The latter is important to avoid overlaying seed and seedling images with reflections and shadows caused by the Petri plates’ rims.

Anti-reflection lids are designed to reduce reflections from seeds and seedlings that are usually visible in the Petri plates’ lids. Although such reflections might not be an issue during imaging, their presence is detrimental for automated image processing, as some of them are difficult to automatically differentiate from actual biological samples (for more information see **Fig. 2** in **File S4**).

**Figure 2.**
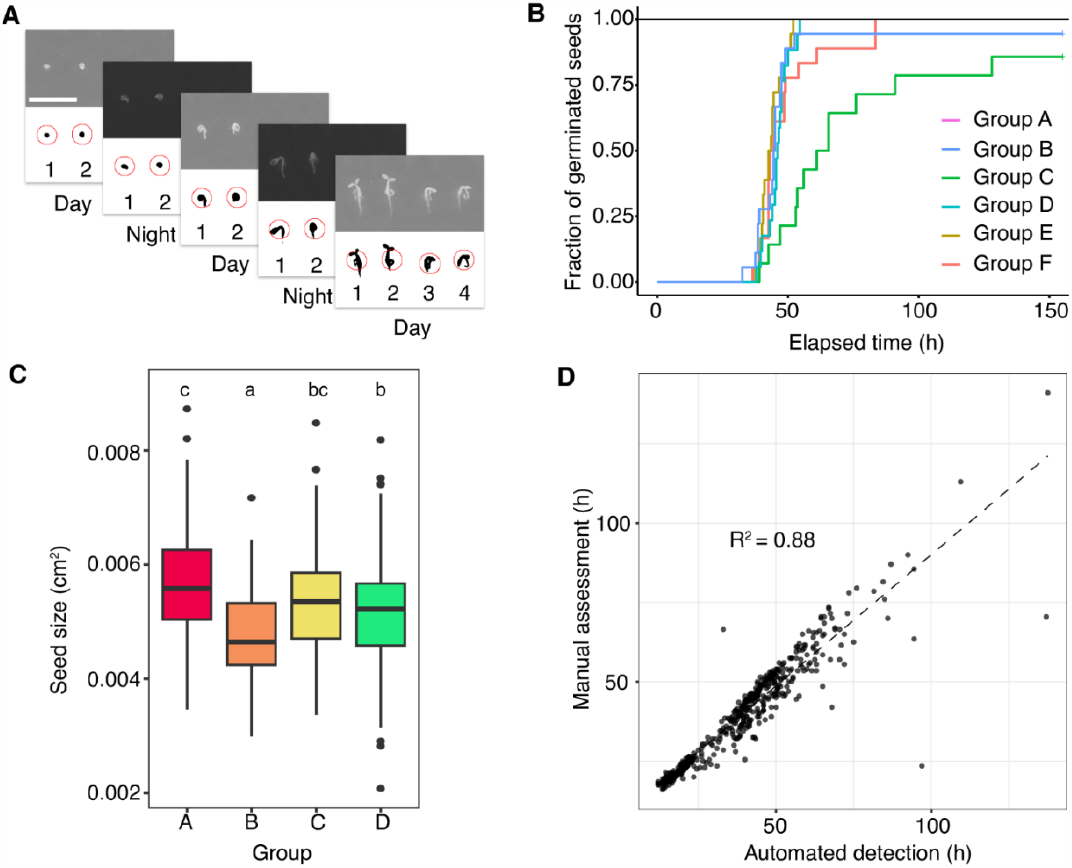
SPIRO Seed Germination Assay. **A**. The graphical output of the SPIRO Seed Germination Assay includes a time-lapse stack file, in which each frame contains the original photo and the result of its segmentation, i.e., a mask for each recognized seed annotated by a number and a circle. Scale bar, 500 μm. **B**. The assay provides Kaplan–Meier test results for germination of all groups of seeds included into analysis. Additional parameters for germination are calculated using the *germinationmetrics* package for R (Aravind et al., 2020) (for more information see SPIRO Assay manual, **File S3**). **C**. The assay also provides a t-test comparison of the mean seed size for the analyzed groups. Boxes represent interquartile range, the horizontal line denotes the median value, circles indicate outliers. Means of the groups that are not significantly different are annotated with the same small case letters, n = 401 seeds. **D**. Automatic detection of seed germination using SPIRO Seed Germination Assay provides results very similar to manual assessment of the germination (n=172).

### Assembling SPIRO

SPIRO was developed specifically to be buildable by a person with no expertise or training in engineering, electronics, or 3D printing. The complete list of components, step by step instructions, and tutorial videos for assembly are provided in **Tables S1** and **S2**, and **Files S1** and **S2**. However, we highly recommend checking the SPIRO Hardware Repository for potential updates before starting the assembly.

The SPIRO hardware includes a set of standard parts, like aluminium profiles, screws and electronic components that need to be purchased. The complete list of these components and links with suggestions on where to purchase them is provided in **Table S1** and the SPIRO Hardware Repository. Our experience of ordering hardware to build SPIRO prototypes in Germany and Sweden reproducibly showed that the most challenging part to acquire are the correct screws. At the moment, we cannot provide a plausible explanation for this peculiar phenomenon and will try to upgrade the SPIRO specifications to reduce the requirements for screws. Notably, approximately one quarter of purchase costs were covering shipping expenses. Furthermore, some parts, such as the LED strips, had to be ordered in excess. Thus, building several SPIROs lowers the price per robot.

The STL, 3MF and F3D files for 3D-printable parts of SPIRO and the printing settings recommendations are provided in **Table S2, File S2**, and the SPIRO Hardware Repository. SPIRO’s hardware was tested and optimized to be printed using PLA (polylactic acid) filament, which is the least expensive and sufficiently robust type of printable plastic. Printing in PETG (polyethylene terephthalate glycol) and ABS (acrylonitrile butadiene styrene) plastic is technically possible, but would require adjustment in scaling and printing settings, as the printed parts might shrink or warp significantly. Printing all SPIRO parts using one 3D printer takes about seven days. The prototyping was done using Prusa i3 MK2/S and MK3S printers. Nevertheless, the pre-arranged component sets we provide (**Table S2**, SPIRO Hardware Repository) can be printed on any type of 3D printer with a printing volume of 25x21x20 cm or more. In our experience, the assembly procedure can be completed by a determined untrained person in approximately two full-time workdays, while an experienced builder can assemble SPIRO in about four hours.

### Software and Installation

Since SPIRO was designed to be used within plant growth cabinets, we developed software that allows remotely controlling SPIRO via the internet. Besides convenience of use, remote control of the robot is essential to enable setting up imaging parameters under the conditions that will be used during the experiment. SPIRO’s software is used for adjusting the camera focus, setting up ISO (the camera’s sensitivity to light) and shutter speeds for day and night imaging conditions, defining the name and the duration of an experiment and the frequency of imaging, starting and terminating experiments and downloading imaging data (for detailed information please refer to **File S4** and the SPIRO Software Repository^6^.

SPIRO’s software has an intuitive web-based graphical user interface that can be easily accessed from any web browser (**Fig. 1B**). The layout of the software was optimized with the help of several beta-testers to ensure that the interface is sufficiently self-explanatory, does not require training prior to use and contains the complete minimal number of essential features. The program and detailed installation instructions are provided in **File S2** and the SPIRO Software Repository. The installation procedure requires SPIRO to be connected to the network, which can be done via the Ethernet port of the Raspberry Pi computer (**Fig. 1A**) or by connecting to a Wi-Fi network. For convenience, we recommend assigning SPIRO a DNS hostname or a static IP address, if this is possible within the user’s network. After installation is complete, it is possible to activate a Wi-Fi hotspot from SPIRO’s Raspberry Pi computer and use it for future connections to the robot (for instructions see **File S2** and the SPIRO Software Repository.

While setting up and starting a SPIRO experiment is done via an internet connection or Wi-Fi hotspot, running the experiment does not require the robot to be online. However, internet connectivity provides access to images during the experiment run and to the data from the previously completed experiments. The SPIRO software is open source, written in Python 3, and released under a 2-clause Berkeley Software Distribution (BSD) license, allowing re-distribution and modification of the source code as long as the original license and copyright notice are retained. SPIRO’s simple and versatile program for image acquisition includes features making automated imaging possible under conditions that might vary between experiments and laboratories:

- Imaging cycles are carried out at a user-defined frequency and duration. Before each imaging cycle, the stepper motor makes a pre-defined number of steps after the positional microswitch sensor was detected in order to place the stage in the “home” position.
- Image acquisition is preceded by assessment of illumination intensity. If the average pixel intensity on a sampled image is less than 10 (out of a maximum of 255), the software triggers acquisition under user-defined “night” settings and the LEDs are switched on for the duration of acquisition (less than one second). If the average pixel intensity on the sample image is higher than 10, the image is acquired with user-defined “day” settings.
- Full resolution RGB photos of each of the four plate holders are saved as PNG files on the microSD card mounted on the Raspberry Pi computer, accessible via the web-based user interface of the SPIRO software.
- While idling between imaging cycles, the stepper motor positions the stage at alternating 45° to ensure that all plates are evenly exposed to the conditions in the incubator.
- The experiment will be automatically terminated if there is no more space available for saving files on the microSD card. To prevent such truncations of experiments we introduced a feature into the software that provides the user with the estimated size of the experiment folder and the available disk space prior to the experiment start.

### Setting up an experiment

SPIRO was originally designed for imaging Arabidopsis seeds and seedlings. We provide detailed guidelines for casting agar plates, sterilizing and plating seeds, and adjusting imaging settings in **File S2**. For potential updates please refer to the SPIRO Assays Repository^7^.

### SPIRO assays

To thoroughly assess the data quality acquired using SPIRO and further enhance applicability of the imaging platform for the plant biology community, we developed complete pipelines for two commonly used phenotyping assays that greatly benefit from automated imaging: seed germination and root growth assays. The assays comprise image processing steps carried out by designated macro scripts in FIJI (Schindelin et al., 2012) (a distribution of ImageJ) and quantitative data processing steps carried out by custom R scripts. Step by step instructions and scripts are provided in **Files S4** and **S5**. Please note that updates are published in the SPIRO Assays Repository.

Each assay starts with pre-processing SPIRO raw data to combine individual images into time-lapse stack files with a set scale. The preprocessed data is then subjected to semi-automated image segmentation with identification of the objects of interest and measurement of their physical parameters, e.g., perimeter, length, and area, at successive time points. The quantitative data is further processed by R scripts to first ensure data quality and then apply custom-designed algorithms that determine seed size, germination time, root length and perform suitable statistical analyses.

The assays were designed to enable applicability for a broad range of experiment layouts and customization for specific uses, thus we introduced several user-guided steps that allow combining seeds or seedlings into groups of interests for analysis, trimming time ranges, renaming samples and removing outliers.

Each assay provides the user with graphical and quantitative outputs and suitable statistical analysis of the data. The assay design ensures quick verification of image segmentation accuracy and identification of potential outliers.

Furthermore, to make combining data from several experiments for the statistical analysis user-friendly, we developed the SPIRO Assay Customizer^8^. Customizer takes the quantitative data from one or several experiments cleaned by the quality control R scripts and provides the user with an intuitive graphical interface for merging experimental data, regrouping, renaming, or removing samples.

The stepwise design of each assay with outputs at intermediate stages allow the user to choose between relying on the provided algorithms or taking over at any point to perform their own analysis of the data.

#### SPIRO Seed Germination Assay

The germination process has been previously described as a two-phase process, where phase I corresponds to rapid water uptake, causing swelling of the seed and restarting of metabolic activity. Phase II is associated with protein synthesis and mitochondrial biogenesis, during which seed size remains constant (Bewley, 1997). The conclusion of germination and the commencement of the post-germination phase can be identified by the emergence of the embryonic root as it breaks through its surrounding tissues, a stage known as radicle emergence (Bewley, 1997). A later study suggested that an additional increase in seed size is associated with seed coat burst, which precedes Arabidopsis radicle emergence (Joosen et al., 2010). SPIRO Seed Germination Assay is based on the simple concept that the perimeter of a seed will steadily increase after germination, i.e., radicle emergence, has taken place. Hence, the image segmentation part of the assay detects individual seeds on the photos (**Fig. 2A, Movie S2**), and tracks changes in the perimeter of each seed within a user-defined time range. The data is then gathered into user-defined groups, e.g., genotypes or treatments, and subjected to a clean-up using a designated R script. During clean-up, objects too large or small to be a single Arabidopsis seed are removed from analysis. After this, for each seed the earliest time point of steady increase in perimeter is detected and identified as the time of germination. The range of the perimeter increase corresponding to the radicle emergence was manually estimated for Arabidopsis seeds and, if needed, can be adjusted to suit other plant species or to determine pre-germination changes of seed size. The assay is optimized for an imaging frequency of every 30 minutes and thus allows tracking minute differences in germination times. To take into account the effect of imbibition on the seed perimeter and also to compensate for the natural variation in Arabidopsis seed sizes, the germination algorithm normalizes perimeter changes for each seed by comparing it to the initial perimeter, which is determined as the average value for the first five images in the time-lapse data. For full details on the image analysis steps and germination detection algorithm, see **File S4**.

The significance of difference between mean germination times for the user-selected groups of seeds is then assessed by survival analysis, using the Kaplan–Meier test (**Fig. 2B**), evaluating differences using the log-rank test with false discovery rate multiple-testing correction. Other germination parameters that might be valuable for the user, such as rate-of-germination curve, time at maximum germination rate; time required to achieve 50% of total germination efficacy, time required for 50% of total seeds to germinate, are calculated by fitting a four-parameter Hill function to the data (El-Kassaby et al., 2008); for detailed information see **File S4** and the SPIRO Assays repository). Furthermore, the assay provides information about the size of individual seeds and the results of t-tests comparing seed sizes between user-defined groups (**Fig. 2C**). These data allow investigating correlations between seed size and germination parameters.

We verified the robustness of the semi-automated SPIRO Seed Germination Assay by comparing its results with germination time points detected manually on the same imaging data. The comparison of germination time points for 172 seeds revealed that the automated assay provides results similar to manual assessment (*R*^*2*^=0.88, **Fig. 2D**). For these experiments, sample preparation and imaging were done according to the instructions provided in **File S4**.

While developing the assay, we optimized the SPIRO hardware and protocol for seed plating and introduced seed plating guides that demark positions for placing seeds at optimal distance from each other and plate rims. As a result, when using four 12 cm square Petri plates, it is possible to detect germination for up to 2300 seeds in a single experiment. Additionally, we strongly recommend using SPIRO anti-reflection lids to reduce image segmentation artifacts caused by reflections (**Fig. 2** in **File S4**).

#### SPIRO Root Growth Assay

Quantifying primary root lengths of Arabidopsis seedlings is frequently used as a readout for physiological response to mutations or environmental stimuli (Lucas et al., 2011; Patterson et al., 2016; Satbhai et al., 2017). SPIRO is an excellent platform for seedling root phenotyping. We first tested processing of SPIRO images by existing automated image analysis tools for detection of single roots and root systems on time-lapse data (Betegón-Putze et al., 2019; Lobet et al., 2013; Satbhai et al., 2017). As these algorithms were optimized for a certain type of imaging data, their applicability for SPIRO-acquired images was limited.

Therefore, we developed the designated SPIRO Root Growth Assay, which uses SPIRO time-lapse data to track primary root length for individual seedlings starting from the germination time-point of the corresponding seed (**Fig. 3A, Movie S2**), builds a root growth rate model for user-defined groups of seedlings (**Fig. 3B**, described in detail in **File S4**), and then performs statistical analysis comparing root lengths and growth rates for the groups (**Fig. 3C**; complete details on the algorithmic and statistical procedures are given in **File S4** and the SPIRO Assays repository). Like the SPIRO Seed Germination Assay, the Root Growth Assay provides the user with a graphical output that shows the results of image segmentation for each user-selected group (**Fig. 3A**) and a quantitative output. The latter comprises (i) measurements of the segmented objects performed by the ImageJ macro; (ii) the measurements data cleaned up using a designated R script; (iii) germination time detected for each seed; (iv) charts plotting seedling’s root lengths *vs* absolute time or normalized to individual germination times; (v) charts displaying model fits for root growth rates of user-selected groups of seedlings (**Fig. 3B**); (vi) bar charts showing predicted root lengths for the groups of seedlings at 24-h intervals (**Fig. 3C**) and (vii) results of the statistical analysis comparing growth rates and root lengths between user-defined groups. For more details, please refer to the assay manual in **File S4** and the SPIRO Assays repository.

**Figure 3.**
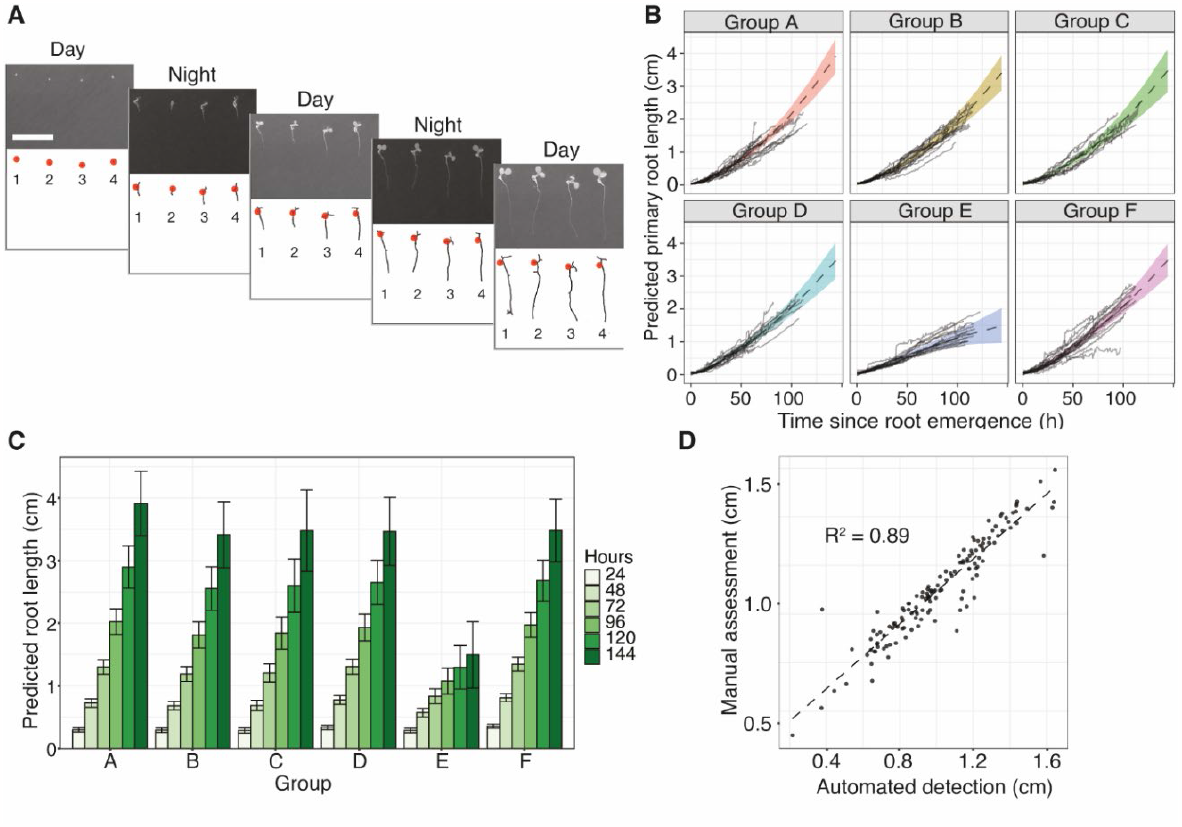
SPIRO Root Growth Assay. **A**. The graphical output of the SPIRO Root Growth Assay includes a time-lapse stack file, wherein each frame contains the original photo and the result of its segmentation, i.e., a skeletonized mask for each recognized seedling with denoted root start coordinate (red dot) annotated with a seedling number. Scale bar, 500 μm. **B**. The assay builds a model for root growth of each analyzed group (for more information see the SPIRO Assay manual, **File S4**). For each root, the elapsed time is calculated from the time of germination for the corresponding seed. Black solid lines indicate root lengths plotted vs time for each seedling, the dotted black line is the predicted root length, and the colored area indicates standard error. **C**. Based on the models shown in B, the assay also predicts average root length for each group at 24 h intervals. Error bars show standard error. **D**. Automatic detection of root length using SPIRO Root Growth Assay provides results very similar to manual assessment of the root lengths (n=141 seedlings). Manual assessment was done using the segmented line tool in ImageJ.

Comparison of manual and automated measurements of 141 Arabidopsis roots revealed that the SPIRO Root Growth Assay provides accuracy comparable with human performance (*R*^*2*^ *= 0*.*89*, **Fig. 3D**).

### Proof of concept experiments

To illustrate some of the advantages of SPIRO for plant biology we conducted two proof of concept experiments. First, we showcase the benefits of utilizing the SPIRO Root Growth Assay, which calculates root growth starting from the germination timepoint of each corresponding seed, rather than the seed plating time. This approach effectively corrects for potential variations in root lengths that may arise due to differences in germination. Second, we demonstrate the suitability of SPIRO for experimental layouts requiring uninterrupted imaging in the dark. By utilizing SPIRO, we were able to discover novel aspects of seed germination dependency on nutrient availability and light stimuli for genotypes deficient for autophagy or phytochromes.

#### SPIRO Root Growth Assay compensates for variations caused by differences in germination

Assessing the differences in root length between wild-type and mutant genotypes under various conditions is frequently used to evaluate the roles of specific genes in root development and growth enabling understanding their molecular and physiological functions (Lucas et al., 2011; Patterson et al., 2016; Satbhai et al., 2017). To ensure accurate interpretation of observed differences, it is important to account for possible confounding factors that might impact root length throughout the experiment. For example, if an analyzed genotype exhibits retarded or accelerated seed germination. This discrepancy will lead to a corresponding delayed or earlier initiation of root growth, eventually resulting in root length variations that are not representative of root cell division or elongation rates.

Changes in seed germination speed can be caused by mutations corrupting signaling of plant hormones such as abscisic acid (ABA) and gibberellins (GAs) (Abley et al., 2021). Additionally, peroxisome-related mutations are known to impact seed longevity and germination (Carrera et al., 2007; Pan et al., 2019). Most importantly, even for experiments that do not focus on such genotypes, the age of the seed batch and its storage conditions have drastic impact on germination efficacy (Trusiak et al., 2023). Therefore, accounting for germination timepoints when evaluating root length data is crucial to ensure accurate and meaningful results. SPIRO Root Growth Assay processes time-lapse data to monitor changes for each seed throughout the whole experiment and can accurately detect germination timepoint for normalizing observed root length changes.

To demonstrate the effect of delayed germination on root length assays, we conducted an experiment using a two-year old batch of autophagy-deficient (autophagy-related gene 5 knockout, *atg5-1*, (Thompson et al., 2005)). Arabidopsis seeds that had been stored at room temperature and varying humidity. SPIRO was used to image the seeds plated on standard 0.5xMS every half hour for a duration of 7 days. Expectedly, the suboptimal storage conditions caused deterioration of seed quality and resulted in an asynchronous germination phenotype, causing high variation in the root lengths of the seedlings at later time points (**Fig. 4A**). The time-lapse data was then processed using SPIRO Root Growth Assay to obtain two types of root length values: raw and normalized to germination time points (**Fig. 4B-D**).

**Figure 4.**
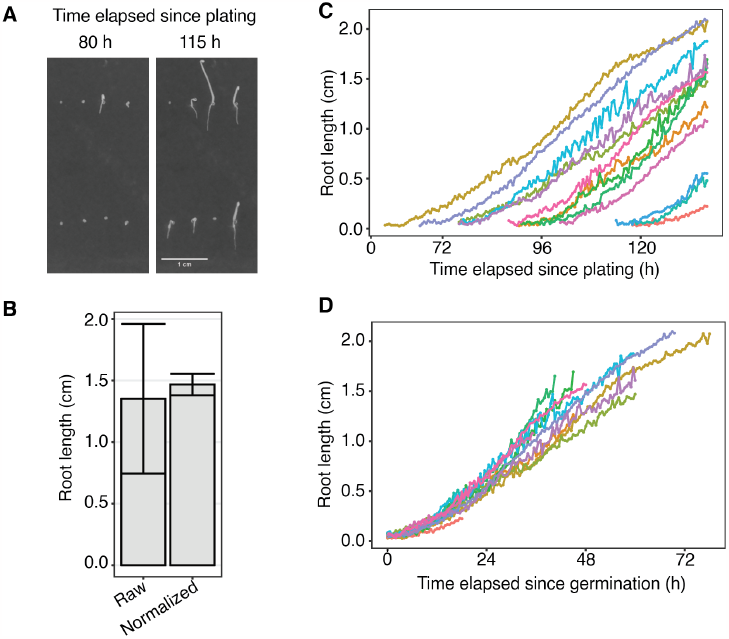
Root length normalization to the germination time point improves data accuracy. **A**. A cropped SPIRO photo illustrating unsynchronized seed germination (80 h since seed plating, left panel) resulting in varying root lengths detected later in the experiment (115 h since seed plating, right panel). A two-year-old batch of *atg5-1* seeds was germinated on the standard 0.5xMS plant agar medium. Plates with seeds mounted on SPIRO were incubated in the dark in a plant growth cabinet for a week and imaged every 30 min. Scale bar, 1 cm. **B**. Root lengths of the same seedlings measured with and without normalization to the corresponding germination times: “normalized” and “raw”, respectively. Normalized root lengths correspond to the values predicted by the SPIRO Root Growth Assay model for 46 h after germination. Raw root lengths correspond to the values detected by the SPIRO Root Growth macro on the last photo of the experiment. The error bars represent standard error for the normalized data and standard deviation for the raw data, n = 13. **C**. Raw root length values plotted for each analyzed seedling vs time elapsed since seed plating. **D**. Normalized root length values plotted for each analyzed seedling vs time elapsed since the corresponding seed germination detected by SPIRO Root Growth macro.

Illustratively, raw root length values measured at a time point elapsed since plating showed significant variation (**Fig. 4B, C**), whereas lengths of the same roots measured starting from the corresponding germination timepoints were consistent (**Fig. 4B, D**). In summary, this experiment exemplifies how using SPIRO and its dedicated Root Growth Assay allows obtaining accurate and biologically relevant information on Arabidopsis root growth.

#### Utilization of SPIRO reveals a novel role of autophagy in photoblastic response

Next, we explored possible uses of SPIRO for tracking seed germination in the dark. The green LED of SPIRO enables acquisition of high-quality images without impacting Arabidopsis root growth or germination (**Fig. 2, Table 1**).

Seed germination in Arabidopsis is an intricate process crucial for the organism’s survival and propagation. It is tightly regulated by internal programs and their interaction with external stimuli such as light and nutrients (Abley et al., 2021; Ibarra et al., 2013). For example, phytochromes A and B are the main light sensors regulating Arabidopsis seed germination (Shinomura et al., 1996), while nitrate and sucrose stimulate and repress germination, respectively (Vidal et al., 2014). One of the major pathways implicated in plant nutrient sensing and remobilization is autophagy, which plays a significant role in seed establishment (Chen et al., 2019; Erlichman et al., 2023). The exact roles autophagy plays in seed preparation for dormancy and germination are topics of actively ongoing investigation (Chen et al., 2019; Erlichman et al., 2023; Iglesias-Fernández & Vicente-Carbajosa, 2022). Here, we took advantage of SPIRO to simultaneously monitor potential effects of all three factors: autophagy, light, and macronutrients, on Arabidopsis seed germination.

To achieve this, we compared germination phenotypes of wild-type, autophagy-deficient (*atg4a/b, atg5, atg7*) and phytochrome-deficient (*phyA, phyB*) Arabidopsis seeds under 16 h illumination conditions (long day) and with turned off lamps (continuous night) in the presence or absence of macronutrients (**Fig. 5, Fig. S1, Movie S3**). All used seeds were of the Col-0 ecotype, seed batches were synchronized and ripened post-harvest prior to the experiment, therefore neither primary nor secondary dormancy were expected to be detectable (Chahtane et al., 2017; Stawska & Oracz, 2019). Indeed, under favourable control conditions all genotypes germinated synchronously (**Fig. 5A**, long day/control medium), with the anticipated exception of the *phyA* mutant. Remarkably, we observed a significant reduction in the ability of autophagy-deficient seeds to germinate in the absence of both nitrogen and illumination (**Fig. 5 A-C**), whereas the mean germination time of *ATG* knockouts was impacted much less than their germination rate (**Fig. 5C and D**). In contrast, depriving these seeds of either light or nitrogen alone did not result in such a pronounced effect on their ability to germinate. Importantly, additional removal of carbon from the medium did not significantly improve germination rate of these mutants (– N/–C, **Fig. 5**), indicating that not C:N ratio but rather only the presence of nitrogen is needed for normal germination rate of *ATG* knockouts under dark conditions (**Fig. 5**). These results reveal a novel potential role of autophagy in Arabidopsis photoblastic response and its crosstalk with nitrogen sensing.

**Figure 5.**
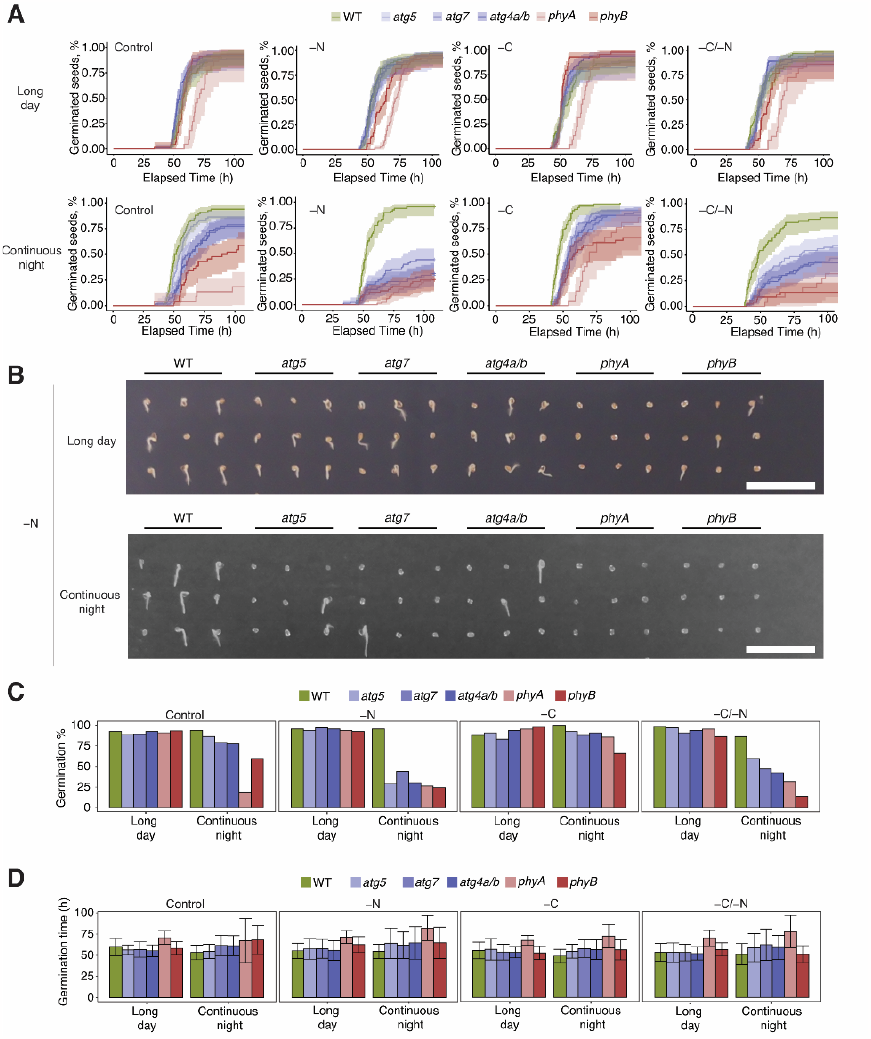
Utilization of SPIRO reveals a novel role of autophagy and nitrogen in seed germination. **A**. Kaplan–Meier plots produced by the SPIRO Seed Germination Assay showing germination dynamics of wild-type (WT), autophagy-deficient (*atg4a/b, atg5, atg7*) and phytochromes-deficient (*phyA, phyB*) Arabidopsis seeds. Seeds were plated on standard 0.5xMS medium (Control), 0.5xMS medium lacking nitrogen (–N) or sucrose (–C), or both (–N/–C). Plates with seeds were imaged by SPIROs inside plant growth cabinets set for 16 h light program (long day) or with turned off lights (continuous night). The charts represent a single experiment performed using two SPIROs. Total number of analyzed seeds = 2770, 2160 out of those germinated (1 seed was removed from the assay during QC check). **B**. A cropped SPIRO photo illustrating germination phenotypes on the –N medium quantified in (**A**). The photo corresponds to 56 h elapsed since plating time. Scale bars, 1 cm. **C**. Bar charts showing germination rate from the same experiment. Autophagy-deficient mutants show a strong decrease in the germination rate in the absence of nitrogen and light, while neither absence of nitrogen nor light alone has a discernible effect. **D**. Bar charts showing mean germination time from the same experiment demonstrate only small genotype-dependent differences compared to germination rate, indicating that breaking dormancy might be the most affected factor. Error bars indicate ± sd. Results of Tukey’s HSD test are shown in **Fig. S1**.

## Discussion

In this publication we thoroughly describe how to build and use the Smart Plate Imaging Robot, SPIRO. SPIRO provides excellent quality images, acquired under desired growth conditions at regular time intervals. It is an affordable, versatile, and robust imaging platform optimal for everyday use in plant biology and microbiology.

Open-source engineering solutions can have a great impact on improving the quality, reproducibility, and transparency of research (Pearce, 2020) but the benefit of their implementation is habitually undervalued. For instance, the number of working hours required for manual time-lapse imaging of Petri plates is often underestimated by calculating only the time needed for acquiring images. However, the daily chore of photographing or scanning plates interferes with other work tasks that must be fitted between imaging sessions and, depending on the desired imaging frequency, can become quite stressful. SPIRO not only frees up users’ time by taking over the imaging workload, but it also improves the quality of the acquired data and provides remote access to it from anywhere in the world. The latter feature turned out to be surprisingly useful during COVID-19 lockdown (Korbel & Stegle, 2020).

We strongly believe that reducing the barrier of entry is crucial to the adoption of open science hardware, especially in non-engineering disciplines. In our experience, one of the most significant bottlenecks in implementing engineering solutions custom-designed for laboratory use is the requirement of expertise in subjects that are not necessarily popular among biologists, such as mechanical engineering and electronics. SPIRO was developed specifically for biologists, and its design has at its core the concept of being simple and intuitive enough to be assembled and operated with no training in engineering, 3D printing, programming or using the SPIRO per se. Furthermore, we strived to develop an imaging platform that can be built anywhere in the world without requiring access to rare, specialized infrastructure and would be affordable also for research groups with limited funding. During preparation of this manuscript, numerous SPIROs have already been constructed in more than a dozen laboratories worldwide, confirming the need for such platform and validating our approach to making its construction feasible.

The current configuration of SPIRO is optimized for image acquisition under conditions commonly used in plant biology and microbiology. Moreover, the underlying Raspberry Pi platform is ideal for further expanding the system. A large variety of Raspberry Pi-compatible sensors and other input/output modules could be incorporated into derivative SPIRO systems to accommodate different research needs. For example, cheap sensors for temperature and humidity can be connected to unused general-purpose input/output (GPIO) pins on the computer board. Such modifications could be valuable when using SPIRO in a growth chamber that does not provide control or logging of these parameters.

We based our design on the use of 3D-printed structural components to further facilitate customization of SPIRO. We provide F3D model files that can be easily modified using Autodesk Fusion 360 software (**File S1**). As of 2023, Autodesk offers a free license tier for academic users, which includes training material. Furthermore, use of 3D-printed parts warrants reproducibility and robustness of the structural components, enables easy maintenance and repairs, and ensures that assembly can be done without access to rare, specialized infrastructure. For instance, not only have 3D printers become very affordable, with the cheapest models costing less than €300, but prints can also be ordered using commercial services or at public makerspace facilities.

We demonstrate analysis of high-throughput SPIRO-acquired data in two semi-automated assays for seed germination and seedling root growth. The detailed protocols for these assays comprise the complete procedure from preparing Petri plates to statistical analysis of the data. Both assays provide data analysis accuracy closely matching human performance (**Fig. 2D** and **3D**). Moreover, the SPIRO Seed Germination Assay enables very high temporal resolution enabling the detection of minute differences in germination times. Importantly, despite its small footprint, the platform still provides practical throughput: a single SPIRO can image up to 2300 Arabidopsis seeds in a single experiment for the Germination Assay, and 190 seedlings for the Root Growth Assay. In their current versions, the assays are implemented using ImageJ and R, software commonly used in biology labs, which should make their use and customization relatively uncomplicated. We have developed them as “plug-and-play” image analyses that are optimized for SPIRO data and provide outstanding results for the intended purpose.

Although the provided assays represent common use cases for SPIRO in plant biology, it did not escape our attention that SPIRO represents an ideal platform for novel assays including monitoring nyctinastic movements of Arabidopsis cotyledons (**Movie S4**), etiolation assays requiring night-imaged time-resolved data (**Movie S5**), root obstacle avoidance response, and other assays where high-temporal resolution data or normalization of phenotypic data to events such as germination or infection is required. SPIRO is also easily extended. As an example, RoPod chambers allow using SPIRO for normalizing microscopy data to other phenotypic data (Guichard et al., 2021). Moreover, the consistent and high quality of images acquired with SPIRO may reduce training requirements for machine learning approaches (Dobos et al., 2019; Gaggion et al., 2020; Yasrab et al., 2019), which might eventually enable more advanced analyses of SPIRO-acquired data such as detangling of crossing roots, time-resolved chlorosis assays, and measurement of lateral roots, hypocotyls and cotyledons under day and night conditions.

In the proof of concept experiments we illustrate the benefit of SPIRO Root Growth Assay that normalizes root length data to the corresponding seed germination, thus providing more accurate and biologically meaningful measurements than an endpoint imaging experiment. Furthermore, by tracking germination dynamics of autophagy-deficient mutants under different conditions, we unveiled a fascinating crosstalk between plant autophagy and responses to light and nitrogen during seed germination. Further in-depth investigations will be necessary to delve into this newly discovered branch of an already complex system that regulates Arabidopsis seed dormancy and germination.

It is worth noting for the sake of reproducibility for the future studies on the topic: although Arabidopsis seeds are photoblastic (Oh et al., 2009), i.e., require light stimuli for germination, we did not observe the expected decrease in germination of the wild-type seeds under continuous night conditions (**Fig. 5, Movie S3**). It could be that powering down the lamps did not completely deplete the light in the cabinet but set it below sensitivity of our luminometer. Indeed, later experiments under the same conditions but using extremely long exposure and a super-sensitive camera for bioluminescence detection revealed leakage of a small amount of light through the ventilation holes on the bottom of the cabinet. Thus, it is plausible that “continuous night” conditions are better described as Very Low Light Fluency conditions (Yanovsky et al., 1997). This could also explain the phenotype of *phyA* knockout showing higher sensitivity to the darkness, as PHYA can be activated by low light intensity triggering Very Low Fluency Response (VLFR), while PHYB requires higher intensity of red light (Ibarra et al., 2013; Oh et al., 2009; Yanovsky et al., 1997).

Downstream action of phytochromes at least partially relies on the activity of the nuclear ubiquitin-proteasome system (UPS) (Shen et al., 2005). This signaling might be less efficient if nuclear 26S proteasome would be exported into the cytoplasm for further degradation, which has been observed previously under nitrogen starvation conditions (Marshall et al., 2016). It is conceivable that autophagy-deficient seeds have a reduced internal nitrogen content, caused by inadequate remobilization of the macronutrient during seed establishment (Chen et al., 2019). Consequently, in the absence of an external nitrogen source, these seeds may indeed experience conditions resembling starvation, leading to a decrease in their UPS activity and subsequently affecting the functionality of phytochromes. This scenario would provide an explanation for the heightened dependence of autophagy-deficient mutants on light for germination and their potential transition into a secondary dormancy state when exposed to conditions characterized by limited nitrogen and low light availability. Further work is required to comprehend the mechanism underpinning this interesting phenomenon. In summary, our findings made in the proof-of-concept experiments highlight the power of using time-resolved SPIRO data to reveal dynamic and context-dependent phenotypes that might be overlooked in traditional endpoint imaging approaches.

The demand for affordable automated platforms for Petri plate imaging is clearly illustrated by the number of publications describing various systems (Ding et al., 2015; Fernandez et al., 2022; Gaggion et al., 2020; Lube et al., 2022; Nagel et al., 2020; Slovak et al., 2014b; Yasrab et al., 2019; Yazdanbakhsh & Fisahn, 2009). We have synthesized the key advantages that we identified in the previously published systems and ensured that SPIRO is affordable, compatible with standard growth cabinets, provides high-quality data and comes with instructions comprehensible for a person not trained in engineering. Our aim was to further facilitate universal access to automated high-quality imaging for all research groups regardless of their level of training or funding resources and enable easy integration of the automated approach into ongoing research.

For the sake of posterity, the current version of the complete information about SPIRO’s construction and use is provided in the supplementary information of this publication. However, SPIRO is under active development, and all updates are made available on the designated GitHub repositories (see Data availability section). We release all components of SPIRO as open-source material under licenses allowing unlimited use and modifications as long as proper attribution is present. The assays provided here are only two examples of the broad variety of possible SPIRO applications. The outstanding quality and fidelity of SPIRO’s images forms an excellent basis for any Petri plate-based imaging assay. We hope that SPIRO will help alleviate some pains of routine lab work and will also become a steppingstone for advancement of users’ interests in developing further solutions. We encourage users to further customize the platform, develop image-analysis pipelines suited for their own research and share optimizations with the scientific community.

## Acknowledgments

We would like to express our deepest gratitude to: Sebastian Mai (Heidelberg University) who provided valuable advice, resistors and terminal ports for SPIRO hardware development; all members of Prof. Karin Schumacher’s research group (Heidelberg University, Germany) for their patience and moral support and especially to Rainer Waadt for the name SPIRO; August Bergh, Thomas Danielsson, and Ping Wu (Ångström laboratory, Uppsala University, Sweden) for their contribution to the development of the early prototype; and to Johan Roxendal (Gothenburg University) and Dan Rosén (Uppsala University) for their contributions to the SPIRO software.

This study was supported by the funding from EU Horizon 2020 MSCA IF (799433 to EA Minina), Carl Tryggers Foundation (CTS 14 326 and 20:287 to EA Minina), The Swedish Research Council Formas (2021-01812 to EA Minina; 2019-01565 to AN Dauphinee; 2017-00541 to PV Bozhkov), DFG (CRC1101, TPA02 to K Schumacher), Formas (2019-01565 to AN Dauphinee), Formas (2016-20031 to M Sandgren), Knut and Alice Wallenberg Foundation (2018.0026 to PV Bozhkov), the Swedish Research Council VR (621-2013-4707 to PV Bozhkov), the Swedish Foundation for Strategic Research (Oil Crops for the Future to PV Bozhkov), and Crops for the Future Research Programme at the Swedish University of Agricultural Sciences.

## Author Contributions

JAO: concept, significant contribution to hardware development, main contribution to software development, significant contribution to assays development, experiments analysis, beta-testing the prototypes, manuscript preparation. JXL: main contribution to assays development, beta testing the final prototype, manuscript preparation. PHE: concept, beta-testing the prototypes and manuals, manuscript preparation. FB: beta testing the final prototype and manuals, contribution to the proof-of-concept experiments. SH: contribution to the proof-of-concept experiments. AND: beta testing the final prototype and manuals, figure preparation. ML, GH, JJ: contribution to hardware development, SB: contribution to root growth analysis. MS, KS, PVB: manuscript preparation. EAM: concept, main contribution to hardware development, significant contribution to software development, significant contribution to assays development, design, performance and analysis of the experiments, main contribution to writing the manuscript and manuals, project supervision.

## Data availability

The latest information about SPIRO is available on GitHub repositories dedicated to hardware (https://github.com/AlyonaMinina/SPIRO.Hardware), software (https://github.com/jonasoh/spiro), semi-automated assays (https://github.com/jiaxuanleong/SPIRO.Assays) and SPIRO Assay Customizer (https://github.com/jonasoh/spiroassay-customizer). The latest updates of the manuals provided in the supplementary material of this article are also available on the corresponding repositories.

## List of supplementary material

| **Supplementary material** | **Description** |
| --- | --- |
| **Figure S1** | P-values for Tukey's HSD for mean seed germination time. |
| **Figure S2** | Implication of SPIRO reveals accelerated shoot etiolation in autophagy deficient seedlings with access to sucrose. |
| **Table S1** | List of components to be purchased and suggestions of vendors |
| **Table S2** | List of 3D-printable components and instructions for printing settings |
| **Supplementary File 1** | Printable (STL, 3MF), and editable (F3D) model files |
| **Supplementary File 2** | SPIRO Assembly Instructions and links to tutorial videos |
| **Supplementary File 3** | SPIRO Software GitHub repository archive file |
| **Supplementary File 4** | SPIRO Assays Manual |
| **Supplementary File 5** | SPIRO Assays GitHub repository archive file |
| **Supplementary File 6** | SPIRO Assay Customizer archive file |
| **Supplementary Figure 1** | *P*-values for Tukey's HSD test of seed germination times shown in the Figure 5 |
| **Supplementary Video 1** | SPIRO Overview. A video showing two SPIRO systems placed in a plant growth incubator and the time-lapse video recorded using them |
| **Supplementary Video 2** | Demonstration of the image quality acquired by SPIRO and the capabilities of the two companion SPIRO Assays. The movie shows raw and preprocessed data, as well as illustrates the output of the Germination and Root Growth Assays. |
| **Supplementary Video 3** | Germination of *Arabidopsis thaliana* seeds under long day and continuous night conditions on the nitrogen depleted medium. |
| **Supplementary Video 4** | Nyctinastic movements of *Arabidopsis thaliana* cotyledons detected using SPIRO. |
| **Supplementary Video 5** | Etiolation of wild-type (WT) and autophagy-deficient (*atg4a/b*, *atg5*) *Arabidopsis thaliana* seedlings. |

## Materials and Methods

### Plant material

All experiments were performed using *Arabidopsis thaliana* Col-0 ecotype. All autophagy-deficient lines were described previously: *atg4a/b, atg4a-1/atg4b-1*(Chung et al., 2010); *atg5* knockout, *atg5-1* (Thompson et al., 2005), *atg7* knockout, *atg7-2* (Hofius et al., 2009), *nbr1* (Zhou et al., 2013). Seeds of *phyA-211* (Nagatani et al., 1993) and *phyB-9* (Reed et al., 1993; Yoshida et al., 2018) mutants were kindly provided by Dr. Maria Eriksson (Umeå University). Seed batches were synchronized by harvesting them from the plants grown simultaneously under long day conditions: 150 μE m^-2^ s^-1^ light for 16 h, 8 h dark, 22°C. Synchronized seed batches were subjected to post-harvest ripening by storing them at 22°C in the dark, 30– 50% humidity for 13 months prior to the experiment.

### Plant growth

Seed sterilization was performed using ethanol method and plated using SPIRO seed plating guides described in **File S4**.

Arabidopsis seedlings were grown in 12 cm square plates with 35 mL of standard 0.5xMS: 0.5xMurashige & Skoog medium including vitamins (M0222, Duchefa Biochemie, Haarlem, The Netherlands), 10mM MES (M1503, Duchefa Biochemie, Haarlem, The Netherlands), 1% sucrose, 0.8% Plant agar (P1001, Duchefa Biochemie, Haarlem, The Netherlands) pH5.8. –C medium composition was identical to 0.5xMS but lacking sucrose. – N medium was prepared using 0.5x MS basal salt micronutrient solution (M0529-1L, Sigma-Aldrich, Saint Louis, MO, USA) supplemented with 1.5mM CaCl_2_, 0.75 MgSO_4_, 0.625 KH_2_PO_4_ and 2.5 KCl, pH5.8.

Plates were closed with sterilized SPIRO lids and imaged while vertically mounted on SPIRO inside Sanyo plant growth cabinet set to long day conditions: 150 μE m^-2^ s^-1^ light for 16 h, 8 h dark, 22 °C. Continuous night conditions were created by turning off lamps in the growth cabinet.

### Assessing the influence of green light on germination and root growth

To determine whether the green light illumination used for imaging in the dark affected plant growth, germination and root growth rates were assessed in A. thaliana seeds and seedlings. This experiment also served to evaluate reproducibility of results between different SPIRO builds and cameras. Two separate SPIRO systems were used, one with an Arducam camera and one with a Raspberry Pi camera. In four separate experiments, replicate plates containing the same genotypes and media were imaged on both systems, with one system having its LEDs disabled, so that each system used its LEDs for two out of four experiments. Germination was evaluated using the SPIRO Germination Assay. Due to the constraints imposed on germination detection by the lack of night images, the germination rate at t = 50 h was used. Results were evaluated using ANOVA with genotype, system, and LED as the independent variables.

For root growth, two SPIRO systems were used, one with and one without LEDs, using replicate genotypes and media. Root growth was measured using the SPIRO Root Growth Assay. Two models were fitted to the data: the first was the standard root growth model as described in the SPIRO Assay Manual (File S4), and the second was the same model but with the fixed effect of LED and all its interaction effects added. Models were then compared using the anova function in R.

### Image analysis

Unless stated otherwise image analysis was performed using dedicated SPIRO assays. Charts for figures were built using ggplot.

## Footnotes

1 https://www.raspberrypi.org/

2 https://Github.Com/AlyonaMinina/SPIRO.Hardware

3 https://www.Makerspaces.Com/What-Is-a-Mak-erspace

4 https://www.hubs.com/

5 https://tutorials-raspberrypi.com/Raspberry-Pi-Sensors-Overview-50-Important-Components/

6 https://github.com/Jonasoh/Spiro

7 https://github.com/jiaxuanleong/SPIRO.Assays

8 https://github.com/jonasoh/spiro-assay-customizer

## References

Abley, K., Formosa-Jordan, P., Tavares, H., Chan, E. Y. T., Afsharinafar, M., Leyser, O., & Locke, J. C. W. (2021). An aba-ga bistable switch can account for natural variation in the variability of arabidopsis seed germination time. ELife, 10. 10.7554/eLife.59485

Aravind, J., Vimala Devi, S., Radhamani, J., Jacob, S. R., & Kalyani, S. (2020). Germinationmetrics: Seed Germination Indices and Curve Fitting. R package version 0.1.3.9000.

Betegón-Putze, I., González, A., Sevillano, X., Blasco-Escámez, D., & Caño-Delgado, A. I. (2019). MyROOT: a method and software for the semiautomatic measurement of primary root length in Arabidopsis seedlings. Plant Journal, 98(6), 1145–1156. 10.1111/tpj.14297

Bewley, J. D. (1997). Seed Germination and Dormancy. The Plant Cell, 44(8), 1055–1066. 10.1105/tpc.9.7.1055

Carrera, E., Holman, T., Medhurst, A., Peer, W., Schmuths, H., Footitt, S., Theodoulou, F. L., & Holdsworth, M. J. (2007). Gene expression profiling reveals defined functions of the ATP-binding cassette transporter COMATOSE late in phase II of germination. Plant Physiology, 143(4). 10.1104/pp.107.096057

Chahtane, H., Kim, W., & Lopez-Molina, L. (2017). Primary seed dormancy: A temporally multilayered riddle waiting to be unlocked. In Journal of Experimental Botany (Vol. 68, Issue 4). 10.1093/jxb/erw377

Chen, Q., Shinozaki, D., Luo, J., Pottier, M., Havé, M., Marmagne, A., Reisdorf-Cren, M., Chardon, F., Thomine, S., Yoshimoto, K., & Masclaux-Daubresse, C. (2019). Autophagy and nutrients management in plants. In Cells (Vol. 8, Issue 11). 10.3390/cells8111426

Chung, T., Phillips, A. R., & Vierstra, R. D. (2010). ATG8 lipidation and ATG8-mediated autophagy in Arabidopsis require ATG12 expressed from the differentially controlled ATG12A and ATG12B loci. Plant Journal, 62(3), 483–493. 10.1111/j.1365-313X.2010.04166.x

De Ligne, L., Vidal-Diez de Ulzurrun, G., Baetens, J. M., Van den Bulcke, J., Van Acker, J., & De Baets, B. (2019). Analysis of spatio-temporal fungal growth dynamics under different environmental conditions. IMA Fungus, 10(1), 1–13. 10.1186/s43008-019-0009-3

Ding, X., Vogel, M., Boschke, E., Bley, T., & Lenk, F. (2015). PetriJet Platform Technology: An Automated Platform for Culture Dish Handling and Monitoring of the Contents. Journal of Laboratory Automation, 20(4), 447–456. 10.1177/2211068215576191

Dobos, O., Horvath, P., Nagy, F., Danka, T., & Viczián, A. (2019). A deep learningbased approach for high-throughput hypocotyl phenotyping. Plant Physiology, 181(4), 1415–1424. 10.1104/pp.19.00728

El-Kassaby, Y. A., Moss, I., Kolotelo, D., & Stoehr, M. (2008). Seed germination: Mathematical representation and parameters extraction. Forest Science, 54(2), 220–227. 10.1093/forestscience/54.2.220

Eriksson, M. E., & Millar, A. J. (2003). The circadian clock. A plant’s best friend in a spinning world. Plant Physiology, 132(2), 732–738. 10.1104/pp.103.022343

Erlichman, O. A., Weiss, S., Abu Arkia, M., Ankary-Khaner, M., Soroka, Y., Jasinska, W., Rosental, L., Brotman, Y., & Avin-Wittenberg, T. (2023). Autophagy in maternal tissues contributes to Arabidopsis seed development. Plant Physiology. 10.1093/plphys/kiad350

Fernandez, R., Crabos, A., Maillard, M., Nacry, P., & Pradal, C. (2022). Highthroughput and automatic structural and developmental root phenotyping on Arabidopsis seedlings. Plant Methods, 18(1). 10.1186/s13007-022-00960-5

Folta, K. M., & Maruhnich, S. A. (2007). Green light: A signal to slow down or stop. Journal of Experimental Botany, 58(12), 3099–3111. 10.1093/jxb/erm130

Gaggion, N., Ariel, F., Daric, V., Lambert, É., Legendre, S., Roulé, T., Camoirano, A., Milone, D. H., Crespi, M., Blein, T., & Ferrante, E. (2020). ChronoRoot: Highthroughput phenotyping by deep segmentation networks reveals novel temporal parameters of plant root system architecture. BioRxiv. 10.1101/2020.10.27.350553

Guichard, M., Holla, S., Wernerová, D., Grossmann, G., & Minina, E. A. (2021). RoPod, a customizable toolkit for noninvasive root imaging, reveals cell type-specific dynamics of plant autophagy. 1–21.

Hofius, D., Schultz-Larsen, T., Joensen, J., Tsitsigiannis, D. I., Petersen, N. H. T., Mattsson, O., Jørgensen, L. B., Jones, J. D. G., Mundy, J., & Petersen, M. (2009). Autophagic Components Contribute to Hypersensitive Cell Death in Arabidopsis. Cell, 137(4), 773–783. 10.1016/j.cell.2009.02.036

https://800ezmicro.com/equipment/anaerobicmicroaerobic-systems/32-petrifoto-plateimaging-system.html. (n.d.).

Ibarra, S. E., Auge, G., Sánchez, R. A., & Botto, J. F. (2013). Transcriptional programs related to phytochrome a function in arabidopsis seed germination. Molecular Plant, 6(4). 10.1093/mp/sst001

Iglesias-Fernández, R., & Vicente-Carbajosa, J. (2022). A View into Seed Autophagy: From Development to Environmental Responses. Plants, 11(23). 10.3390/plants11233247

Joosen, R. V. L., Kodde, J., Willems, L. A. J., Ligterink, W., van der Plas, L. H. W., & Hilhorst, H. W. M. (2010). Germinator: a software package for high-throughput scoring and curve fitting of Arabidopsis seed germination. The Plant Journal, 62(1), 148–159. 10.1111/j.1365-313X.2009.04116.x

Korbel, J. O., & Stegle, O. (2020). Effects of the COVID-19 pandemic on life scientists. In Genome Biology (Vol. 21, Issue 1). 10.1186/s13059-020-02031-1

Lobet, G., Draye, X., & Périlleux, C. (2013). An online database for plant image analysis software tools. Plant Methods, 9(1). 10.1186/1746-4811-9-38

Lube, V., Noyan, M. A., Przybysz, A., Salama, K., & Blilou, I. (2022). MultipleXLab: A high-throughput portable live-imaging root phenotyping platform using deep learning and computer vision. Plant Methods, 18(1), 38. 10.1186/s13007-02200864-4

Lucas, M., Swarup, R., Paponov, I. A., Swarup, K., Casimiro, I., Lake, D., Peret, B., Zappala, S., Mairhofer, S., Whitworth, M., Wang, J., Ljung, K., Marchant, A., Sandberg, G., Holdsworth, M. J., Palme, K., Pridmore, T., Mooney, S., & Bennett, M. J. (2011). SHORT-ROOT regulates primary, lateral, and adventitious root development in Arabidopsis. Plant Physiology, 155(1), 384–398. 10.1104/pp.110.165126

Maia Chagas, A. (2018). Haves and have nots must find a better way: The case for open scientific hardware. PLoS Biology, 16(9), e3000014. 10.1371/jour-nal.pbio.3000014

Marshall, R. S., McLoughlin, F., & Vierstra, R. D. (2016). Autophagic Turnover of Inactive 26S Proteasomes in Yeast Is Directed by the Ubiquitin Receptor Cue5 and the Hsp42 Chaperone. Cell Reports, 16(6). 10.1016/j.celrep.2016.07.015

Nagatani, A., Reed, J. W., & Chory, J. (1993). Isolation and initial characterization of Arabidopsis mutants that are deficient in phytochrome A. Plant Physiology, 102(1). 10.1104/pp.102.1.269

Nagel, K. A., Lenz, H., Kastenholz, B., Gilmer, F., Averesch, A., Putz, A., Heinz, K., Fischbach, A., Scharr, H., Fiorani, F., Walter, A., & Schurr, U. (2020). The platform GrowScreen-Agar enables identification of phenotypic diversity in root and shoot growth traits of agar grown plants. Plant Methods, 16(1), 1–17. 10.1186/s13007-020-00631-3

Oh, E., Kang, H., Yamaguchi, S., Park, J., Lee, D., Kamiya, Y., & Choi, G. (2009). Genome-wide analysis of genes targeted by PHYTOCHROME INTERACTING FACTOR 3-LIKE5 during seed germination in arabidopsis. Plant Cell, 21(2). 10.1105/tpc.108.064691

Pan, R., Liu, J., & Hu, J. (2019). Peroxisomes in plant reproduction and seed-related development. In Journal of Integrative Plant Biology (Vol. 61, Issue 7). 10.1111/jipb.12765

Patterson, K., Walters, L. A., Cooper, A. M., Olvera, J. G., Rosas, M. A., Rasmusson, A. G., & Escobar, M. A. (2016). Nitrate-regulated glutaredoxins control arabidopsis primary root growth. Plant Physiology, 170(2), 989–999. 10.1104/pp.15.01776

Pearce, J. M. (2016). Return on investment for open source scientific hardware development. Science and Public Policy, 43(2), 192–195. 10.1093/scipol/scv034

Pearce, J. M. (2020). Economic savings for scientific free and open source technology: A review. HardwareX, 8, e00139. 10.1016/j.ohx.2020.e00139

Quint, M., Delker, C., Franklin, K. A., Wigge, P. A., Halliday, K. J., & Van Zanten, M. (2016). Molecular and genetic control of plant thermomorphogenesis. Nature Plants, 2(January), 1–9. 10.1038/nplants.2015.190

Reed, J. W., Nagpal, P., Poole, D. S., Furuya, M., & Chory, J. (1993). Mutations in the gene for the red/far-red light receptor phytochrome B alter cell elongation and physiological responses throughout arabidopsis development. Plant Cell, 5(2). 10.1105/tpc.5.2.147

Satbhai, S. B., Göschl, C., & Busch, W. (2017). Automated high-throughput root phenotyping of Arabidopsis thaliana under nutrient deficiency conditions. Methods in Molecular Biology, 1610, 135–153. 10.1007/978-1-4939-7003-2_10

Schindelin, J., Arganda-Carreras, I., Frise, E., Kaynig, V., Longair, M., Pietzsch, T., Preibisch, S., Rueden, C., Saalfeld, S., Schmid, B., Tinevez, J. Y., White, D. J., Hartenstein, V., Eliceiri, K., Tomancak, P., & Cardona, A. (2012). Fiji: An opensource platform for biological-image analysis. Nature Methods, 9(7), 676–682. 10.1038/nmeth.2019

Shen, H., Moon, J., & Huq, E. (2005). PIF1 is regulated by light-mediated degradation through the ubiquitin-26S proteasome pathway to optimize photomorphogenesis of seedlings in Arabidopsis. Plant Journal, 44(6). 10.1111/j.1365-313X.2005.02606.x

Shinomura, T., Nagatani, A., Hanzawa, H., Kubota, M., Watanabe, M., & Furuya, M. (1996). Action spectra for phytochrome A-and B-specific photoinduction of seed germination in Arabidopsis thaliana. Proceedings of the National Academy of Sciences of the United States of America, 93(15). 10.1073/pnas.93.15.8129

Slovak, R., Göschl, C., Su, X., Shimotani, K., Shiina, T., & Busch, W. (2014a). A scalable open-source pipeline for large-scale root phenotyping of Arabidopsis. Plant Cell, 26(6), 2390–2403. 10.1105/tpc.114.124032

Slovak, R., Göschl, C., Su, X., Shimotani, K., Shiina, T., & Busch, W. (2014b). A scalable open-source pipeline for large-scale root phenotyping of Arabidopsis. Plant Cell, 26(6), 2390–2403. 10.1105/tpc.114.124032

Stawska, M., & Oracz, K. (2019). Phyb and hy5 are involved in the blue light-mediated alleviation of dormancy of arabidopsis seeds possibly via the modulation of expression of genes related to light, ga, and aba. International Journal of Molecular Sciences, 20(23). 10.3390/ijms20235882

Subramanian, R., Spalding, E. P., & Ferrier, N. J. (2013). A high throughput robot system for machine vision based plant phenotype studies. Machine Vision and Applications, 24(3), 619–636. 10.1007/s00138-012-0434-4

Thompson, A. R., Doelling, J. H., Suttangkakul, A., & Vierstra, R. D. (2005). Autophagic nutrient recycling in Arabidopsis directed by the ATG8 and ATG12 conjugation pathways. Plant Physiology, 138(4). 10.1104/pp.105.060673

Topham, A. T., Taylor, R. E., Yan, D., Nambara, E., Johnston, I. G., & Bassel, G. W. (2017). Temperature variability is integrated by a spatially embedded decisionmaking center to break dormancy in Arabidopsis seeds. Proceedings of the National Academy of Sciences of the United States of America, 114(25), 6629–6634. 10.1073/pnas.1704745114

Tovar, J. C., Hoyer, J. S., Lin, A., Tielking, A., Callen, S. T., Elizabeth Castillo, S., Miller, M., Tessman, M., Fahlgren, N., Carrington, J. C., Nusinow, D. A., & Gehan, M. A. (2018). Raspberry Pi-powered imaging for plant phenotyping. Applications in Plant Sciences, 6(3), e1031. 10.1002/aps3.1031

Trusiak, M., Plitta-Michalak, B. P., & Michalak, M. (2023). Choosing the Right Path for the Successful Storage of Seeds. In Plants (Vol. 12, Issue 1). 10.3390/plants12010072

Vidal, E. A., Moyano, T. C., Canales, J., & Gutiérrez, R. A. (2014). Nitrogen control of developmental phase transitions in Arabidopsis thaliana. In Journal of Experimental Botany (Vol. 65, Issue 19, pp. 5611–5618). Oxford University Press. 10.1093/jxb/eru326

Wenzel, T. (2023). Open hardware: From DIY trend to global transformation in access to laboratory equipment. PLoS Biology, 21(1). 10.1371/jour-nal.pbio.3001931

Wientjes, E., Philippi, J., Borst, J. W., & van Amerongen, H. (2017). Imaging the Photosystem I/Photosystem II chlorophyll ratio inside the leaf. Biochimica et Biophysica Acta - Bioenergetics, 1858(3), 259–265. 10.1016/j.bbabio.2017.01.008

Yanovsky, M. J., Casal, J. J., & Luppi, J. P. (1997). The VLF loci, polymorphic between ecotypes Landsberg erecta and Columbia, dissect two branches of phytochrome A signal transduction that correspond to very-low-fluence and high-irradiance responses. The Plant Journal, 12(3). 10.1046/j.1365-313x.1997.00659.x

Yasrab, R., Atkinson, J. A., Wells, D. M., French, A. P., Pridmore, T. P., & Pound, M. P. (2019). RootNav 2.0: Deep learning for automatic navigation of complex plant root architectures. GigaScience, 8(11), 1–16. 10.1093/gigascience/giz123

Yazdanbakhsh, N., & Fisahn, J. (2009). High throughput phenotyping of root growth dynamics, lateral root formation, root architecture and root hair development enabled by PlaRoM. Functional Plant Biology, 36(11), 938–946. 10.1071/FP09167

Yoshida, Y., Sarmiento-Mañús, R., Yamori, W., Ponce, M. R., Micol, J. L., & Tsukaya, H. (2018). The arabidopsis phyB-9 mutant has a second-site mutation in the VENOSA4 gene that alters chloroplast size, photosynthetic traits, and leaf growth. In Plant Physiology (Vol. 178, Issue 1). 10.1104/pp.18.00764

Zhou, J., Wang, J., Cheng, Y., Chi, Y. J., Fan, B., Yu, J. Q., & Chen, Z. (2013). NBR1-Mediated Selective Autophagy Targets Insoluble Ubiquitinated Protein Aggregates in Plant Stress Responses. PLoS Genetics, 9(1). 10.1371/journal.pgen.1003196

