## Supplementary Figure 1 for "SPIRO – the automated Petri plate imaging platform designed by biologists, for biologists"

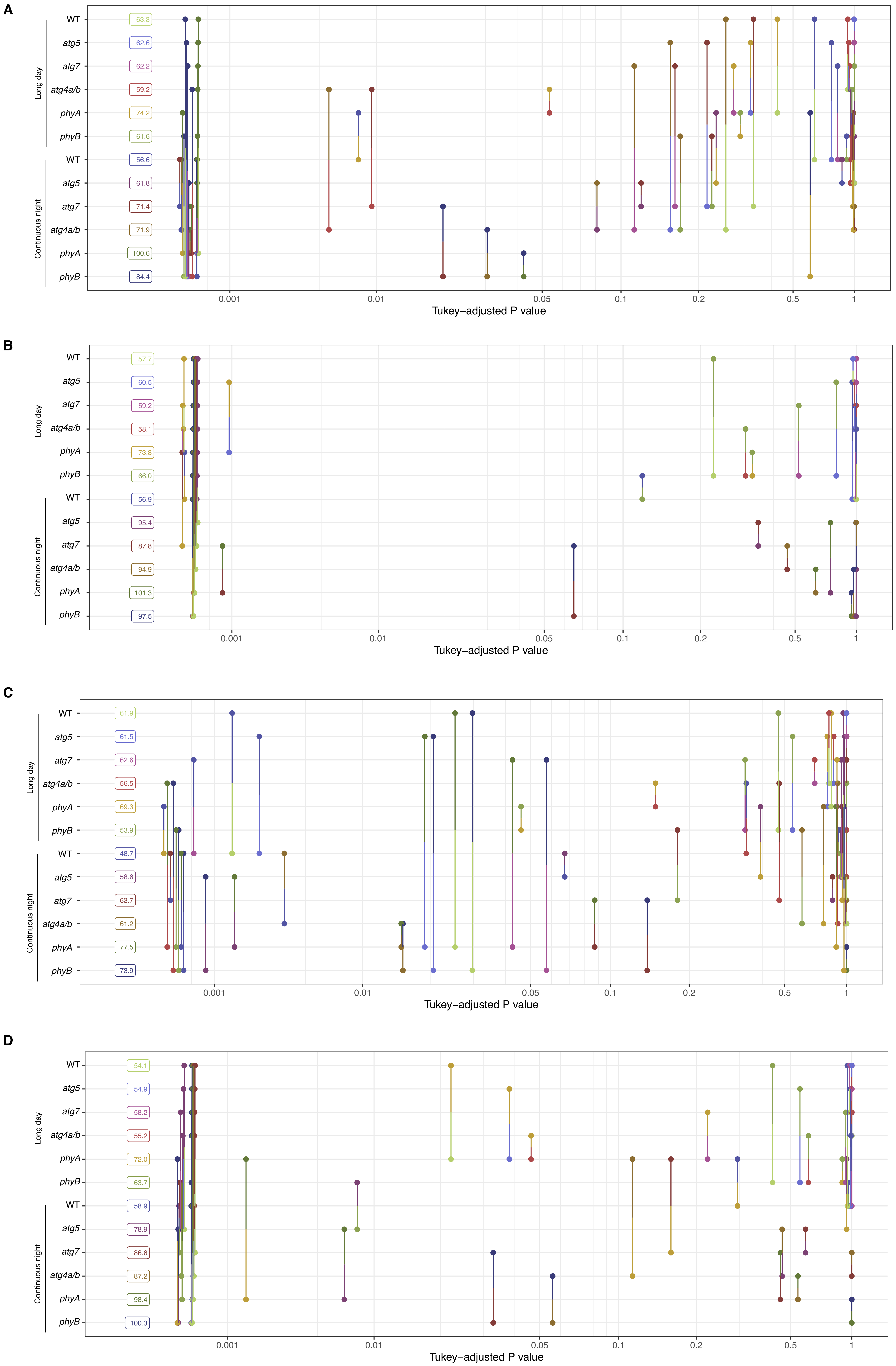


**Figure S1. P-values for Tukey's HSD test of seed germination times.**
Total number of analyzed seeds = 2770, 2160 out of those germinated, and one germinated seed did not pass the QC check. Germination of all genotypes was compared under two illumination conditions on different types of growth media:
(A) Control medium; (B) Nitrogen-deprived medium; (C) Carbon-deprived medium; (D) Medium depleted of both nitrogen and carbon.
