## Supplementary figures and images for "SPIRO – the automated Petri plate imaging platform designed by biologists, for biologists"

### anti-reflection lid for 9 cm round Petri plates v5.png

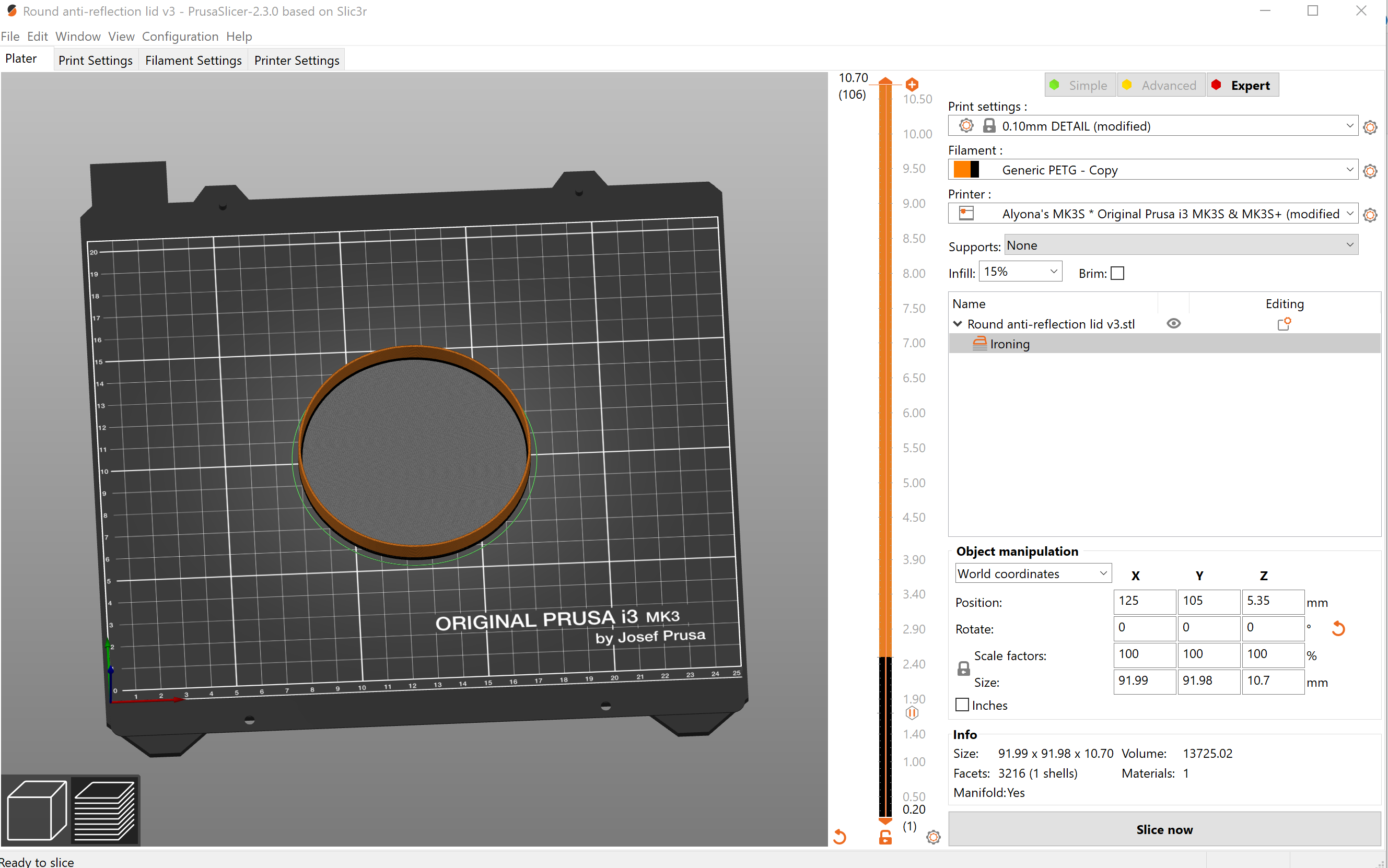

### anti-reflection lid for 12 cm square Petri plates v5.png

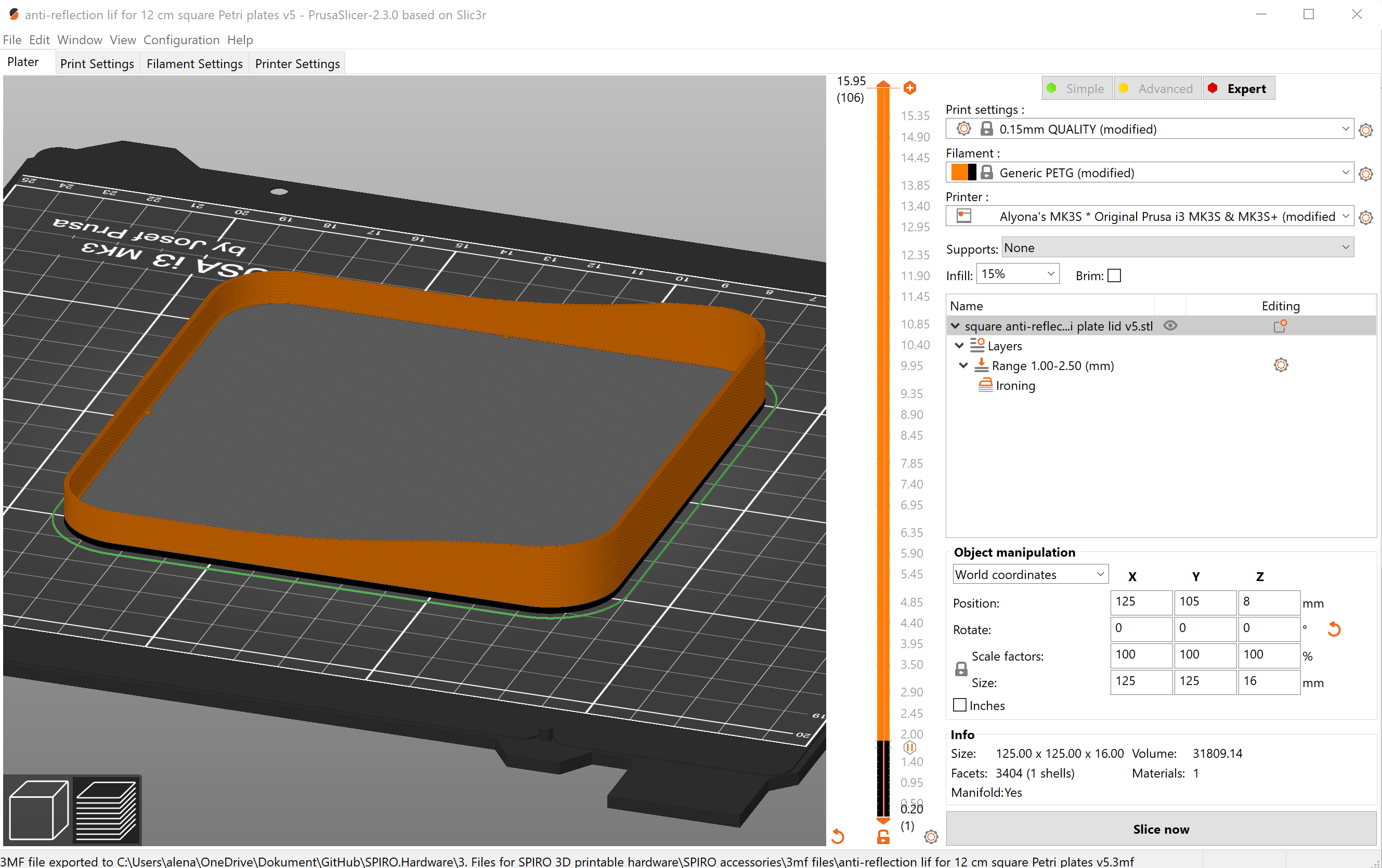

### apple-touch-icon-152x152.png

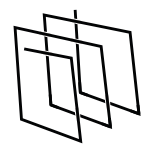

### apple-touch-icon.png

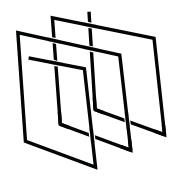

### empty.png

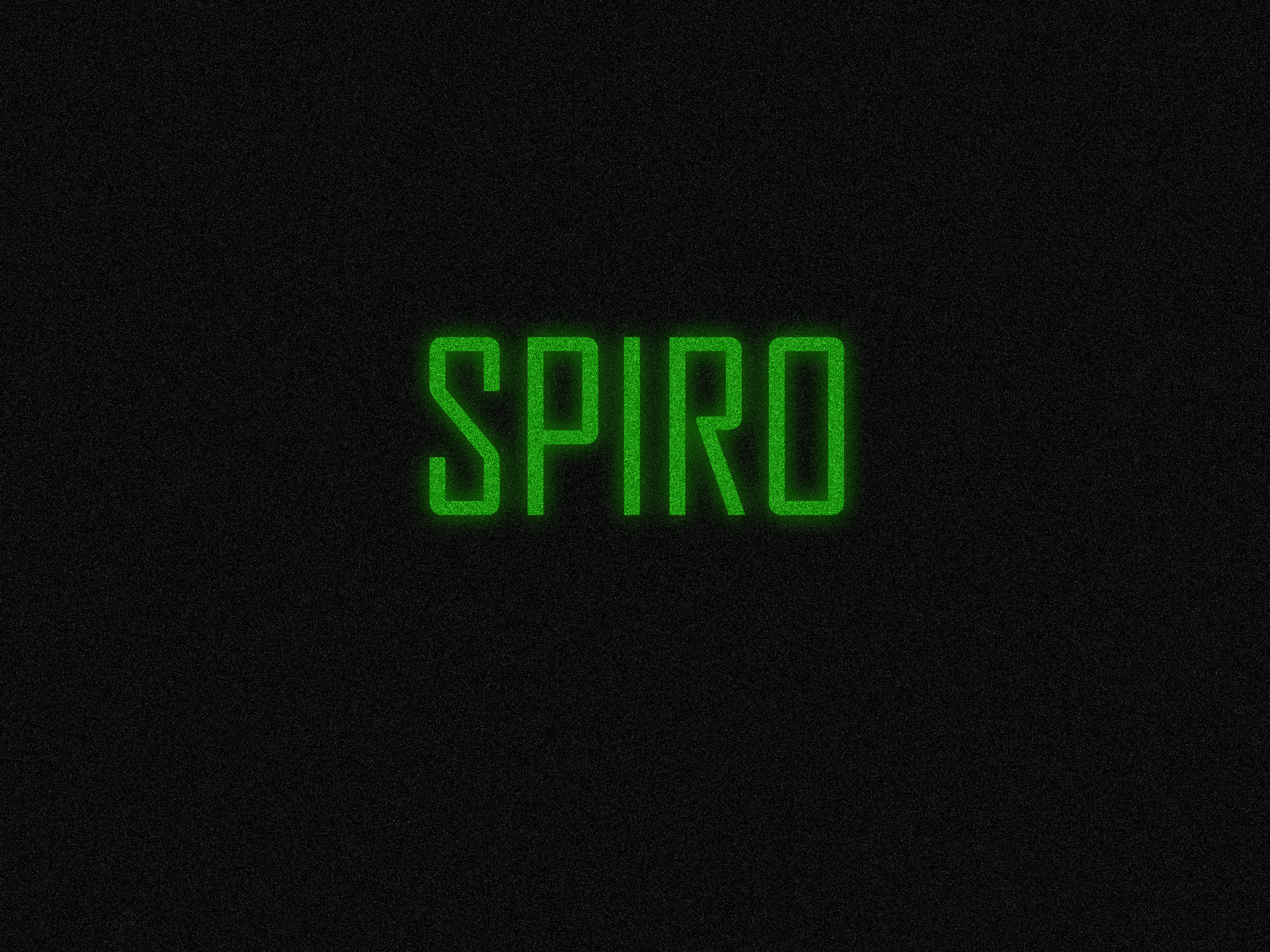

### favicon-16x16.png

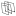

### favicon-32x32.png

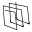

### set 01.jpg

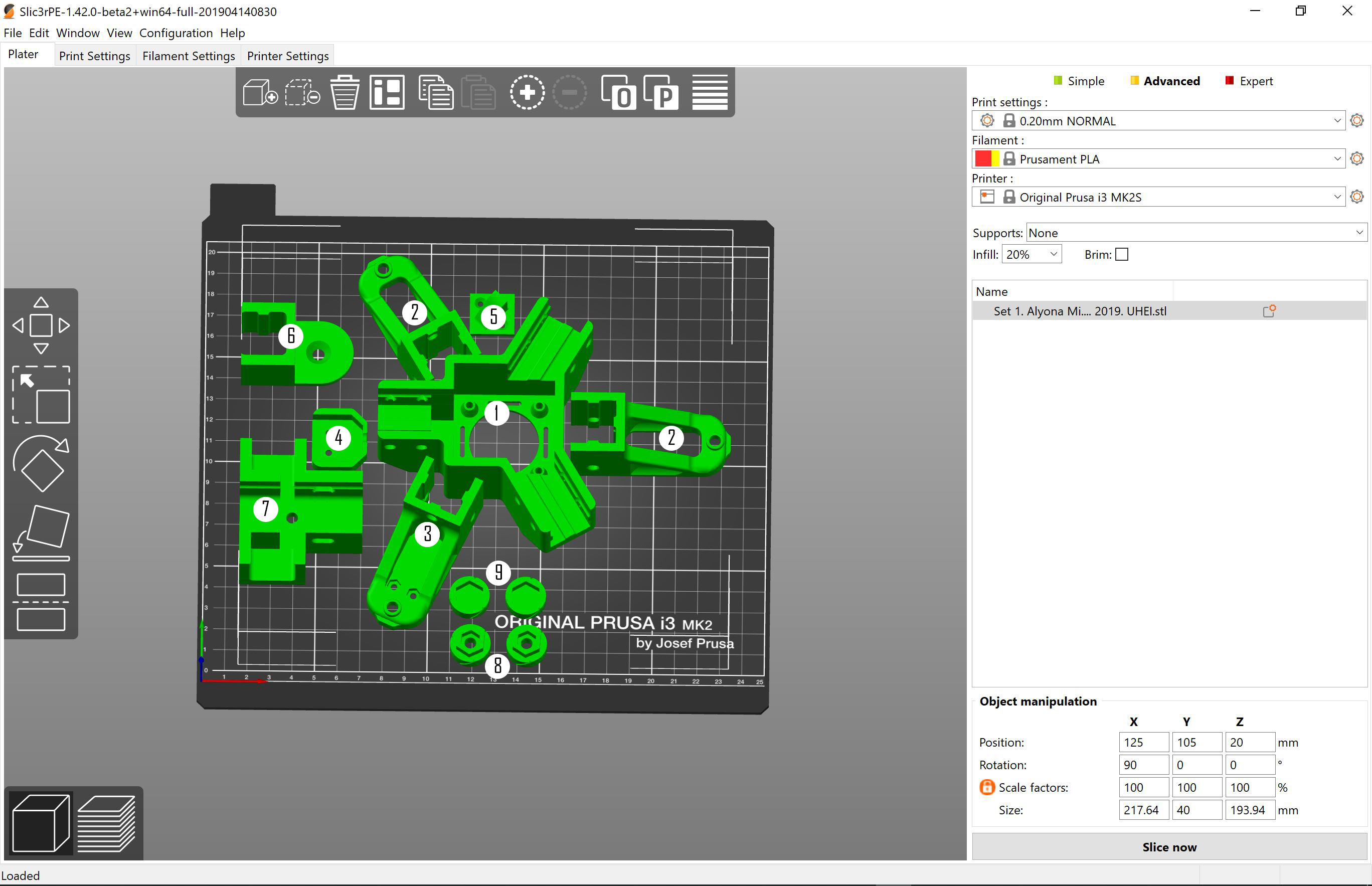

### set 02.jpg

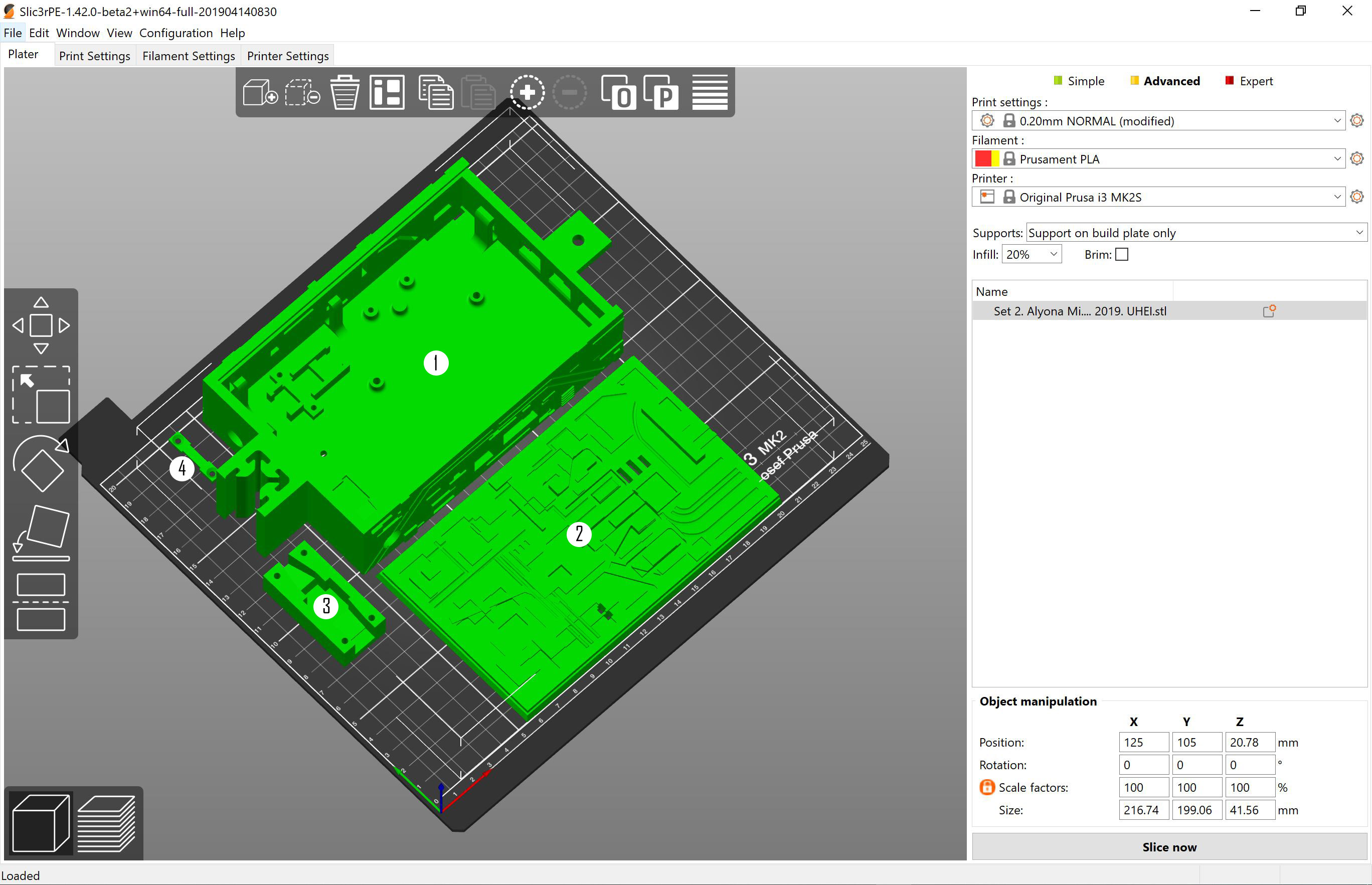

### set 03a.PNG

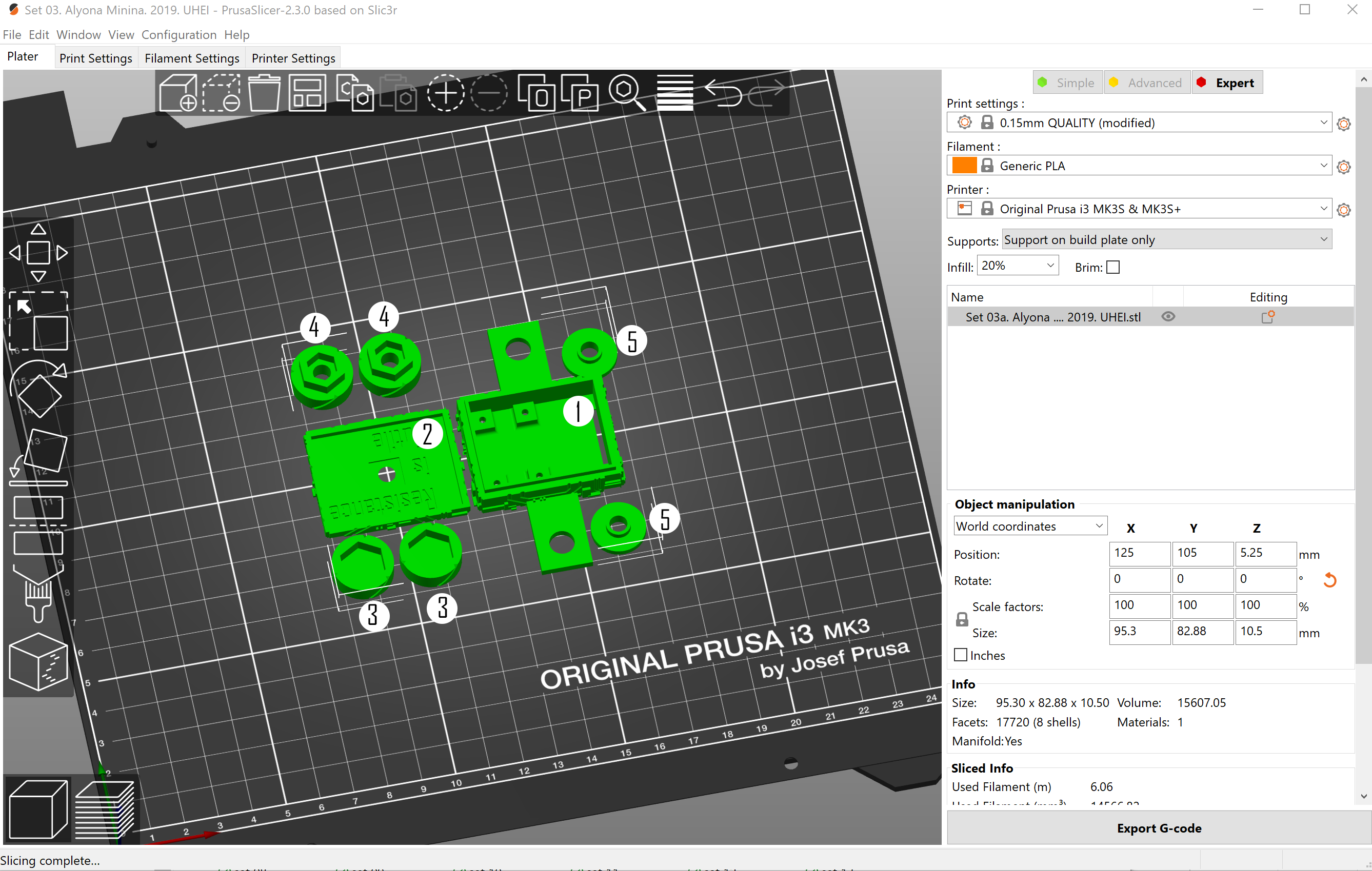

### set 03b.PNG

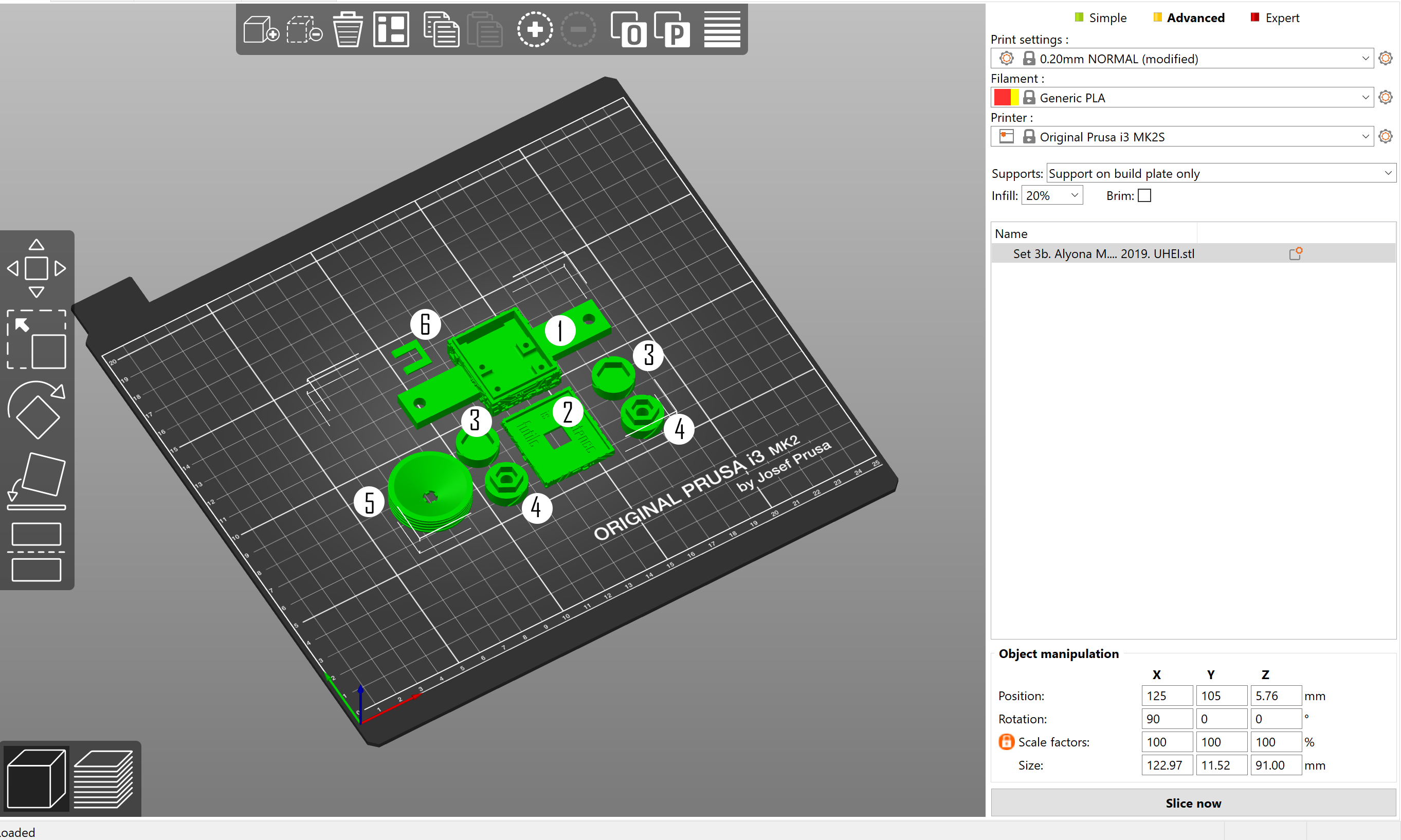

### set 04.jpg

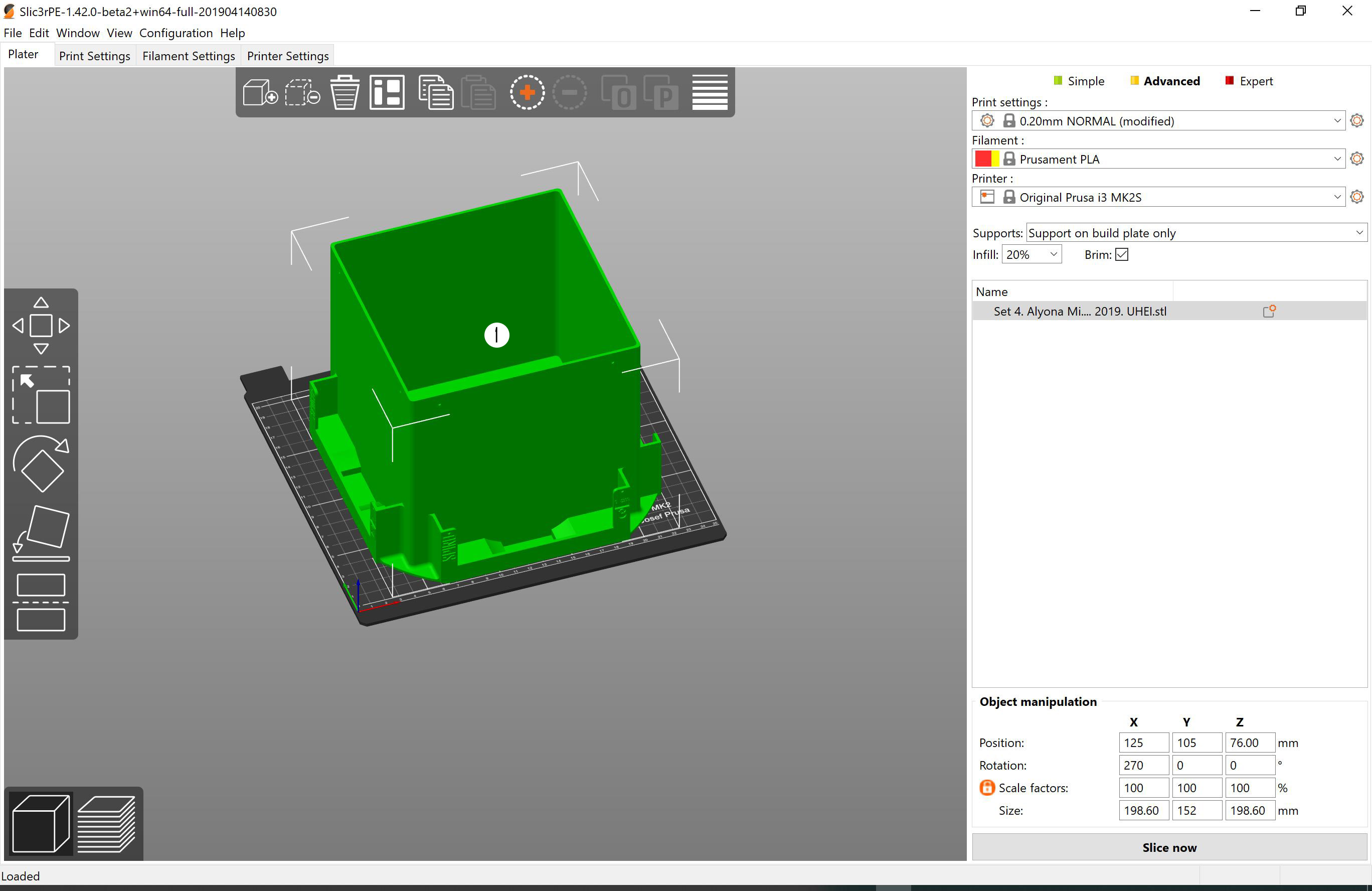

### set 05.jpg

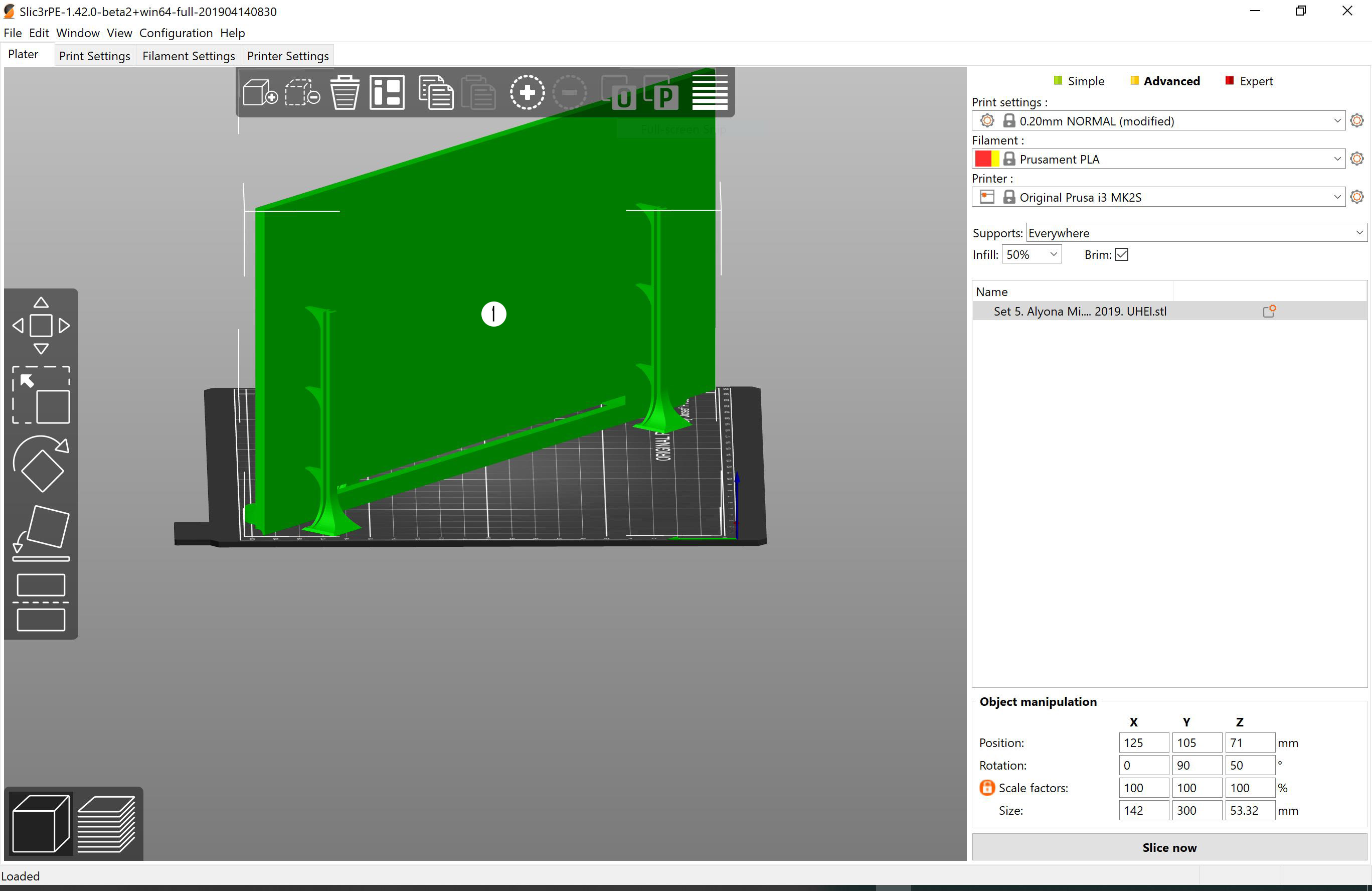

### set 06.jpg

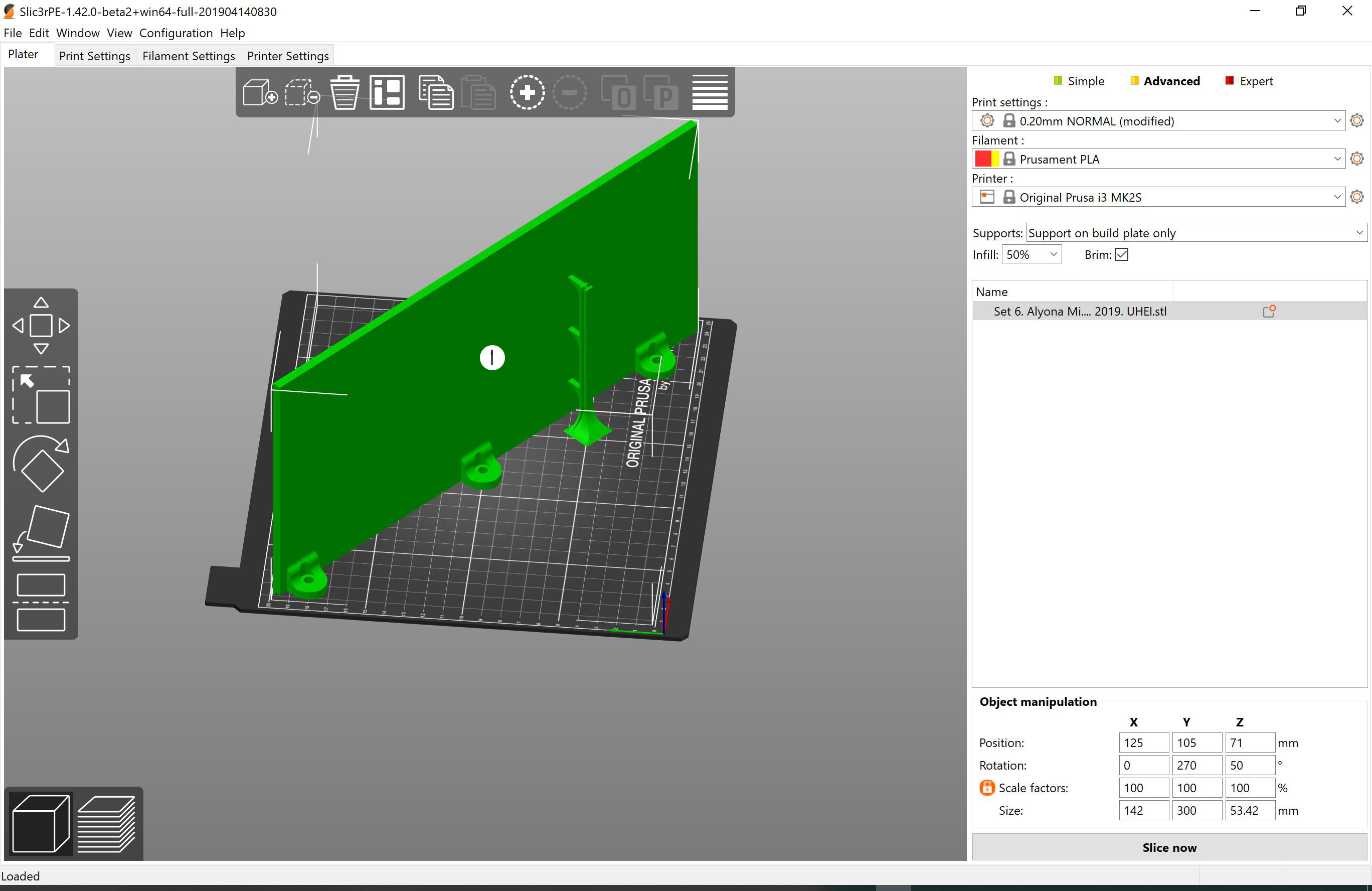

### set 07.jpg

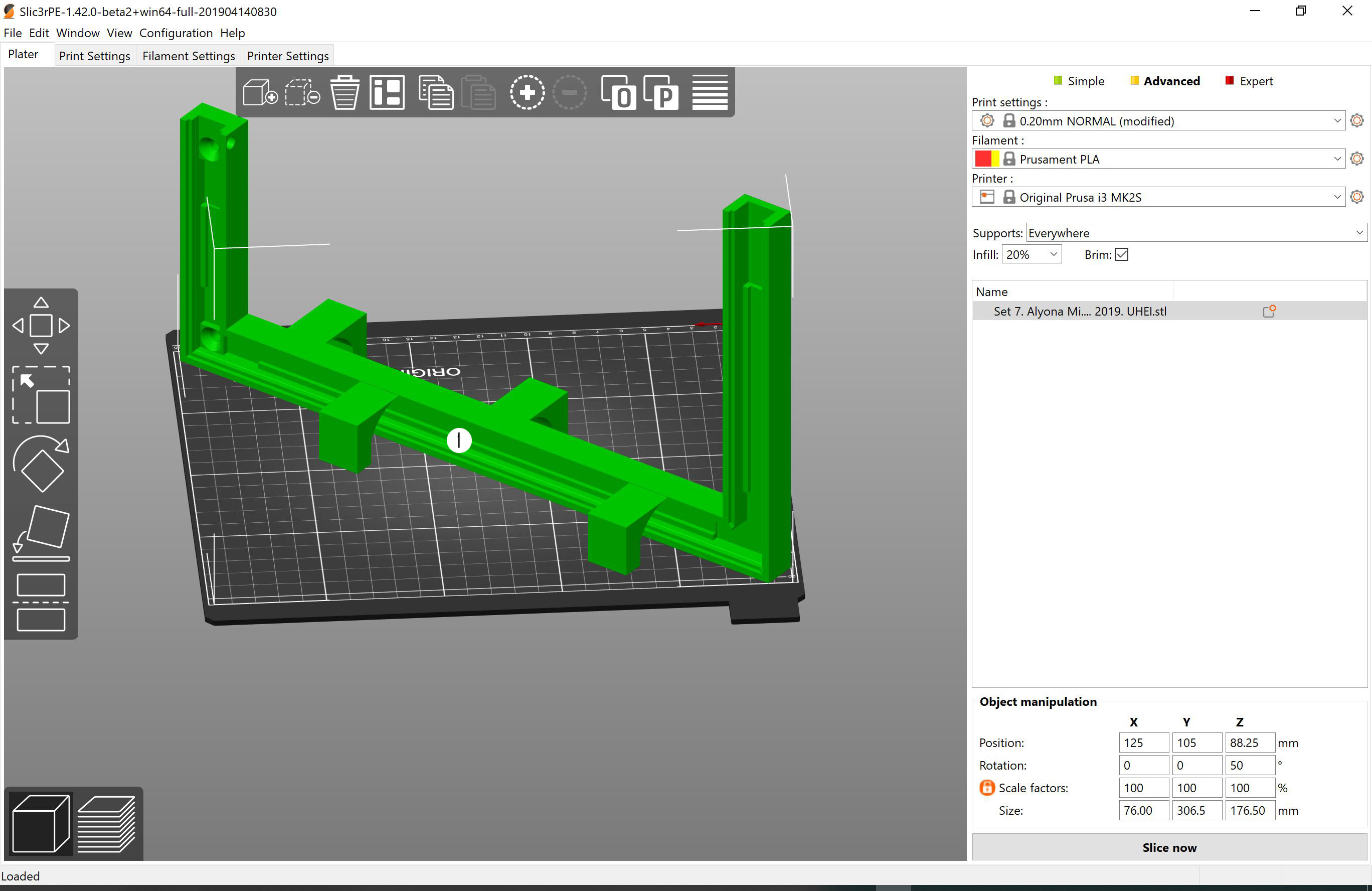

### set 08.jpg

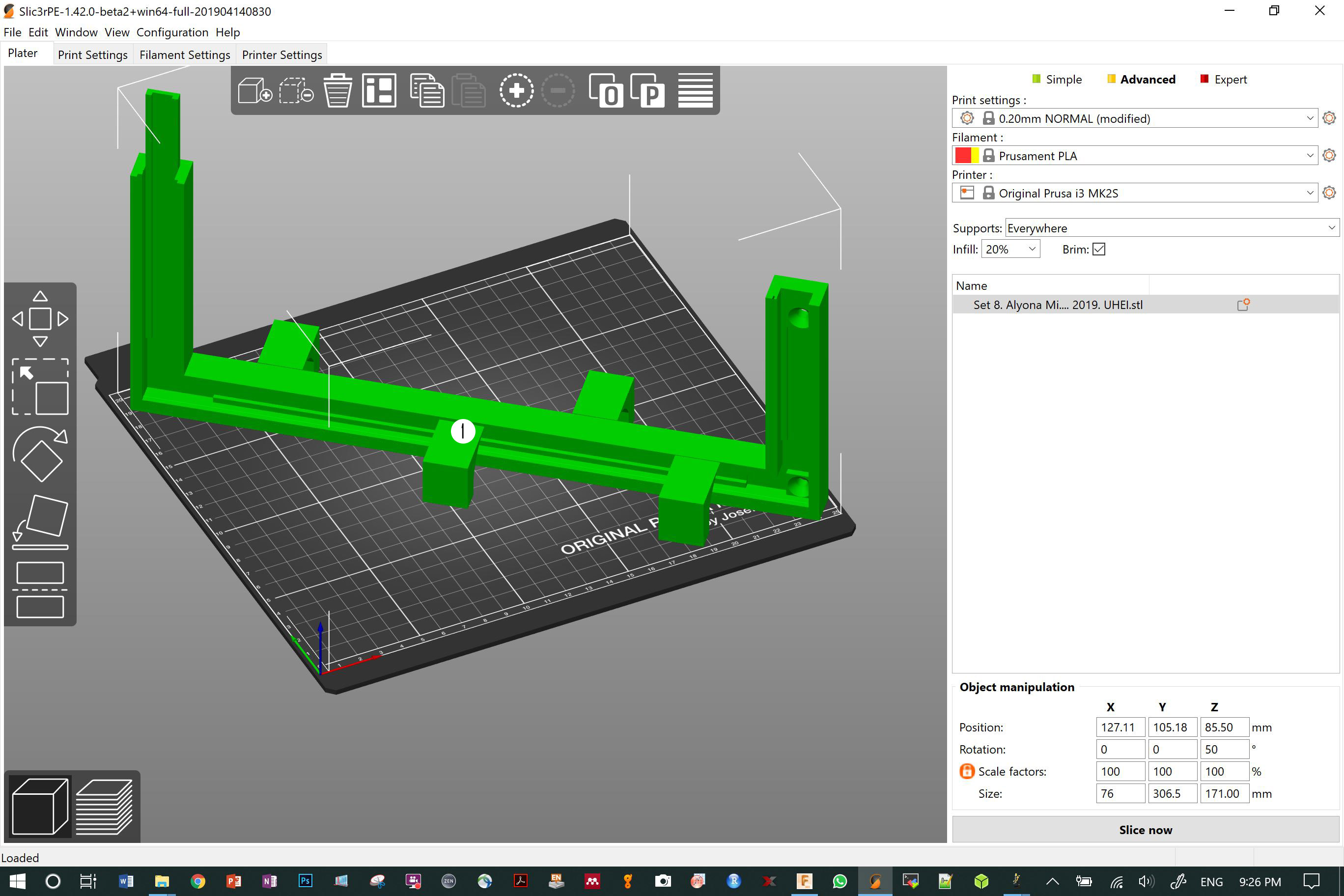

### set 09.jpg

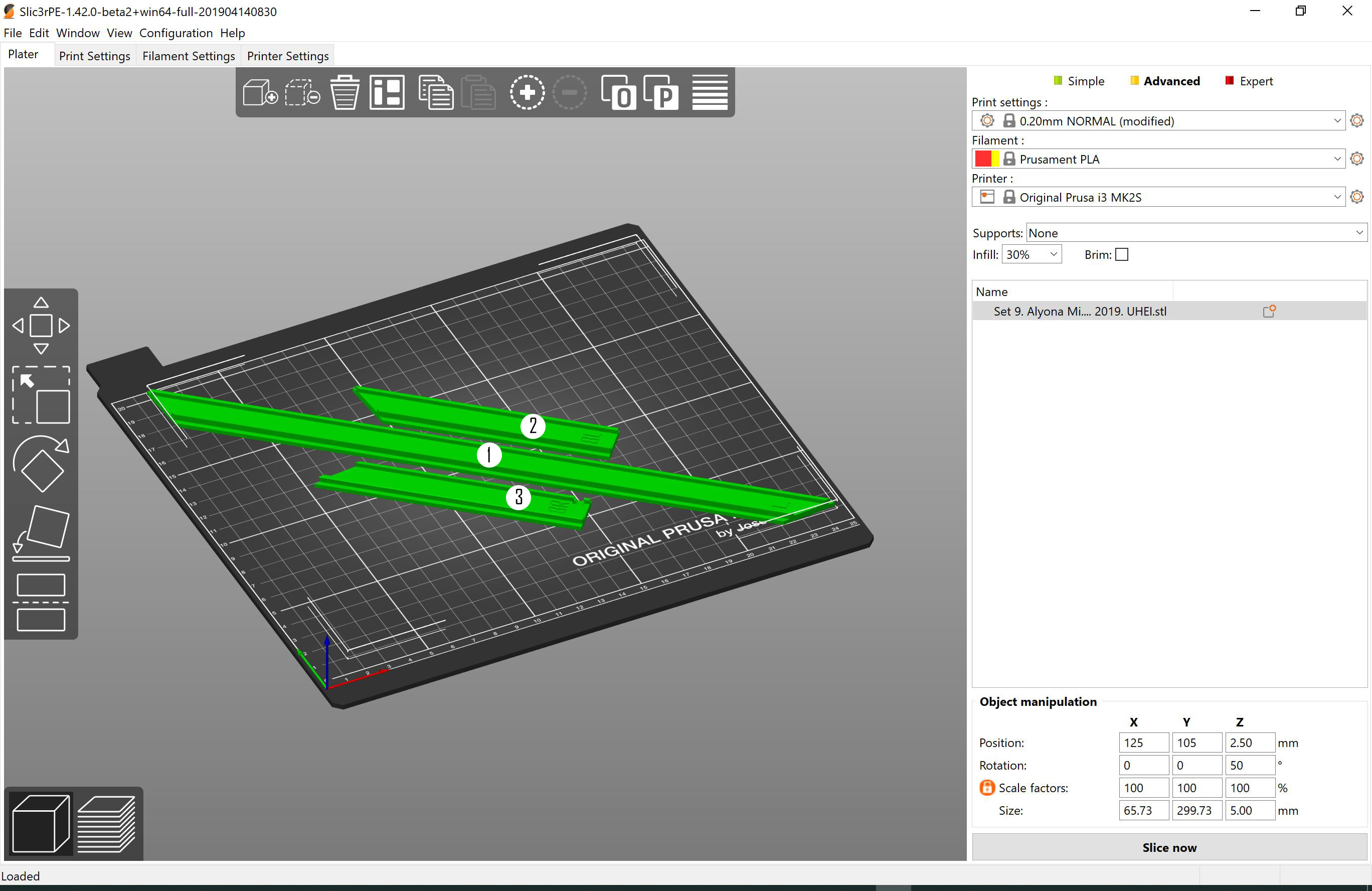

### set 10.jpg

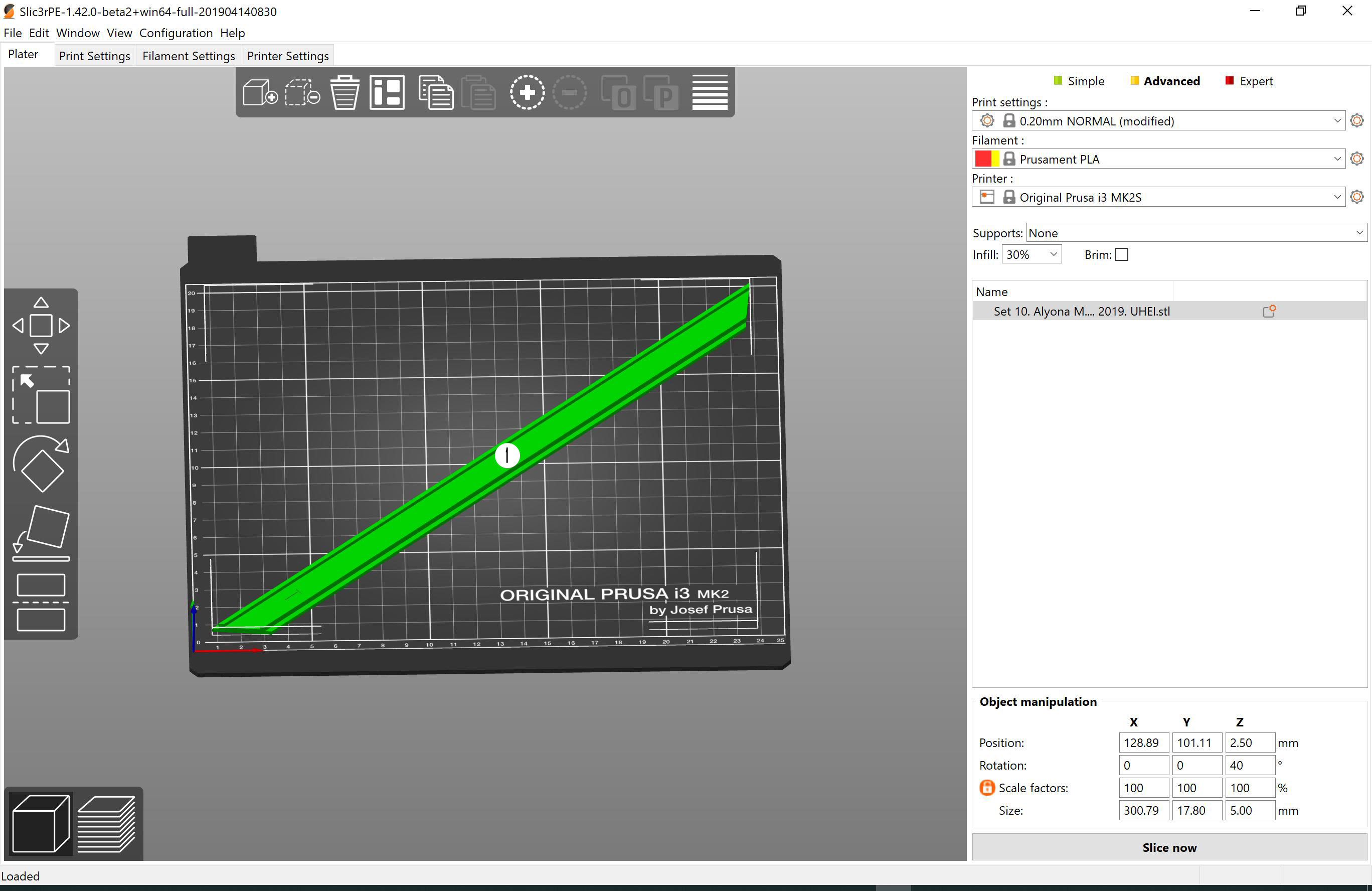

### set 11.jpg

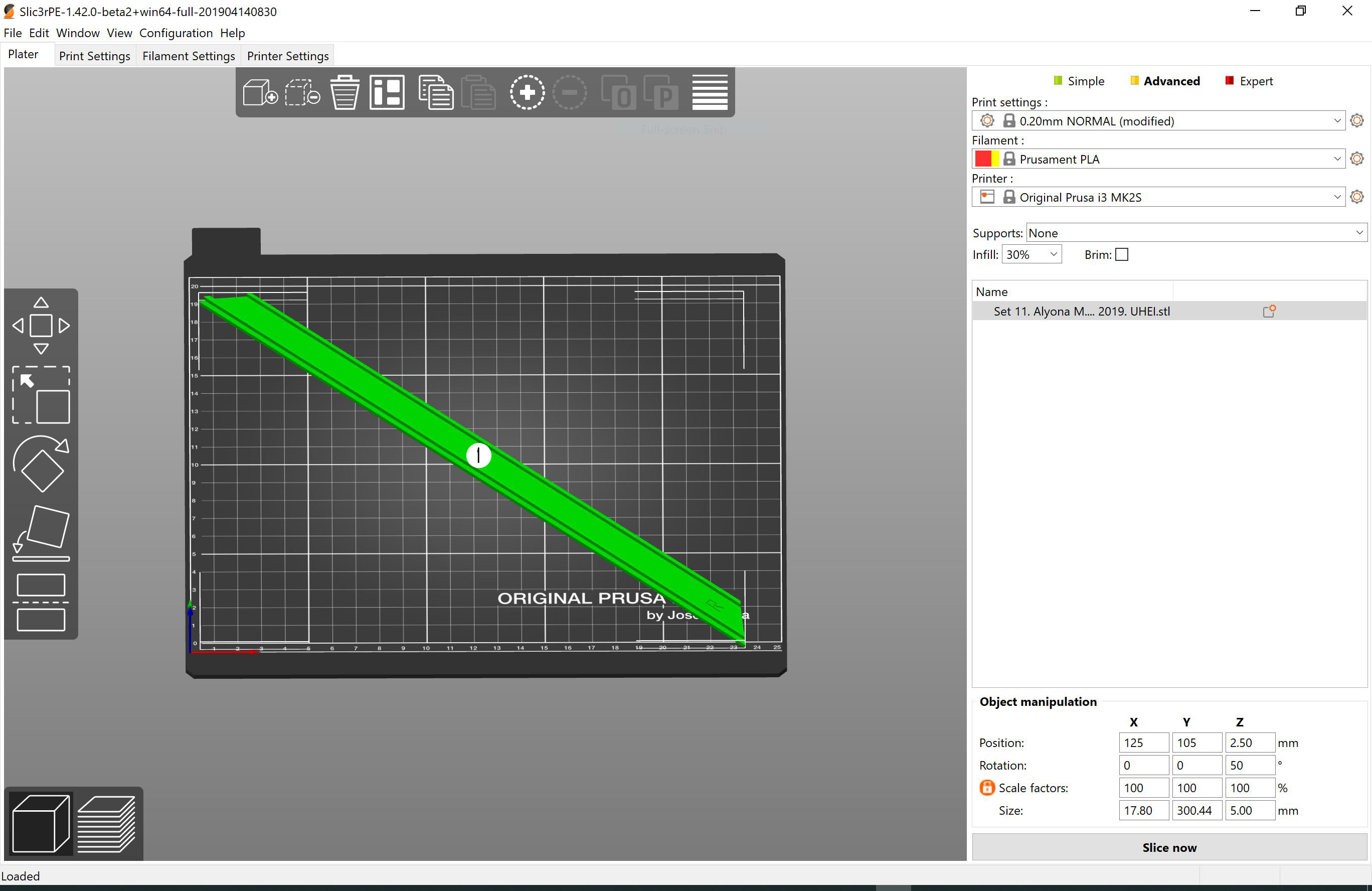

### set 12.PNG

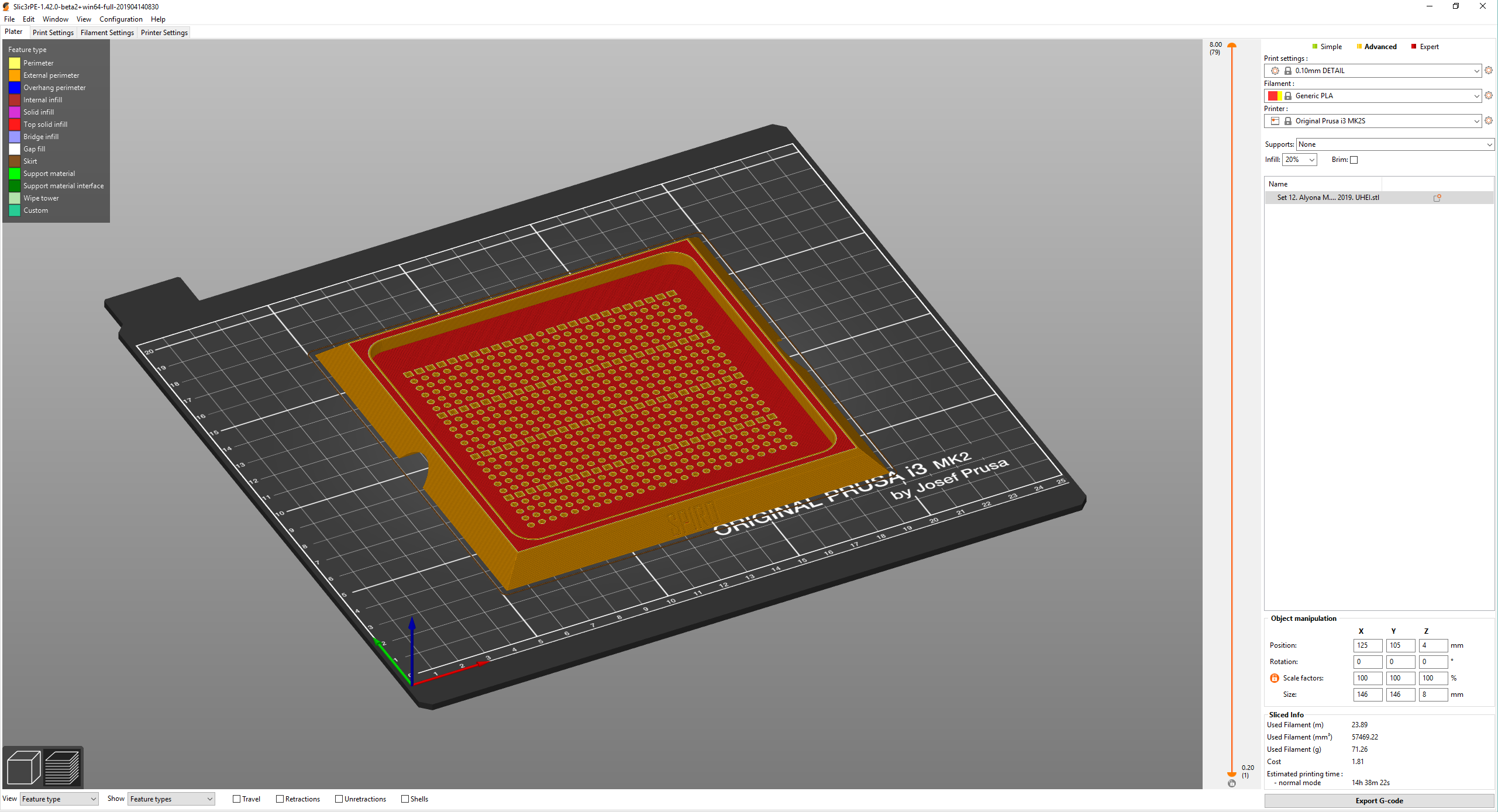

### set 13.PNG

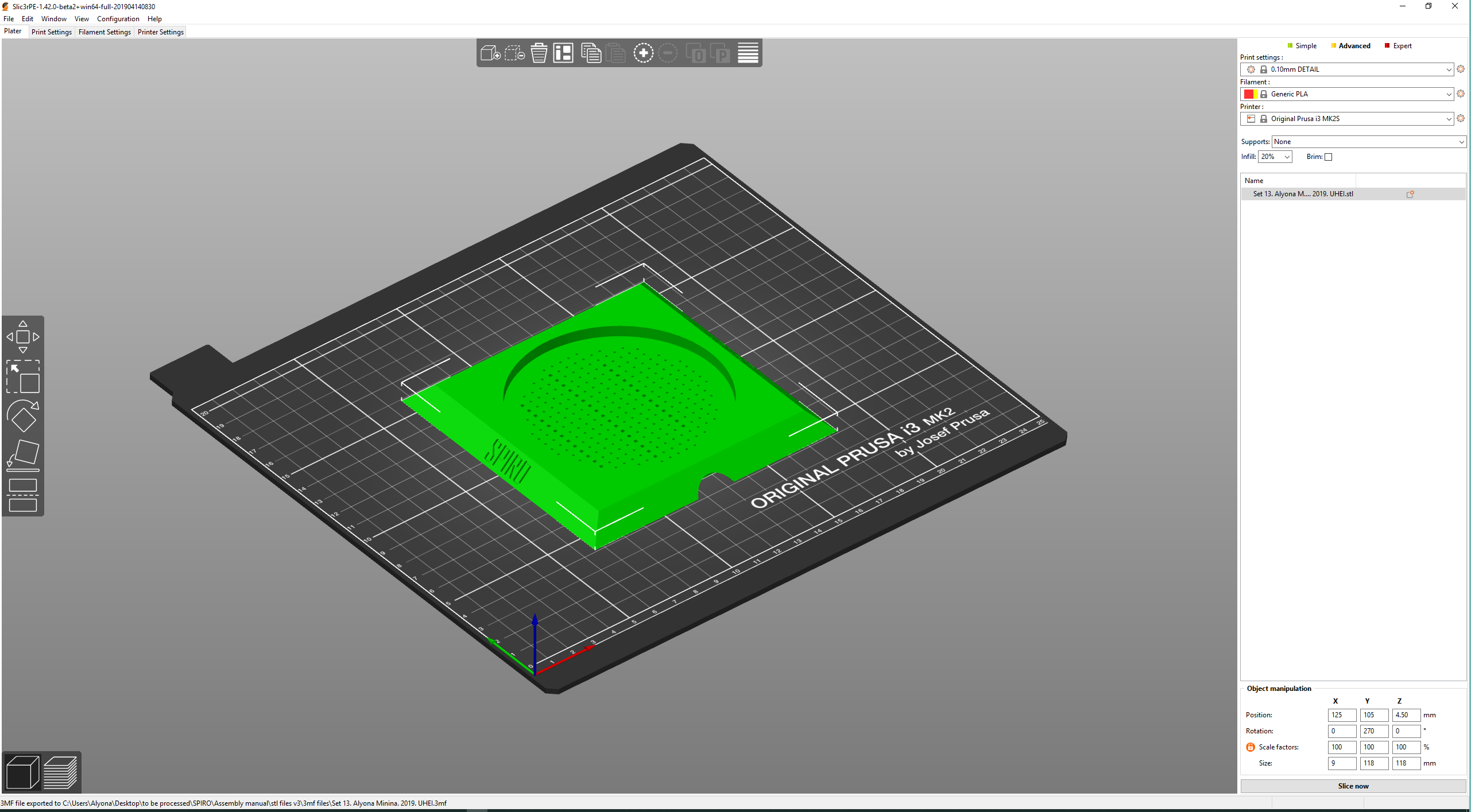

### SPIRO seed plating guide for 9 cm round plates.png

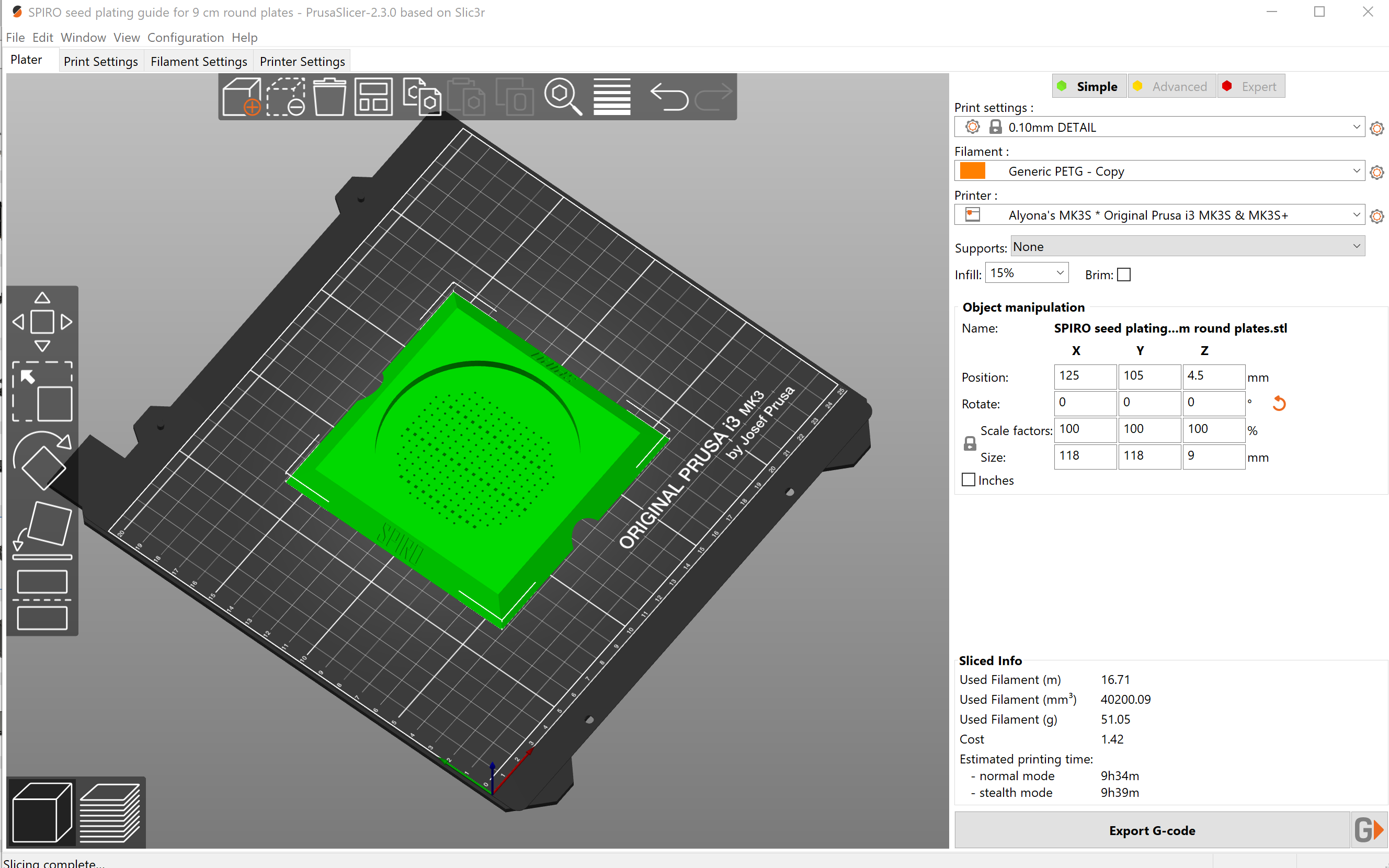

### SPIRO seed plating guide for 12 cm square plates.png

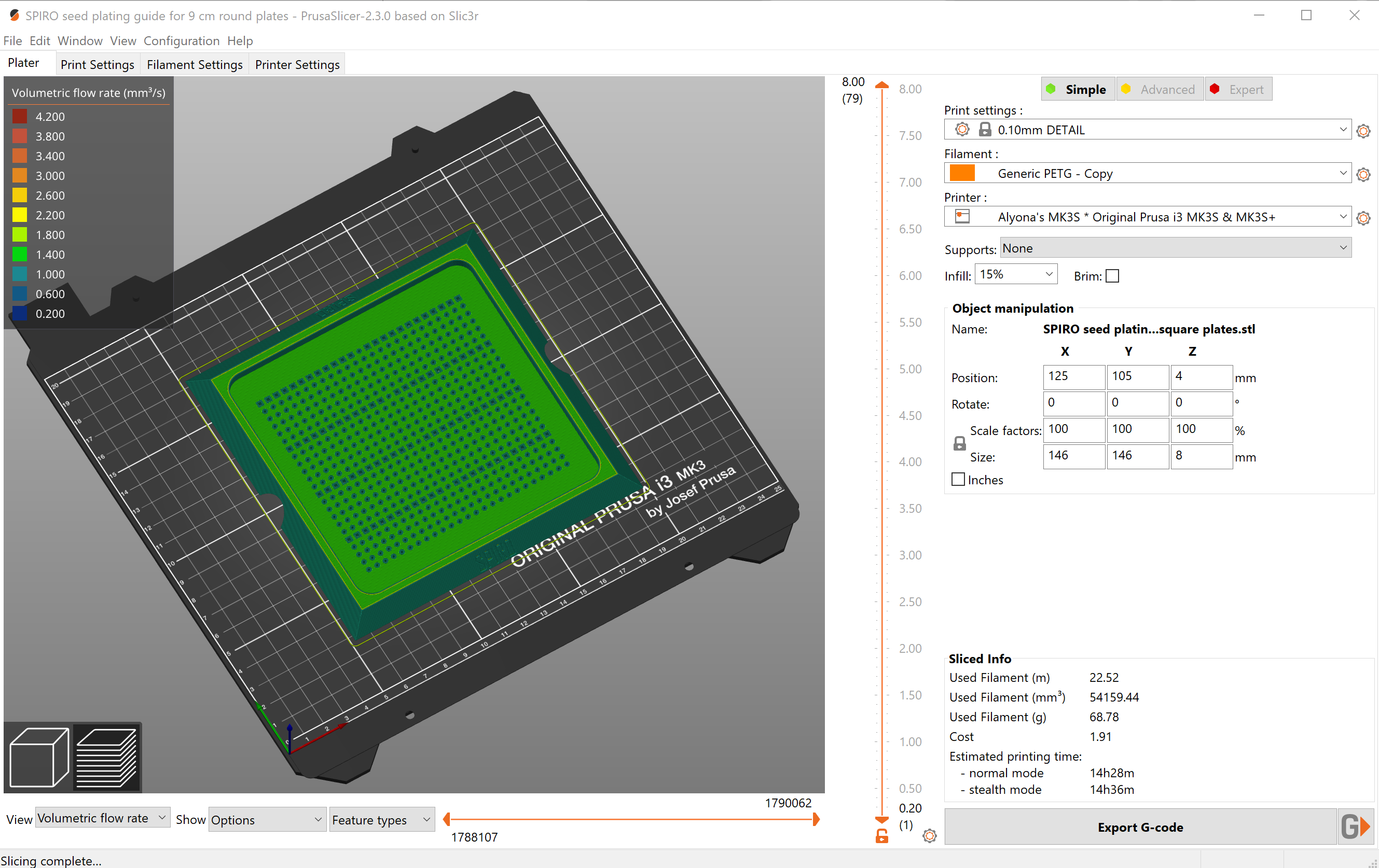

### spiro-folder.png

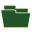
