## Supplementary File 2 for "SPIRO – the automated Petri plate imaging platform designed by biologists, for biologists"

SPIRO

Assembly instructions

#### Overview

SPIRO, the **S**mart **P**late **I**maging **Ro**bot, is a simple imaging system that comprises only a handful of electronics components (**Fig. 1A-B**) and has 3D-printable structural parts (**Fig. 1C-D**). It is designed for automated time-lapse imaging of samples growing on vertically positioned Petri plates, e.g., plant seeds and seedlings, calli, bacterial colonies, fungi, and moss.

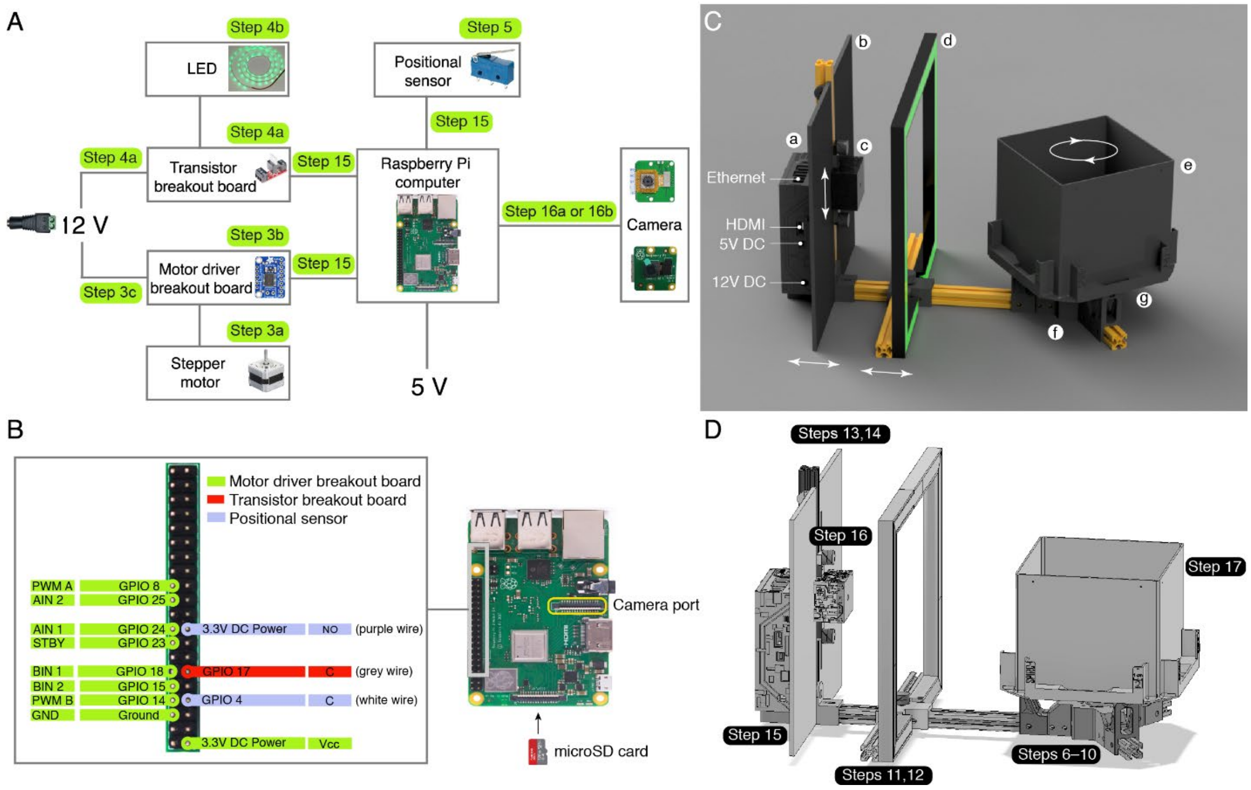

**Figure 1. SPIRO electronics components.**
(**A**) SPIRO wiring scheme. The system can be equipped with either an Arducam camera module with motorized focus mounted on the Raspberry Pi v2 camera (step 16a) or a Raspberry Pi camera with manual focus (step 16b). (**B**) Pinout schematic. The camera is powered and controlled via the camera port on the Raspberry Pi board (yellow frame). (**C**) Overview of SPIRO structural parts: a, Borg's nest harboring the electronics; b, vertical rail with light screen (position can be adjusted along the horizontal rail); c, camera house (position vertically adjustable); d, LED frame with diffusors (horizontally adjustable); e, rotating stage to hold Petri plates; f, motor hub harboring stepper motor and connecting the horizontal rails; g, positional sensor (mini microswitch) used to align the position of the rotating stage with the camera. (**D**) Overview of steps to assemble the structural parts.

A single Raspberry Pi computer controls the rest of SPIRO's electronic components: camera, positional sensor, stepper motor and LEDs. Here we describe two possible configurations of SPIRO with or without motorized focus for the camera. SPIRO is controlled remotely via a web-based user interface, and the motorized camera module allows convenient focus adjustment from the software. However, this configuration is about 25 euros more expensive. A positional microswitch ensures that the Petri plates will be placed straight in front of the camera during imaging. LEDs are powered by a 12 V power supply unit, and switched on/off via a transistor. The stepper motor rotating the stage is controlled via the motor driver breakout board and powered by the same 12 V power supply unit. The SPIRO program and underlying operating system are installed on a microSD card.

SPIRO's laconic configuration was fine-tuned for assembly without any previous experience in 3D printing, use of Raspberry Pi, and mechanical engineering. The user will need to acquire basic knowledge of soldering, which was verified to be achievable by watching a tutorial YouTube video[1].

SPIRO is assembled in 17 main steps, as described below. The time required for assembly (starting from the step 3) can range from several hours to several days depending on the user's skills in wielding a screwdriver. The time required for printing the structural parts is typically seven days, although it can be shorter when using more than one printer.

We strongly recommend to check for potential updates in the SPIRO Hardware Repository[2] prior to purchasing or printing components.

#### 1. Purchase components from Table S1

SPIRO includes a number of standard structural and electronic components that need to be purchased. The complete list of components, required amounts and suggestions for distributors can be found in **Table S1** of the manuscript and **in the** SPIRO Hardware Repository[2].

Please note that shipment costs comprise about a quarter of the building price for a single SPIRO. Additionally, some of the components can be ordered only in superfluous amounts, e.g., LED strip. Thus, building several SPIROs simultaneously will likely reduce the cost per robot.

#### 2. Print structural parts of SPIRO

The printable structural parts of SPIRO are combined into 11 sets (**Table S2**, **File S1**, SPIRO Hardware Repository[2]). According to our experience, printing all sets takes approximately seven days and requires about 1.5 kg of matte black PLA plastic and 100 g of semi-transparent PLA plastic.

All components were printed using Prusa i3 MK2S printer and additionally tested on a Prusa i3 MK3. If you do not have access to a 3D printer, it is possible to find publicly available printers in centers like Makerspaces or, alternatively, to order 3D prints from online services (3dhubs or similar).

We have thoroughly verified and optimized printing settings for each set of parts. It is highly recommended to follow the printing instructions provided in the table to achieve the best possible quality. As of today, SPIRO hardware has been tested using only PLA plastic. It is sufficiently robust, is the cheapest type of filament and also the easiest one to print with. If desired, SPIRO can be also printed using PETG or ABS filament, however it might require adjustments in scaling of the models to compensate for shrinkage of the printed parts.

For 3D printing, model files need to be converted into g-code files, which then can be read by a 3D printer to print the component. Such conversion can be done in a slicing software of users' choice. For this project we used PrusaSlicer v2.1[3]. In addition to the physical parameters of a printable object, g-code files also contain information about the type of filament and the printer to be used, how much plastic should be used for filling in the object's volume (infill), position of the model on the print bed etc.

**File S1** contains .stl, .3mf and .f3d type of files for each hardware set of SPIRO. Additionally, we include screenshots of creating g-code files for each SPIRO set in PrusaSlicer.

**.stl** files can be used to create g-code files in any slicing software. Please use the printing settings recommended in **Table S2** of the manuscript or **the** SPIRO Hardware Repository[2].

**.3mf** project files already contain recommended settings optimized for printing using Prusa i3 MK3/S. These files can be used for example in the PrusaSlicer software to adjust filament and printer settings and generate g-codes files.

**.f3d** can be used for modifying our design. This can be done using the Autodesk Fusion 360 software, which is distributed with free license for educational facilities.

#### 3. Wire the stepper motor

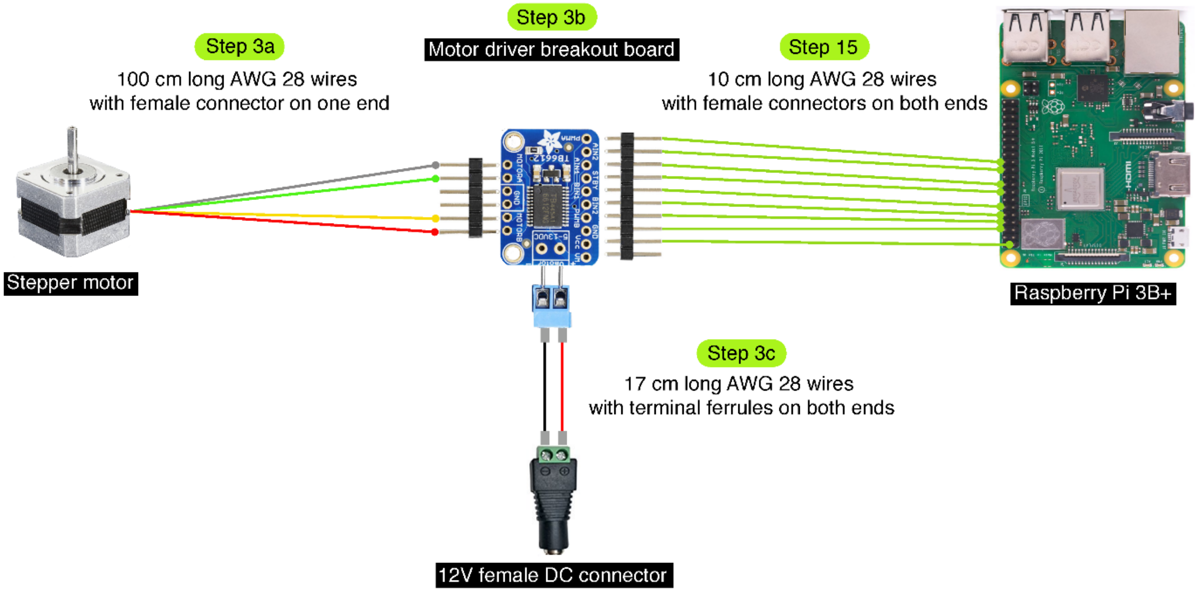

**Figure 2. Wiring the stepper motor.**
This step comprises three sub-steps: 3a, soldering four wires to the motor to have a total length of 100 cm and adding female connectors. 3b, soldering components to the motor driver breakout board (see Fig. 3): the pin headers (b) and the two-pin screw terminal (c) should be soldered to the board (a). Note the orientation of the parts in the description in the corresponding section. 3c, wiring the 12 V DC female connector to the screw terminal on the motor driver breakout board. Connecting the wires from the motor to the driver and from the driver to the Raspberry Pi is done in a later (step 15).

#### 3a. Solder stepper motor wires

Each of the four wires of the motor should be extended to a total length of 100 cm and have a female connector at the end.

- The type of solder wire we have used contains 60% tin and 40% lead with a rosin core. However, due to health reasons it is generally recommended to use lead free solder wire.
- When extending the wires, please use jumper cables with colors corresponding to the wires already attached to the motor: grey, green, yellow, red (**Fig. 2**).
- Total length of each wire: wire already connected to the stepper motor + AWG 28 cable + jumper cable with one female connector = 100 cm.
- Measure the length of the wires already attached to the motor.
- Cut off one connector from the jumper cable to leave only one female connector attached.
- Cut the AWG 28 cables to the length required so that each wire is 100 cm in total length.
- Follow the instructions in video 3a. Stepper motor wires for wire connection and insulation.

| **Component** | **# in Table S1** |
|---|---|
| *NEMA 17 stepper motor* | 4 |
| *AWG 28 color cables* | 15 |
| *AWG 28 jumper cables with female connector* | 11 |
| *Heat-shrink tube 1–2mm in diameter* | 13 |

| **Tool** | **# in Table S1** |
|---|---|
| *Soldering iron + solder* | 46 |
| *Lighter* | – |
| *Scissors* | – |
| *Wire stripper* | 44 |

#### 3b. Solder the motor driver breakout board

- Solder the pin headers to the board with the long pins extending from the labeled side of the board (**Fig. 3A**). The pin headers are included into the Motor driver breakout board kit as a single strip that needs to be broken into two shorter strips with 6 and 10 pins.

- Solder the two-pin screw terminal to the *Vmotor* holes. Make sure that the terminal and the long pins are on the same side of the board and that the metal ports of the terminal are facing away from the board (**Fig. 3B**).

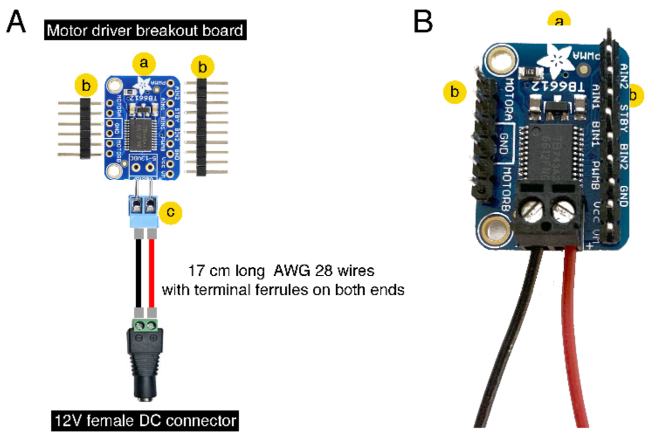

**Figure 3. Soldering the motor driver breakout board components.** (**A**) Components to be soldered: a, PCB; b, pin headers; c, two-pin screw terminal. (**B**) Soldered driver with power wires connected to the screw terminal.

| Component | # in Table S1 |
|---|---|
| *Motor driver breakout board kit* | 3 |
| *Two-pin screw terminal 3.5 mm pitch* | 10 *(if not included in the kit)* |

| Tool | # in Table S1 |
|---|---|
| *Soldering iron + solder* | 46 |

#### 3c. Wire the motor driver to the DC connector

- Use two 17 cm long AWG 28 wires (black and red).
- Secure terminal ferrules on both ends of each wire. For instructions see the video **Securing terminal ferrules**.
- Connect the "+" screw port of the motor driver and the DC connector using the red wire, and the "–" port with the black wire.
- Tighten the screws to secure the wires in place.
- Please note that there will be another pair of wires going into the ports of the DC connector to power the SparkFun MOSFET power control kit (**Fig. 5D**).

| **Component** | **# in Table S1** |
|---|---|
| *AWG 28 color cables* | 15 |
| *Terminal ferrules 0.5 mm² (not insulated)* | 14 |
| *Heat-shrink tube 2 mm in diameter* | 13 |

| **Tool** | **# in Table S1** |
|---|---|
| *Lighter* | – |
| *Scissors* | – |
| *Wire stripper* | 44 |
| *Ferrule crimper* | 45 |

### 4. LED wiring

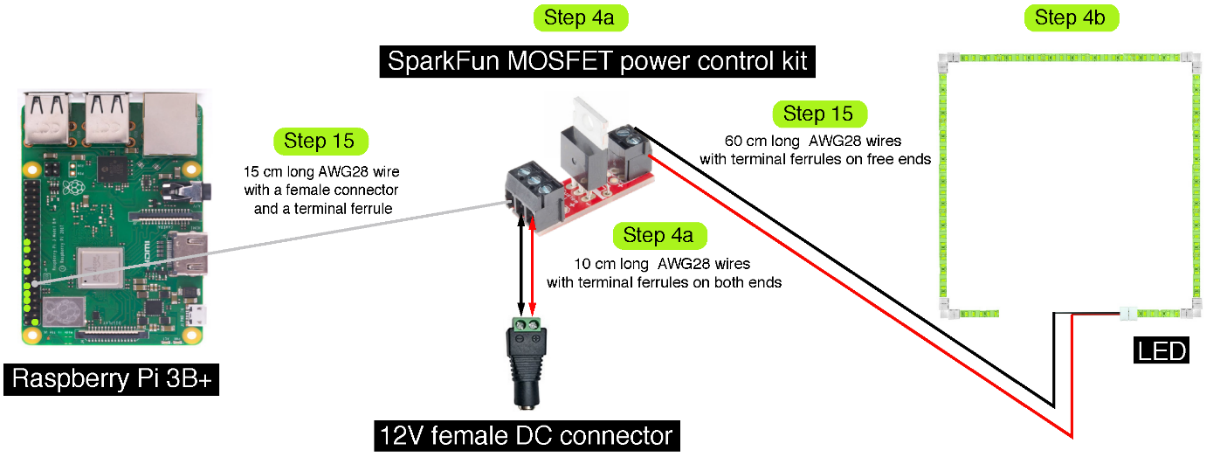

**Figure 4. LED wiring.**

These steps will include soldering the transistor that will be used to control the LED (SparkFun MOSFET power control kit, step 4a) and connecting the transistor to the 12V DC power supply. The LED strip is assembled into a square using 90° corner connectors (step 4b) and a terminal LED connector. The LED is attached to the SparkFun MOSFET power control kit later during SPIRO assembly (step 15).

#### 4a. Assemble the SparkFun MOSFET power control kit

- Solder the resistor, transistor and both screw terminals to the SparkFun MOSFET power control kit board (**Fig. 5**).

> Important! Don't be like Jonas, take extra care to solder part **b** in the correct direction (**Fig. 5B and C**).

- Trim the ends of **b** and **c** after the soldering is complete (**Fig. 5**).

- Connect the three-pin screw terminal to the 12V DC connector:
  - Use two 10 cm long 28 AWG wires (black and red).
  - Secure electric wire ferrules on both ends of each wire. For instructions see the video [Securing terminal ferrules].
  - Use the red wire to connect the "+" port of the three-pin screw terminal of the SparkFun kit to the "+" port of the DC connector.
  - Use the black wire to connect the "–" port of the three-pin screw terminal of the SparkFun kit to the "–" port of the DC connector.
  - Tighten the screws to secure wires in place.

- Use a 15 cm long grey jumper cable with a female connector. Secure an electric wire ferrule on the free end of the wire. For instructions, see the video [Securing terminal ferrules].

- Place the ferrule protected end of this wire into the "C" port of the three-pin screw terminal.

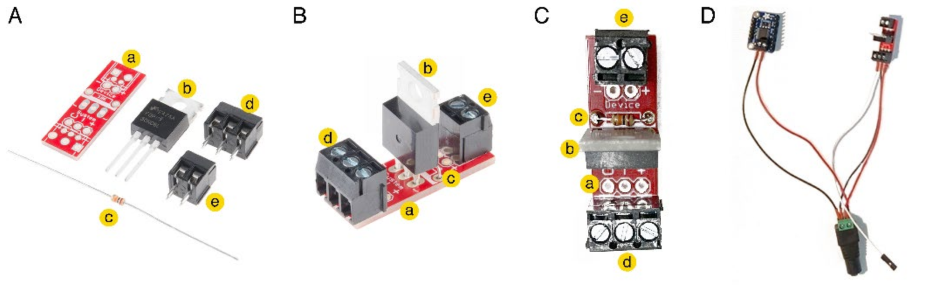

**Figure 5. Assembling the SparkFun MOSFET power control kit**

(**A**) Components to be soldered: a, PCB; b, transistor; c, resistor, d, three-pin screw terminal; e, two-pin screw terminal (the photo is modified from SparkFun.com). (**B**) The side view of the soldered kit. Please note the orientation of the transistor (b) and the screw terminals (d and e). (**C**) Top view of the soldered kit, note the orientation of the transistor (b) and position of the resistor (c). (**D**) SparkFun MOSFET power control kit and motor driver wired to the 12V female DC connector.

| **Component** | **# in Table S1** |
|---|---|

| | |
|---|---|
| *SparkFun MOSFET power control kit* | 7 |
| *AWG 28 color cables* | 15 |
| *Terminal ferrules 0,5 mm² (not insulated)* | 14 |
| *Heat-shrink tube 2mm in diameter* | 13 |
| **Tool** | **# in Table S1** |
| *Soldering iron + solder* | 46 |
| *Lighter* | – |
| *Scissors* | – |
| *Wire stripper* | 44 |
| *Ferrule crimper* | 45 |

#### 4b. Solder wires to the LED terminal connector and assemble the LED strip

- Cut the LED strip into three 25 cm long segments (these will be the left, top and right parts of the LED square) and two 5 cm long segments (these will be the bottom left and bottom right parts of the LED square). Connect the segments using the corner connectors. Make sure the "+" contacts on the corners are connected to the "+" contacts on the LED strip (**Fig. 4**).

- Extend the two wires already attached to the LED terminal connectors so that they have a total length of 50 cm.

- Crimp a ferrule onto the free end of each wire and secure it with the heat-shrink tube. For instructions, see the video **Securing terminal ferrules**.

- Bind both wires together with a coiled cable protection. Cover the 40 cm length starting from the connector. Leave the last 10 cm free of cover.

- Attach the terminal connector to the right bottom strip of the LED square. For instructions, see the video **LED strip assembly**.

| **Component** | **# in Table S1** |
|---|---|
| *8 mm green LED strip* | 8 |
| *4x 8 mm 2-pin LED strip corner connectors* | 9 |
| *8 mm 2-pin LED strip terminal connector* | 9 |
| *AWG 28 color cables* | 15 |
| *Terminal ferrules 0,5 mm² (not insulated)* | 14 |
| *Heat-shrink tube 2mm in diameter* | 13 |
| *Coiled cable protection* | 16 |

| **Tool** | **# in Table S1** |
|---|---|
| *Soldering iron + solder* | 46 |
| *Lighter* | – |
| *Scissors* | – |
| *Wire stripper* | 44 |
| *Ferrule crimper* | 45 |

#### 5. Solder the positional sensor (mini microswitch) wires

- Solder two 110 cm long AWG 28 wires to the microswitch contacts (**Fig. 6**):
    - The purple wire should be attached to contact "NO".
    - The white wire should be attached to contact "C".
    - Each wire should end with a female connector.
- Follow the instructions in video 3a. Stepper motor wires for wire connection and insulation.

| **Component** | **# in Table S1** |
|---|---|
| *Positional sensor (mini microswitch)* | 19 |
| *AWG 28 color cables* | 15 |
| *AWG 28 jumper cables with a female connector* | 11 |
| *Heat-shrink tube 2 mm in diameter* | 13 |

| **Tool** | **# in Table S1** |
|---|---|
| *Soldering iron + solder* | 46 |
| *Lighter* | – |
| *Scissors* | – |
| *Wire stripper* | 44 |

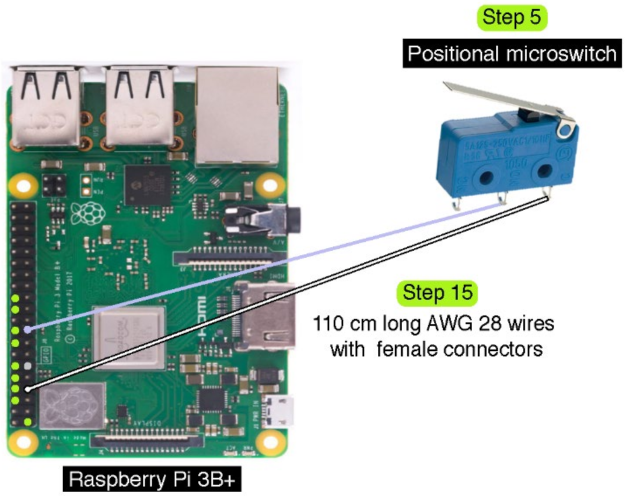

**Figure 6. Wiring the positional microswitch.** The positional microswitch acts as a bridge between the 3.3V pin on the Raspberry Pi (purple wire) and its GPIO pin (white wire). When the switch is activated, the bridge is closed, indicating the position of the imaging stage.

#### 6. Mount the motor hub

Follow instructions in video 6. Mounting the motor hub.

| Component | # in Table S1 |
|---|---|
| *The stepper motor wired in the step 3* | - |
| *4x M3 screws, 12 mm long* | 28 |

| Component | # in Table S2 |
|---|---|
| *Motor hub* | Set 1 component 1 |

| Tool | # in Table S1 |
|---|---|
| *Allen key for M3* | 48 |

#### 7. Mount the motor hub on the aluminium profiles

Follow instructions in video [7. Mounting the motor hub on the aluminium profiles.]

| Component | # in Table S1 |
|---|---|
| *The motor hub assembled in step 6* | - |
| *1x T2020 black aluminium profile 40 cm long* | 20 |
| *2x T2020 black aluminium profile 8 cm long* | 20 |
| *12x M5 8 mm screws, button head, hex socket, black* | 33 |
| *12x M5 T-nuts for T2020 aluminium profile* | 37 |

| Tool | # in Table S1 |
|---|---|
| *Allen key for M5* | 48 |

#### 8. Mount bearings on the bearing guides

Follow instructions in video 8. Mounting the bearings.

| Component | # in Table S1 |
|---|---|
| *3x ball bearings* | 22 |
| *6x M5 washers* | 39 |
| *3x M5 hexagonal nuts* | 38 |
| *3x M5 x 16 mm screws, button head, hex socket, black* | 35 |

| Component | # in Table S2 |
|---|---|
| *3x bearing guides* | Set 1 components 2 and 3 |

| Tool | # in Table S1 |
|---|---|
| *Allen key for M5* | 48 |

### 9. Mount the positional sensor on the bearing guide

Follow the instructions in video [9. Mounting the mini microswitch.]

| Component | # in Table S1 |
|---|---|
| *The mini microswitch soldered in the step 5* | – |
| *2x M2.5 12 mm screws, flat head countersunk, black* | 25 |
| *2x M2.5 hexagonal nuts* | 27 |
| *The bearing guide assembled in the step 8* | – |

| Component | # in Table S2 |
|---|---|
| *Mini microswitch housing* | Set 1 component 4 |

| Tool | # in Table S1 |
|---|---|
| *Screwdriver for M2.5* | 47 |

#### 10. Mount bearing guides on the aluminium profiles and install the slot cover

Follow instructions in video 10a. Mounting bearing guides and video 10b. Aluminium profile slot cover.

| Component | # in Table S1 |
|---|---|
| *Bearing guides assembled in steps 8 and 9* | – |
| *9x M5 8 mm screws, button head, hex socket, black* | 33 |
| *9x M5 T-nuts for T2020 aluminium profile* | 37 |
| *2020 Aluminium Profile Slot Cover black, ca 1 m* | 21 |

| Tool | # in Table S1 |
|---|---|
| *Allen key for M5* | 48 |

#### 11. Mount the aluminium rail for the LED frame

Follow instructions in video 11. Mounting aluminium rail for LED.

| Component | # in Table S1 |
|---|---|
| *2x Aluminium profiles T2020, black, 14 cm long* | 20 |
| *M5 16mm bolt, hex head* | 35 |
| *6x M5 8 mm screws, buttonhead, black* | 33 |
| *3x M5 T-nuts* | 37 |
| *2x M5 slide-in brackets for T2020 aluminium profile* | 40 |

| Component | # in Table S2 |
|---|---|
| *Holder for the LED frame rail* | Set 1 component 7 |
| *1x bottom for a thumb screw* | Set 1 component 8 |
| *1x cap for a thumb screw* | Set 1 component 9 |

| Tool | # in Table S1 |
|---|---|
| *Allen key for M5* | 48 |

Optional: it is advisable to use slots cover on the bottom slot of the profile to enhance stability of the frame.

### 12. Mount the LED frame

Follow instructions in video 12. Assembling of the LED illumination:

- Attach the LED strip to the LED frame.
- Mount the frame to the aluminium rail.

| Component | # in Table S1 |
|---|---|
| *LED strip assembled in step 4b* | – |
| *4x M5 12 mm screws, flat head countersunk* | 32 |
| *4x M5 T-nuts* | 37 |
| *Super glue* | 50 |

| Component | # in Table S2 |
|---|---|
| *LED frame, right half* | Set 7 |
| *LED frame, left half* | Set 8 |
| *Diffusor, left part (L)* | Set 9 component 1 |
| *Diffusor, bottom left part (BL)* | Set 9 component 2 |
| *Diffusor, bottom right part (BR)* | Set 9 component 3 |
| *Diffusor, top part (T)* | Set 10 |
| *Diffusor, right part (R)* | Set 11 |

| Tool | # in Table S1 |
|---|---|
| *M5 Allen key* | 48 |

### 13. Mount the vertical rail

Follow instructions in video [13. Mounting the vertical rail]

| Component | # in Table S1 |
|---|---|
| *2x Aluminium profiles T2020, black, 30 cm long* | 20 |
| *1x M5 16mm bolt, hex head* | 35 |
| *1x M5 8 mm screw, button head, hex socket, black* | 33 |
| *1x M5 slide-in bracket for T2020 aluminium profile* | 40 |

| Component | # in Table S2 |
|---|---|
| *Holder for vertical rail* | Set 1 component 6 |
| *1x bottom for a thumb screw* | Set 1 component 8 |
| *1x cap for a thumb screw* | Set 1 component 9 |

| Tool | # in Table S1 |
|---|---|
| *M5 Allen key* | 48 |

### 14. Mount the light screen

Follow instructions in video 14. Mounting the light screen.

| Component | # in Table S1 |
|---|---|
| *5x M5 11 mm screws, button head, hex socket, black* | 41 |
| *5x M5 T-nuts* | 37 |

| Component | # in Table S2 |
|---|---|
| *Left part of the light screen* | Set 6 |
| *Right part of the light screen* | Set 5 |

| Tool | # in Table S1 |
|---|---|
| *M5 Allen key* | 48 |

#### 15. Assemble and mount the Borg's nest

Follow instructions in the video 15. Assembling and mounting of the Borgs nest. The pinout scheme for all components is available in **Fig. 1B**. Additionally, connections for individual components are illustrated in **Figs. 2**, **4** and **6**.

We recommend to prepare the microSD card according to the instructions in step 18 of this manual and to insert it into the Raspberry Pi before assembly.

| **Component** | **# in Table S1** |
|---|---|
| *1x Raspberry Pi 3B+* | 1 |
| *Motor driver + SparkFun power control kit + DC connector wired in steps 3 and 4* | – |
| *2x M2.5 12 mm screws, flat head countersunk* | 25 |
| *2x M2.5 12 mm screws, button head* | 24 |
| *4x M2.5 hex nuts* | 27 |
| *4x M3 12 mm screws, button head* | 28 |
| *4x M3 20 mm screws, flat head countersunk* | 29 |
| *8x M3 hex nuts* | 30 |
| *1x M5 11 mm screw, button head, black* | 41 |
| *2x M5 T-nuts* | 37 |

| **Component** | **# in Table S2** |
|---|---|
| *1x Borgs nest* | Set 2 component 1 |
| *1x Lid for the Borg's nest* | Set 2 component 2 |
| *1x DC connector lid* | Set 2 component 3 |
| *1x SparkFun holder* | Set 2 component 4 |

| **Tool** | **# in Table S1** |
|---|---|
| *M2.5 screwdriver* | 47 |
| *M3 and M5 Allen keys* | 48 |
| *Tweezers* | - |

#### 16. Mount and connect the camera

Here we provide instructions for two configurations of SPIRO that differ solely in the focusing solutions for the camera: (i) manual focus; (ii) motorized focus.

The motorized focus configuration requires a camera assembled from a Raspberry Pi v2 camera, in which the original camera module is replaced by the Arducam camera IMX219 8 MP Auto Focus camera module (**Table S1**). The solution with manual focusing requires unmodified Raspberry Pi v2 camera only (**Table S1**).

Both configurations provide the same quality of images. The motorized focus configuration is approximately 25 Euros more expensive than the configuration with the manual focus adjustment. However, the motorized option allows focus adjustment remotely from SPIRO software and thus enables considerably more convenient operation of the robot.

#### 16a. For the configuration with manual focus (alternative to 16b)

Follow instructions in the video 16. Mounting and connecting the camera.

| Component | # in Table S1 |
|---|---|
| *1x Raspberry Pi V2 Camera* | 18a |
| *1x Flex cable, ca 25 cm long* | 17 |
| *4x M2.5 8mm screws, flat head countersunk* | 26 |
| *4x M2.5 hex nuts* | 27 |
| *2x M5 16mm bolts, hex head* | 35 |
| *2x M5 T-nuts* | 37 |

| Component | # in Table S2 |
|---|---|
| *1x Housing for Pi camera* | Set 3b component 1 |
| *1x Lid for camera housing* | Set 3b component 2 |
| *2x Bottom for a thumb screw* | Set 3b component 3 |
| *2x Cap for a thumb screw* | Set 3b component 4 |
| *1x Stabilizer for the camera* | Set 3b component 5 |

| Tool | # in Table S1 |
|---|---|
| *M2.5 screwdriver* | 47 |
| *M3 Allen key* | 48 |

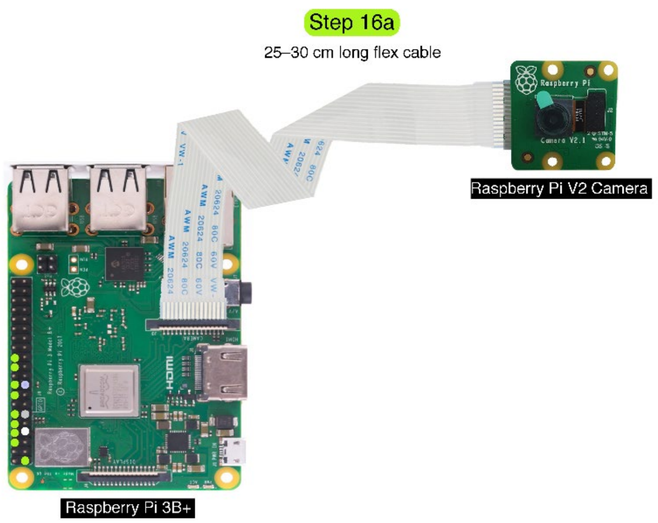

**Figure 7. Connecting the Raspberry Pi camera v2 with manual focus**
The Raspberry Pi V2 camera is connected to the camera port on the Raspberry Pi computer via a white flex cable. Please note that the metal surfaces on the flex cable should be facing the metal pins in the ports on the Raspberry Pi computer and on the camera PCB.

#### 16b. For the configuration with motorized focus (alternative to 16a)

- Replace the original module of the Raspberry Pi V2 camera with the Arducam module. Please follow the instructions provided by Arducam[4].
- Connect the camera board to the Raspberry Pi computer. The procedure is the same as for the 16a step shown in the video 16. Mounting and connecting the camera.
- Optionally, it is possible to also use printed washers placed on the thumb screws between the camera house and the aluminium profile. They increase the space left for the flex cable and thus reduce mechanical stress on it when the camera is moved along the vertical axis.

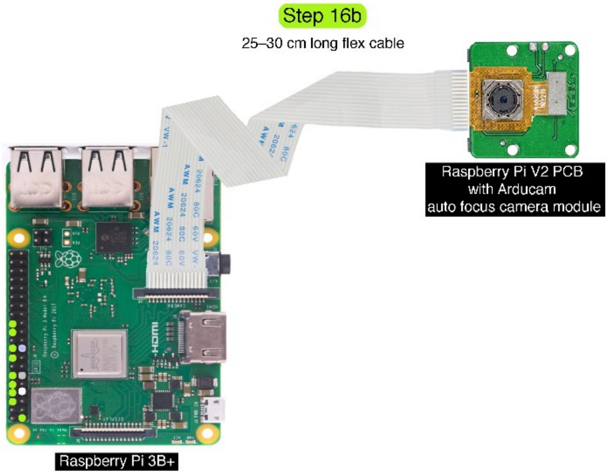

**Figure 8. Connecting the camera with the Arducam motorized camera module.** Firstly, the original Raspberry Pi V2 camera module is replaced by the Arducam module. Secondly, the Raspberry Pi V2 camera board is connected to the camera port on the Raspberry Pi computer via a white flex cable. Please note that the metal surfaces on the flex cable should be facing the metal pins in the ports on the Raspberry Pi computer and on the camera PCB.

| Component | # in Table S1 |
|---|---|
| *1x Raspberry Pi V2 Camera* | 18b |
| *1x Arducam IMX219 8MP Auto Focus Camera Module* | 18b |
| *1x Flex cable, ca 25 cm long* | 17 |
| *4x M2.5 8mm screws, flat head countersunk* | 26 |
| *4x M2.5 hex nuts* | 27 |
| *2x M5 16mm bolts, hex head* | 35 |
| *2x M5 T-nuts* | 37 |

| Component | # in Table S2 |
|---|---|
| *1x Housing for Arducam camera module* | Set 3a component 1 |
| *1x Lid for camera housing* | Set 3a component 2 |
| *2x Cap for a thumb screw* | Set 3a component 3 |
| *2x Bottom for a thumb screw* | Set 3a component 4 |
| *2x washers* | Set 3a component 5 |

| Tool | # in Table S1 |
|---|---|
| *M2.5 screwdriver or allen key* | 47 |

### 17. Assemble and mount the stage

Follow instructions in video **17. Assembling and mounting of the cube**. While mounting the shaft coupling, make sure that more than half of it is covering the motor shaft. It is of utmost importance to fix the screws on the coupling as tightly as possible to prevent unwanted wobbling during the stage rotation.

It is possible to prevent the vibrations of the stepper motor from resonating inside the cube. For this, prepare a washer made out of Parafilm (four layers of 2x2cm Parafilm squares suffices) to be fitted between the hex head of the M5 bolt and the cube bottom. It is crucial to ensure that the hex head of M5 is completely sunken into the well designed for it (not shown in the video).

Optionally, the bolt can be additionally secured in its position with a washer and a screw nut (M5) fastened under the cube.

| **Component** | **# in Table S1** |
|---|---|
| *1x Rigid shaft coupling M5 to M5* | 23 |
| *2x M2.5 8mm screws, flat head countersunk* | 26 |
| *2x M2.5 hex nuts* | 27 |
| *1x M5 20 mm bolt, hex head* | 36 |

| **Component** | **# in Table S2** |
|---|---|
| *1x Cube* | Set 4 |
| *1x Mini microswitch pin (tiny dick)* | Set 1 component 5 |

| **Tool** | **# in Table S1** |
|---|---|
| *M2.5 screwdriver* | 47 |

SPIRO assembly is complete!

#### 18. Software installation

SPIRO software should be installed on a formatted microSD card that is then placed into a slot on the Raspeberry Pi board (**Fig. 9**). For installation, please follow the instructions provided in the SPIRO Software Repository[5]. The acquired data will be stored on the same card, thus the capacity of the microSD card will determine how many images you can store. As a rule of thumb, one image will take up 8 MB. A 64 GB card costs around €10 at the time of writing this manual and can hold approximately 8000 images. Choose a well-known brand, as these cards are more reliable. Cards rated *A*, e.g. A1, are the best choice for performance.

Please note, that for completing the installation you will need to temporarily connect a keyboard to the USB port (**Fig. 9A**) and a monitor to the HDMI port (**Fig. 9**) of the Raspberry Pi computer.

##### Installing from the Supplementary File

**This step is only needed if the software repository is not accessible!** In case the SPIRO Software Repository is not available, installation from the provided Supplementary File is possible. This file also contains the installation instructions in the file README.md. In this case, the supplementary file must be transferred to the Raspberry Pi. This is most easily accomplished using an SFTP client, such as MobaXterm (Windows only) or FileZilla (Windows/Linux/Mac). After transferring the zip file (`spiro-software.zip`), enter the following command to install the SPIRO software and its dependencies:

```
sudo pip3 install spiro-software.zip
```

This command replaces the corresponding command stated in the online installation instructions (starting with `sudo pip3 install`).

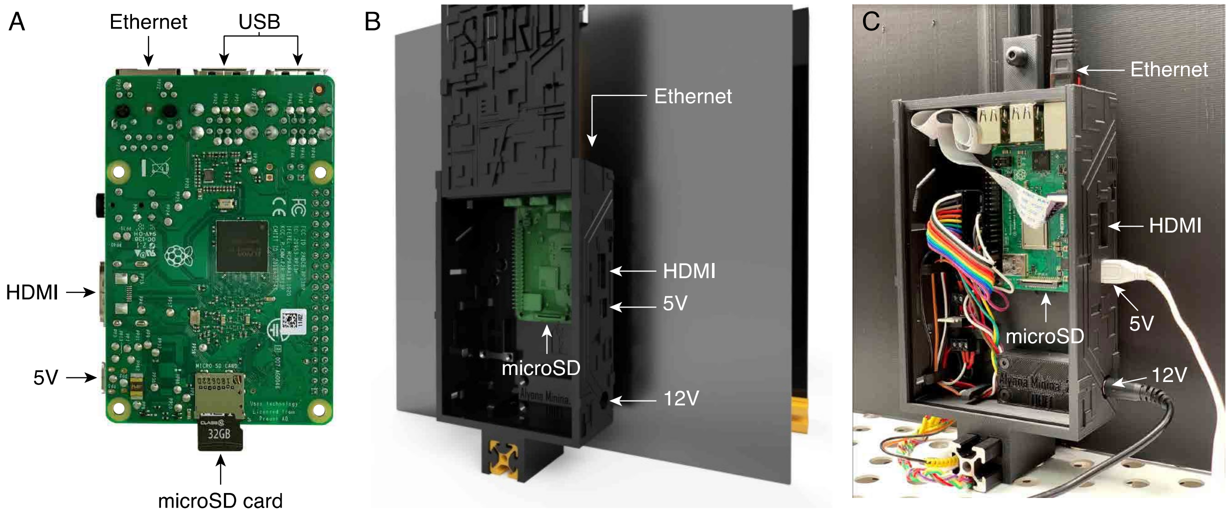

**Figure 9. Connecting power and network cables and placing the microSD card**
(**A**) Bottom view of the Raspberry Pi 3B+ with indicated slot for microSD card and ports for Ethernet, power and HDMI cables. (**B**) Rendering of assembled SPIRO showing placement of microSD card and ports for connecting power, Ethernet and HDMI cables. (**C**) Photo of assembled SPIRO showing the components indicated on (**A**) and (**B**).

| **Component** | **# in Table S1** |
|---|---|
| *1x microSD card* | 12 |
| *1x Ethernet cable* | 51 |

| **Tool** |
|---|
| *A computer with microSD card reader* |
| *A monitor with an HDMI cable* |
| *USB keyboard* |
| *Internet access, either via WiFi or ethernet* |

#### 19. Network setup

The SPIRO software is primarily controlled over the network using a web interface. Three basic modes of network connectivity are possible:

1. connecting to an existing network via an Ethernet cable
2. connecting to an existing network via Wi-Fi
3. acting as a Wi-Fi hotspot without connecting to an existing network.

The choice of network connection mode depends on the local network settings, such as firewall or other network policies and availability of network connections in the facility where SPIRO is physically deployed.

Connection modes 1 and 2 are the most flexible, in that they allow for online updating of the software and for remotely interacting with the software. However, local network policies may not allow for the use of these modes. For accessing the SPIRO software, TCP ports 22 and 8080 need to be reachable from the client. Check with your network administrator whether this is the case.

Please note, that by default the Raspberry Pi computer will be assigned a dynamic IP address. This means that every time the robot is disconnected and reconnected to the network, the address may change and you would need to physically plug a screen and keyboard into the SPIRO HDMI and USB ports to find out its current address. Therefore, we strongly recommend to request a static IP (or a dynamic hostname) for your SPIRO from your local network administrator.

Connection mode 3, hotspot mode, allows for interacting with the system without a accessing an existing network. Instead, the user connects to the Wi-Fi network named *SPIRO-XXXXXX* (where XXXXXX is a pseudorandom identifier). After connecting to this Wi-Fi network, the SPIRO system can be found at the convenience address *spiro.local* (i.e., the web interface may be reached at *http://spiro.local:8080*). The Wi-Fi hotspot does not route any traffic, and thus does not provide access to any other internet resources.

Wi-Fi hotspot mode can be enabled at the same time as the system is connected to an existing network. This option may be desirable if there is a network connection available, but firewall policies restrict access to the system. In this case, the network connection can be used for system updates, whereas the hotspot connection is used for interacting with the software. Hotspot mode is enabled by following the instruction in the online installation instructions.

### Links

1. https://www.youtube.com/watch?v=Qps9woUGkvI&ab_channel=oneTesla.
2. https://github.com/AlyonaMinina/SPIRO.Hardware.
3. https://www.prusa3d.com/prusaslicer/.
4. https://youtu.be/Zy1FYLadOfA.
5. https://github.com/jonasoh/spiro.
