## Supplementary File 3 for "SPIRO – the automated Petri plate imaging platform designed by biologists, for biologists": LICENSE.txt

Copyright (c) 2019, Jonas Ohlsson < jonas.ohlsson .at. slu.se >
All rights reserved.

Redistribution and use in source and binary forms, with or without
modification, are permitted provided that the following conditions are met:

1. Redistributions of source code must retain the above copyright notice, this
   list of conditions and the following disclaimer.
2. Redistributions in binary form must reproduce the above copyright notice,
   this list of conditions and the following disclaimer in the documentation
   and/or other materials provided with the distribution.

THIS SOFTWARE IS PROVIDED BY THE COPYRIGHT HOLDERS AND CONTRIBUTORS "AS IS" AND
ANY EXPRESS OR IMPLIED WARRANTIES, INCLUDING, BUT NOT LIMITED TO, THE IMPLIED
WARRANTIES OF MERCHANTABILITY AND FITNESS FOR A PARTICULAR PURPOSE ARE
DISCLAIMED. IN NO EVENT SHALL THE COPYRIGHT OWNER OR CONTRIBUTORS BE LIABLE FOR
ANY DIRECT, INDIRECT, INCIDENTAL, SPECIAL, EXEMPLARY, OR CONSEQUENTIAL DAMAGES
(INCLUDING, BUT NOT LIMITED TO, PROCUREMENT OF SUBSTITUTE GOODS OR SERVICES;
LOSS OF USE, DATA, OR PROFITS; OR BUSINESS INTERRUPTION) HOWEVER CAUSED AND
ON ANY THEORY OF LIABILITY, WHETHER IN CONTRACT, STRICT LIABILITY, OR TORT
(INCLUDING NEGLIGENCE OR OTHERWISE) ARISING IN ANY WAY OUT OF THE USE OF THIS
SOFTWARE, EVEN IF ADVISED OF THE POSSIBILITY OF SUCH DAMAGE.
