## Supplementary File 3 for "SPIRO – the automated Petri plate imaging platform designed by biologists, for biologists": README.md

<p align="center">
  <img src="https://user-images.githubusercontent.com/6480370/61460733-a965dd00-a96f-11e9-820a-352e4eebd1e5.png" alt="SPIRO logo">
</p>

SPIRO is an imaging platform designed for making highly reproducible, high temporal resolution timelapse sequences of biological samples grown on Petri dishes: plant seedlings, fungal mycelium, bacterial colonies, etc. SPIRO supports imaging of up to four plates at the same time. It is designed to be suitable for the two most poular Petri dish formats: round 9 cm plates and square 12 cm plates.

The image below is a link to a YouTube video showing some of its design and features.

[![SPIRO intro](https://user-images.githubusercontent.com/6480370/60589568-1e46ed80-9d9a-11e9-96ae-08fe85d8b415.png)](http://www.youtube.com/watch?v=fh5NMvDNjNc "SPIRO intro")

## Table of Contents

* [Hardware](#hardware)
* [Examples](#example-images)
* [3D printer models](#3d-printer-models)
* [Automated data analysis](#automated-data-analysis)
* [Installation](#installation)
  * [Enabling the Wi-Fi hotspot](#enabling-the-wi-fi-hotspot)
* [Usage](#usage)
  * [Working with SPIRO](#working-with-spiro)
  * [Connecting to the web interface](#connecting-to-the-web-interface)
  * [Setting up imaging](#setting-up-imaging)
  * [Starting an experiment](#starting-an-experiment)
  * [Downloading images](#downloading-images)
* [Maintaining the system](#maintaining-the-system)
  * [Restarting the software](#restarting-the-software)
  * [Shutting down the system](#shutting-down-the-system)
  * [Keeping software up to date](#keeping-software-up-to-date)
* [Troubleshooting](#troubleshooting)
  * [Viewing the software log](#viewing-the-software-log)
  * [Testing the LED and motor](#testing-the-led-and-motor)
* [Licensing](#licensing)

## Hardware

SPIRO is based around [Raspberry Pi 3 B+](https://www.raspberrypi.org/products/raspberry-pi-3-model-b-plus/), using 3D printed parts to hold everything together. It works with both the [official camera module](https://www.raspberrypi.org/products/camera-module-v2/) as well as with third-party cameras. We use the [Arducam Motorized Focus camera](http://www.arducam.com/programmable-motorized-focus-camera-raspberry-pi/), which allows focusing the images via the built-in live view web server.

The system basically consists of a camera, a green LED illuminator for imaging in the dark, and a motor-controlled imaging stage, as shown below. 

![SPIRO close-up](https://user-images.githubusercontent.com/6480370/60957134-6a4ae280-a304-11e9-8a03-0d854267297b.jpeg)

It is relatively cheap and easy to assemble multiple systems for running larger experiments.

![Multiple SPIROs](https://user-images.githubusercontent.com/6480370/60957494-0bd23400-a305-11e9-85d2-895cf8936120.jpeg)

## Example images

Here are a couple of examples showcasing the image quality of day and night images. Click for full resolution.

![Example day image](https://user-images.githubusercontent.com/6480370/77849203-2ff2df80-71ca-11ea-9285-0ebd909cd601.png)

![Example night image](https://user-images.githubusercontent.com/6480370/77849217-4b5dea80-71ca-11ea-8270-8f839e69e6f8.png)

Below is a timelapse sequence of *Arabidopsis* seedlings growing on agar. The image has been heavily downscaled due to size constraints.

<p align="center">
  <img src="https://user-images.githubusercontent.com/6480370/60673063-350f4200-9e77-11e9-9fad-f9ec3140b05c.gif" alt="Timelapse example">
</p>

## 3D printer models

Models for the 3D printed hardware components can be found at [AlyonaMinina/SPIRO.Hardware](https://github.com/AlyonaMinina/SPIRO.Hardware).

## Automated data analysis

Image data acquired using SPIRO is highly reproducible, and well suited for automated image analysis. We have so far developed pipelines for automated assessment of germination and root growth rates, which can be found at [jiaxuanleong/SPIRO.Assays](https://github.com/jiaxuanleong/SPIRO.Assays).

## Installation

First, prepare the SD card with a fresh release of Raspberry Pi OS Lite (follow the official [instructions](https://www.raspberrypi.org/documentation/installation/installing-images/README.md)). 

**Note**: If using the [ArduCam drop-in replacement camera module](https://www.arducam.com/product/arducam-imx219-auto-focus-camera-module-drop-in-replacement-for-raspberry-pi-v2-and-nvidia-jetson-nano-camera/), you need to add the following line to the file config.txt on the newly prepared SD card:
```
dtparam=i2c_vc=on
```

Connect the Raspberry Pi to a screen and keyboard. Log in using the default credentials (username `pi`, password `raspberry`). Start the system configuration:

```
sudo raspi-config
```

In the raspi-config interface, make the following changes:
* **Change the password**. The system will allow network access, and a weak password **will** compromise your network security **and** your experimental data.
* After changing the password, connect the network cable (if you are using wired networking).
* Under *Interfacing*, enable *Camera*, *I2C*, and *SSH*. 
* In the *Advanced Options*, set *Memory Split* to 256.
* Under *Localisation Options*, make sure to set the *Timezone*. Please note that a working network connection is required to maintain the correct date.
* If needed, configure *Network* and *Localization* options here as well. Set a *Hostname* under Network if you plan on running several SPIROs.
* Finally, select *Finish*, and choose to reboot the system when asked. 
* After reboot, the system shows a message on the screen showing its IP address ("My IP address is: *a.b.c.d*"). Make a note of this address as you will need it to access the system over the network. Make sure that your network allows access to ports 8080 on this IP address. (Alternatively, see [Enabling the Wi-Fi hotspot](#enabling-the-wifi-hotspot))

Next, make sure the system is up to date, and install the required tools (answer yes to any questions):

```
sudo apt update
sudo apt upgrade
sudo apt install python3-pip git i2c-tools wiringpi libatlas-base-dev zip
```

Then, install the SPIRO software and its dependencies:

```
sudo pip3 install git+https://github.com/jonasoh/spiro#egg=spiro
```

Finally, instruct the system to automatically run the SPIRO control software on boot:

```
spiro --install
systemctl enable --user spiro
sudo loginctl enable-linger pi
systemctl --user start spiro
```

You may now place the system in the setting in which you will be using it for your experiments.

### Enabling the Wi-Fi hotspot

For situations where the web UI cannot be used via the network, e.g. if using the system is used where a network connection is not available or where firewall policies block access to the web UI or SSH, SPIRO can be configured to act as a Wi-Fi hotspot. In this mode, a Wi-Fi network with the name spiro-XXXXXX is provided, which provides access only to SPIRO.

To enable the hotspot, run the following command from the terminal or via SSH (note that the system must be connected to a network when initially enabling the hotspot, as it needs to be able to download several software packages):

```
sudo spiro --enable-hotspot
```

After installing and configuring the required services, the details for connecting to the hotspot are given:

```
Access point configured and enabled. Below are the details for connecting to it:

SSID:     spiro-feed13
Password: 517824ee

Connect to the web interface using the address http://spiro.local:8080
```

To reach the web UI or SSH when connected to the hotspot, use the convenience address `spiro.local` and the normal ports (22 for SSH, 8080 for web UI). Please note that the hotspot only allows access to the SPIRO system, and otherwise provides no internet connectivity.

The hotspot can be disabled using the command:

```
sudo spiro --disable-hotspot
```

## Usage

### Working with SPIRO

Initiating experiments and controlling imaging parameters is performed using a web-based UI which can be controlled using any web browser. The screenshot shows the main view of an idle system:

![Web UI overview](https://user-images.githubusercontent.com/6480370/78147448-d0980800-7433-11ea-9bdc-13d733346f59.png)

The live view allows correcting the focus of the lens as well as placement of Petri plates and distance of the illuminator for optimal image quality. The UI also allows for calibration of the motor and setting up day and night imaging parameters (shutter speed and ISO) separately.

![Experiment control](https://user-images.githubusercontent.com/6480370/78147828-561bb800-7434-11ea-8f5f-105af09fa559.png)

In the experiment control view, parameters such as the duration of the experiment and frequency of imaging is configured. The system dynamically estimates the required size of the images, given the configured parameters.

![Running experiment](https://user-images.githubusercontent.com/6480370/78148080-b3b00480-7434-11ea-9c6c-fd01ceb1d394.png)

During a running experiment, most functions of the web UI are disabled. Browsing to the UI instead displays the status of the running experiment, and provides details on time remaining and the current image quality.

For administration of SPIRO (e.g., for updating the software and diagnosing problems), an SSH client is very useful. Below are a few choices:

**Windows**
* [MobaXterm](https://mobaxterm.mobatek.net/) is a popular SSH client that also supports file transfer. Recommended for beginners. 
* [PuTTY](https://www.chiark.greenend.org.uk/~sgtatham/putty/latest.html) is a lightweight and popular SSH client.

**Mac/Linux**
* On Mac and Linux, an SSH client is built into the system. Using the *Terminal*, connect to SPIRO using the command `ssh pi@1.2.3.4` (where `1.2.3.4` is the IP address of your system).

### Connecting to the web interface

SPIRO is controlled via its web interface. To access the system, point your web browser to the address *http://**a.b.c.d:8080**/*, where *a.b.c.d* is the IP address you noted previously (the server always runs on port 8080). At the first access, you are asked to set a password for the system. **Do not use the same password as you set previously!** This password is for the web interface and should be regarded as a low-privilege password.

### Setting up imaging

After logging in to the system, you are presented with the *Live view*. Here, you can adjust the image in real time, allowing you to make sure that the camera is at a proper distance from the plate, that the focus is set correctly, and that the LED illuminator is working and placed in a position where reflections are not an issue.

Under *Day image settings* and *Night image settings*, you can adjust the exposure time for day and night images in real time. For night images, make sure that representative conditions are used (i.e., turn off the lights in the growth chamber). When the image is captured according to your liking, choose *Update and save*.

For locating the initial imaging position, the system turns the cube until a positional switch is activated. It then turns the cube a predefined amount of steps (*calibration value*). The check that the calibration value is correct, go to the *Start position calibration* view. Here, you may try out the current value, as well as change it. Make sure to click *Save value* if you change it.

### Starting an experiment

When imaging parameters are set up to your liking, you are ready to start your experiments. In the *Experiment control* view, choose a name for your experiment, as well as the duration and imaging frequency. After you choose *Start experiment*, the system will disable most of the functionality of the web interface, displaying a simple status window containing experiment parameters, as well as the last image captured.

### Downloading images

Images can be downloaded from the web interface under *File manager*. The File manager also allows deleting files to free up space on the SD card.

## Maintaining the system

### Restarting the software

If for some reason the software ends up in an unusable state, you can often restart it from the web UI (**System settings -> Restart web UI**). The web UI can also be restarted via SSH, by issuing the following command:

```
systemctl --user restart spiro
```

It has happened that the camera has become unresponsive, due to bugs in the underlying library. In this case, rebooting the system is the only way of getting it back into a usable state. This can be done via the web UI (**System settings -> Reboot system**), or by issuing the following command via SSH:

```
sudo shutdown -r now
```

### Shutting down the system

If you want to power down the system, always perform a clean shutdown to ensure that no damage is caused to the filsesystem. This can either be done via the web UI (**System settings -> Power off system**), or by issuing the following command over SSH:

```
sudo shutdown -h now
```

After a few seconds, when only the red LED on the Raspberry Pi is lit, you may pull the power.

### Keeping software up to date

You should regularly update the underlying operating system to make sure that it has the latest security patches. To do so, log in using your SSH client (or connect a screen and keyboard), and issue the following commands, answering *Y* to any queries:

```
sudo apt update
sudo apt upgrade
```

To update the SPIRO control software, issue the commands:

```
sudo pip3 install -U git+https://github.com/jonasoh/spiro#egg=spiro
```

After updating the SPIRO software, either use the web UI (**System settings -> Restart web UI**), or issue the following command in an SSH prompt:

```
systemctl --user restart spiro
```

## Troubleshooting

### Viewing the software log

To view the last 50 entries in the SPIRO log:

```
journalctl --user-unit=spiro -n 50
```

Or, to view the entire log:

```
journalctl --user-unit=spiro
```

### Testing the LED and motor

To check whether the Raspberry Pi can control the LED illuminator, first set the LED control pin to output mode:

```
gpio -g mode 17 out
```

You may then toggle it on and off using the command

```
gpio -g toggle 17
```

If it doesn't respond to this command, this may indicate either miswiring, or that either the LED strip or the MOSFET is non-functional.

Similarly, you can turn on and off the motor:

```
gpio -g mode 23 out
gpio -g toggle 23
```

When GPIO pin 23 is toggled on, the cube should be locked in position. If it is not, check that your wiring looks good, that the power supply is connected, and that the shaft coupler is firmly attached to both the cube and the motor. 

If the motor is moving jerkily during normal operation, there is likely a problem with the wiring of the coil pins (Ain1&2 and Bin1&2).

## Licensing

The code for SPIRO is licensed under a 2-clause BSD license, allowing redistribution and modification of the source code as long as the original license and copyright notice are retained. SPIRO includes the fonts [Aldrich](https://fonts.google.com/specimen/Aldrich) and [Saira Condensed](https://github.com/Omnibus-Type/Saira), for which the licenses can be found under the `doc/licenses` directory.
