## Supplementary File 3 for "SPIRO – the automated Petri plate imaging platform designed by biologists, for biologists": setup.py

```
from setuptools import setup, find_packages
setup(name = 'spiro',
      version = '1.0',
      packages = find_packages(),
      scripts = ['bin/spiro'],
      install_requires = ['picamera==1.13', 'RPi.GPIO==0.6.5', 'Flask==1.1.1', 'waitress==1.4.3', 'numpy==1.18.1'],
      author = 'Jonas Ohlsson',
      author_email = '',
      description = 'Control software for the SPIRO biological imaging system',
      url = 'https://github.com/jonasoh/spiro',
      include_package_data = True,
      zip_safe = False,
      )
```
