## Supplementary File 3 for "SPIRO – the automated Petri plate imaging platform designed by biologists, for biologists": config.py

```
import json
import os
import sys

def log(msg):
    sys.stderr.write(msg + '\n')
    sys.stderr.flush()


class Config(object):
    defaults = {
        'calibration' : 8,   # number of steps taken after positional sensor is lit
        'sensor': 4,         # pin for mini microswitch positional sensor
        'LED': 17,           # pin for turning on/off led
        'PWMa': 8,           # first pwm pin
        'PWMb': 14,          # second pwm pin
        'coilpin_M11': 25,   # ain2 pin
        'coilpin_M12': 24,   # ain1 pin
        'coilpin_M21': 18,   # bin1 pin
        'coilpin_M22': 15,   # bin2 pin
        'stdby': 23,         # stby pin
        'focus': 250,	     # default focus distance
        'password': '',      # an empty password will trigger password initialization for web ui
        'secret': '',        # secret key for flask sessions
        'dayshutter': 100,   # day exposure time in fractions of a second, e.g. 100 means 1/100
        'dayiso': 50,        # daytime iso values
        'nightshutter': 10,  # night exposure time
        'nightiso': 400,     # night iso
        'name': 'spiro',     # the name of this spiro instance
        'debug': False,      # debug logging
    }

    config = {}

    def __init__(self):
        self.cfgdir = os.path.expanduser("~/.config/spiro")
        self.cfgfile = os.path.join(self.cfgdir, "spiro.conf")
        self.read()
        if os.path.exists(self.cfgfile):
            st = os.stat(self.cfgfile)
            self.mtime = st.st_mtime
        else:
            self.mtime = 0


    def read(self):
        os.makedirs(self.cfgdir, exist_ok=True)
        if os.path.exists(self.cfgfile):
            try:
                with open(self.cfgfile, 'r') as f:
                    self.config = json.load(f)
            except Exception as e:
                log("Failed to read or parse config file: " + str(e))


    def write(self):
        try:
            with open(self.cfgfile + ".tmp", 'w') as f:
                json.dump(self.config, f, indent=4)
            os.replace(self.cfgfile + ".tmp", self.cfgfile)
        except OSError as e:
            log("Failed to write config file: " + e.strerror)

    def get(self, key):
        if os.path.exists(self.cfgfile):
            st = os.stat(self.cfgfile)
            mt = st.st_mtime
            if mt > self.mtime:
                # config file was changed on disk -- reload it
                self.read()

        return self.config.get(key, self.defaults.get(key))


    def set(self, key, value):
        self.config[key] = value
        self.write()


    def unset(self, key):
        if key in self.config:
            del self.config[key]
            self.write()
```
