## Supplementary File 3 for "SPIRO – the automated Petri plate imaging platform designed by biologists, for biologists": experimenter.py

```
import os
import time
import threading
import numpy as np
from datetime import date
from statistics import mean
from collections import deque
from spiro.config import Config
from spiro.logger import log, debug

class Experimenter(threading.Thread):
    def __init__(self, hw=None, cam=None):
        self.hw = hw
        self.cam = cam
        self.cfg = Config()
        self.delay = 60
        self.duration = 7
        self.dir = os.path.expanduser('~')
        self.starttime = 0
        self.endtime = 0
        self.running = False
        self.status = "Stopped"
        self.daytime = "TBD"
        self.quit = False
        self.stop_experiment = False
        self.status_change = threading.Event()
        self.next_status = ''
        self.last_captured = None
        self.nshots = 0
        self.idlepos = 0
        threading.Thread.__init__(self)


    def stop(self):
        self.status = "Stopping"
        self.next_status = ''
        self.stop_experiment = True
        log("Stopping running experiment...")


    def getDefName(self):
        '''returns a default experiment name'''
        today = date.today().strftime('%Y.%m.%d')
        return today + ' ' + self.cfg.get('name')


    def isDaytime(self):
        '''algorithm for daytime estimation.
           if the average pixel intensity is less than 10, we assume it is night.
           this may be tweaked for special use cases.'''
        oldres = self.cam.resolution
        self.cam.resolution = (320, 240)
        self.cam.iso = self.cfg.get('dayiso')
        self.cam.shutter_speed = 1000000 // self.cfg.get('dayshutter')
        output = np.empty((240, 320, 3), dtype=np.uint8)
        self.cam.capture(output, 'rgb')
        self.cam.resolution = oldres
        debug("Daytime estimation mean value: " + str(output.mean()))
        return output.mean() > 10


    def setWB(self):
        debug("Determining white balance.")
        self.cam.awb_mode = "auto"
        time.sleep(2)
        g = self.cam.awb_gains
        self.cam.awb_mode = "off"
        self.cam.awb_gains = g


    def takePicture(self, name):
        filename = ""
        prev_daytime = self.daytime
        self.daytime = self.isDaytime()
        
        if self.daytime:
            time.sleep(0.5)
            self.cam.shutter_speed = 1000000 // self.cfg.get('dayshutter')
            self.cam.iso = self.cfg.get('dayiso')
            self.cam.color_effects = None
            filename = os.path.join(self.dir, name + "-day.png")
        else:
            # turn on led
            self.hw.LEDControl(True)
            time.sleep(0.5)
            self.cam.shutter_speed = 1000000 // self.cfg.get('nightshutter')
            self.cam.iso = self.cfg.get('nightiso')
            filename = os.path.join(self.dir, name + "-night.png")
        
        if prev_daytime != self.daytime and self.daytime and self.cam.awb_mode != "off":
            # if there is a daytime shift, AND it is daytime, AND white balance was not previously set,
            # set the white balance to a fixed value.
            # thus, white balance will only be fixed for the first occurence of daylight.
            self.setWB()

        debug("Capturing %s." % filename)
        self.cam.exposure_mode = "off"
        self.cam.capture(filename)
        self.last_captured = filename
        self.cam.color_effects = None
        self.cam.shutter_speed = 0
        # we leave the cam in auto exposure mode to improve daytime assessment performance
        self.cam.exposure_mode = "auto"
       
        if not self.daytime:
            # turn off led
            self.hw.LEDControl(False)


    def run(self):
        '''starts experiment if there is signal to do so'''
        while not self.quit:
            self.status_change.wait()
            if self.next_status == 'run':
                self.next_status = ''
                self.status_change.clear()
                self.runExperiment()


    def go(self):
        '''signals intent to start experiment'''
        self.next_status = 'run'
        self.status_change.set()                


    def runExperiment(self):
        '''main experiment loop'''
        if self.running:
            raise RuntimeError('An experiment is already running.')

        try:
            debug("Starting experiment.")
            self.running = True
            self.status = "Initiating"
            self.starttime = time.time()
            self.endtime = time.time() + 60 * 60 * 24 * self.duration
            self.last_captured = None
            self.delay = self.delay or 0.001
            self.nshots = self.duration * 24 * 60 // self.delay
            self.cam.exposure_mode = "auto"
            self.cam.shutter_speed = 0
            self.hw.LEDControl(False)

            if self.dir == os.path.expanduser('~'):
                # make sure we don't write directly to home dir
                self.dir = os.path.join(os.path.expanduser('~'), self.getDefName())

            for i in range(4):
                platedir = "plate" + str(i + 1)
                os.makedirs(os.path.join(self.dir, platedir), exist_ok=True)

            while time.time() < self.endtime and not self.stop_experiment:
                loopstart = time.time()
                # need to use time-based loop control as we do not know how long a rotation takes
                nextloop = time.time() + 60 * self.delay
                if nextloop > self.endtime:
                    nextloop = self.endtime
                
                for i in range(4):
                    # rotate stage to starting position
                    if i == 0:
                        self.hw.motorOn(True)
                        self.status = "Finding start position"
                        debug("Finding initial position.")
                        self.hw.findStart(calibration=self.cfg.get('calibration'))
                        debug("Found initial position.")
                    else:
                        self.status = "Imaging"
                        # rotate cube 90 degrees
                        debug("Rotating stage.")
                        self.hw.halfStep(100, 0.03)

                    # wait for the cube to stabilize
                    time.sleep(0.5)

                    now = time.strftime("%Y%m%d-%H%M%S", time.localtime())
                    name = os.path.join("plate" + str(i + 1), "plate" + str(i + 1) + "-" + now)
                    self.takePicture(name)

                self.nshots -= 1
                self.hw.motorOn(False)
                self.status = "Waiting"

                if self.idlepos > 0:
                    # alternate between resting positions during idle, stepping 45 degrees per image
                    self.hw.motorOn(True)
                    self.hw.halfStep(50 * self.idlepos, 0.03)
                    self.hw.motorOn(False)

                self.idlepos += 1
                if self.idlepos > 7:
                    self.idlepos = 0

                while time.time() < nextloop and not self.stop_experiment:
                    time.sleep(1)

        finally:
            log("Experiment stopped.")
            self.cam.color_effects = None
            self.status = "Stopped"
            self.stop_experiment = False
            self.running = False
            self.cam.exposure_mode = "auto"
            self.cam.meter_mode = 'spot'
```
