## Supplementary File 3 for "SPIRO – the automated Petri plate imaging platform designed by biologists, for biologists": hostapd.py

```
#!/usr/bin/env python3
#
# hostapd.py -
#   configure system as an AP directing all client queries to the web ui
#

import subprocess
import textwrap
import uuid
from spiro.logger import log, debug


def init():
    """makes sure everything runs smoothly later on."""
    p = subprocess.run(["systemctl", "daemon-reload"], stderr=subprocess.STDOUT, stdout=subprocess.PIPE)


def install_reqs():
    """installs hostapd and dnsmasq. returns a tuple of (installation status, output of failed command)."""
    reqs = ['dnsmasq', 'hostapd']
    for r in reqs:
        p = subprocess.run(['dpkg', '-l', r], stdout=subprocess.PIPE, stderr=subprocess.STDOUT)
        if p.returncode != 0:
            # install requirement
            p = subprocess.run(["apt", "install", "-y", r], stderr=subprocess.STDOUT, stdout=subprocess.PIPE)
            if p.returncode != 0:
                # installation failed
                return (False, p.stdout)
            # stop the service as it is not properly configured yet
            p = subprocess.run(["systemctl", "stop", r], stderr=subprocess.STDOUT, stdout=subprocess.PIPE)

    return (True, None)


def config_hostapd():
    """sets up hostapd config. returns a tuple of (network name, password)."""
    # use last 6 digits of MAC address as unique id
    u = str(uuid.uuid1())
    id = "spiro-" + u[30:36]
    pwd = u[0:8]

    with open("/etc/hostapd/hostapd.conf", "w") as f:
        f.write(textwrap.dedent("""\
                                # auto-generated by spiro software
                                interface=wlan0
                                driver=nl80211
                                ssid={0}
                                hw_mode=g
                                channel=7
                                wmm_enabled=0
                                macaddr_acl=0
                                auth_algs=1
                                ignore_broadcast_ssid=0
                                wpa=2
                                wpa_passphrase={1}
                                wpa_key_mgmt=WPA-PSK
                                wpa_pairwise=TKIP
                                rsn_pairwise=CCMP
                                """.format(id, pwd)))
    return(id, pwd)


def config_dnsmasq():
    """sets up dnsmasq config. all dns queries are forwarded to the local ip (192.168.138.1)"""
    with open("/etc/dnsmasq.conf", "w") as f:
        f.write(textwrap.dedent("""\
                                # auto-generated by spiro software
                                interface=wlan0
                                dhcp-range=192.168.138.10,192.168.138.254,12h
                                address=/#/192.168.138.1
                                no-resolv
                                no-poll
                                no-hosts
                                domain=spiro.local
                                resolv-file=/dev/null
                                """))
    with open("/etc/default/dnsmasq", "w") as f:
        f.write(textwrap.dedent("""\
                                ENABLED=1
                                DNSMASQ_EXCEPT=lo
                                """))


def config_dhcpcd(enable):
    """enables/disables static ip for the wlan0 interface. replaces the system dhcpcd.conf with our own."""
    with open("/etc/dhcpcd.conf", "w") as f:
        f.write(textwrap.dedent("""\
                                # auto-generated by spiro software
                                hostname
                                clientid
                                persistent
                                option rapid_commit
                                option domain_name_servers, domain_name, domain_search, host_name
                                option classless_static_routes
                                option interface_mtu
                                option ntp_servers
                                require dhcp_server_identifier
                                slaac private
                                """))
        if enable == True:
            f.write(textwrap.dedent("""\
                                    interface wlan0
                                    static ip_address=192.168.138.1/24
                                    nohook wpa_supplicant
                                    """))


def restart_services():
    """restarts the required services after config updates. returns False if any restart command returns
    a non-zero exit code."""
    services = ['dhcpcd', 'dnsmasq', 'hostapd']
    codes = []

    for s in services:
        p = subprocess.run(["systemctl", "restart", s], stderr=subprocess.STDOUT, stdout=subprocess.PIPE)
        debug("Restarted service {0} with status code {1}.".format(s, p.returncode))
        codes.append(p.returncode)

    return(all(c == 0 for c in codes))


def enable_services():
    # make sure wlan interface is not blocked
    p = subprocess.run(['rfkill', 'unblock', 'wlan'], capture_output=True)

    services = ['dnsmasq', 'hostapd']
    for s in services:
        p = subprocess.run(['systemctl', 'unmask', s], capture_output=True)
        p = subprocess.run(['systemctl', 'enable', s], capture_output=True)
        

def disable_services():
    services = ['dnsmasq', 'hostapd']
    for s in services:
        p = subprocess.run(['systemctl', 'stop', s], capture_output=True)
        p = subprocess.run(['systemctl', 'disable', s], capture_output=True)


def start_ap():
    log("Setting up dependencies...")
    init()
    install_reqs()
    log("Configuring system...")
    config_dhcpcd(enable=True)
    config_dnsmasq()
    ssid, pwd = config_hostapd()
    log("Starting services...")
    enable_services()
    r = restart_services()
    if not r:
        log("Failed to restart services.")
        log("Setting up access point failed.")
        return(1)
    else:
        log("Access point configured and enabled. Below are the details for connecting to it:")
        log("\nSSID:     " + ssid)
        log("Password: " + pwd)
        log("\nConnect to the web interface using the address http://spiro.local")
        return(0)


def stop_ap():
    log("Disabling services...")
    init()
    config_dhcpcd(enable=False)
    p = subprocess.run(['systemctl', 'restart', 'dhcpcd'], capture_output=True)
    disable_services()
    log("Access point disabled.")
```
