## Supplementary File 3 for "SPIRO – the automated Petri plate imaging platform designed by biologists, for biologists": hwcontrol.py

```
import RPi.GPIO as gpio
import time
import os
from spiro.config import Config
from spiro.logger import log, debug

class HWControl:
    def __init__(self):
        gpio.setmode(gpio.BCM)
        self.cfg = Config()
        self.pins = {
            'LED' : self.cfg.get('LED'),
            'sensor' : self.cfg.get('sensor'),
            'PWMa' : self.cfg.get('PWMa'),
            'PWMb' : self.cfg.get('PWMb'),
            'coilpin_M11' : self.cfg.get('coilpin_M11'),
            'coilpin_M12' : self.cfg.get('coilpin_M12'),
            'coilpin_M21' : self.cfg.get('coilpin_M21'),
            'coilpin_M22' : self.cfg.get('coilpin_M22'),
            'stdby' : self.cfg.get('stdby')
        }
        self.led = False
        self.GPIOInit()


    def GPIOInit(self):
        gpio.setwarnings(False)
        gpio.setup(self.pins['LED'], gpio.OUT)
        gpio.setup(self.pins['sensor'], gpio.IN, pull_up_down=gpio.PUD_DOWN)
        gpio.setup(self.pins['PWMa'], gpio.OUT)
        gpio.setup(self.pins['PWMb'], gpio.OUT)
        gpio.setup(self.pins['coilpin_M11'], gpio.OUT)
        gpio.setup(self.pins['coilpin_M12'], gpio.OUT)
        gpio.setup(self.pins['coilpin_M21'], gpio.OUT)
        gpio.setup(self.pins['coilpin_M22'], gpio.OUT)
        gpio.setup(self.pins['stdby'], gpio.OUT)
        gpio.output(self.pins['PWMa'], True)
        gpio.output(self.pins['PWMb'], True)
        self.LEDControl(False)
        self.motorOn(False)


    def cleanup(self):
        gpio.cleanup()


    def findStart(self, calibration=None):
        """rotates the imaging stage until the positional switch is activated"""
        calibration = calibration or self.cfg.get('calibration')
        timeout = 60
        starttime = time.time()

        # make sure that switch is not depressed when starting
        if gpio.input(self.pins['sensor']):
            while gpio.input(self.pins['sensor']) and time.time() < starttime + timeout:
                self.halfStep(1, 0.03)

        while not gpio.input(self.pins['sensor']) and time.time() < starttime + timeout:
            self.halfStep(1, 0.03)

        if time.time() < starttime + timeout:
            self.halfStep(calibration, 0.03)
        else:
            log("Timed out while finding start position! Images will be misaligned.")


    # sets the motor pins as element in sequence
    def setStepper(self, M_seq, i):
        gpio.output(self.pins['coilpin_M11'], M_seq[i][0])
        gpio.output(self.pins['coilpin_M12'], M_seq[i][1])
        gpio.output(self.pins['coilpin_M21'], M_seq[i][2])
        gpio.output(self.pins['coilpin_M22'], M_seq[i][3])


    # steps the stepper motor using half steps, "delay" is time between coil change
    # 400 steps is 360 degrees
    def halfStep(self, steps, delay, keep_motor_on=False):
        time.sleep(0.005) # time for motor to activate
        for i in range(0, steps):
            self.setStepper(self.halfstep_seq, self.seqNumb)
            self.seqNumb += 1
            if(self.seqNumb == 8):
                self.seqNumb = 0
            time.sleep(delay)


    # sets motor standby status
    def motorOn(self, value):
        gpio.output(self.pins['stdby'], value)


    # turns on and off led
    def LEDControl(self, value):
        gpio.output(self.pins['LED'], value)
        self.led = value


    # focuses the ArduCam motorized focus camera
    # code is from ArduCam GitHub repo
    def focusCam(self, val):
        value = (val << 4) & 0x3ff0
        data1 = (value >> 8) & 0x3f
        data2 = value & 0xf0
        if os.path.exists('/dev/i2c-0'):
            os.system("i2cset -y 0 0x0c %d %d" % (data1,data2))
        if os.path.exists('/dev/i2c-1'):
            os.system("i2cset -y 1 0x0c %d %d" % (data1,data2))


    # my copy of the pinout
    pins = {}
    
    # state of stepper motor sequence
    seqNumb = 0
    
    # sequence for one coil rotation of stepper motor using half step
    halfstep_seq = [(1,0,0,0), (1,0,1,0), (0,0,1,0), (0,1,1,0),
                    (0,1,0,0), (0,1,0,1), (0,0,0,1), (1,0,0,1)]
```
