## Supplementary File 3 for "SPIRO – the automated Petri plate imaging platform designed by biologists, for biologists": logger.py

```
import sys
from spiro.config import Config

cfg = Config()

def log(msg):
    sys.stderr.write(msg + '\n')
    sys.stderr.flush()

def debug(msg):
    if cfg.get('debug'):
        sys.stderr.write(msg + '\n')
        sys.stderr.flush()
```
