## Supplementary File 3 for "SPIRO – the automated Petri plate imaging platform designed by biologists, for biologists": spiro.py

```
#!/usr/bin/env python3
#
# spiro.py -
#   script for time-lapse imaging of Petri dishes, with a focus on plants
#   (i.e. it is adapted to day/night cycles).
#
# - Jonas Ohlsson <jonas.ohlsson .a. slu.se>
#

import os
from picamera import PiCamera
from spiro.hwcontrol import HWControl
from spiro.config import Config
from spiro.logger import log, debug
import spiro.hostapd as hostapd
import spiro.webui as webui
import argparse
import textwrap
import sys
import signal
import RPi.GPIO as gpio

parser = argparse.ArgumentParser(
             description=textwrap.dedent("""\
                                         SPIRO control software.
                                         Running this command without any flags starts the web interface.
                                         Specifying flags will perform those actions, then exit."""))
parser.add_argument('--reset-config', action="store_true", dest="reset",
                    help="reset all configuration values to defaults")
parser.add_argument('--reset-password', action="store_true", dest="resetpw",
                    help="reset web UI password")
parser.add_argument('--install-service', action="store_true", dest="install",
                    help="install systemd user service file")
parser.add_argument('--toggle-debug', action="store_true", dest="toggle_debug",
                    help="toggles additional debug logging on or off")
parser.add_argument('--enable-hotspot', action="store_true", dest="enable_ap",
                    help="enables the wi-fi hotspot")
parser.add_argument('--disable-hotspot', action="store_true", dest="disable_ap",
                    help="disables the wi-fi hotspot")
options = parser.parse_args()

def initCam():
    cam = PiCamera()
    # cam.framerate dictates longest exposure (1/cam.framerate)
    cam.framerate = 5
    cam.iso = 50
    cam.resolution = cam.MAX_RESOLUTION
    cam.rotation = 90
    cam.image_denoise = False
    hw.focusCam(cfg.get('focus'))
    return cam


def installService():
    try:
        os.makedirs(os.path.expanduser('~/.config/systemd/user'), exist_ok=True)
    except OSError as e:
        print("Could not make directory (~/.config/systemd/user):", e)
    try:
        with open(os.path.expanduser('~/.config/systemd/user/spiro.service'), 'w') as f:
            if (os.path.exists('/home/pi/.local/bin/spiro')): exe = '/home/pi/.local/bin/spiro'
            else: exe = '/usr/local/bin/spiro'
            f.write(textwrap.dedent("""\
                [Unit]
                Description=SPIRO control software
                [Service]
                ExecStart={}
                Restart=always
                [Install]
                WantedBy=default.target
                """).format(exe))
    except OSError as e:
        print("Could not write file (~/.config/systemd/user/spiro.service):", e)
    print("Systemd service file installed.")


def terminate(sig, frame):
    global shutdown
    if sig == signal.SIGALRM:
        # force shutdown
        debug("Shut down time-out, force-quitting.")
        debug("If the software locks up at this point, a reboot is needed.")
        debug("This is due to a bug in the underlying camera code.")
        cam.close()
        sys.exit()

    if not shutdown:
        # give the app 10 seconds to shut down, then force it
        shutdown = True
        signal.alarm(10)

    log("Signal " + str(sig) + " caught -- shutting down.")
    webui.stop()
    hw.motorOn(False)
    cam.close()
    hw.cleanup()
    sys.exit()


shutdown = False
cfg = Config()
cam = None
hw = HWControl()
for sig in [signal.SIGTERM, signal.SIGINT, signal.SIGQUIT, signal.SIGHUP, signal.SIGALRM]:
    signal.signal(sig, terminate)

# start here.
def main():
    if options.reset:
        print("Clearing all configuration values.")
        try:
            os.remove(os.path.expanduser('~/.config/spiro/spiro.conf'))
        except OSError as e:
            print("Could not remove file (~/.config/spiro/spiro.conf):", e.strerror)
            raise
    if options.install:
        print("Installing systemd service file.")
        installService()
    if options.resetpw:
        print("Resetting web UI password.")
        cfg.set('password', '')
    if options.toggle_debug:
        cfg.set('debug', not cfg.get('debug'))
        if cfg.get('debug'):
            print("Debug mode on.")
        else:
            print("Debug mode off")
    if options.enable_ap:
        hostapd.start_ap()
    if options.disable_ap:
        hostapd.stop_ap()
    if any([options.reset, options.resetpw, options.install, options.toggle_debug,
            options.enable_ap, options.disable_ap]):
        sys.exit()

    # no options given, go ahead and start web ui
    global cam
    gpio.setmode(gpio.BCM)
    hw.GPIOInit()
    cam = initCam()
    log('Starting web UI.')
    webui.start(cam, hw)
```
