## Supplementary File 3 for "SPIRO – the automated Petri plate imaging platform designed by biologists, for biologists": spiro.js

```
function bgGet(url) {
    var x=new XMLHttpRequest;
    x.open('GET', url);
    x.send();
    return(x.responseText);
}

function toggleLED(value) {
    if(value) {
        bgGet('/led/on');
    } else {
        bgGet('/led/off');
    }
}

function updateFocusNumber(value) {
    focusnumber = document.getElementById('focusnumber');
    focusnumber.value = value;
    bgGet('/focus/' + value);
}

function updateFocusSlider(value) {
    focusslider = document.getElementById('focusslider');
    focusslider.value = value;
    bgGet('/focus/' + value);
}

function updateShutter(value) {
    shutterrange = document.getElementById('shutterrange');
    shutternumber = document.getElementById('shutternumber');
    bgGet('/shutter/live/' + value);
    shutterrange.value = value;
    shutternumber.value = value;
}

function calcDiskSpace() {
    disk = document.getElementById('disk');
    delaydom = document.getElementById('delay');
    delay = parseFloat(delaydom.value);
    duration = document.getElementById('duration');
    duration = parseFloat(duration.value);
    req = 4 * 10 * duration * 24 * 60 / delay / 1024;
    disk.innerHTML = req.toFixed(1) + " GB";
    avail = document.getElementById('diskavail').innerHTML;
    avail = parseFloat(avail.split(" ")[0]);
    if (req > avail) {
        console.log("req > avail");
        if (!disk.classList.contains("diskfull")) {
            console.log("add diskfull class");
            disk.classList.add("diskfull");
        }
    } else {
        console.log("req < avail");
        if (disk.classList.contains("diskfull")) {
            console.log("remove diskfull class");
            disk.classList.remove("diskfull");
        }
    }
}

function tryCalibration() {
    calib = document.getElementById('calibration');
    bgGet('/findstart/' + calib.value);
}
```
