## Supplementary File 3 for "SPIRO – the automated Petri plate imaging platform designed by biologists, for biologists": calibrate.html

{% extends "layout.html" %}
{% block title %}Main view{% endblock %}
{% block content %}

{{ name }}

- Live view
- Day image settings
- Night image settings
- Calibrate motor
- Experiment control
- System settings
- File manager
- Log out

Set up start position

Try value
Save value

{% endblock %}
