## Supplementary File 3 for "SPIRO – the automated Petri plate imaging platform designed by biologists, for biologists": experiment.html

{% extends "layout.html" %}
{% block title %}Experiment view{% endblock %}
{% block content %}
{% if running %}
{% if status == "Waiting" %}
{% else %}
{% endif %}
{% endif %}

{{ name }}

{% if running %}- Live view
- Set up day image
- Set up night image
- Calibrate motor
{% else %}- Live view
- Day image settings
- Night image settings
- Calibrate motor
{% endif %}- Experiment control
- System settings
- File manager
- Log out

{% if running %}

### Experiment info

**Status:** {{ status }}

**Folder:** {{ directory }}

**Started:** {{ starttime }}

**Ends:** {{ endtime }}

**Images remaining/plate:** {{ nshots }}

**Disk required:** {{ diskreq }} GB

**Disk available:** {{ diskspace }} GB

Stop experiment

{% else %}

Start new experiment

Directory

Duration

days

Image every

minutes

Disk space required:{{ (4 \* 4 \* duration \* 24 \* 60 / delay / 1024)|round(1) }} GB

Disk space available:{{ diskspace }} GB

Start experiment

{% endif %}

{% if running %}

{% endif %}

{% endblock %}
