## Supplementary File 3 for "SPIRO – the automated Petri plate imaging platform designed by biologists, for biologists": exposure.html

{% extends "layout.html" %}
{% block title %}Exposure controls{% endblock %}
{% block content %}

{{ name }}

- Live view
{% if time == 'day' %}- Day image settings
- Night image settings
{% else %}- Day image settings
- Night image settings
{% endif %}- Calibrate motor
- Experiment control
- System settings
- File manager
- Log out

Live view

Exposure parameters

Shutter speed (1/n)


ISO
{% if time == 'day' %}
{% else %}
{% endif %}

Update and save

{% if time == 'day' %}


{% if dayshutter %}

Day image captured at 1/{{ dayshutter }} s, ISO {{ dayiso }}

{% endif %}
{% else %}


{% if nightshutter %}

Night image captured at 1/{{ nightshutter }} s, ISO {{ nightiso }}

{% endif %}
{% endif %}

{% endblock %}
