## Supplementary File 3 for "SPIRO – the automated Petri plate imaging platform designed by biologists, for biologists": filemanager.html

{% extends "layout.html" %}
{% block title %}File manager{% endblock %}
{% block content %}

{{ name }}

{% if running %}- Live view
- Set up day image
- Set up night image
- Calibrate motor
{% else %}- Live view
- Day image settings
- Night image settings
- Calibrate motor
{% endif %}- Experiment control
- System settings
- File manager
- Log out

File manager

{% for dir in dirs %}- {{ dir }} [Download] [Delete]
  {% endfor %}

---

**Disk available:** {{ diskspace }} GB

{% endblock %}
