## Supplementary File 3 for "SPIRO – the automated Petri plate imaging platform designed by biologists, for biologists": layout.html

SPIRO: {% block title %}{% endblock %}


{% with messages = get\_flashed\_messages() %}
{% if messages %}

{% for message in messages %}- {{ message }}
{% endfor %}
{% endif %}
{% endwith %}
{% block content %}{% endblock %}
