## Supplementary File 3 for "SPIRO – the automated Petri plate imaging platform designed by biologists, for biologists": newpass.html

{% extends "layout.html" %}
{% block title %}Set new password{% endblock %}
{% block content %}

Set new password for {{ name }}
{% if nopass %}
{% else %}

Current password

{% endif %}

Password

Password again

Set password

{% endblock %}
