## Supplementary File 3 for "SPIRO – the automated Petri plate imaging platform designed by biologists, for biologists": shutdown.html

{% extends "layout.html" %}
{% block title %}Shutting down{% endblock %}
{% block content %}

### Powering off system

It is safe to unplug the power when the LEDs on the system board are turned off.

{% endblock %}
