## Supplementary File 3 for "SPIRO – the automated Petri plate imaging platform designed by biologists, for biologists": webui.py

```
# webui.py -
#   main web ui controller for spiro
#
# - Jonas Ohlsson <jonas.ohlsson .a. slu.se>
#

import io
import os
import time
import shutil
import signal
import hashlib
import subprocess
from threading import Thread, Lock, Condition

from waitress import serve
from flask import Flask, render_template, Response, request, redirect, url_for, session, flash, abort

from spiro.config import Config
from spiro.logger import log, debug
from spiro.experimenter import Experimenter

app = Flask(__name__)
app.jinja_env.trim_blocks = True
app.jinja_env.lstrip_blocks = True

class Rotator(Thread):
    def __init__(self, value):
        Thread.__init__(self)
        self.value = value
    
    def run(self):
        lock.acquire()
        try:
            hw.motorOn(True)
            time.sleep(0.5)
            hw.halfStep(self.value, 0.03)
            time.sleep(0.5)
        finally:
            hw.motorOn(False)
            lock.release()


class StreamingOutput(object):
    def __init__(self):
        self.frame = None
        self.buffer = io.BytesIO()
        self.condition = Condition()

    def write(self, buf):
        if buf.startswith(b'\xff\xd8'):
            # New frame, copy the existing buffer's content and notify all
            # clients it's available
            self.buffer.truncate()
            with self.condition:
                self.frame = self.buffer.getvalue()
                self.condition.notify_all()
            self.buffer.seek(0)
        return self.buffer.write(buf)


class StillOutput(object):
    def __init__(self):
        self.frame = None
        self.buffer = io.BytesIO()

    def write(self, buf):
        if buf.startswith(b'\xff\xd8'):
            self.buffer.truncate()
            self.frame = self.buffer.getvalue()
            self.buffer.seek(0)
        return self.buffer.write(buf)


class ZoomObject(object):
    def __init__(self):
        self.roi = 1
        self.x = 0.5
        self.y = 0.5
    
    def set(self, x=None, y=None, roi=None):
        '''convenience function for setting zoom/pan'''
        if x is not None: self.x = x
        if y is not None: self.y = y
        if roi is not None: self.roi = roi
        self.apply()

    def apply(self):
        '''checks and applies zoom/pan values on camera object'''
        self.roi = max(min(self.roi, 1.0), 0.2)
        limits = (self.roi / 2.0, 1 - self.roi / 2.0)
        self.x = max(min(self.x, limits[1]), limits[0])
        self.y = max(min(self.y, limits[1]), limits[0])
        camera.zoom = (self.y - self.roi/2.0, self.x - self.roi/2.0, self.roi, self.roi)


def public_route(decorated_function):
    '''decorator for routes that should be accessible without being logged in'''
    decorated_function.is_public = True
    return decorated_function


def not_while_running(decorated_function):
    '''decorator for routes that should be inaccessible while an experiment is running'''
    decorated_function.not_while_running = True
    return decorated_function


@app.before_request
def check_route_access():
    '''checks if access to a certain route is granted. allows anything going to /static/ or that is marked public.'''
    if not request.endpoint: abort(404)
    if cfg.get('password') == '' and not any([request.endpoint == 'newpass', request.endpoint == 'static']):
        return redirect(url_for('newpass'))
    if any([request.endpoint == 'static',
            checkPass(session.get('password')),
            getattr(app.view_functions[request.endpoint], 'is_public', False)]):
        if experimenter.running and getattr(app.view_functions[request.endpoint], 'not_while_running', False):
            return redirect(url_for('empty'))
        return  # Access granted
    else:
        return redirect(url_for('login'))


def checkPass(pwd):
    if pwd:
        hash = hashlib.sha1(pwd.encode('utf-8'))
        if hash.hexdigest() == cfg.get('password'):
            return True
    return False


@app.route('/index.html')
@app.route('/')
def index():
    if experimenter.running:
        return redirect(url_for('experiment'))
    if restarting:
        return render_template('restarting.html', refresh='60; url=/', message="Rebooting system...")
    return render_template('index.html', live=livestream, focus=cfg.get('focus'), led=hw.led, name=cfg.get('name'))


@app.route('/empty')
def empty():
    return render_template('unavailable.html'), 409


@public_route
@app.route('/login', methods=['GET', 'POST'])
def login():
    if request.method == 'POST':
        pwd = request.form['password']
        if checkPass(pwd):
            session['password'] = pwd
            log("Web user successfully logged in from IP " + request.remote_addr)
            return redirect(url_for('index'))
        else:
            flash("Incorrect password.")
            log("Incorrect password in web login from IP " + request.remote_addr)
            return redirect(url_for('login'))
    else:
        return render_template('login.html', name=cfg.get('name'))


@public_route
@app.route('/logout')
def logout():
    session['password'] = ''
    return redirect(url_for('login'))


@public_route
@app.route('/newpass', methods=['GET', 'POST'])
def newpass():
    if request.method == 'POST':
        currpass = request.form['currpass']
        pwd1 = request.form['pwd1']
        pwd2 = request.form['pwd2']

        if cfg.get('password') != '':
            if not checkPass(currpass):
                flash("Current password incorrect.")
                return render_template('newpass.html', name=cfg.get('name'))

        if pwd1 == pwd2:
            hash = hashlib.sha1(pwd1.encode('utf-8'))
            cfg.set('password', hash.hexdigest())
            session['password'] = pwd1
            flash("Password was changed.")
            log("Password was changed by user with IP " + request.remote_addr)
            return redirect(url_for('index'))
        else:
            flash("Passwords do not match.")
            log("Password change attempt failed by user with IP " + request.remote_addr)
            return redirect(url_for('newpass'))
    else:
        return render_template('newpass.html', nopass=cfg.get('password') == '', name=cfg.get('name'))


@not_while_running
@app.route('/zoom/<int:value>')
def zoom(value):
    zoomer.set(roi=float(value / 100))
    return redirect(url_for('index'))


@not_while_running
@app.route('/pan/<dir>/<value>')
def pan(dir, value):
    if dir == 'x':
        zoomer.set(x = zoomer.x + float(value))
    elif dir == 'y':
        zoomer.set(y = zoomer.y + float(value))
    return redirect(url_for('index'))


@not_while_running
@app.route('/live/<value>')
def switch_live(value):
    if setLive(value):
        zoomer.set(0.5, 0.5, 1)
    if value == 'on':
        camera.shutter_speed = 0
        camera.exposure_mode = "auto"
    return redirect(url_for('index'))


def setLive(val):
    global livestream
    prev = livestream
    if val == 'on' and livestream != True:
        livestream = True
        camera.resolution = "2592x1944"
        camera.start_recording(liveoutput, format='mjpeg', resize='1024x768')
    elif val == 'off' and livestream == True:
        livestream = False
        camera.stop_recording()
        camera.resolution = camera.MAX_RESOLUTION
    return prev != livestream


@not_while_running
@app.route('/led/<value>')
def led(value):
    if value == 'on':
        hw.LEDControl(True)
    elif value == 'off':
        hw.LEDControl(False)
    return redirect(url_for('index'))


@not_while_running
@app.route('/rotate/<int:value>')
def rotate(value):
    if value > 0 and value <= 400:
        rotator = Rotator(value)
        rotator.start()
    return redirect(url_for('index'))


@not_while_running
@app.route('/findstart')
@app.route('/findstart/<int:value>')
def findstart(value=None):
    hw.motorOn(True)
    if not value:
        hw.findStart()
    elif value > 0 and value < 400:
        hw.findStart(calibration=value)
    time.sleep(0.5)
    hw.motorOn(False)
    return redirect(url_for('index'))


def liveGen():
    while True:
        with liveoutput.condition:            
            got_frame = liveoutput.condition.wait(timeout=0.1)
        if got_frame:
            yield (b'--frame\r\nContent-Type: image/jpeg\r\n\r\n' + liveoutput.frame + b'\r\n')
        else:
            # failed to acquire an image; return nothing instead of waiting
            yield b''
            

@not_while_running
@app.route('/stream.mjpg')
def liveStream():
    return Response(liveGen(), mimetype='multipart/x-mixed-replace; boundary=frame')


@app.route('/nightstill.png')
def nightStill():
    if nightstill.seek(0, io.SEEK_END) == 0:
        return redirect(url_for('static', filename='empty.png'))
    nightstill.seek(0)
    return Response(nightstill.read(), mimetype="image/png")


@app.route('/daystill.png')
def dayStill():
    if daystill.seek(0, io.SEEK_END) == 0:
        return redirect(url_for('static', filename='empty.png'))
    daystill.seek(0)
    return Response(daystill.read(), mimetype="image/png")


@app.route('/lastcapture.png')
def lastCapture():
    if not experimenter.last_captured:
        return redirect(url_for('static', filename='empty.png'))
    else:
        try:
            with open(experimenter.last_captured, 'rb') as f:
                return Response(f.read(), mimetype="image/png")
        except Exception as e:
            print("Could not read last captured image:", e)
            return redirect(url_for('static', filename='empty.png'))


def takePicture(obj):
    obj.truncate()
    obj.seek(0)
    camera.capture(obj, format="png")
    obj.seek(0)


def grabExposure(time):
    global dayshutter, nightshutter
    if time in ['day', 'night']:
        if time == 'day':
            takePicture(daystill)
            dayshutter = camera.shutter_speed
        else:
            #camera.color_effects = (128, 128)
            takePicture(nightstill)
            camera.color_effects = None
            nightshutter = camera.shutter_speed
        return redirect(url_for('exposure', time=time))
    else:
        abort(404)


@not_while_running
@app.route('/focus/<int:value>')
def focus(value):
    value = min(1000, max(10, value))
    hw.focusCam(value)
    cfg.set('focus', value)
    return redirect(url_for('index'))


@app.route('/experiment', methods=['GET', 'POST'])
def experiment():
    if request.method == 'POST':
        if request.form['action'] == 'start':
            if experimenter.running:
                flash("Experiment is already running.")
            else:
                if request.form.get('duration'): experimenter.duration = int(request.form['duration'])
                else: experimenter.duration = 7
                if request.form.get('delay'): experimenter.delay = int(request.form['delay'])
                else: experimenter.delay = 60
                if request.form.get('directory'): experimenter.dir = os.path.expanduser(os.path.join('~', request.form['directory'].replace('/', '-')))
                else: experimenter.dir = os.path.expanduser('~')
                setLive('off')
                zoomer.set(roi=1.0)
                log("Starting new experiment.")
                experimenter.next_status = 'run'
                experimenter.status_change.set()
                # give thread time to start before presenting template
                time.sleep(1)
        elif request.form['action'] == 'stop':
            experimenter.stop()
            time.sleep(1)

    if os.path.exists(experimenter.dir):
        df = shutil.disk_usage(experimenter.dir)
    else:
        df = shutil.disk_usage(os.path.expanduser('~'))

    diskspace = round(df.free / 1024 ** 3, 1)
    diskreq = round(experimenter.nshots * 4 * 10 / 1024, 1)
    return render_template('experiment.html', running=experimenter.running, directory=experimenter.dir, 
                           starttime=time.ctime(experimenter.starttime), delay=experimenter.delay,
                           endtime=time.ctime(experimenter.endtime), diskspace=diskspace, duration=experimenter.duration,
                           status=experimenter.status, nshots=experimenter.nshots + 1, diskreq=diskreq, name=cfg.get('name'),
                           defname=experimenter.getDefName())


@not_while_running
def exposureMode(time):
    if time == 'day':
        camera.iso = cfg.get('dayiso')
        camera.shutter_speed = 1000000 // cfg.get('dayshutter')
        camera.exposure_mode = "off"
        hw.LEDControl(False)
        return redirect(url_for('exposure', time='day'))
    elif time == 'night':
        camera.iso = cfg.get('nightiso')
        camera.shutter_speed = 1000000 // cfg.get('nightshutter')
        camera.exposure_mode = "off"
        hw.LEDControl(True)
        return redirect(url_for('exposure', time='night'))
    elif time == 'auto':
        camera.shutter_speed = 0
        camera.iso = 0
        camera.exposure_mode = "auto"
        return redirect(url_for('index'))
    abort(404)


@not_while_running
@app.route('/shutter/<time>/<int:value>')
def shutter(time, value):
    if time in ['day', 'night', 'live']:
        value = max(10, min(value, 1000))
        camera.shutter_speed = 1000000 // value
        return redirect(url_for('index'))
    else:
        abort(404)


@not_while_running
@app.route('/exposure/<time>', methods=['GET', 'POST'])
def exposure(time):
    if not time in ['day', 'night']: abort(404)
    ns=None
    ds=None

    if request.method == 'POST':
        shutter = request.form.get('shutter')
        if shutter:
            shutter = int(shutter)
            shutter = max(10, min(shutter, 1000))
            cfg.set(time + 'shutter', shutter)
            flash("New shutter speed for " + time + " images: 1/" + str(shutter))
        iso = request.form.get('iso')
        if iso:
            iso = int(iso)
            iso = max(50, min(iso, 800))
            cfg.set(time + 'iso', iso)
            flash("New ISO for " + time + " images: " + str(shutter))

        exposureMode(time)
        grabExposure(time)
    else:
        exposureMode(time)
        setLive('on')
        camera.exposure_mode = "off"

    if nightshutter:
        ns = 1000000 // nightshutter
    if dayshutter:
        ds = 1000000 // dayshutter

    return render_template('exposure.html', shutter=cfg.get(time+'shutter'), time=time, 
                           nightshutter=ns, dayshutter=ds, name=cfg.get('name'), iso=camera.iso,
                           dayiso=cfg.get('dayiso'), nightiso=cfg.get('nightiso'))


@not_while_running
@app.route('/calibrate', methods=['GET', 'POST'])
def calibrate():
    if request.method == 'POST':
        value = request.form.get('calibration')
        if value:
            value = int(value)
            value = max(0, min(value, 399))
            cfg.set('calibration', value)
            flash("New value for start position: " + str(value))
    exposureMode('auto')
    setLive('on')
    return render_template('calibrate.html', calibration=cfg.get('calibration'), name=cfg.get('name'))


@not_while_running
@app.route('/exit')
def exit():
    global restarting
    restarting = True
    signal.alarm(1)
    return redirect(url_for('wait_for_restart'))


@not_while_running
@app.route('/reboot')
def reboot():
    global restarting
    restarting = True
    subprocess.Popen(['sudo', 'shutdown', '-r', 'now'])
    return redirect(url_for('index'))


@not_while_running
@app.route('/shutdown')
def shutdown():
    subprocess.run(['sudo', 'shutdown', '-h', 'now'])
    return render_template('shutdown.html')


@app.route('/settings', methods=['GET', 'POST'])
def settings():
    if request.method == 'POST':
        if request.form.get('name'):
            cfg.set('name', request.form.get('name'))
    return render_template('settings.html', name=cfg.get('name'), running=experimenter.running)


@not_while_running
@app.route('/restarting')
def wait_for_restart():
    if restarting:
        return render_template('restarting.html', refresh=5,
                               message="Restarting Web UI...")
    else:
        return redirect(url_for('index'))


@app.route('/files')
def file_browser():
    dirs = []
    dir = os.path.expanduser('~')
    df = shutil.disk_usage(dir)
    diskspace = round(df.free / 1024 ** 3, 1)

    for entry in os.scandir(dir):
        if entry.is_dir() and os.path.dirname(entry.path) == dir and not entry.name.startswith('.'):
            dirs.append(entry.name)
    return render_template('filemanager.html', dirs=sorted(dirs), diskspace=diskspace, name=cfg.get('name'), running=experimenter.running)


@app.route('/get/<exp_dir>.zip')
def make_zipfile(exp_dir):
    'creates a zipfile on the fly, and streams it to the client'
    dir = os.path.expanduser('~')
    zip_dir = os.path.abspath(os.path.join(dir, exp_dir))
    if verify_dir(zip_dir):
        p = subprocess.Popen(['/usr/bin/zip', '-r', '-0', '-', os.path.basename(zip_dir)], stdout=subprocess.PIPE, cwd=dir)
        return Response(stream_popen(p), mimetype='application/zip')
    else:
        abort(404)


@app.route('/delete/<exp_dir>', methods=['GET', 'POST'])
def delete_dir(exp_dir):
    dir = os.path.expanduser('~')
    del_dir = os.path.abspath(os.path.join(dir, exp_dir))

    if request.method == 'GET':
        return render_template('delete.html', dir=exp_dir)
    else:
        if os.path.abspath(experimenter.dir) == del_dir and experimenter.running:
            flash('Cannot remove active experiment directory. Please stop experiment first.')
            return redirect(url_for('file_browser'))
        if verify_dir(del_dir):
            shutil.rmtree(del_dir)
            flash(f'Directory {exp_dir} deleted.')
            return redirect(url_for('file_browser'))


def verify_dir(check_dir):
    '''checks that the directory is
       1. immediately contained within the appropriate parent dir
       2. does not contain initial dots, and
       3. is indeed a directory'''
    check_dir = os.path.abspath(check_dir)
    dir = os.path.expanduser('~')
    return os.path.dirname(check_dir) == dir and not os.path.basename(check_dir).startswith('.') and os.path.isdir(check_dir)


def stream_popen(p):
    '''generator for sending STDOUT to a web client'''
    data = p.stdout.read(128*1024)
    while data:
        yield data
        data = p.stdout.read(128*1024)


liveoutput = StreamingOutput()
nightstill = io.BytesIO()
daystill = io.BytesIO()
zoomer = ZoomObject()
cfg = Config()
lock = Lock()

experimenter = None
nightshutter = None
dayshutter = None
camera = None
hw = None

restarting = False
livestream = False

def start(cam, myhw):
    global camera, hw, experimenter
    camera = cam
    hw = myhw
    experimenter = Experimenter(hw=hw, cam=cam)
    experimenter.start()
    if cfg.get('secret') == '':
        secret = hashlib.sha1(os.urandom(16))
        cfg.set('secret', secret.hexdigest())
    app.secret_key = cfg.get('secret')
    try:
        camera.meter_mode = 'spot'
        camera.rotation = 90
        setLive('on')
        #app.run(host="0.0.0.0", port=8080, debug=False)
        # use a tcp timeout of 20 seconds to improve hanging behavior in live view
        serve(app, listen="*:8080", threads=8, channel_timeout=20)
    finally:
        stop()

def stop():
    experimenter.stop()
    experimenter.quit = True
    experimenter.status_change.set()
```
