## Supplementary File 4 for "SPIRO – the automated Petri plate imaging platform designed by biologists, for biologists"

### SPIRO

#### Assays Manual

#### Contents

1. Overview ... 3
2. Setting up SPIRO experiments ... 4
  2.1 Plate preparation ... 4
  2.2 SPIRO software settings ... 7
  2.3 Accessing SPIRO Data ... 10
  2.4 Troubleshooting ... 10
3. Semi-automated data processing in SPIRO assays ... 11
  3.1 Overview ... 11
  3.3 Software installation ... 13
    3.3.1 Software for quantitative data analysis ... 13
    3.3.2 Software for image analysis ... 13
    3.3.3 SPIRO Assay Customizer ... 14
  4.4 Image preprocessing ... 15
    4.4.1 Overview ... 15
    4.4.2 Using the Preprocessing macro ... 16
  4.5 SPIRO Seed Germination Assay ... 18
    4.5.1 Overview ... 18
    4.5.2 Using the Germination Macro ... 21
    4.5.3 Using R scripts for the Germination Assay ... 23
  4.6 SPIRO Root Growth Assay ... 26
    4.6.1 Overview ... 26
    4.6.2 Using the Root Growth Macro ... 27
    4.6.3 Using Root growth R scripts ... 30
  4.7 SPIRO Assay Customizer ... 33
    4.7.1 Merge assays ... 33
    4.7.2 Customize assay ... 34
Appendix 1: Quick start guides ... 35
  Macro installation ... 35
  Preprocessing macro ... 35
  Germination macro ... 35
  Root Growth macro ... 36

Analysis in R ........................................................ 36
References ........................................................ 37

### 1. Overview

This document provides general information and specific instructions on the use and functionality of SPIRO, the **S**mart **P**late **I**maging **Ro**bot.

Protocols for preparing Petri plates with samples, adjusting imaging settings in the SPIRO software, downloading the data and managing the system storage are described in Section 2.

Section 3 describes the use and functionality of the semi-automated assays for tracking seed germination and root growth.

Appendix 1 provides a quick guide for the assays that can be helpful for experienced or impatient users.

Please note that this document links to a number of online resources, such as trouble-shooting tables and GitHub repositories, which are continuously updated. If you encounter an issue with using the SPIRO system or its software, please report these issues in the appropriate GitHub repository (specific links are provided in the corresponding sections below).

### 2. Setting up SPIRO experiments

#### 2.1 Plate preparation

- The SPIRO plate holders are designed for two standard plate types: ∅ 9 cm round Petri plates (Sarstedt, 82.1473.001) and 12 cm square plates (Greiner Bio-One, 688102).

- For Arabidopsis thaliana seeds/seedlings grown in 12 cm square plates, use 35 mL of 0.5x MS medium solidified with plant agar. 9 cm round plates can be used with 15 mL of 0.5xMS medium.

  Conditions in most growth cabinets for plants cause extensive water condensation on the Petri plates' lids, rendering them too opaque for imaging. To circumvent this issue, we recommend to image from the bottom side of the plates through the medium. Hence it is important for the medium layer to be as transparent as possible. In our experience, using the above recommended volumes of the medium allows for sufficient image quality compatible with the automated analysis and does not cause discernible growth or germination phenotypes. Using more transparent solidifying agents, such as Phytagel (Sigma, P8169), did not influence the quality of the analysis.

- We strongly recommend using custom printed seed plating guides (**Table S2** of this manuscript and in the SPIRO Hardware Repository[1]) for positioning the seeds on plates.

  For germination assays, we recommend plating seeds at least 4 mm apart. For root growth assays, it is preferable to have ca 5–10 mm distance between seeds in a single row to decrease the probability of roots from neighboring seedlings crossing each other. Additionally, avoid placing seeds closer than 1.5 cm towards the edges of a plate, where reflections might interfere with seed recognition (**Fig. 1**).

- For convenience of seed plating, we recommend sterilizing *Arabidopsis thaliana* seeds using the ethanol method.

  Incubate seeds in an Eppendorf tube with 70% ethanol with 0.05% TritonX-100 (20 minutes, at RT, agitating). Replace the solution with 96% ethanol, repeat the washing step twice to get rid of Triton X-100 (avoid incubating seeds in 96% ethanol for longer than 10 minutes). Remove all ethanol from the tube and leave the tube open for the seeds to air dry under sterile conditions.

- While plating the seeds, avoid damaging the medium surface. Imperfections in the medium might generate noise in the automated image analysis.

  In our experience, transferring seeds onto plates using sterile toothpicks is the most convenient method.

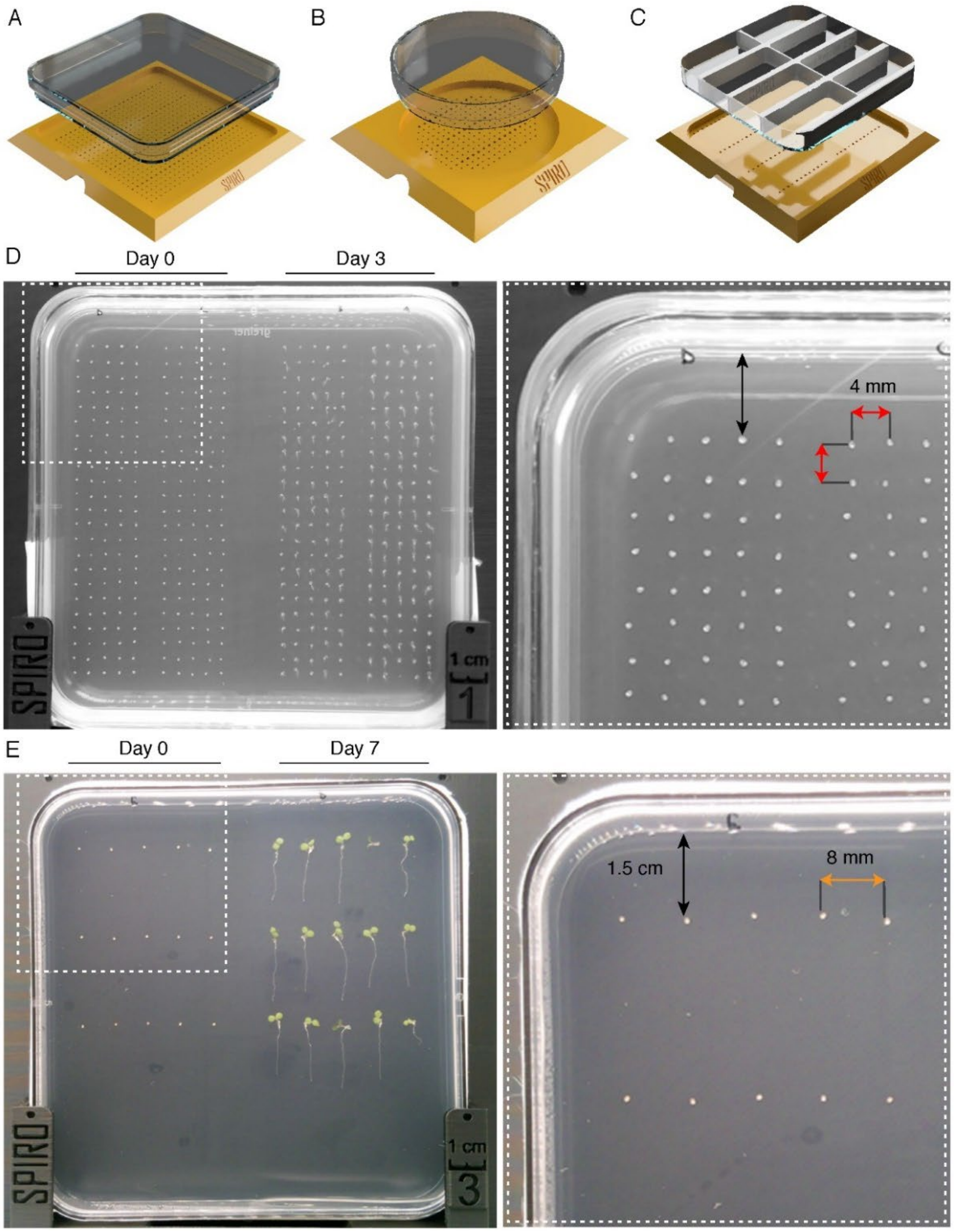


**Figure 1. Examples of plating seeds for germination and root growth assays**.
(**A**) 3D printed seed plating guide for 12 cm square plates.(**B**) An example of a plate imaged by SPIRO for germination assay on the first and third days of the experiment. Seeds are placed 1.5 cm away from plate edges and 4 mm away from each other. (**C**) 3D printed seed plating guide for 9 cm round plates. (**D**) An example of a plate imaged by SPIRO for the root growth assay at the first and the seventh day of the experiment. Seeds are placed 1.5 cm away from plate edges and 8 mm away from each other. (**E**) 3D printed insets that can be glued into Petri plates to create multi-chamber compartments applicable for different treatments. The insets can be used with the corresponding seed plating guides.

- Use custom lids printed with black filament to eliminate reflections (**Table S2** of this manuscript and in the SPIRO Hardware Repository[1]).

Reflections in the plastic of normal lids are detrimental to automated image analysis, especially during root detection (**Fig. 2**). The printed lids should preferably be sterilized using UV light. Sterilization in 70% ethanol is also possible, but must be followed by a prolonged drying of the lids. Ethanol left in pores of the plastic might impact seedling growth.

- For long time-lapse experiments, plates should be properly sealed to prevent evaporation.

Please note that plate holders are quite tight and will not allow for bulky layers of Parafilm or tape. We recommend using plastic seal (Phytotechlab, Product ID: A003).

- Place the plates into the holders of SPIRO with lids towards the cube.

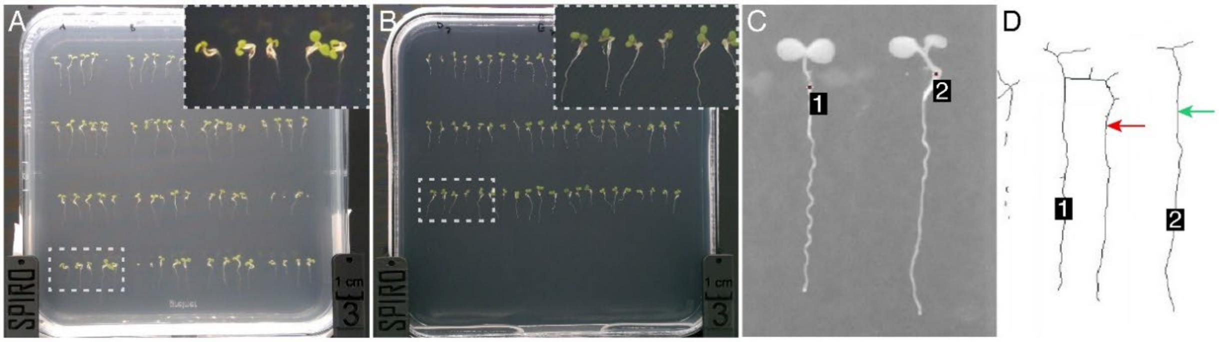


**Figure 2. Reducing reflections in the Petri plate lids is crucial for automated image processing.** Examples of images acquired using normal (**A**) and 3D-printed anti-reflection (**B)** lids. (**C**) A magnified part of the photo shown in (A) processed by the SPIRO root growth assay, seedlings recognized by the software are indicated with numbers. (**D**) The result of image segmentation by the SPIRO root growth assay performed on the data shown in (C). Background is shown in white, while objects recognized as seedlings are shown as black lines with ID numbers. Green arrow, seedling 2; red arrow, seedling 2 reflection being mistakenly annotated as a part of seedling 1.

#### 2.2 SPIRO software settings

- SPIRO is controlled by the designated software on the Raspberry Pi computer and operated by the user via a web-based user interface. Thus, to enable access to the SPIRO software during experiment setup, the robot must be connected to the internet either via Wi-Fi or Ethernet cable, or set up to act as a Wi-Fi hotspot.

  Since plant growth cabinets are often located where an internet connection is not available, the SPIRO computer can be used as a Wi-Fi hotspot (for instructions see **File S2** of this manuscript and SPIRO Software Repository[2]).

- To access the SPIRO web UI, open a web browser and type in the address http://x.y.z.w:8080/ (where x.y.z.w is IP address of your Raspberry Pi computer). Log in using the password you provided during SPIRO Web UI initial setup (**File S2** of this manuscript and SPIRO Software Repository[2]).

  Note that in Wi-Fi hotspot mode, any address will work (although we recommend using the address spiro.local).

- In the **Live view** tab (**Fig. 3**), use the focus slider or type in numbers to adjust the **focus** (for SPIROs with motorized cameras), **Zoom** and **pan** (icon with four arrows) tools might be helpful for verification of the focus. Please note, that the acquired images must contain the complete 1 cm scale bar engraved on SPIRO if they are to be used with the provided automated assays.

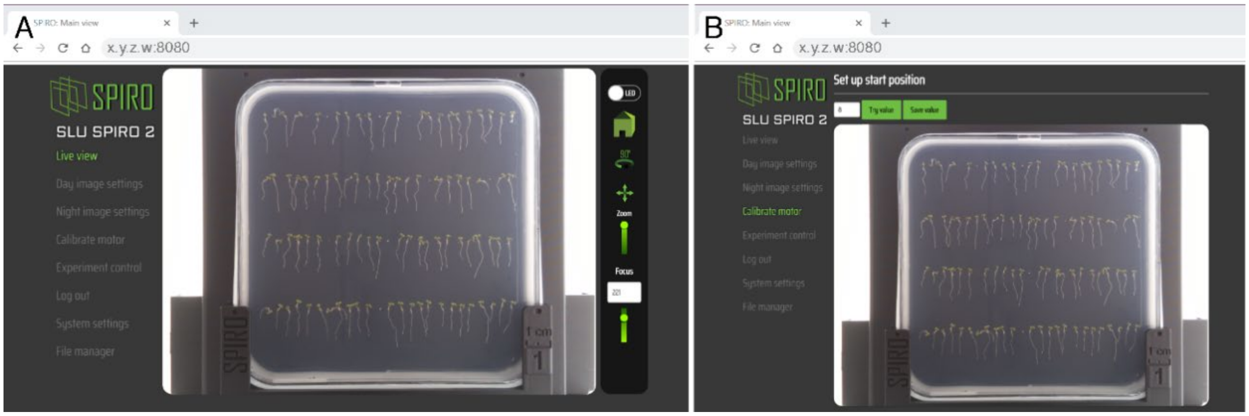


**Figure 3. Using the web interface to set up imaging.**
Live view (**A**) and Calibrate motor (**B**) tabs of the SPIRO web UI should be used to adjust the focus and alignment of the cube position with the camera.

- Press the **Home** button on the right panel of the Live view and check if the first plate is moved to face the camera. If the plate is not facing the camera, you can adjust and verify the optimal value for cube rotation angle in the **Calibrate motor** tab.

- **Day** and **night image settings** tabs allow adjustment of ISO and camera shutter speed to optimize image light intensity under desired imaging conditions.

As the accuracy of image processing for semi-automated assays depends greatly on the image quality, avoid acquiring over- or under-exposed images (**Fig. 4**).

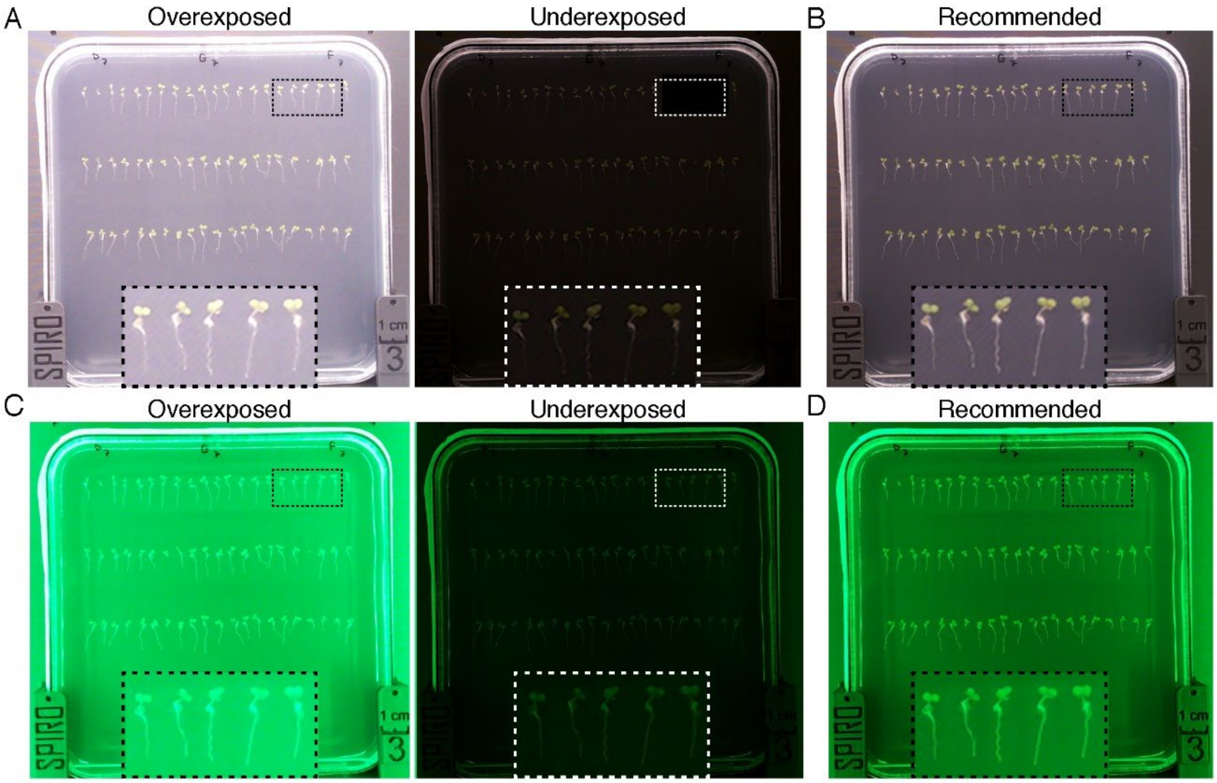


**Figure 4. Impact of the exposure settings on the image quality.** Day and night image settings tabs allow adjustment of ISO and shutter speed for optimal image light intensity under day and night conditions. (**A, C**) Examples of over- and under-exposed day and night images that impacts quality of data. (**B, D**) Examples of images with correct exposure, optimal for the semi-automated assays.

- In the **Experiment** control tab, you will be prompted to enter experiment folder name.

Please note that this name will later be used during data analysis to create a unique ID for each seed/seedling. If you wish to combine data from different experiments for analysis it is important to have unique experiment folder names. You will also be asked to provide imaging frequency and duration of your experiment. SPIRO is capable of imaging at the maximum rate of 4 plates/ca 2 minutes. **SPIRO germination and root growth semi-automated assays were optimized for imaging every 30 and 60 minutes, respectively**.

- The software will estimate required and available storage space. If amount of available space is less than is required, it will be shown in red.

Please note that the experiment will terminate, when no more storage space will be available. To free up space, you can remove unnecessary files from the Raspberry Pi computer even during a running experiment (see Section 2.3).

#### 2.3 Accessing SPIRO Data

- The experiment folder with acquired images will be saved on the microSD card of the SPIRO Raspberry Pi computer.

- To download SPIRO experiment data, click on the menu item *File manager* in the web interface.

  The File manager presents a list of all experiment folders stored on the microSD card. For each item, there are buttons for downloading and deleting data. Downloaded data comes as a ZIP file, which needs to be extracted before using it for the semi-automated SPIRO assays.

- SPIRO captures images with the following directory structure and file naming scheme (**Fig. 5**):

*<experiment name>/<plateN>/<date>-<time>-[day/night].png*

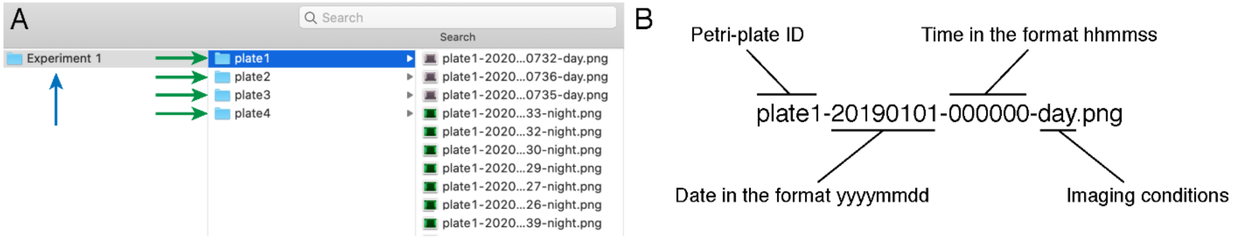


**Figure 5. SPIRO image data file structure.**
For each experiment all images of an individual plate are grouped into a subfolder (green arrow). All subfolders are contained within a single main folder (blue arrow). (**B**) Naming images for the analysis is automatically done by SPIRO.

- To free up space on the system, directories can be deleted from the *File manager* tab in the web interface. Note that deleting a directory which is currently being used for imaging is not allowed. The File Manager will also display how much disk space is remaining.

#### 2.4 Troubleshooting

- For troubleshooting, please refer to this online table.

- Please submit reports about errors and suggestions for improvement as a GitHub issue.

### 3. Semi-automated data processing in SPIRO assays

#### 3.1 Overview

SPIRO Germination and Root Growth Assays were specifically designed for semi-automated analyses of high-throughput data acquired using SPIRO and optimized for *Arabidopsis thaliana* seeds and seedlings, but can be tuned for use with other species.

The SPIRO Germination and Root Growth Assays comprise three major parts, with user-guided[U] and automated[A] steps that facilitate the assays' applicability for a broad range of experimental layouts. A simplified version of the SPIRO Germination

**Image preprocessing**

- Setting the scale in cm[U]
- Reducing the size of images by cropping the unnecessary parts[U]
- Creating a time-lapse stack file[A]
- Correction for drift (if needed)[A]
- Conversion of color images into greyscale 8-bit format[A]

**Image analysis**

- Defining groups of seeds or seedlings to be analyzed, e.g., different genotypes, treatments[U]
- Defining the time-range of interest from the time-lapse data[U]
- Identifying seeds on the images[A]
- Obtaining data for detecting seed germination time points[A]
- *(Root growth assay only)* Detecting coordinates for the root start of each seedling[A]
- *(Root growth assay only)* Segmenting images to identify each seedling[A]
- *(Root growth assay only)* Recording changes in root length over time for each seedling[A]
- Generating TIFF time-lapse files with graphical output of the analysis[A]

- Generating TSV files with objects measurements at each time point[A]

**Data processing in R**

- Filtering out data points that do not conform to the required quality and detecting germination time point for each seed[A]
- Statistical analysis of germination parameters and seed sizes[A]
- *(Root growth assay only)* Building polynomial fit models to determine mean root length and root growth rate for each group[A]
- *(Root growth assay only)* Statistical analysis of the root length and growth data[A]
- Relabeling the groups and samples for the statistical analysis is possibly using SPIRO Assay Customizer[U]

Although the general workflow for both the germination and root growth assays is similar, it is important to note a few differences. Firstly, the number of biological replicates that can be fitted into a single plate will be an order of magnitude lower for the root growth assay when compared to germination assay (see section **2.1**). Secondly, although the root growth assay alone will also generate the same quantitative analysis and statistics about seed germination as the germination assay would, but unlike the dedicated germination assay it will not provide graphical macro output. **Importantly, the R scripts for statistical analysis will not work if both germination and root growth macros have been run on the same dataset**. Thus, if it is indeed desired to run both analyses on a single dataset, it is advisable to make a copy of the experiment folder after preprocessing and then run each analysis on a separate copy.

For impatient and/or experienced users, quick start guides are available in **Appendix 1.**

#### 3.3 Software installation

The macros for image analysis were written in the ImageJ Macro Language and tested on ImageJ version 1.52p.

The R scripts were tested on R versions 3.6.1 and 4.0.2. For the sake of convenience, we highly recommend using RStudio to run the scripts, and Git version control for downloading and keeping scripts and macros up to date.

##### *3.3.1 Software for quantitative data analysis*

a. Download and install R.

b. Download and install Git.

c. Download and install R Studio and dependencies:

- Open R Studio and enable Git.
- Create a new project with version control. In the menu, go to *File → New project→ Version control → Git*.
    - Repository URL: https://github.com/jiaxuanleong/SPIRO.Assays.
    - Type in the desired directory name (e.g. SPIRO.Assays).
    - Pick the location suitable for keeping the scripts and macros for the assays (e.g., the user Documents folder).

##### *3.3.2 Software for image analysis*

a. Download and install Fiji (ImageJ with plugins).

- Allow updates.
- It is advisable to allocate 75% of available RAM to be used by ImageJ during high-throughput data processing. In Fiji, go to *Edit → Option → Memory & Threads*. Type in the memory amount in MB that ImageJ will be permitted to use.

b. Download and install the TurboReg plugin.

c. Download and install the MultiStackReg plugin.

d. Macros can be installed permanently by copy pasting the code into the Startup macro.fjii.ijm file (accessed via the menu item *Plugins → Macros → Startup Macros...*) or each time Fiji is restarted. For the latter option:

- Start Fiji.
- Go to *Plugins → Macros → Install*. Locate the folder downloaded while creating new project in R studio → locate and open the SPIRO_macros.ijm file.
- Macros will be available on the bottom of the option list under *Plugins → Macros.*

e. Note that if the macros are installed by pasting them into the startup macros file, they will need to be replaced whenever the macros are updated.

##### *3.3.3 SPIRO Assay Customizer*

SPIRO Assay Customizer is a useful companion tool for running the SPIRO Assays. It allows merging data from several experiments, adjusting grouping, and excluding seeds or seedlings from analysis. Although all these tasks can be performed in an Excel file, we recommend using the Customizer to avoid accidental mislabeling and deletion of data points.

The latest version of the Customizer can be downloaded from SPIRO Assay Customizer Repository[3]. Note that opening Customizer for the first time may trigger a default warning about suspicious software, which requires manual intervention. **On Windows systems**, if presented with the "Windows protected your computer" message, click on *More info* and then *Run anyway*. **On MacOS systems**, right-click on the application icon and choose *Open*. A popup will appear, urging you to not run the software. Press *Open* again and the software will run.

#### 4.4 Image preprocessing

##### *4.4.1 Overview*

The first step in either of the SPIRO assays is preprocessing of the raw data. It is used for:

- Creating a time-lapse file for each analyzed Petri plate.
- Setting scale in cm.
- Reducing file size by cropping off unnecessary background in images.
- Extracting green channel information from the RGB images and saving it as 8-bit greyscale data to facilitate further segmentation.
- If needed, correcting for drift between images caused by cube rotation.

To reduce RAM requirements, images are analyzed in batches, meaning that only a subset of images is processed at each step. The macro will eventually process all images present in the experiment folder. The default size of each batch is set to 350 images, but can be adjusted to the capabilities of your computer (for more info, see the troubleshooting table). The preprocessed output is an individual TIFF file for each plate, containing all acquired images for the plate. In ImageJ technical terms, such file is called an *image stack* and the individual images within it are henceforth referred to as *slices*. The main benefit of a stack file is the ease of navigating through the time-lapse series.

##### 4.4.2 Using the Preprocessing macro

a. Download the image data (see Section 2.3). The folder containing the images will be further referred to as the *Experiment folder*.

b. Make sure that all installation steps are complete (see Section 3.3).

c. Install SPIRO_Macros.ijm as described in the 3.3.2d and open the SPIRO_Preprocessing macro (*Plugins → Macros → SPIRO_Preprocessing*).

d. Follow the instructions provided by the macro.

- In the file picker, open the Experiment folder containing subfolders with images.
- You will be asked if the automatically selected area for cropping needs correction. Please make sure that the selected area includes the engraved scale bar and SPIRO logo (**Fig. 6A)**, as those features are needed for efficient drift correction.
- Please note that registration for drift correction is a time-consuming process and can be skipped if no drift is visible while scrolling through the slices of the stack.

e. The results of preprocessing will be saved in the experiment folder (**Fig. 6B**):

Experiment folder/Results/Preprocessing/platen_preprocessed.tif

f. For troubleshooting, please refer to this table.

g. Please submit reports about errors and suggestions for optimizations as a GitHub issue.

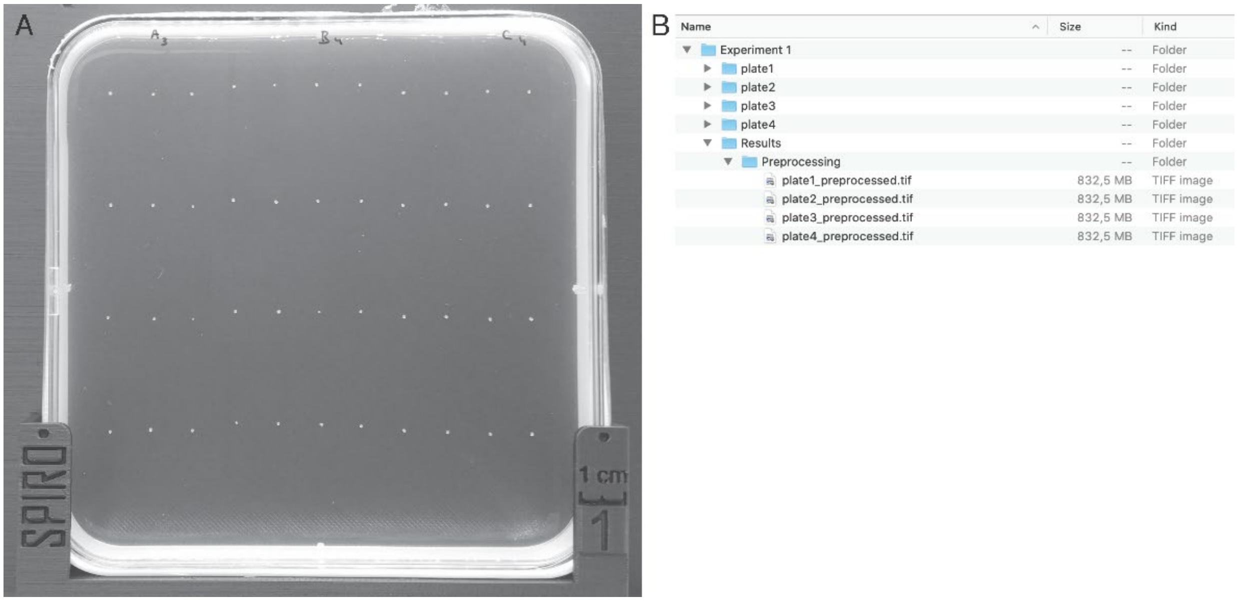


**Figure 6**. **Preprocessing macro output.**
**(A)** A correctly cropped image, including the engraved SPIRO marking. **(B)** The results of preprocessing will be saved as time-lapse TIFF stack files with names corresponding to images of plates being analyzed. The original imaging data will be kept unmodified. Preprocessed data can be further used for detection of germination and for tracking root length. The macros for these tasks will automatically analyze data for all preprocessed plates in the experiment folder.

#### 4.5 SPIRO Seed Germination Assay

##### *4.5.1 Overview*

The Germination Assay comprises automated image analysis using an ImageJ macro and subsequent data quality control followed by germination detection and statistical analysis using the corresponding R scripts. SPIRO Assay Customizer can be used to merge cleaned up data of multiple experiments to be analyzed together, regroup or rename samples, or to remove outliers.

To make the macro applicable for a wide range of experimental setups we included three user-guided steps into image processing:

1. The macro will prompt the user to select a subset of images representing the time interval of interest.
2. It will also prompt the user to select groups of seeds that should be analyzed together, e.g., the same genotype.
3. It will prompt manual control over removing or adding objects recognized as seeds on the image for further analysis.

The SPIRO Germination Assay was tuned for use with Arabidopsis. To enable its use for other species, seed detection parameters will likely need tweaking. For more info, see the Debug Mode Section in the SPIRO Assays Repository[4].

For each selected group the macro will identify the position of each individual seed at the first time point and track changes in each seed's perimeter and area over time.

The macro output is subsequently processed by two R scripts. First, data not conforming to the quality requirements is filtered out by cleanup_germination_data.R.

QC comprises the following rules, which are evaluated on a per-seed basis:

- Objects identified as seeds may not have an area exceeding 0.02 $cm^2$ (this area is larger than a recently germinated Arabidopsis seed). Time-lapse data for each seed is truncated at the time point where the area limit is met.
- Objects with areas of less than 0.002 $cm^2$ (smaller than a typical dry seed of Arabidopsis) are removed from analysis.

- If multiple objects remain within a seed position area after filtering are applied, the data for this seed is truncated at the point of the first occurrence of multiple objects (e.g., neighboring roots growing into the ROI).

After QC, the germination data is analyzed using the script process_germination_data.R. Seed germination is defined as radicle emergence event[5] that would inevitably lead to a progressive increase in the seeds' size. Thus, germination time is identified as the time point after which a steady increase in seed perimeter is observed. The same object imaged under day and night illumination conditions might be detected as having slightly different perimeters. To compensate for this, the script quantifies for each seed the difference between perimeter values detected during daytime conditions (the mean value for 8 h within the first 24 h of the experiment) and night-time conditions (the mean value for 8 h within the first 24 h of the experiment). Data for all time points are then normalized based on the mean perimeter difference.

For each seed, the script determines the following parameters to assess whether the seed has germinated:

- Initial seed perimeter: $P_{initial} = \frac{\sum_{i=1}^{5} P_i}{5}$, where P$i$ is the perimeter of the seed on slice number $i$.
- Perimeter change at timepoint $i$: $\Delta P_i = \frac{\sum_{x=1}^{r} P_{i+x}}{r-i} - P_{initial}$. Note that this value includes a forward-looking moving average of r slices. r is defined as either 2.5 h or 5 slices, whichever is highest.
- Per-timepoint perimeter change: $\Delta\Delta P_i = \Delta P_{i+1} - \Delta P_i$.
- Range for assessing constant increase in perimeter, s, which is either 5 h or 10 slices, whichever is highest.
- Magnitude of increase over the range s: $\Delta s = \Delta P_{i+s} - \Delta P_i$
- Magnitude of increase over the range 2$s$: $\Delta 2s = \Delta P_{i+2s} - \Delta P_i$

The time point $i$ is considered to be the germination time point if:

- $\Delta P_i$ is positive
- $\Delta\Delta P_i$ is positive for at least 80% of consecutive timepoints within the range s, starting from i
- $\Delta s$ is > 0.05 cm.

- $\Delta 2s$ is > 0.1 cm.
- The above statements are the first occurrence in the data for the seed.

Please note that the algorithm requires perimeter data to be available for approximately 12.5 h after germination has occurred. For assessing certain germination parameters, the R package germinationmetrics[6] is used.

##### *4.5.2 Using the Germination Macro*

a. Make sure that all installation steps are complete (see Section 3.3.2).

b. Make sure that preprocessing was performed on the dataset (see Section 4.4.2).

c. Install SPIRO_Macros.ijm as described in 3.3.2d and open the SPIRO_Germination macro (*Plugins → Macros → SPIRO_Germination*).

d. Follow the instructions provided by the macro.

- In the file picker, open the Experiment folder with the preprocessed data.
- Reducing the number of slices/time points included into analysis will significantly speed up the process. **While truncating the time-lapse stack, it is very important to keep in mind that the germination detection algorithm requires presence of at least 2.5 h before and approximately 12.5 h after a seed has germinated. Also include as much data as possible preceding germination, as this is used for perimeter normalization.**
- You will be prompted to select groups of seeds, e.g., different genotypes and/or treatments. This step is crucial for downstream statistical analysis of the quantitative data. If needed, grouping can be manually modified after data quality control to enable different comparisons without re-running image analysis (see Section 4.7)
- While selecting the groups of seeds that will be analyzed together, avoid including areas with reflections or imperfections on the medium surface, as these might be detected as seed-like objects.
- The macro will suggest using the same area selection and names for groups in all plates. These can be manually edited and adjusted. Please save the adjustments using the “update” button in the ROI manager.
- To enable analysis of samples imaged under various illumination conditions we added a user-guided step that allows selection/deselection of objects missed by the algorithm or falsely recognized as seeds.
- Results of the analysis for each group will be saved in the experiment folder:

  Experiment name/Results/Germination/plate*n*/*group name*

e. The macro will produce two output files for each group of seeds (**Fig. 7A**):

- *group name* germinationlabelled.tif is the graphical output of the macro. It contains a time-lapse stack of side-by-side comparison of original and segmented images for the corresponding group with numbered identified seeds. This information can be helpful to verify that image processing indeed identified seeds and only seeds (**Fig. 7B**).

- *group name* germination analysis.tsv is the quantitative output of the macro, it contains perimeter and area data for each seed at each time point.

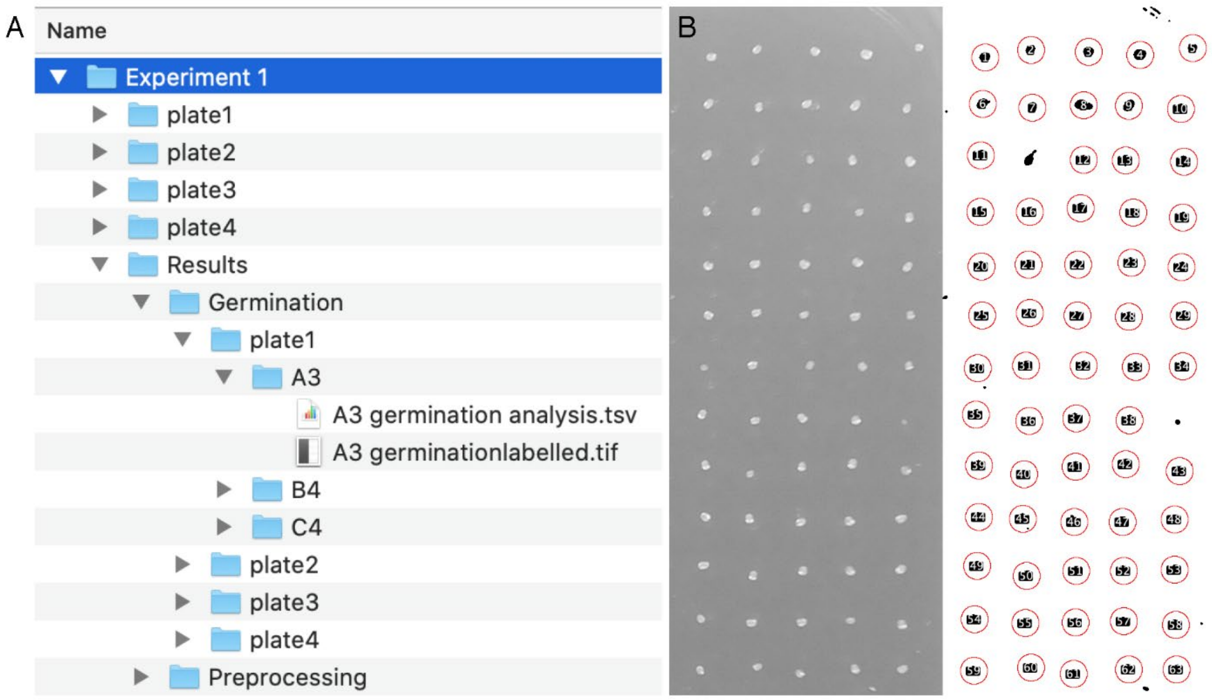


**Figure 7. Output of the Germination Macro.**
(**A**) The results of the germination macro will be saved in the Results/Germination/ Plate*n*/Group_name subfolder. The preprocessed file cropped according to the group selection and user-selected time truncation together with the results of image segmentation will be provided in the group_name germinationlabelled.tif. This file can be used to assess the quality of seed recognition. The file group_name germination analysis.tsv will contain quantitative data about size and perimeter changes for each seed over time. (**B**) An example of a single frame from a group_name germinationlabelled.tif stack file. The left part is the photo acquired by SPIRO and cropped by the user, and the right part is the result of this image segmentation by the Germination macro. Each red circle is centered around an object recognized as a seed, and the numbers indicate the number of a seed that can be then tracked in the TSV file.

##### *4.5.3 Using R scripts for the Germination Assay*

The Germination Assay includes two R scripts for data quality control (QC) and germination detection combined with the statistical analysis.

a. Make sure that all installation steps are complete (see Section 3.3.2).

b. Open RStudio.

c. If the SPIRO Assays project is not open, go to *File → Open Project*. In the file picker, locate the project that you saved during installation steps (see Section 3.3.2).

d. Open cleanup_germination_data.R (click on the corresponding file name in the Files tab in the bottom right panel of R studio).

e. Run the script: in the top menu find *Code → Source.* Select your experiment folder when prompted. For Windows users, this can be done via a file picker (please note, that RStudio may place this window behind the R Studio window). For MacOS users, right click on your experiment folder and hold the Option (⌥) key and select the "Copy *path* as Pathname" option, then copy paste the path to your experiment folder into the console of R Studio.

f. In the Console, messages indicating what seeds were removed from the final output file, and for what reason, will be printed.

g. Results of the QC step are saved in two files in the experiment folder in two files (**Fig. 8A**):

- In germination.postQC.log.tsv you can find a summary of which seeds were processed normally and which have been filtered out by the QC.
- germination.postQC.tsv contains quantitative macro output data that passed quality control.

The script will create a unique ID for each seed that will contain information about seed number, group name, plate number and experiment name. Unique IDs allow backtracking the quantitative data to the graphical output of the macro and also simultaneous analysis of data from several experiments. Importantly, in this file you can modify group names if you wish to perform different grouping for statistical analysis (see Section 4.7). Please note that editing data using Microsoft Excel is not recommended as this is an error-prone approach.

h. Make sure that groups reflect the comparisons you wish to make. If merging data from several experiments, relabeling/regrouping or removing of outliers is required, we recommend to use SPIRO Assay Customizer (see Section 4.7).

i. Run process_germination_data.R (click on the corresponding file name in the right bottom panel of R studio, then *Code* → *Source*). If needed, open the project again (see 4.5.3c).

j. Point the script to the experiment folder (see 4.5.3d)

k. Analysis can be performed several times on the same experiment, and results of each analysis run will be saved in a separate subfolder (**Fig. 8A**). The analysis produces the following output:

- descriptive_stats.tsv contains T50, mean germination time, number of germinated and ungerminated seeds for each group and mean seed size per group.
- germination-perseed.tsv contains germination times (expressed in h and slice number) as well as the seed size for each seed. This file also includes the QC log note.
- germination.kaplan-meier_test.tsv contains the log-rank *p* values for germination comparison between groups using Kaplan-Meier analysis.
- seedsize.t-tests.tsv contains group comparisons of seed sizes using Student's *t* test, expressed as raw and FDR-corrected values.
- A folder Germination Plots, containing illustrative plots of several germination parameters for each group (**Fig. 8B**).
- A folder Kaplan-Meier Plots, containing Kaplan-Meier plots for each pair of groups, and one for all groups combined (**Fig. 8C**).

l. For running further analyses, the file germination-perseed.tsv can be analyzed using any statistical software, including but not limited to R.

m. For troubleshooting, please refer to this table. Please submit reports about errors and suggestions for optimizations as a GitHub issue.

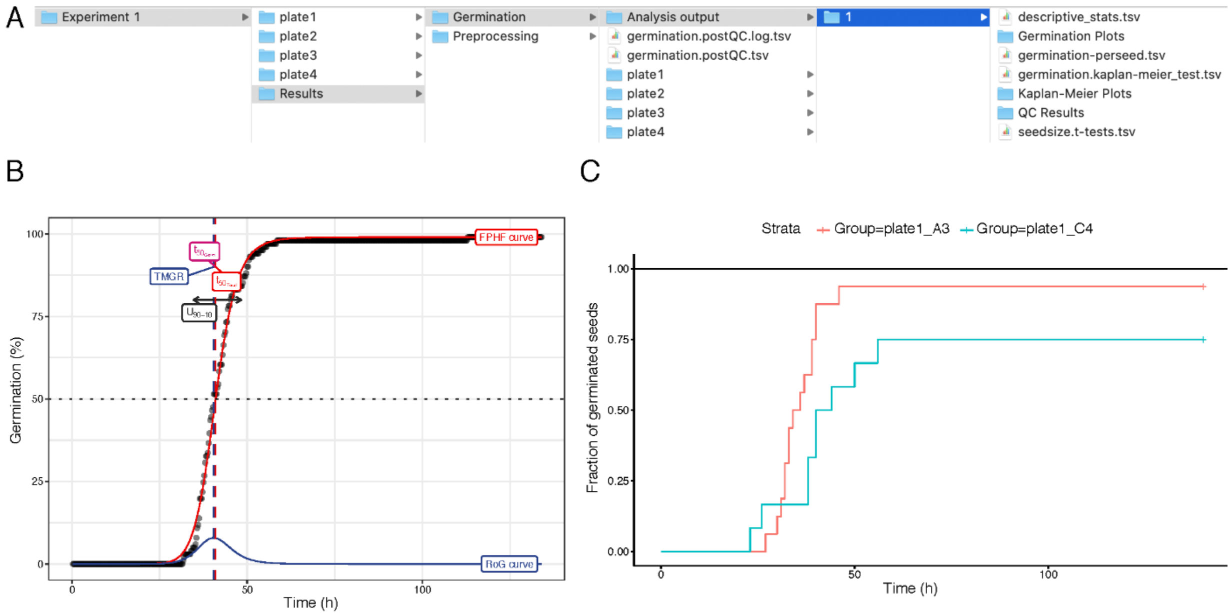


**Figure 8. Results of analysis output from the germination R scripts.**

(**A**) The results of the cleanup_germination_data.R script will be saved in two tsv files: germination.postQC.log.tsv and germination.postQC.tsv. The latter one can be used to manually modify group names while changing comparisons for the statistical analysis by process_germination_data.R. Results of each round of statistical analysis will be saved in a numbered subfolder within *Analysis* output folder. (**B**) Germination plot for individual group. FPHF curve, four-parameter Hill function curve; RoG curve, rate-of-germination curve; TMGR, time at maximum germination rate; T50 Germ, time required for 50% of germinated seeds to germinate; T50 total, time required for 50% of total seeds to germinate. Importantly, for this chart it is calculated accounting only viable seeds for the group, while the mean germination time provided in the statistical analysis in the .tsv files is calculated for all seeds detected for the group; $U_{90-10}$, uniformity defined as the time between the timepoints where 10% and 90% of seeds are germinated. Detailed description of the parameters is available in the germinationmetrics documentation (**C**) Kaplan–Meier "survival curve" plot shows proportion of germinated seeds for each group plotted against time. The test compares mean germination time calculated for all seeds present in the group. Results of the test are summarized in the file germination.kaplan-meier_test.tsv.

#### 4.6 SPIRO Root Growth Assay

##### *4.6.1 Overview*

This assay comprises automated image analysis (detection of each seed and root on the time-lapse images), and subsequent data quality control followed by germination detection, tracking of root length increase, building predictive models for root growth for groups of seedlings and finally, statistical analysis.

The workflow comprises a number of user-guided and automated steps. First, the designated ImageJ macro will prompt the user to select a subset of images representing the time interval of interest. Second, it will ask the user to select groups of seedlings that should be analyzed together, e.g., the same genotype. Third, it will prompt the user to manually adjust automated recognition of seeds. This step allows removing outliers or adding seeds missed by image segmentation. The duration of the image analysis will largely depend on the number of time-points, number of seedlings and RAM of the computer. The user-guided steps typically take less than an hour. The rest of the image analysis is automated and might take up to several hours. If for some reason, the analysis was not complete, rerunning the macro on the same experiment folder will resume from the step at which analysis was aborted.

For each selected group, the macro will identify the position of each individual seed and the border between shoot and root (further referred to as *root start coordinate*, *rsc*) in the time-lapse data and records individual seedlings root lengths for each time point. The macro output is subsequently processed by two R scripts.

Root growth data is cleaned using the following algorithm: a 5-slice left-aligned moving average is constructed from the raw primary root length, in order to remove fluctuations inherent to the measurement of root length. For each timepoint, the difference in moving average relative to the preceding timepoint is calculated, and the sign of the change recorded (i.e., –1 for a negative change, +1 for a positive change, and 0 for no change). The root is considered to have plateaued if the sign is 0 for 7 consecutive slices. Plateauing is generally caused by the root growing outside of its ROI. A "root length jump" is detected when the absolute change in root length between two consecutive timepoints is larger than 0.5 cm. "Jumps" are generally caused by two roots crossing. Both plateaus and jumps lead to truncation of data at the point of occurrence. Root growth is recorded relative to the seeds' germination time, with germination detection considered $t = 0$, in order to normalize root growth rates between seedlings.

For comparison of the root growth performance of different seed types, a mixed-effect second-order polynomial model is fitted using the R package glmmTMB[7]. From the estimated model, the average growth slope of each seedling group is compared using the R package emmeans. The model consists of two parts: a fixed part with an intercept and second order polynomials for each seedling group, and a random part allowing the intercept and the polynomial coefficients to vary over the individual seedlings to properly treat the variation between seedlings that is not due to the seedling group as random variation in the post-hoc average slope comparison between seedling groups. The choice of a second degree polynomial model is motivated by assessment of growth curves from previous experiments, properties of the growth curve such as monotonicity, but not linearity, and computational issues with more complex models.

##### *4.6.2 Using the Root Growth Macro*

a. Make sure that all installation steps are complete (see Section 3.3.2).

b. Install SPIRO_Macros.ijm as described in 3.3.2d and open the SPIRO_Germination macro (*Plugins → Macros → SPIRO_RootGrowth)*.

c. In the file picker, open the Experiment folder with the preprocessed data.

d. Follow the instructions provided by the macro:

- Reducing the number of slices/time points included into analysis allows omitting unnecessary time point where the roots might overlap. Please note that the analysis requires image data for a few hours prior to seeds germinating.

- You will be prompted to select groups of seedlings, e.g., different genotypes and/or treatments. This step is crucial for downstream statistical analysis of the quantitative data. If needed, grouping can be modified after data quality control (see Section 4.7) to enable different comparisons without re-running image analysis.

- While selecting the groups of seedlings that will be analyzed together, avoid including areas with reflections or imperfections on the medium surface and empty space.

- Selected groups can be later further rearranged for statistical analysis.

e. The macro will suggest to use the same selection/names for groups for all plates. The area and the group names can be manually edited and adjusted.

f. The macro will automatically detect seeds on the first slice of the time-lapse stack and prompt the user to verify the automated selection by showing the first and the last slices of the user selected time range. At this step you may deselect any seeds/seedlings that you do not wish to include into analysis and manually select any seeds missed by the automated image segmentation.

g. Results of the analysis for each group files will be saved in the experiment folder (**Fig. 9A**):

Experiment name/Results/Root Growth/plate*n*/*group name*

h. The macro will produce four output files for each group of seeds:

- group name rootgrowthdetection.tif is the graphical output of the macro (**Fig. 9B**). It contains time-lapse side by side comparison of original and results of image segmentation for the corresponding group with numbered identified seedlings. This information can be helpful to verify that accuracy of image processing.
- group name rootgrowthmeasurement.tsv is the quantitative output of the macro. It contains root length for each seedling over time.
- group name germination analysis.tsv contains perimeter and area data for each seed at each time point, which will be used for identifying the germination time point for each seed.

i. Image processing using Root Growth macro might be a lengthy procedure, depending on the size of data and RAM of the computer. To prevent loss of time, we introduced a run recovery feature. If *Root Growth* macro run was interrupted, upon restart the script will automatically detect the last completed analysis step and recover the run from the following step.

j. For troubleshooting, please refer to this table.

k. Please submit reports about errors and suggestions for optimizations as a GitHub issue.

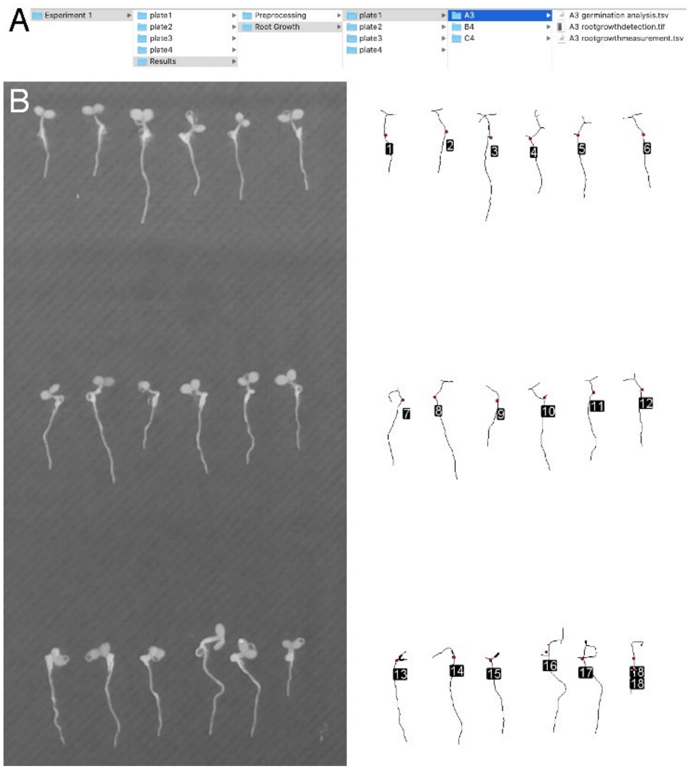


**Figure 9. Output of the root growth macro.**
(**A**) The results of the root growth macro will be saved in the subfolder Results/Root Growth/platen/group_name. The file group_name rootgrowthdetection.tif containing graphical output of root growth estimation, group_name rootgrowthmeasurement.tsv, contains quantitative results of root growth estimation, and group_name germination analysis.tsv, contains quantitative data about size and perimeter changes for each seed over time. (**B**) An example of a single frame from a group_name rootgrowthdetection.tif stack file, which show side by side raw image used for the analysis (left side) and results of this image segmentation (right side). Detected seedlings are presented as skeletonized lines, red dots denote position of the detected root start, the numbers indicate the number of a seedling that can be tracked to the the data in the .tsv file.

##### *4.6.3 Using Root growth R scripts*

a. Make sure that all installation steps are complete (see Section 3.3.2).

b. Open RStudio.

c. If the SPIRO Assays project is not open, go to *File → Open Project*. In the file picker, locate the project that you saved during installation steps (see Section 3.3.2).

d. Open cleanup_germination_data.R (click on the corresponding file name in Files tab, the right bottom panel of R studio).

e. Run the script: in the top menu find *Code → Source.* Select your experiment folder when prompted. For Windows users, this can be done via file picker (please note, that R Studio may place this window behind the R Studio window). For MacOS users, right click on your experiment folder and hold the Option (⌥) key and select the "Copy *path* as Pathname" option, then copy paste the path to your experiment folder into the console of R Studio.

f. In the Console, messages indicating what seeds were removed from the final output file, and for what reason, will be printed.

g. Results of the QC step are saved in two files in the experiment folder (for description see 4.5.3g):

   Experiment name/Results/Root Growth/germination.postQC.log.tsv
   Experiment name/Results/Root Growth/germination.postQC.tsv.

h. Run process_germination_data.R (click on the corresponding file name in the right bottom panel of R studio, then *Code → Source*). If needed, open the project again (see 4.5.3c).

i. Point the script to the experiment folder (see 4.5.3d)

j. The script will output the file germination-perseed.tsv to the Results/Root Growth folder. This file contains germination times for each seedling. These values are used for estimating the initiation of root growth.

k. After running germination QC and processing, proceed with the root growth analysis. Run consolidate_rootgrowth_data.R.

l. Inspect the Pre-analysis graphs in the directory Results/Root Growth/Pre-Analysis Graphs. These graphs can indicate problems such as misidentified germination. If needed, manually correct the values for germination by editing the file germination-perseed.tsv and rerun the script. The graphs will also indicate any measurement error not filtered during consolidation, and suggest problematic seedlings which may be omitted from analysis (see Section 4.7)

m. If needed, adjust grouping for the analysis and exclude mismeasured seedlings (see Section 4.7).

n. Run the script process_rootgrowth_data.R.

o. Analysis can be performed several times on the same experiment, and results of each analysis run will be saved in a separate directory in the Analysis results directory (**Fig. 10A**):

- Model prediction overview.pdf contains a graphical overview of the fitted model (**Fig. 10B**). This figure shows the model fit for each group, overlaid on the raw data. The line indicates the predicted root length versus time for each group, and the shaded area indicates the prediction interval of the model.

- Pairwise p-value plot.pdf visually indicates the Tukey-adjusted *p* values for each comparison.

- Predicted Root Lengths.tsv contains the predicted primary root lengths (in cm) at 24-h intervals.

- Predicted Root Growth Rates.tsv contains the predicted root growth rates (in µm/h) at 24 h intervals.

- Root growth rate barchart.pdf and Root length barchart.pdf contain illustrative charts depicting predicted root growth rates and lengths (Fig. 10C,D).

- Tukey CLD.tsv contains the Tukey compact letter display (CLD) values for comparing all groups. Non-shared numbers indicate that the groups are different at the $p < 0.05$ level.

p. For running custom statistical analyses, the file rootgrowth.postQC.tsv contains all data that is used for fitting the model. This file can be analyzed using any statistical software, not just R. A copy of this file is included in each Analysis results subdirectory, indicating the state (e.g., seedling grouping) used for the particular analysis.

q. For troubleshooting, please refer to this table.

r. Please submit reports about errors and suggestions for optimizations as a GitHub issue.

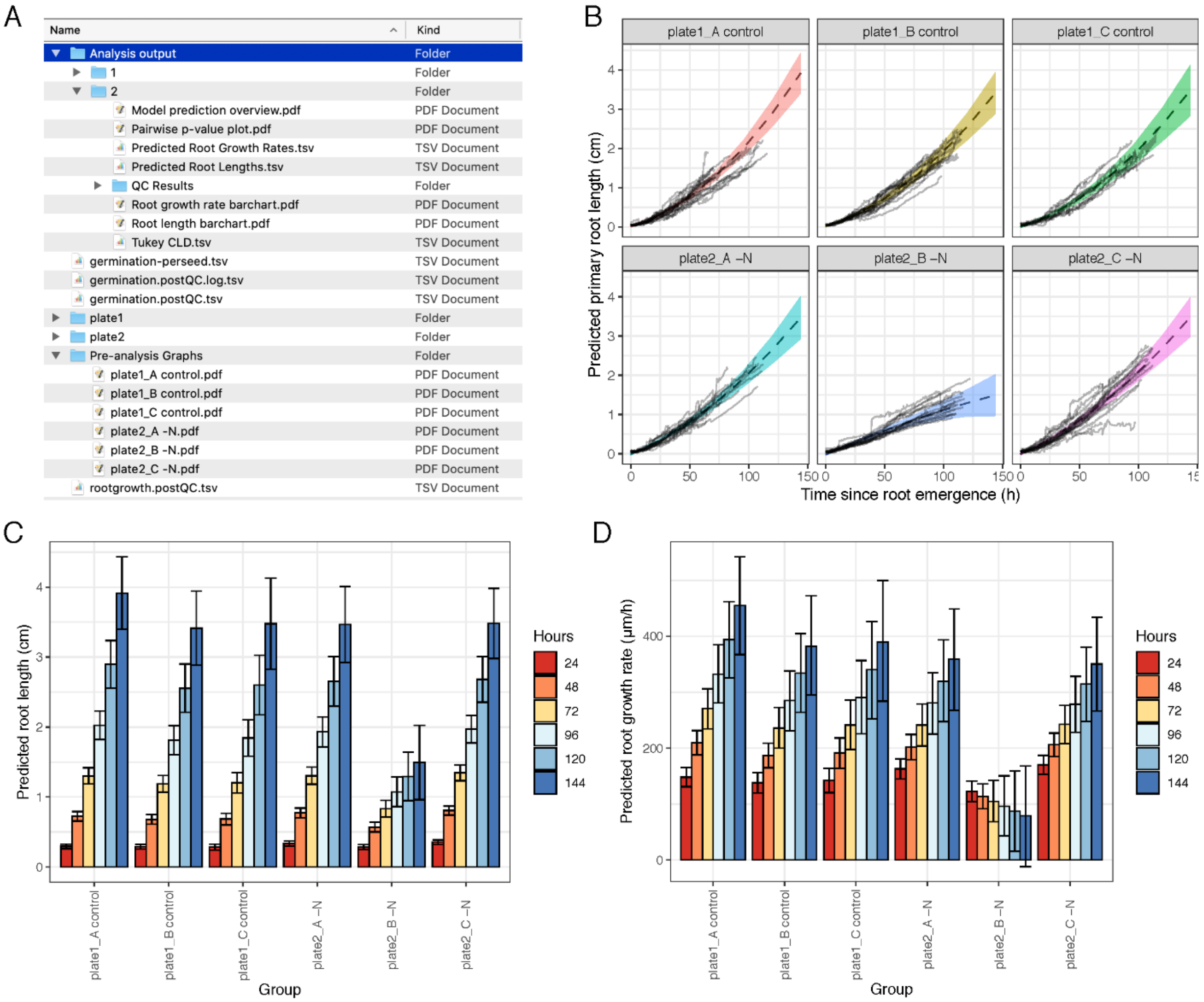


**Figure 10. Output from the root growth R scripts.**

**(A)** Folder structure after running all R scripts, showing locations of the pre-analysis graphs and the analysis output folders. **(B)** Model prediction overview indicating the model fit with standard error for each group. **(C)** Predicted root lengths at 24 h intervals for each group. **(D)** Predicted root growth rates at 24 h intervals for each group.

#### 4.7 SPIRO Assay Customizer

SPIRO Assay Customizer is a companion utility for merging and editing experimental data after quality control but prior to statistical analysis. The tool can:

- merge data from several experiments
- reassign seedlings to existing or new groups
- exclude seedlings from analysis

The two main modes, *Merge Assays* and *Customize Assay*, are described below.

##### *4.7.1 Merge assays*

In the Merge Assays mode, you can combine data from several independently analyzed experiments. Experiments are joined after performing data quality control and normalization, i.e. either cleanup_germination_data.R (for germination assay), or after running consolidate_rootgrowth_data.R (root growth assay). After choosing the Merge Assays mode, you are presented with a simple interface (**Fig. 11**).

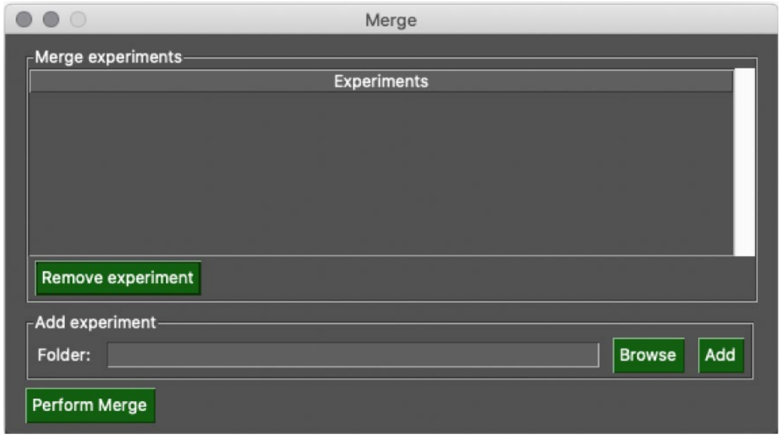


**Figure 11. SPIRO Assay Customizer *Merge Assays* mode.**
For each experiment you want to include in the merged output, click on *Browse*, and locate **either the *Root Growth* or *Germination* subfolder** of the *Results* subfolder of the main Experiment folder. After locating the folder, click *Add*. When all experiments are present in the *Merge Experiments* pane, click *Perform Merge*. The program will ask you to choose a folder. Create a new empty folder and select it. The merged data will be saved to this folder, and you may now run the appropriate processing script on the merged folder (or further customize the assay, see below).

##### *4.7.2 Customize assay*

In *Customize Assay* mode, you will be presented with the following window after opening a rootgrowth.postQC.tsv or germination.postQC.tsv file (**Fig. 12**). In the pane to the left, all included seedlings are displayed together with their current group assignment. In the right pane, the groups available for assignment are listed.

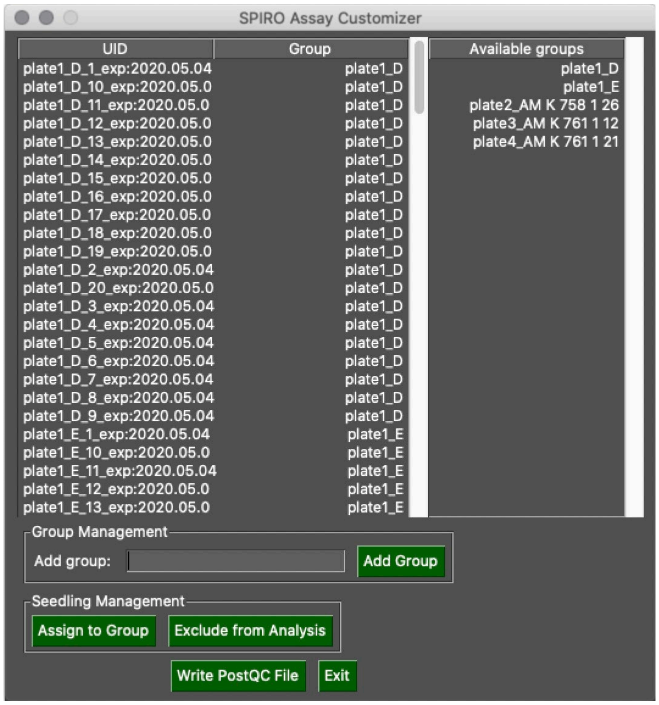


**Figure 12. SPIRO Assay Customizer *Customize Assay* mode.**

- **Adding a new group:** To make a new group available for assignment, type the name of the new group into the *Add group* textbox, then click the *Add Group* button. This makes the new group appear in the right pane.
- **Excluding a seedling from analysis:** To exclude a seedling, select it in the left pane and click *Exclude from Analysis*. Its group assignment will change to *NA*. Please note that several seedlings can be selected if either Shift or Control are held down during selection.
- **Reassigning seedlings to new groups:** With one or more seedlings selected in the left pane, and the desired group selected in the right pane, click *Assign to Group*. The assigned group will change.
- **Saving your changes:** Click on *Write PostQC File* to save your changes.

### Appendix 1: Quick start guides

#### Macro installation

Plugins → Install → select the SPIRO_macros.ijm file.

Install TurboReg and MultiStackReg into Fiji.

#### Preprocessing macro

**Input:** image data folder from SPIRO

1. Check if drift between images is present.
   Open your unprocessed images as a stack, by dragging and dropping a folder into the FIJI toolbar. Scroll through the stack to check for drift.
2. Plugins → Macros → SPIRO_Preprocessing
3. If drift is present, enable registration when prompted by macro. Otherwise, disable.
4. Follow instructions of the macro.

*Note: Preprocessing takes a lot of time and Fiji may look inactive. If in doubt, run the "Monitor Events" function of Fiji, if the macro is running, the Log window will print lines of currently running events. Close the Log window to cancel "Monitor Events" after checking.*

**Output:** plate*n*_preprocessed.tif

#### Germination macro

**Input:** plate*n*_preprocessed.tif

1. Plugins → Macros → SPIRO_Germination.
2. Select substack:
   **From** at least 2.5 hrs before germination of first seed.
   **To** time point well beyond (e.g., 12 h) when all seeds have germinated.
3. Select and name groups (e.g., seedlings from the same genotype, to be analysed as a group).
4. Verify automated seed detection by deleting wrongly detected seeds from ROI Manager, or adding ROIs around undetected seeds to the ROI Manager.
5. Allow macro to continue with automated analysis.

#### Root Growth macro

**Input:** plate*n*_preprocessed.tif

1. Plugins → Macros → SPIRO_RootGrowth
2. Select substack:
   **From** at least 2.5 hrs before germination.
   **To** the time point where data quality goes down due to too many seedlings that touch each other.
3. Select and name groups (e.g., seedlings from the same genotype).
4. Verify automated seed detection by deleting wrongly detected seeds from ROI Manager, or adding ROIs around undetected seeds to the ROI Manager.
5. Allow macro to continue with automated analysis.

**Output:** *groupname* rootgrowthdetection.tif; *groupname* rootgrowthmeasurements.tsv; and *groupname* germination analysis.tsv for each processed group.

#### Analysis in R

1. Create a new project with version control. In the menu, go to *File → New project→ Version control → Git*.
2. Input repository URL: https://github.com/jiaxuanleong/SPIRO.Assays.
3. Type in the desired directory name (e.g., SPIRO.Assays).
4. Pick the location suitable for keeping the scripts and macros for the assays (e.g., the user Documents folder).
5. Run the scripts in the following order: cleanup_germination_data.R, process_germination_data.R, and for the root growth assay, additionally run consolidate_rootgrowth_data.R followed by process_rootgrowth_data.R.

#### References

1. https://github.com/AlyonaMinina/SPIRO.Hardware.

2. https://github.com/jonasoh/spiro.

3. https://github.com/jonasoh/spiro-assay-customizer/releases.

4. https://github.com/jiaxuanleong/SPIRO.Assays.

5. Bewley, J. D. Seed Germination and Dormancy. *Plant Cell* **44**, 1055–1066 (1997).

6. Aravind, J., Vimala Devi, S., Radhamani, J., Jacob, S. R. & Kalyani, S. Germination-metrics: Seed Germination Indices and Curve Fitting. R package version 0.1.3.9000. (2020).

7. Brooks, M. E. *et al.* glmmTMB balances speed and flexibility among packages for zero-inflated generalized linear mixed modeling. *R Journal* **9**, (2017).
