## Supplementary File 5 for "SPIRO – the automated Petri plate imaging platform designed by biologists, for biologists": cleanup_germination_data.R

```
# cleanup_germination_data.R -
#
#   imports data from germination macro (or root growth macro germination detection) and cleans it up a bit.
#   after running this script, adjust the groups in the file germination.postQC.tsv, 
#   then run process_germination_data.R to get statistics.

# clean slate
rm(list=ls())
source('R scripts/common.R')
p_load(dplyr, foreach, doParallel, data.table, zoo, RcppRoll, rlang)

# below are cutoffs for area filtering
upper_area_threshold = 0.02
lower_area_threshold = 0.002

# main function for extracting data from files
processfile <- function(file, logdir, expname) {
  r <- SeedPos <- Date <- ImgSource <- startdate <- ElapsedHours <- plates <- DayNight <- NULL
  dirparams <- unlist(strsplit(dirname(file), '/', fixed=T))
  plate <- dirparams[length(dirparams)-1]
  group <- dirparams[length(dirparams)]
  
  resultfile <- fread(file)
  resultfile <- dplyr::select(resultfile, -V1)
  
  if ("X1" %in% names(resultfile)) {
    resultfile <- dplyr::select(resultfile, -X1)
  }
  resultfile <- as.data.frame(resultfile)

  for (i in 1:nrow(resultfile)) {
    row <- resultfile[i,]
    # first, go through the file and make a list of rois and timepoints
    params <- unlist(strsplit(row$Label, ':', fixed=T))
    
    if (length(params) == 3) {
      # assume that if the label contains two colons (i.e. 3 extracted elements), it is the initial image
      # for that seed, i.e., we are starting with a new seed here. set initial parameters like seed no etc.
      r <- params[2]
      SeedPos[[i]] <- r
      d <- startdate <- getdate(params[3])
      Date[[i]] <- as.character(d)
      DayNight[[i]] <- getdaynight(params[3])
    } else {
      # this is not the first record for a seed
      SeedPos[[i]] <- r
      d <- getdate(params[2])
      Date[[i]] <- as.character(d)
      DayNight[[i]] <- getdaynight(params[2])
    }
    ElapsedHours[[i]] <- elapsed(startdate, d)
  }
  data <- NULL
  data$UID <- paste0(plate, '_', group, '_', SeedPos, '_exp:', expname)
  data$Group <- paste0(plate, '_', group)
  resultfile <- dplyr::select(resultfile, -Label)
  data <- cbind(data, resultfile, SeedPos=unlist(SeedPos), Date=unlist(Date), ElapsedHours=unlist(ElapsedHours), 
                DayNight=unlist(DayNight))
  data <- check_duplicates(data, paste0(logdir, '/', basename(file), '.log'))
  return(data)
}

check_duplicates <- function(data, logfile) {
  # check for rois with duplicate measurements
  error_uids <- error_types <- NULL
  
  for(uid in unique(data$UID)) {
    # check data per uid (will only be one group per file)
    # ds = data subset
    ds <- data[which(data$UID == uid),]

    # trim data to first occurence of large area
    largearea <- which(ds$Area > upper_area_threshold)
    if (length(largearea)) {
      ds <- ds[1:largearea[1]-1,]
      if(largearea[1] < 50) {
        error_uids <- c(error_uids, uid)
        error_types <- c(error_types, 'EARLY_LARGE_AREA')
      }
    }
    
    # filter out large and small objects
    ds %>% filter(Area >= lower_area_threshold) %>% filter(Area <= upper_area_threshold) -> ds
    
    # we need to check if there are multiple measurements left for any slice
    ds %>% group_by(Slice) %>% mutate(n=n()) -> ds
    
    # truncate data after first occurrence of n>1 (keep first slice)
    if (length(which(ds$n > 1) > 0) & !is_empty(which(ds$n > 1))) {
      ds <- ds[1:max(which(ds$n > 1)[1] - 1, 1),]
    }

    # remove this uid from data, so we can add modified subset in its place
    data <- data[data$UID != uid,]

    # if there are anomalous objects in the first slice, remove the seed from analysis
    if (length(ds$n) > 0) {
      if(max(ds$n) > 1) {
        error_uids <- c(error_uids, uid)
        error_types <- c(error_types, 'DUPE')
      } else {
        ds <- dplyr::select(ds, -n)
        # need to cast to data frame or this fails
        data <- rbind(as.data.frame(data), as.data.frame(ds))
      }
    }
  }
  
  if (length(error_uids > 0)) {
    errors <- data.frame(UID=error_uids, Type=error_types)
    fwrite(errors, file=logfile, sep='\t')
  }
  
  return(data)
}

dir <- choose_dir()
resultsdir <- paste0(dir, '/Results')
germdir <- paste0(resultsdir, '/Germination')
rootdir <- paste0(resultsdir, '/Root Growth')

# get all matching .tsv files in the directories
germfiles <- list.files(path = germdir, pattern = ' germination analysis.tsv$', full.names = TRUE, recursive = TRUE, ignore.case = TRUE, no.. = TRUE)
rootfiles <- list.files(path = rootdir, pattern = ' germination analysis.tsv$', full.names = TRUE, recursive = TRUE, ignore.case = TRUE, no.. = TRUE)

if (length(rootfiles) == 0 & length(germfiles) > 0) {
  outdir <- germdir
  files <- germfiles
} else if (length(germfiles) == 0 & length(rootfiles) > 0) {
  outdir <- rootdir
  files <- rootfiles
} else if (length(germfiles) == 0 & length(rootfiles) == 0) {
  stop("No germination data found in that directory.")
} else {
  stop("Germination data found in both Germination and Root Growth folders. This script needs germination data from exactly one assay.")
}

allout <- toolarge <- err_largeobj <- err_multiobj <- NULL

if (!exists('germination.debug')) {
  num_cores <- max(1, detectCores() - 1)
  cl <- makeCluster(num_cores)
  registerDoParallel(cl)
  
  if (num_cores > 1) {
    core_plural <- 'threads'
  } else {
    core_plural <- 'thread'
  }
  expname <- basename(dir)
  cat(paste0("Performing germination QC for experiment << ", expname, " >>\n\n"))
  cat(paste0("Processing files and performing basic quality control, using ", 
             length(cl), ' ', core_plural, "...\n"))
  
  allout <- foreach(f=files, .combine=rbind, .multicombine=T, .packages=c('dplyr', 'data.table', 'zoo', 'rlang')) %dopar%
    processfile(f, outdir, expname)
  stopCluster(cl)
} else {
  allout <- NULL
  expname <- basename(dir)
  for (f in files) {
    allout <- rbind(allout, processfile(f, outdir, expname))
  }
}

# check the logs
logfiles <- list.files(path = outdir, pattern = ' germination analysis.tsv.log$', full.names = TRUE, recursive = TRUE, ignore.case = TRUE, no.. = TRUE)
log <- NULL
for (f in logfiles) {
  logfile <- fread(f)
  log <- rbind(log, logfile)
  file.remove(f)
}

for (errtype in unique(log$Type)) {
  if (errtype == 'EARLY_LARGE_AREA') {
    cat('Large non-seed object detected in ROI. Time range has been truncated.\n')
    cat('Affected seeds:\n ')
    cat(paste0(log$UID[log$Type == 'EARLY_LARGE_AREA'], '\n'))
    cat('\n')
    err_largeobj <- data.frame(UID=log$UID[log$Type == 'EARLY_LARGE_AREA'], Note='Large non-seed object detected in ROI. Time range was truncated.')
  } else if (errtype == 'DUPE') {
    cat('Multiple objects remained after filtering. Affected seeds were removed from analysis.\n')
    cat('Affected seeds:\n ')
    cat(paste0(log$UID[log$Type == 'DUPE'], '\n'))
    cat('\n')
    err_multiobj <- data.frame(UID=log$UID[log$Type == 'DUPE'], Note='Multiple objects remained after filtering. Seed was removed from analysis.')
  }
}
errors <- rbind(err_largeobj, err_multiobj)
errors$UID <- as.character(errors$UID)
errors$Note <- as.character(errors$Note)

# round off ElapsedHours to ensure compatibility with spreadsheet software
allout$ElapsedHours <- round(allout$ElapsedHours, 4)

# rearrange columns
allout <- allout[,c(1:2, 8, 3:7, 9)]

fwrite(allout, file=paste0(outdir, "/germination.postQC.tsv"), sep='\t')
if (!grepl('Root Growth$', outdir)) {
  cat(paste0("Saving cleaned and collated data to '", outdir, "/germination.postQC.tsv", "'.\nPlease edit that file to set up correct grouping for your experiment.\n"))
} else {
  prominent_message("Now run process_germination_data.R.")
}

# create error log
seedlog <- data.frame(UID=unique(allout$UID), Note='Seed processed normally.')
seedlog$UID <- as.character(seedlog$UID)
seedlog$Note <- as.character(seedlog$Note)
seedlog <- merge(seedlog, errors, by="UID", all=T)
seedlog$Note.y <- as.character(seedlog$Note.y)
seedlog %>% mutate(Note = coalesce(Note.x, Note.y)) %>% dplyr::select(c(UID, Note)) -> seedlog

fwrite(seedlog, file=paste0(outdir, "/germination.postQC.log.tsv"), sep='\t')
```
