## Supplementary File 5 for "SPIRO – the automated Petri plate imaging platform designed by biologists, for biologists": common.R

```
# common.R -
# some functions common to other scripts

options(dplyr.summarise.inform = FALSE)

if (!require('pacman')) {
  install.packages('pacman')
  library(pacman)
}

spiro_common_included <- function() {
  return(TRUE)
}

choose_dir <- function() {
  # there is no support for directory picker under non-windows platforms
  if (.Platform$OS.type == 'unix') {
    dir <- readline(prompt = "Enter directory: ")
  } else {
    dir <- choose.dir(getwd(), "Choose folder to process")
  }
  return(dir)
}

create_rundir <- function(outdir) {
  # set up output dir
  if (dir.exists(paste0(outdir, '/Analysis output'))) {
    runs <- list.dirs(paste0(outdir, '/Analysis output'), full.names=F, recursive=F)
    run_number <- length(runs) + 1
    while (file.exists(paste0(outdir, '/Analysis output/', run_number))) {
      run_number <- run_number + 1
    }
  } else {
    dir.create(paste0(outdir, '/Analysis output'), recursive=T)
    run_number <- 1
  }
  
  rundir <- paste0(outdir, '/Analysis output/', run_number)
  dir.create(rundir, showWarnings=F)
  return(rundir)
}

# return elapsed time in hours from two datetime strings
elapsed <- function(from, to) {
  f <- strptime(from, format="%Y-%m-%d %H:%M:%S")
  t <- strptime(to, format="%Y-%m-%d %H:%M:%S")
  return(as.numeric(difftime(t, f, units='hours')))
}

# extract datetime from strings such as "plate1-20190602-082944-day"
getdate <- function(name) {
  params <- unlist(strsplit(name, '-', fixed=T))
  x <- paste0(params[2:3], collapse='')
  return(strptime(x, format='%Y%m%d%H%M%S'))
}

# extract day/night from filename
getdaynight <- function(name) {
  return(unlist(strsplit(name, '-', fixed=T))[4])
}

# show prominent message
prominent_message <- function(message) {
  cat(paste0(rep('*', nchar(message)), collapse=''))
  cat(paste0('\n', message, '\n', collapse=''))
  cat(paste0(rep('*', nchar(message)), collapse=''))
}
```
