## Supplementary File 5 for "SPIRO – the automated Petri plate imaging platform designed by biologists, for biologists": consolidate_rootgrowth_data.R

```
# consolidate_rootgrowth_data.R -
# 
#   cleans up root growth data, and merges it with germination data

# clean slate
rm(list=ls())
source('R scripts/common.R')

p_load(dplyr, ggplot2, ggrepel, patchwork, zoo, data.table)

# ggplot theme
th <- theme_bw() + theme(legend.position="bottom", legend.text=element_text(size=8))

# function for plotting unprocessed data
plotfile <- function(file) {
  r <- fread(file)
  names(r) <- c('Slice', 'Label', 'Rootno', 'Length')
  d <- dirname(file)
  dirparams <- unlist(strsplit(d, '/', fixed=T))
  r$GID <- paste0(dirparams[length(dirparams)-1], '_', dirparams[length(dirparams)])
  r$UID <- paste0(r$GID, '_', r$Rootno)
  params <- unlist(strsplit(r$Label, '-', fixed=TRUE))
  r$date <- as.POSIXct(strptime(paste0(params[seq(2, length(params), 4)], params[seq(3, length(params), 4)]), format='%Y%m%d%H%M%S'))
  
  # add elapsed times
  r %>% group_by(UID) %>%
    arrange(date) %>%
    mutate(elapsed=elapsed(date[1], date)) -> r

  # get rid of divide-by-zero
  r.tmp <- r[r$Slice > 1,]
  TimePerSlice <- mean(r.tmp$elapsed / (r.tmp$Slice-1))
  
  return(ggplot(r, aes(x=elapsed, y=Length, color=UID, group=UID)) + 
           geom_point() + 
           geom_line() +
           scale_x_continuous(sec.axis=sec_axis(~./TimePerSlice, breaks=seq(min(r$elapsed/TimePerSlice), max(r$elapsed/TimePerSlice), 20), name="Slice")) +
           labs(title=paste0("Unprocessed graph for ", r$GID), x="Elapsed time (h)", y="Root length (cm)") + th +
           geom_label_repel(data=r %>% group_by(UID) %>% arrange(elapsed) %>% summarize(elapsed=last(elapsed), Length=last(Length)), 
                            aes(label=UID)) +
           theme(legend.position="none"))
}

processfile <- function(file, expname) {
  r <- fread(file)
  names(r) <- c('Slice', 'Label', 'Rootno', 'Length')
  d <- dirname(file)
  dirparams <- unlist(strsplit(d, '/', fixed=T))
  r$GID <- paste0(dirparams[length(dirparams)-1], '_', dirparams[length(dirparams)])
  r$UID <- paste0(r$GID, '_', r$Rootno, '_exp:', expname)
  params <- unlist(strsplit(r$Label, '-', fixed=TRUE))
  r$date <- as.POSIXct(strptime(paste0(params[seq(2, length(params), 4)], params[seq(3, length(params), 4)]), format='%Y%m%d%H%M%S'))
  
  # find out which seeds were processed normally by germination script
  r %>% filter(UID %in% normseeds) -> r
  
  # add elapsed times
  r %>% group_by(UID) %>%
    arrange(date) %>%
    mutate(elapsed=elapsed(date[1], date)) -> r

  r %>% mutate(delta = Length - lag(Length)) -> r

  r %>% arrange(elapsed) %>%
    group_by(UID) %>%
    mutate(mablright = rollapply(Length, 5, mean, align="left", fill=NA),
           mabldelta = mablright - lag(mablright),
           mablsign = sign(mabldelta)) -> r
  
  r %>% arrange(elapsed) %>%
    group_by(UID) %>%
    mutate(signsum = rollapply(mablsign, 9, sum, na.rm=T, align="left", fill=NA)) -> r

  # use germination data to assess root growth
  r %>% group_by(UID) %>%
    filter(Slice >= germtimes$slice[germtimes$UID == UID[1]]) -> r

  # construct a normalized elapsed time value based on root growth start
  r %>% arrange(elapsed) %>%
    group_by(UID) %>% 
    mutate(normtime = elapsed - elapsed[1], dbg=elapsed[1]) -> r

  # identify plateaus:
  # if there is exactly zero growth for 7 consecutive timepoints, it is a plateau
  r %>% arrange(elapsed) %>%
    group_by(UID) %>%
    mutate(still=rollapply(mablsign, 7, function(x) { return(all(x == 0)) }, align="left", fill=NA)) -> r
  
  # truncate after plateau detected
  r$still[which(r$still == FALSE)] <- NA
  r %>% arrange(elapsed) %>%
    group_by(UID) %>%
    mutate(still=na.locf(still, na.rm=F)) -> r
  r$still[which(is.na(r$still))] <- FALSE
  r %>% filter(still!=TRUE) -> r

  # mark big jumps (length increase > 0.5 cm)
  r$jump <- NA
  r$jump[abs(r$delta) > 0.5] <- TRUE

  r %>% arrange(elapsed) %>%
    group_by(UID) %>%
    mutate(jump = na.locf(jump, na.rm=F)) -> r
  r$jump[which(is.na(r$jump))]<-FALSE
  r %>% filter(jump!=TRUE) -> r
  
  return(r)
}

dir <- choose_dir()

resultsdir <- paste0(dir, '/Results')
outdir <- paste0(resultsdir, '/Root Growth')
rundir <- paste0(outdir, '/Pre-analysis Graphs')
if (!dir.exists(rundir)) {
  dir.create(rundir)
}

# get all .tsv files in the directory
files <- list.files(path = outdir, pattern = ' rootgrowthmeasurement.tsv$', full.names = TRUE, recursive = TRUE, ignore.case = TRUE, no.. = TRUE)

if (length(files) < 1) {
  cat("No suitable root analysis results files found in that directory.\n")
  stop()
}

if (!file.exists(paste0(outdir, '/germination-perseed.tsv'))) {
  cat("germination-perseed.tsv not found. Run process_germination_data.R first.\n")
  stop()
}

allout <- NULL

expname <- basename(dir)
cat(paste0("Performing root growth QC for experiment << ", expname, " >>\n\n"))

cat("Processing files, please wait...\n")

germtimes <- fread(paste0(outdir, '/germination-perseed.tsv'))
names(germtimes)[3] <- 'slice'
normseeds <- germtimes$UID[!is.na(germtimes$slice)]
badseeds <- germtimes$UID[is.na(germtimes$slice)]
if(length(badseeds)) {
  cat("The following seedlings do not have a germination time, and are excluded from analysis:\n ")
  cat(paste0(badseeds, '\n'))
}

for (f in files) {
  # clean up file
  out <- processfile(f, expname)
  allout <- rbind(allout, out)
  
  # snip uids
  out$UID <- sub('_exp:.*$', '', out$UID)
  
  # plot unprocessed data
  d <- dirname(f)
  dirparams <- unlist(strsplit(d, '/', fixed=T))
  GID <- paste0(dirparams[length(dirparams)-1], '_', dirparams[length(dirparams)])
  p_before <- plotfile(f)
  out.tmp <- out[out$Slice > 1,]
  TimePerSlice <- mean(out.tmp$elapsed / (out.tmp$Slice-1))
  
  # plot processed data
  p_proc <- ggplot(out, aes(x=elapsed, y=Length, color=UID, group=UID)) + 
    geom_point() +
    geom_line() +
    scale_x_continuous(sec.axis=sec_axis(~./TimePerSlice, breaks=seq(as.integer(min(out$elapsed/TimePerSlice)), as.integer(max(out$elapsed/TimePerSlice)), 20), name="Slice")) +
    labs(title=paste0("Processed graph for ", GID), x="Elapsed time (h)", y="Root length (cm)") + th +
    geom_label_repel(data=out %>% group_by(UID) %>% arrange(elapsed) %>% summarize(elapsed=last(elapsed), Length=last(Length)), 
                     aes(label=UID)) +
    theme(legend.position="none")
  p_norm <- ggplot(out, aes(x=normtime, y=Length, color=UID, group=UID)) + 
    geom_point() +
    geom_line() +
    labs(title=paste0("Normalized-time graph for ", GID), x="Relative elapsed time (h)", y="Root length (cm)") + th +
    geom_label_repel(data=out %>% group_by(UID) %>% arrange(normtime) %>% summarize(normtime=last(normtime), Length=last(Length)), 
                     aes(label=UID)) +
    theme(legend.position="none")
  p <- p_before / p_proc / p_norm
  ggsave(p, filename=paste0(rundir, '/', GID, '.pdf'), width=25, height=30, units='cm')
}

allout %>% arrange(GID, UID, normtime) %>% 
  dplyr::select(c(UID, GID, elapsed, normtime, Length, date)) -> allout

names(allout)[2] <- 'Group'
names(allout)[3] <- 'ElapsedHours'
names(allout)[4] <- 'RelativeElapsedHours'
names(allout)[5] <- 'PrimaryRootLength'
names(allout)[6] <- 'Date'

fwrite(allout, file=paste0(outdir, "/rootgrowth.postQC.tsv"), sep='\t')
cat(paste0("Output saved to ", outdir, "/rootgrowth.postQC.tsv", 
           ". Adjust groups and remove problematic seedlings from this file, then run process_rootgrowth_data.R.\n"))
cat(paste0("Graphs are available for each group in the directory ", rundir, "\n"))
```
