## Supplementary File 5 for "SPIRO – the automated Petri plate imaging platform designed by biologists, for biologists": process_germination_data.R

```
# process_germination_data.R -
#
#   takes data generated by cleanup_germination_data.R and estimates germination times.
#   statistics will be output as tsv files in the data folder.
#   additionally, a graph indicating germination progression is generated for each group.

# clean slate
rm(list=ls())
source('R scripts/common.R')

p_load(dplyr, ggplot2, survival, survminer, RcppRoll, data.table, doParallel)
p_load_gh('aravind-j/germinationmetrics')

kmplot <- function(grp1, grp2, mydata, outdir) {
  ds <- subset(mydata, with(mydata, Group==grp1 | Group==grp2))
  mysurvobj <- Surv(time=ds$`Germination Time (h)`, event=ds$event)
  mysfit <- surv_fit(mysurvobj~Group, data=ds)
  p <- ggsurvplot(mysfit, pval=F, fun=function(y) { return(1-y) }) +
    labs(x='Elapsed Time (h)', y='Fraction of germinated seeds')
  suppressWarnings(ggsave(p$plot + geom_hline(yintercept=1), 
                          filename=paste0(rundir, "/Kaplan-Meier Plots/KaplanMeier-", pvals$Group.1[i], "_vs_", pvals$Group.2[i], ".pdf"),
                          width=25, height=15, units="cm"))
  # get p-value (standard log-rank)
  return(surv_pvalue(mysfit)$pval)
}

dir <- choose_dir()

resultsdir <- paste0(dir, '/Results')
germdir <- paste0(resultsdir, '/Germination')
rootdir <- paste0(resultsdir, '/Root Growth')

if (file.exists(paste0(germdir, "/germination.postQC.tsv")) & !file.exists(paste0(rootdir, "/germination.postQC.tsv"))) {
  outdir <- germdir
} else if (!file.exists(paste0(germdir, "/germination.postQC.tsv")) & file.exists(paste0(rootdir, "/germination.postQC.tsv"))) {
  outdir <- rootdir
} else if (!file.exists(paste0(germdir, "/germination.postQC.tsv")) & !file.exists(paste0(rootdir, "/germination.postQC.tsv"))) {
  stop("germination.postQC.tsv not found in specified directory. Did you run cleanup_germination_data.R?\n")
} else {
  stop("germination.postQC.tsv found in both Germination and Root Growth directories. This script needs germination data from exactly one assay.")
}

data <- fread(paste0(outdir, "/germination.postQC.tsv"))
rundir <- create_rundir(outdir)

# copy postQC input file to rundir for reference
dir.create(paste0(rundir, '/QC Results'))
file.copy(paste0(outdir, "/germination.postQC.tsv"), paste0(rundir, '/QC Results'))
if (file.exists(paste0(outdir, "/germination.postQC.log.tsv"))) {
  file.copy(paste0(outdir, "/germination.postQC.log.tsv"), paste0(rundir, '/QC Results'))
}

cat('Processing germination data...\n')

# if Group is set to NA, do not include in analysis
data <- data[which(!is.na(data$Group)),]

data$dPerim <- data$ddPerim <- data$Germinated <- 0
groups <- uids <- perims <- NULL

# average elapsed time per slice
avgTimePerSlice <- data[Slice > 1, mean(ElapsedHours / (Slice-1))]

# range for averages is either 2.5 h, or 5 slices (whichever is highest)
rollrange <- round(max(5, 2.5 / avgTimePerSlice), 0)

# number of positive ddPerims is either 5 h or 10 slices
ddprange <- round(max(10, 5 / avgTimePerSlice), 0)

# get number of slices available per uid
nrows <- data[, list(rows=nrow(.SD)), by=UID]

# calculate seed size by uid
seedsizes <- data[, .(SeedSize=mean(Area[1:rollrange])), by=.(UID, Group)]

# correct for day/night difference in perimeter
# - we want to use data only during the first 24 h to reduce risk of inflation due to early germination,
# - we want to avoid the very first slices since there may be imbibition going on,
# - and we want to use day/night slices that are adjacent to each other in time
#
# we assume ~8 h night and 24 h cycle duration but the algorithm might work for other values as well

# number day and night slices separately
data[, count:=seq_along(ElapsedHours), by=.(UID, DayNight)]

# get elapsed times for the first and last night slices of the first night
data[DayNight == 'night' & ElapsedHours < 24, .(nightstart=first(ElapsedHours), 
                                                nightend=last(ElapsedHours)), by=UID] -> ranges

ranges[, start := fcase(nightstart <= 4 & nightend > 20, 0,
                        # we have a split night, use all of the range
                        nightstart > 4 & nightstart <= 12, max(nightend - 8, 4),
                        # night is early
                        nightstart > 12, nightstart - 8), by=UID
                        # night is late
       ][, end := fcase(nightstart <= 4 & nightend > 20, 24,
                              nightstart > 4 & nightstart <= 12, nightend + 8,
                              nightstart > 12, min(nightend, nightstart + 8)), by=UID]

diffs <- data[, .(nightmean = mean(`Perim.`[DayNight=='night' & 
                                              ElapsedHours > ranges[UID==unlist(.BY)]$start &
                                              ElapsedHours < ranges[UID==unlist(.BY)]$end][1:16], na.rm=T),
                  daymean =   mean(`Perim.`[DayNight=='day' & 
                                              ElapsedHours > ranges[UID==unlist(.BY)]$start &
                                              ElapsedHours < ranges[UID==unlist(.BY)]$end][1:16], na.rm=T)), by=UID]

diffs$diff <- with(diffs, nightmean - daymean)

# we may be missing some values, e.g. if there is no data for first 24 h
x <- unique(data$UID)[! unique(data$UID) %in% diffs$UID]
if(length(x) > 0) diffs <- rbind(as.data.frame(diffs), data.frame(UID=x, diff=0, stringsAsFactors=F))
diffs$diff[!is.finite(diffs$diff)] <- 0

# subtract day/night difference from night perimeter values
data[DayNight == 'night', `Perim.` := `Perim.` - diffs$diff[diffs$UID == .BY$UID], by=UID]

# moving average perimeter increase
data[, dPerim := roll_meanl(`Perim.`, n=rollrange, na.rm=T, fill=NA) - mean(`Perim.`[1:rollrange]), by=UID]

# set delta delta perimeter (rate of change of perimeter)
data[, ddPerim := dPerim - shift(dPerim), by=UID][1, ddPerim := 0, by=UID]

# dPerim change over the ddP range
data[, ddPchange := shift(dPerim, n=ddprange, type="lead") - dPerim, by=UID]

# dPerim change over 2*ddP range
data[, ddPlongchange := shift(dPerim, n=ddprange*2, type="lead") - dPerim, by=UID]

# rolling average of the sign of ddPerim: basically a measure of the fraction of positive ddPerims
data[, sign_mean := roll_meanl(sign(ddPerim), n=ddprange, fill=NA), by=UID]

# require 0.8 mean sign and increased dPerim of 0.05 over the ddprange for germination
data[sign_mean>=0.8 & ddPchange>0.05 & ddPlongchange>0.1, Germinated := TRUE, by=UID]

# get the germination slice for each uid
germslices <- data[Germinated==TRUE, .(germslice=min(Slice)), by=UID]

# function to return exactly one instance of germination occurrence
set_germination <- function(ds, uid) {
  uid <- unlist(uid)[1]
  out <- rep(0, nrow(ds))
  germslice <- which(ds$Slice == germslices$germslice[germslices$UID == uid]) + 1
  if (! uid %in% germslices$UID) {
    return(out)
  } else if (germslice > nrow(ds)) {
    return(out)
  } else {
    out[germslice] <- 1
    return(out)
  }
}

# remove the old germination data and replace with the single-occurrence one
data[, Germinated := FALSE]
data[, Germinated := set_germination(.SD, .BY), by=UID]

data.peruid <- data %>%
  group_by(Group, UID) %>%
  filter(Germinated == 1) %>%
  arrange(ElapsedHours) %>%
  summarize(GerminationTime = ElapsedHours[1], GerminationSlice = Slice[1],
            SeedSize = seedsizes$SeedSize[seedsizes$UID==UID[1]])

seedsizes.pergroup <- seedsizes %>%
  group_by(Group) %>%
  summarize(SeedSizeSD = sd(SeedSize, na.rm=T), SeedSize = mean(SeedSize, na.rm=T)) %>%
  dplyr::select(Group, SeedSize, SeedSizeSD)

data.peruid <- as.data.frame(data.peruid)

uids <- unique(data$UID)
uids <- uids[order(uids)]
included <- uids %in% data.peruid$UID
for(uid in uids[!included]) {
  group <- data$Group[data$UID == uid][1]
  row = data.frame(Group = group, UID = uid, GerminationTime = NA, GerminationSlice = NA, SeedSize = NA)
  data.peruid <- rbind(data.peruid, row)
}

# sort data naturally
data.peruid$num <- gsub('^plate.*_([0-9]+)_exp:.*$', '\\1', data.peruid$UID)
data.peruid <- data.peruid[order(data.peruid$Group, data.peruid$num),]
data.peruid <- data.peruid %>% dplyr::select(-num)

names(data.peruid)[3] <- 'Germination Time (h)'
names(data.peruid)[4] <- 'Germination Detected on Frame'
names(data.peruid)[5] <- 'Seed Size (cm2)'

# combine with QC notes
if (file.exists(paste0(outdir, "/germination.postQC.log.tsv"))) {
  seedlog <- fread(paste0(outdir, "/germination.postQC.log.tsv"))
  data.peruid <- merge(data.peruid, seedlog, all=T)
  fwrite(data.peruid, file=paste0(rundir, "/germination-perseed.tsv"), sep='\t')
} else {
  data.peruid$Note <- NA
}

if (grepl('Root Growth$', outdir)) {
  rg <- data.peruid[,c(1,2,4,6)]
  fwrite(rg, file=paste0(outdir, "/germination-perseed.tsv"), sep='\t')
}

# merge group and uid so we can keep both in the conversion long->wide->long
data$GroupUID <- paste(data$Group, data$UID, sep="!")

# convert to wide and back again as a hacky way of making sure all seeds have equal number of slices
data.wide <- dcast(data, Slice ~ GroupUID, value.var = "Germinated")
data.long <- melt(data.wide, id.vars="Slice")

# now get all the groups and uids back
groupuids <- unlist(strsplit(as.character(data.long$variable), "!"))
data.long$Group <- groupuids[seq(1, length(groupuids), 2)]
data.long$UID <- groupuids[seq(2, length(groupuids), 2)]
data.long$value[is.na(data.long$value)] <- 0

# calculate germination stats
germstats <- data.long %>%
  group_by(Group, Slice) %>%
  summarize(GermCount = sum(value))

germstats <- germstats %>%
  group_by(Group) %>%
  arrange(Slice) %>%
  mutate(CumGermCount = cumsum(GermCount))

# to get the intervals in hours, we calculate average time per slice and multiply
# slice# by that values. this will hopefully be good enough approximation in most cases. 
germstats$ApproxTime <- round(germstats$Slice * round(avgTimePerSlice, 2), 2)

# now calculate some germination metrics
cat('Calculating germination statistics...\n')
groups <- t50s <- mgts <- mgtses <- nseeds <- nongerms <- NULL
dir.create(paste0(rundir, "/Germination Plots"))
for(group in unique(data.long$Group)) {
  # calculate t50 for summary file
  t50 <- t50(germ.counts = germstats$GermCount[germstats$Group == group], 
             intervals = germstats$ApproxTime[germstats$Group == group], 
             method = "coolbear")
  mgt <- MeanGermTime(germ.counts = germstats$GermCount[germstats$Group == group], 
             intervals = germstats$ApproxTime[germstats$Group == group])
  mgtse <- SEGermTime(germ.counts = germstats$GermCount[germstats$Group == group], 
                      intervals = germstats$ApproxTime[germstats$Group == group])
  nseed <- sum(germstats$GermCount[germstats$Group == group])
  nongerm <- sum(is.na(data.peruid$`Germination Time (h)`[which(data.peruid$Group == group)]))
  
  t50s <- c(t50s, t50)
  groups <- c(groups, group)
  mgts <- c(mgts, mgt)
  mgtses <- c(mgtses, mgtse)
  nseeds <- c(nseeds, nseed)
  nongerms <- c(nongerms, nongerm)
  
  # make germination graph
  pdf(paste0(rundir, "/Germination Plots/germinationplot-", group, ".pdf"), width=7, height=5)
  tryCatch(graph <- FourPHFfit(germ.counts = germstats$GermCount[germstats$Group == group], 
                      intervals = germstats$ApproxTime[germstats$Group == group],
                      total.seeds = length(unique(data$UID[data$Group == group])),
                      tmax = max(germstats$ApproxTime[germstats$Group == group])),
           error=function(e) {
             cat(paste0("Unable to generate germination plot for group ", group, ":\n"))
             cat(paste0(e$message, "\n"))
           })
  print(plot(graph, mgt=F) + scale_x_continuous(breaks=seq(0, max(germstats$ApproxTime[germstats$Group == group]), 24)) + labs(x="Time (h)"))
  dev.off()
}

germstats.pergroup <- data.frame(Group = groups, t50 = t50s, MeanGermTime = mgts, 
                                 MeanGermTimeSE = mgtses, n = nseeds, Ungerminated = nongerms)
germstats.pergroup <- merge(germstats.pergroup, seedsizes.pergroup)
names(germstats.pergroup) <- c('Group', 't50 (h)', 'Mean Germination Time (h)', 'Mean Germination Time SE',
                               'Germinated seeds', 'Ungerminated seeds', 'Seed Size (cm2)', 'Seed Size SD')
fwrite(germstats.pergroup, file=paste0(rundir, "/descriptive_stats.tsv"), sep='\t')

if (length(unique(data.long$Group)) > 1) {
  # generate table for t-tests
  pvals <- as.data.frame(t(combn(unique(data.long$Group), 2)))
  colnames(pvals) <- c('Group.1', 'Group.2')
  pvals$Group.1 <- as.character(pvals$Group.1)
  pvals$Group.2 <- as.character(pvals$Group.2)
  seedsize.pvals <- pvals
  
  for (i in seq(1, nrow(pvals))) {
    # first we check the p value
    tryCatch({ts <- t.test(data.peruid$`Seed Size (cm2)`[data.peruid$Group == pvals$Group.1[i]],
                           data.peruid$`Seed Size (cm2)`[data.peruid$Group == pvals$Group.2[i]])
              seedsize.pvals$p.value[i] <- ts$p.value
             },
             error=function(e) {
                cat(paste0("Unable to compare groups ", pvals$Group.1[i], "(x) to ", pvals$Group.2[i], "(y): "))
                cat(paste0(e$message, "\n"))
                germination.pvals$p.value[i] <- NA
                seedsize.pvals$p.value[i] <- NA
    })
  }
  
  # generate corrected p-values according to the false discovery rate (FDR) method
  # see help(p.adjust) for available methods
  seedsize.pvals$corrected.p.value <- p.adjust(seedsize.pvals$p.value, method="fdr")
  
  names(seedsize.pvals) <- c('Group 1', 'Group 2', 'Raw p value', 'FDR-corrected p value')
  fwrite(seedsize.pvals, file=paste0(rundir, "/seedsize.t-tests.tsv"), sep='\t')
} else {
  cat("Only one group in output, skipping t-tests.\n")
}

if (length(unique(data.long$Group)) > 1) {
  dir.create(paste0(rundir, "/Kaplan-Meier Plots"))
  # generate table for kaplan-meier plots
  # we only compare two groups for each analysis -- 
  #   please make your own custom analysis if this is inappropriate for your data!
  pvals$km.logrank <- NA
  
  # include only seeds that were processed normally or status is unknown (i.e., log file missing)
  data.surv <- data.peruid[data.peruid$Note=='Seed processed normally.' | is.na(data.peruid$Note),]

  data.surv$event <- 1
  data.surv$event[which(is.na(data.surv$`Germination Time (h)`))] <- 0
  # ungerminated seeds have a 0 for event and a time equal to end of assay
  data.surv$`Germination Time (h)`[which(is.na(data.surv$`Germination Time (h)`))] <- max(data$ElapsedHours)

  # make a kaplan-meier plot for all groups
  survobj <- Surv(time=data.surv$`Germination Time (h)`, event=data.surv$event)
  sfit <- survfit(survobj~Group, data=data.surv)
  p <- ggsurvplot(sfit, pval=F, fun=function(y) { return(1-y) }) +
    labs(x='Elapsed Time (h)', y='Fraction of germinated seeds')
  suppressWarnings(ggsave(p$plot + geom_hline(yintercept=1), 
                          filename=paste0(rundir, "/Kaplan-Meier Plots/KaplanMeier-allgroups.pdf"), 
                          width=25, height=15, units="cm"))

  num_cores <- max(1, detectCores() - 1)
  cl <- makeCluster(num_cores)
  registerDoParallel(cl)
  
  pvals$km.logrank <- foreach(i=seq(1, nrow(pvals)), .combine=c, .packages=c('survival', 'survminer', 'data.table')) %dopar% {
    kmplot(pvals$Group.1[i], pvals$Group.2[i], data.surv, paste0(rundir, "/Kaplan-Meier Plots/KaplanMeier-", pvals$Group.1[i], "_vs_", pvals$Group.2[i]))
  }
  
  stopCluster(cl)

  names(pvals) <- c('Group 1', 'Group 2', 'Log-Rank p-value')
  fwrite(pvals, file=paste0(rundir, "/germination.kaplan-meier_test.tsv"), sep='\t')
}

cat(paste0("Statistics have been written to the directory ", rundir, "\n"))
if (grepl('Root Growth$', outdir)) {
  prominent_message("Now run consolidate_rootgrowth_data.R.")
}
```
