## Supplementary File 5 for "SPIRO – the automated Petri plate imaging platform designed by biologists, for biologists": process_rootgrowth_data.R

```
# process_rootgrowth_data.R - 
#
#   takes data generated by consolidate_rootgrowth_data.R and estimates per-group root growth rates.
# 

# clean slate
rm(list=ls())
source('R scripts/common.R')
p_load(glmmTMB, parallel, data.table, ggplot2, dplyr, emmeans, multcomp, multcompView, effects, ggeffects, MASS, tibble, tidyr)

dir <- choose_dir()

resultsdir <- paste0(dir, '/Results')
outdir <- paste0(resultsdir, '/Root Growth')
rundir <- create_rundir(outdir)

if (file.exists(paste0(outdir, "/rootgrowth.postQC.tsv"))) {
  data <- fread(paste0(outdir, '/rootgrowth.postQC.tsv'))
} else {
  stop("rootgrowth.postQC.tsv not found in specified directory. Did you run consolidate_rootgrowth_data.R?\n")
}

# if Group is set to NA, do not include in analysis
data <- data[which(!is.na(data$Group)),]

# copy postQC input file to rundir for reference
dir.create(paste0(rundir, '/QC Results'))
file.copy(paste0(outdir, "/rootgrowth.postQC.tsv"), paste0(rundir, '/QC Results'))
file.copy(paste0(outdir, "/germination-perseed.tsv"), paste0(rundir, '/QC Results'))

# loop to ask the user to choose the control group
groups <- as.character(unique(data$Group))
ngroups <- length(groups)

# fit the mixed model
cat("Fitting the model, hold tight...\n")
num_cores <- max(1, detectCores() - 1)
if (ngroups > 1) {
  fit.glmmTMB <- glmmTMB(data=data,
          formula=PrimaryRootLength ~ 1 + poly(RelativeElapsedHours, 2, raw=TRUE) * Group + (1+poly(RelativeElapsedHours, 2, raw=TRUE) | UID),
          REML=T, control=glmmTMBControl(optimizer=optim, optArgs=list(method="BFGS"), parallel=num_cores))
  fit.glmmTMB %>% emtrends(~Group, var="RelativeElapsedHours") -> fit.trend
  fit.cld <- cld(fit.trend)
  fit.cld %>% as_tibble %>%
    dplyr::select(c(Group, .group)) %>%
    rename(Tukey.CLD = .group) -> fit.cld
  fit.pwpp <- pwpp(fit.trend, values=F) + theme_bw()
  
  fwrite(fit.cld, file=paste0(rundir, '/Tukey CLD.tsv'), sep='\t')
  ggsave(fit.pwpp, filename = paste0(rundir, "/Pairwise p-value plot.pdf"), width=25, height=15, units="cm")
} else {
  fit.glmmTMB <- glmmTMB(data=data,
          formula=PrimaryRootLength ~ 1 + poly(RelativeElapsedHours, 2, raw=TRUE) + (1+poly(RelativeElapsedHours, 2, RAW=TRUE) | UID),
          REML = T)
}

# make trend graph
if (ngroups > 1) {
  fit.glmmTMB %>% ggeffect(terms = c("RelativeElapsedHours [0,24,48,72,96,120,144]","Group")) -> glmmTMB_model_ggeffects
  data %>% rename(group=Group, x=RelativeElapsedHours, predicted=PrimaryRootLength) %>%
    dplyr::select(c(UID, x, predicted, group)) -> plotting
  glmmTMB_model_ggeffects %>% as_tibble %>%
    ggplot(aes(x=x, y=predicted, ymin=conf.low, ymax=conf.high, fill=group, group=group)) +
    geom_line(linetype="dashed") + 
    geom_ribbon(alpha=.5) +
    geom_line(data=plotting, aes(group=UID, ymin=NULL, ymax=NULL), alpha=.3) +
    facet_wrap(~group) +
    labs(fill='Group', y="Predicted primary root length (cm)", x="Time since root emergence (h)") +
    theme_bw() +
    theme(legend.position="none") -> p
  ggsave(p, filename=paste0(rundir, '/Model prediction overview.pdf'), width=25, height=25, units="cm")
} else {
  fit.glmmTMB %>% ggeffect(terms = c("RelativeElapsedHours [0,24,48,72,96,120,144]")) -> glmmTMB_model_ggeffects
  data %>% rename(group=Group, x=RelativeElapsedHours, predicted=PrimaryRootLength) %>%
    dplyr::select(c(UID, x, predicted, group)) -> plotting
  glmmTMB_model_ggeffects %>% as_tibble %>%
    ggplot(aes(x=x, y=predicted, ymin=conf.low, ymax=conf.high)) +
    geom_line(linetype="dashed") + 
    geom_ribbon(alpha=.5) +
    geom_line(data=plotting, aes(group=UID, ymin=NULL, ymax=NULL), alpha=.3) +
    labs(y="Predicted primary root length (cm)", x="Time since root emergence (h)") +
    theme_bw() +
    theme(legend.position="none") -> p
  ggsave(p, filename=paste0(rundir, '/Model prediction overview.pdf'), width=25, height=25, units="cm")
}

glmmTMB_model_ggeffects %>% as_tibble %>% 
  arrange(group, x) %>%
  rename(RelativeHours = x, PredictedRootLength = predicted, Group = group) %>%
  dplyr::select(c(Group, RelativeHours, PredictedRootLength, std.error, conf.low, conf.high)) %>%
  mutate(PredictedRootLength=round(PredictedRootLength, 3), std.error=round(std.error, 3), conf.low=round(conf.low, 3),
         conf.high=round(conf.high, 3)) %>%
  filter(RelativeHours > 0) -> root.table

fwrite(root.table, file=paste0(rundir, '/Predicted Root Lengths.tsv'), sep='\t')

p <- ggplot(root.table, aes(x=Group, y=PredictedRootLength, ymin=conf.low, ymax=conf.high, fill=as.ordered(RelativeHours))) + 
  geom_col(position="dodge", color="black") +
  geom_errorbar(position="dodge") +
  scale_fill_brewer(palette='Greens') +
  theme_bw() +
  labs(fill='Hours', y='Predicted root length (cm)') +
  theme(axis.text.x = element_text(angle = 90, vjust = 0.5))
ggsave(p, filename=paste0(rundir, '/Root length barchart.pdf'), width=25, height=15, units="cm")

# make growth rate estimates by approximating derivative of fitted model
rates <- NULL
for (time in c(24,48,72,96,120,144)) {
  Xp <- stats::model.matrix(object = formula(fit.glmmTMB) %>% lme4::nobars(), 
                            data=tibble(PrimaryRootLength=1, 
                                        RelativeElapsedHours=time,
                                        Group=(fit.glmmTMB$frame$Group) %>% unique))
  Xp2 <- stats::model.matrix(object = formula(fit.glmmTMB) %>% lme4::nobars(), 
                             data=tibble(PrimaryRootLength=1, 
                                         RelativeElapsedHours=time+0.00001,
                                         Group=(fit.glmmTMB$frame$Group) %>% unique))
  
  Vc <- vcov(fit.glmmTMB)$cond
  model_coefs <- fixef(fit.glmmTMB)$cond
  
  Bu <- MASS::mvrnorm(n = 10000, 
                      mu = model_coefs, 
                      Sigma = Vc)
  
  t(Xp%*%t(Bu)) -> sim1
  t(Xp2%*%t(Bu)) -> sim2
  
  sim <- (sim2-sim1)/0.00001
  
  # summarize and convert to um/h
  apply(sim, MARGIN = 2, 
        function(x){data.frame(mean=mean(x),
                               conf.low = quantile(x, type=8, prob=0.025),
                               conf.high = quantile(x, type=8, prob=0.975))}) %>%
    enframe() %>%
    unnest(value) %>%
    mutate(RelativeElapsedHours=time,
           Group=fit.glmmTMB$frame$Group %>% unique,
           mean=mean*10000,
           conf.low=conf.low*10000,
           conf.high=conf.high*10000) %>%
    dplyr::select(Group, RelativeElapsedHours, mean, conf.low, conf.high) -> rate

  rates <- rbind(rates, rate)
}

p <- ggplot(rates, aes(x=Group, y=mean, ymin=conf.low, ymax=conf.high, fill=as.ordered(RelativeElapsedHours))) + 
  geom_col(position="dodge", color="black") +
  geom_errorbar(position="dodge") +
  scale_fill_brewer(palette='Greens') +
  theme_bw() +
  labs(fill='Hours', y=expression('Predicted root growth rate ('*mu*'m/h)')) +
  theme(axis.text.x = element_text(angle = 90, vjust = 0.5))
ggsave(p, filename=paste0(rundir, '/Root growth rate barchart.pdf'), width=25, height=15, units="cm")

names(rates)[3] <- 'GrowthRate_um_h'
fwrite(rates, file=paste0(rundir, '/Predicted Root Growth Rates.tsv'), sep='\t')

cat(paste0('Wrote statistics and graphs to ', rundir, '.\n'))
```
