## Supplementary File 5 for "SPIRO – the automated Petri plate imaging platform designed by biologists, for biologists": README.md

<p align="center">
  <img src="https://github.com/AlyonaMinina/Files_for_SPIRO_reps/blob/master/SPIRO.Assays%20files/SPIRO%20text%20logo.png?raw=true" height="80" title="SPIRO Assays"><br>
  <b>S</b>mart <b>P</b>late <b>I</b>maging <b>Ro</b>bot semi-automated assays
</p>

This repository contains ImageJ macros, R scripts and a detailed [manual](https://github.com/jiaxuanleong/SPIRO.Assays/raw/master/SPIRO%20Assays%20Manual.pdf) for semi-automated high-throughput image analyses for seed germination and root growth assays. Both assays were optimized for the plant model organism *Arabidopsis thaliana* and data acquired using [SPIRO](https://github.com/jonasoh/spiro). 

Each assay comprises three major steps:
* Raw data preprocessing (ImageJ macros)
* Image analysis (ImageJ macros)
* Quality control and statistical analysis (R scripts)

<p align="center">
  <img src="https://github.com/AlyonaMinina/Files_for_SPIRO_reps/blob/master/SPIRO.Assays%20files/spiro-assays-v1-resized-v2.gif" height="400" title="SPIRO assays overview">
</p>

**Figure1.** SPIRO semi-automated assays require preprocessing of the raw data to create 8-bit time-lapse stack files. The preprocessed data can then be used for either seed germination or root growth assays.

# Image preprocessing
<img align="right" width="600" src="https://github.com/AlyonaMinina/Files_for_SPIRO_reps/blob/master/SPIRO.Assays%20files/preprocessing-v2%20flat.png" alt="Preprocessing  overview">

The first step in either of the SPIRO assays is preprocessing of the raw data. **It is used for**:
* Creating a time-lapse file for each analysed Petri plate
* Setting scale in cm
* Reducing file size by cropping off unnecessary background in images
* Converting RGB images into greyscale 8-bit to facilitate further segmentation
* If needed, correcting for accidental drift of object during imaging

# SPIRO Seed Germination Assay

<img align="right" width="600" src="https://github.com/AlyonaMinina/Files_for_SPIRO_reps/blob/master/SPIRO.Assays%20files/germination%20v2%20flat.png" alt="SPIRO Germination assay overview">

This assay was designed for *Arabidopsis thaliana* seed germination analysis by detecting the time point where the seed perimeter starts showing a stable increase.

**At a glance, the assay is performed like this:**
* Preprocessed data is used to select groups of seeds and define the desired time range for analysis
* Perimeters and areas are recorded for each seed and time point
* The data is data is subjected to quality control
* Optionally, grouping can be adjusted using the SPIRO assay customizer
* Data is processed further to determine germination time point for each seed based on dynamics of its perimeter changes

**After processing, results include:**
* Germination time point for each seed
* Size of each seed
* Germination statistics for each user-selected group of seeds (based on the [germinationmetrics](https://aravind-j.github.io/germinationmetrics/index.html) package)
* Kaplan-Meier plots for groupwise comparisons
* T-test results for seed sizes

# SPIRO Root Growth Assay

<img align="right" width="600" src="https://github.com/AlyonaMinina/Files_for_SPIRO_reps/blob/master/SPIRO.Assays%20files/root-grwoth-v2%20flat.png" alt="SPIRO Root Growth assay overview">

This assay allows tracking primary root lengths and growth rates for indivual seedlings and groups of seedlings. Primary root growth is tracked starting from the germination time point, and statistical comparison of root growth between groups is performed using a mixed model. 

**The assay comprises several steps:**
* Preprocessed data is used to select groups of seeds and define the desired time range for analysis
* Seed perimeters and areas are recorded for each time point
* The primary root length is recorded for each seedling at each time point
* Germination time point is detected for each seed
* The primary root length data is normalized to the germination time point and subjected to quality control
* The data can be optionally relabelled and regrouped using the SPIRO assay customizer

**The results of the assay include:**
* Root length for each seedling at each time point
* Bar charts representing per-group root length and growth rates at 24-h intervals
* Plots visualizing the model prediction of root growth for each group
* Plots with raw and normalized (to germination time) root length vs time for each seedling of a group 
* Results of statistical analysis comparing root growth between groups 

# SPIRO Assay Customizer
<img align="right" width="400" src="https://user-images.githubusercontent.com/6480370/86357062-ce289280-bc5c-11ea-816f-a656977b224c.png" alt="SPIRO Assay Customizer">

The [SPIRO Assay Customizer](https://github.com/jonasoh/spiro-assay-customizer) is a companion tool for SPIRO Assays, which enables more user friendly handling of data rearrangment for SPIRO assays. Its use is optional but recommended. 

**The Customizer provides an intuitive user interface for**:
* Merging data from several experiments
* Relabelling samples or groups of samples
* Removing samples or groups of samples from analysis
* Reshuffling samples between groups

# SPIRO Assay DEBUG mode

DEBUG mode can be enabled at the start of each assay by holding down the Ctrl (Control) key.

## Preprocessing

* Batch size during drift correction may be lowered in order to reduce RAM requirements
> Drift correction for large datasets (e.g. more than 200 time points) can take up a lot of RAM. This may cause ImageJ to freeze up when its allocated RAM is too low, thus images are drift-corrected in batches. If the default batch size of 350 is causing ImageJ to freeze up on your computer, you may reduce this number e.g. to 100, when the macro prompts you to do so.

## Seed Germination Assay

* Seed detection parameters, i.e. area and circularity, may be modified
> The default seed detection parameters were optimized using typical *Arabidopsis thaliana* seeds. If your seeds do not fit the default parameters, the expected area and circularity may be modified when the appropriate dialog box appears during macro run in the debug mode. The area and circularity of your seeds can be estimated manually in ImageJ using the same thresholding as in the macro.

> **! Changes in the allowed area size must be also introduced into downstream R germination QC R script to allow for correct data filtering. For this, please change the variables `upper_area_threshold` and `lower_area_threshold` in `cleanup_germination_data.R`.**

## Root Growth Assay

* Seed detection parameters, i.e. area and circularity, can be modified
> See above under “Seed Germination Assay”.

* Overlay skeletons can be enabled
> Lighting conditions may affect image capture and cause some roots to look translucent, making it difficult for the thresholding methods to distinguish roots from growth media. As a result, on some time frames detected roots might have gaps in their outlines. To overcome this, we introduced the “Overlay Skeletons” function which superimposes roots from one time point to the next, thus filling in potential gaps. However, this function might increase background noise.

* Non-essential intermediate output files will not be deleted at the end of the run
> If the results of the root growth macro do not make sense, the intermediate files are useful for troubleshooting root masking and root start coordinates.

# Troubleshooting

Some common errors and workarounds have been summarized in these tables:
* [Preprocessing troubleshooting](https://github.com/AlyonaMinina/Files_for_SPIRO_reps/blob/master/SPIRO.Assays%20files/Preprocessing%20troubleshooting.md)
* [Germination assay troubleshooting](https://github.com/AlyonaMinina/Files_for_SPIRO_reps/blob/master/SPIRO.Assays%20files/Germination%20assay%20troubleshooting.md)
* [Root growth assay troubleshooting](https://github.com/AlyonaMinina/Files_for_SPIRO_reps/blob/master/SPIRO.Assays%20files/Root%20growth%20assay%20troubleshooting.md)
