## Supplementary File 5 for "SPIRO – the automated Petri plate imaging platform designed by biologists, for biologists": SPIRO Assays Manual.pdf

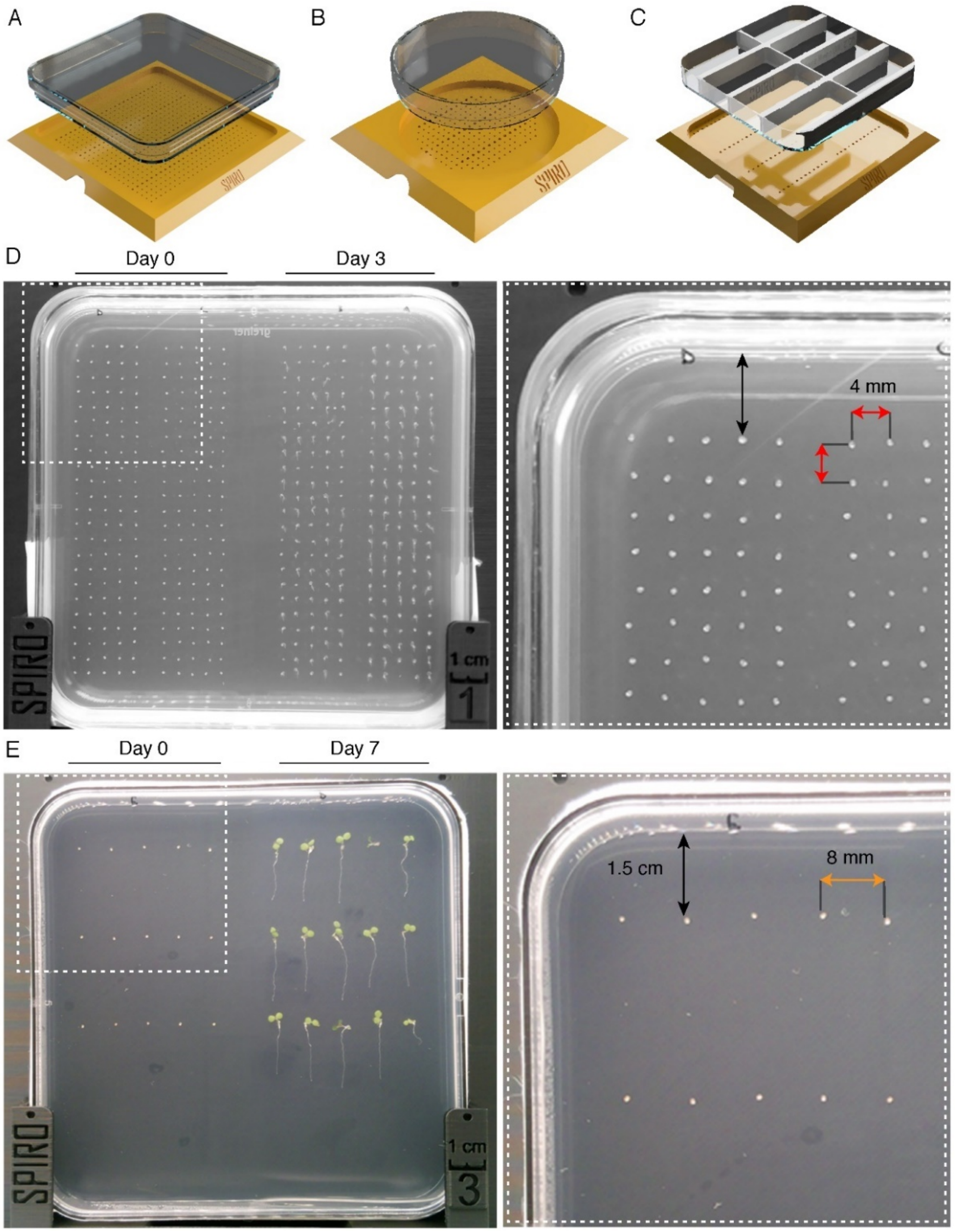


**Figure 1. Examples of plating seeds for germination and root growth assays**.
(**A**) 3D printed seed plating guide for 12 cm square plates.(**B**) An example of a plate imaged by SPIRO for germination assay on the first and third days of the experiment. Seeds are placed 1.5 cm away from plate edges and 4 mm away from each other. (**C**) 3D printed seed plating guide for 9 cm round plates. (**D**) An example of a plate imaged by SPIRO for the root growth assay at the first and the seventh day of the experiment. Seeds are placed 1.5 cm away from plate edges and 8 mm away from each other. (**E**) 3D printed insets that can be glued into Petri plates to create multi-chamber compartments applicable for different treatments. The insets can be used with the corresponding seed plating guides.

- Use custom lids printed with black filament to eliminate reflections (**Table S2** of this manuscript and in the SPIRO Hardware Repository[1]).

  Reflections in the plastic of normal lids are detrimental to automated image analysis, especially during root detection (**Fig. 2**). The printed lids should preferably be sterilized using UV light. Sterilization in 70% ethanol is also possible, but must be followed by a prolonged drying of the lids. Ethanol left in pores of the plastic might impact seedling growth.

- For long time-lapse experiments, plates should be properly sealed to prevent evaporation.

  Please note that plate holders are quite tight and will not allow for bulky layers of Parafilm or tape. We recommend using plastic seal (Phytotechlab, Product ID: A003).

- Place the plates into the holders of SPIRO with lids towards the cube.

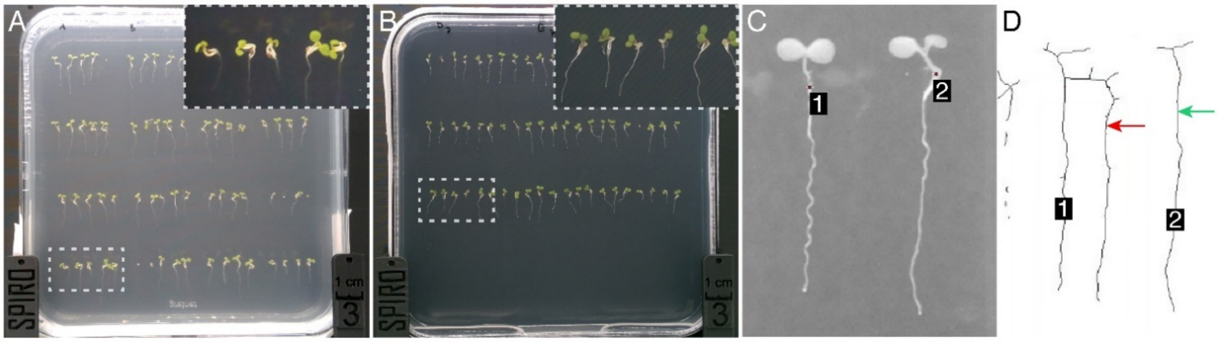


**Figure 2. Reducing reflections in the Petri plate lids is crucial for automated image processing.** Examples of images acquired using normal (**A**) and 3D-printed anti-reflection (**B)** lids. (**C**) A magnified part of the photo shown in (A) processed by the SPIRO root growth assay, seedlings recognized by the software are indicated with numbers. (**D**) The result of image segmentation by the SPIRO root growth assay performed on the data shown in (C). Background is shown in white, while objects recognized as seedlings are shown as black lines with ID numbers. Green arrow, seedling 2; red arrow, seedling 2 reflection being mistakenly annotated as a part of seedling 1.

### 2.2 SPIRO software settings

- SPIRO is controlled by the designated software on the Raspberry Pi computer and operated by the user via a web-based user interface. Thus, to enable access to the SPIRO software during experiment setup, the robot must be connected to the internet either via Wi-Fi or Ethernet cable, or set up to act as a Wi-Fi hotspot.

Since plant growth cabinets are often located where an internet connection is not available, the SPIRO computer can be used as a Wi-Fi hotspot (for instructions see **File S2** of this manuscript and SPIRO Software Repository[2]).

- To access the SPIRO web UI, open a web browser and type in the address http://x.y.z.w:8080/ (where x.y.z.w is IP address of your Raspberry Pi computer). Log in using the password you provided during SPIRO Web UI initial setup (**File S2** of this manuscript and SPIRO Software Repository[2]).

Note that in Wi-Fi hotspot mode, any address will work (although we recommend using the address spiro.local).

- In the **Live view** tab (**Fig. 3**), use the focus slider or type in numbers to adjust the **focus** (for SPIROs with motorized cameras), **Zoom** and **pan** (icon with four arrows) tools might be helpful for verification of the focus. Please note, that the acquired images must contain the complete 1 cm scale bar engraved on SPIRO if they are to be used with the provided automated assays.

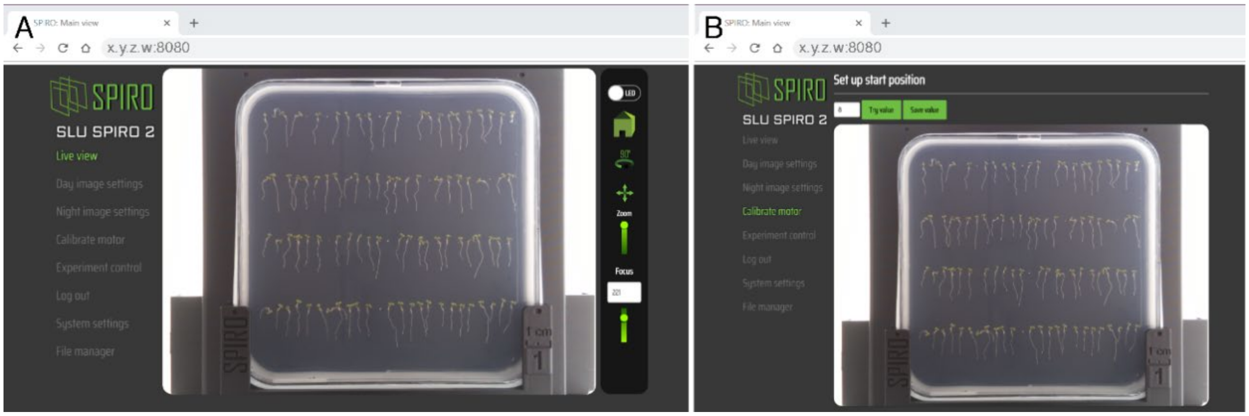


**Figure 3. Using the web interface to set up imaging.**
Live view (**A**) and Calibrate motor (**B**) tabs of the SPIRO web UI should be used to adjust the focus and alignment of the cube position with the camera.

- Press the **Home** button on the right panel of the Live view and check if the first plate is moved to face the camera. If the plate is not facing the camera, you can adjust and verify the optimal value for cube rotation angle in the **Calibrate motor** tab.

- **Day** and **night image settings** tabs allow adjustment of ISO and camera shutter speed to optimize image light intensity under desired imaging conditions.

As the accuracy of image processing for semi-automated assays depends greatly on the image quality, avoid acquiring over- or under-exposed images (**Fig. 4**).

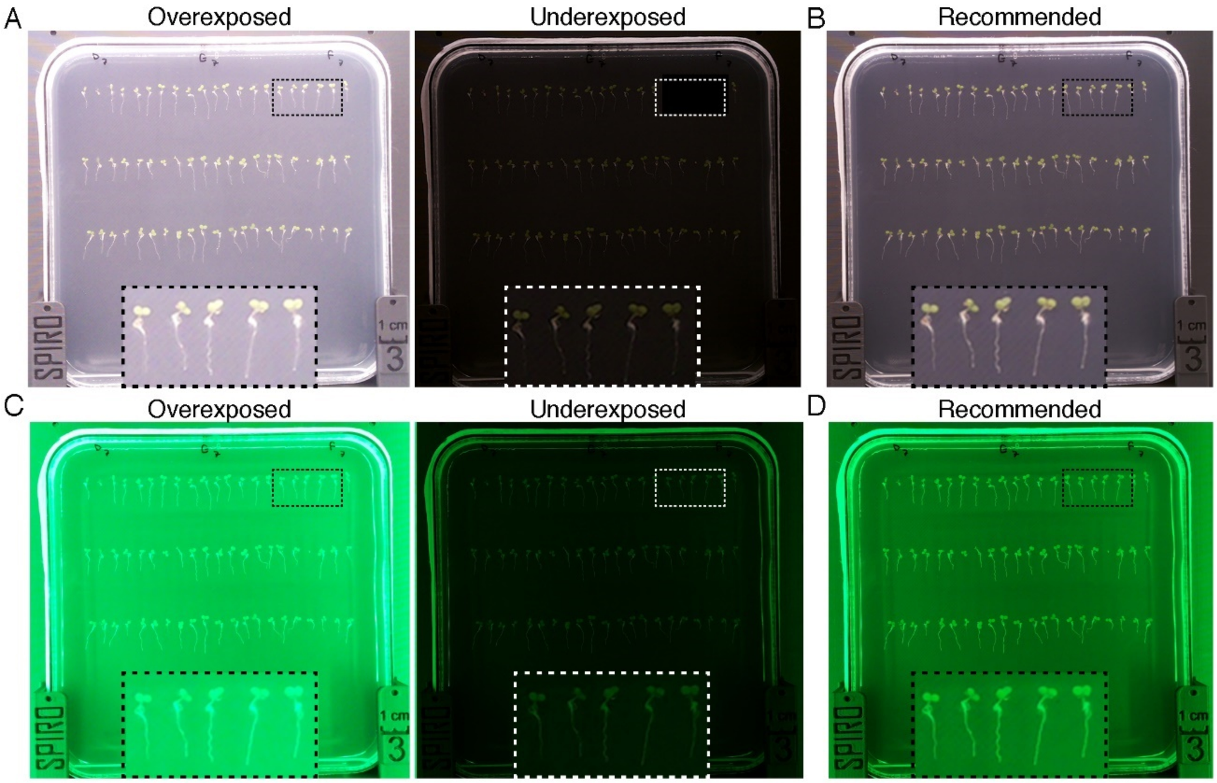


**Figure 4. Impact of the exposure settings on the image quality.** Day and night image settings tabs allow adjustment of ISO and shutter speed for optimal image light intensity under day and night conditions. (**A, C**) Examples of over- and under-exposed day and night images that impacts quality of data. (**B, D**) Examples of images with correct exposure, optimal for the semi-automated assays.

- In the **Experiment** control tab, you will be prompted to enter experiment folder name.

Please note that this name will later be used during data analysis to create a unique ID for each seed/seedling. If you wish to combine data from different experiments for analysis it is important to have unique experiment folder names. You will also be asked to provide imaging frequency and duration of your experiment. SPIRO is capable of imaging at the maximum rate of 4 plates/ca 2 minutes. **SPIRO germination and root growth semi-automated assays were optimized for imaging every 30 and 60 minutes, respectively**.

- The software will estimate required and available storage space. If amount of available space is less than is required, it will be shown in red.

Please note that the experiment will terminate, when no more storage space will be available. To free up space, you can remove unnecessary files from the Raspberry Pi computer even during a running experiment (see Section 2.3).

### 2.3 Accessing SPIRO Data

- The experiment folder with acquired images will be saved on the microSD card of the SPIRO Raspberry Pi computer.

- To download SPIRO experiment data, click on the menu item *File manager* in the web interface.

  The File manager presents a list of all experiment folders stored on the microSD card. For each item, there are buttons for downloading and deleting data. Downloaded data comes as a ZIP file, which needs to be extracted before using it for the semi-automated SPIRO assays.

- SPIRO captures images with the following directory structure and file naming scheme (**Fig. 5**):

*<experiment name>/<plateN>/<date>-<time>-[day/night].png*

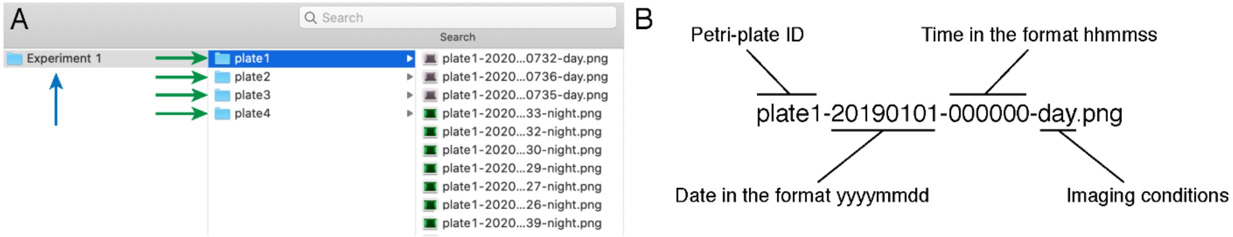


**Figure 5. SPIRO image data file structure.**
For each experiment all images of an individual plate are grouped into a subfolder (green arrow). All subfolders are contained within a single main folder (blue arrow). (**B**) Naming images for the analysis is automatically done by SPIRO.

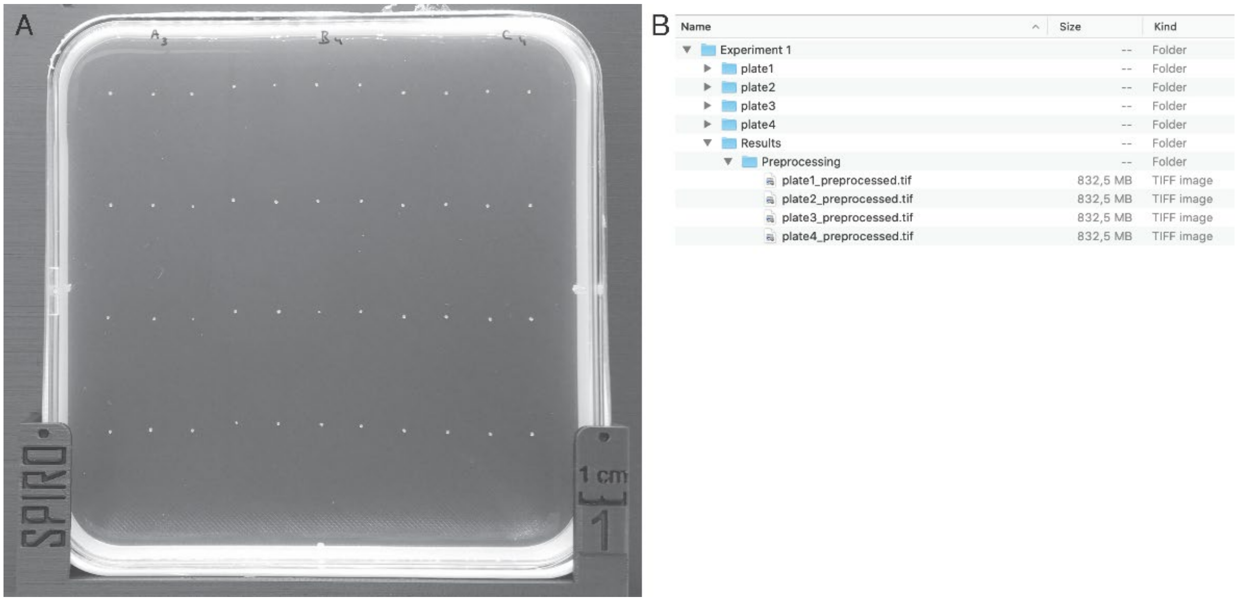


**Figure 6**. **Preprocessing macro output.**
**(A)** A correctly cropped image, including the engraved SPIRO marking. **(B)** The results of preprocessing will be saved as time-lapse TIFF stack files with names corresponding to images of plates being analyzed. The original imaging data will be kept unmodified. Preprocessed data can be further used for detection of germination and for tracking root length. The macros for these tasks will automatically analyze data for all preprocessed plates in the experiment folder.

- *group name* germination analysis.tsv is the quantitative output of the macro, it contains perimeter and area data for each seed at each time point.

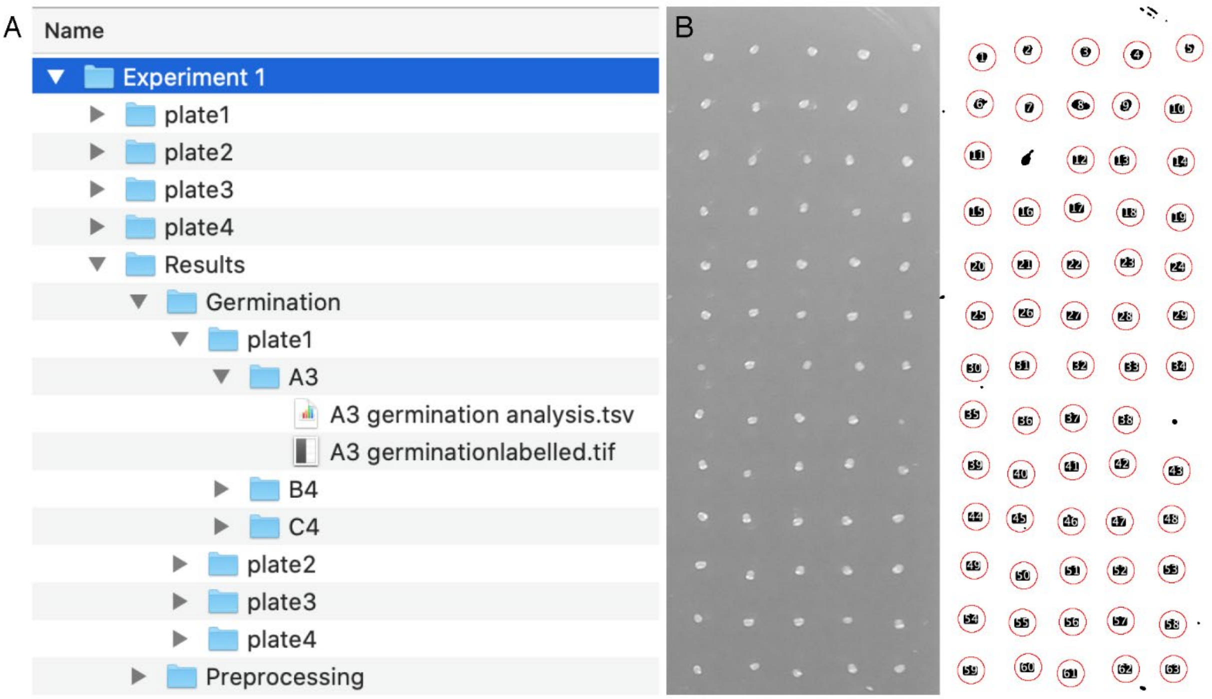


**Figure 7. Output of the Germination Macro.**
(**A**) The results of the germination macro will be saved in the Results/Germination/ Plate*n*/Group_name subfolder. The preprocessed file cropped according to the group selection and user-selected time truncation together with the results of image segmentation will be provided in the group_name germinationlabelled.tif. This file can be used to assess the quality of seed recognition. The file group_name germination analysis.tsv will contain quantitative data about size and perimeter changes for each seed over time. (**B**) An example of a single frame from a group_name germinationlabelled.tif stack file. The left part is the photo acquired by SPIRO and cropped by the user, and the right part is the result of this image segmentation by the Germination macro. Each red circle is centered around an object recognized as a seed, and the numbers indicate the number of a seed that can be then tracked in the TSV file.

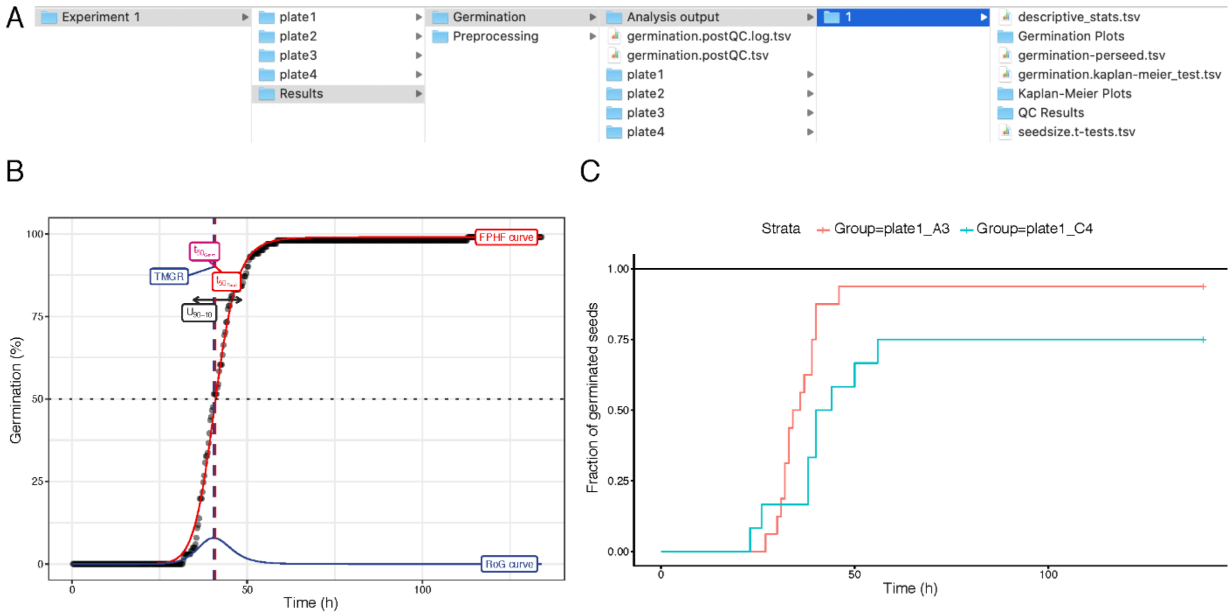


**Figure 8. Results of analysis output from the germination R scripts.**

(**A**) The results of the cleanup_germination_data.R script will be saved in two tsv files: germination.postQC.log.tsv and germination.postQC.tsv. The latter one can be used to manually modify group names while changing comparisons for the statistical analysis by process_germination_data.R. Results of each round of statistical analysis will be saved in a numbered subfolder within *Analysis* output folder. (**B**) Germination plot for individual group. FPHF curve, four-parameter Hill function curve; RoG curve, rate-of-germination curve; TMGR, time at maximum germination rate; T50 Germ, time required for 50% of germinated seeds to germinate; T50 total, time required for 50% of total seeds to germinate. Importantly, for this chart it is calculated accounting only viable seeds for the group, while the mean germination time provided in the statistical analysis in the .tsv files is calculated for all seeds detected for the group; $U_{90-10}$, uniformity defined as the time between the timepoints where 10% and 90% of seeds are germinated. Detailed description of the parameters is available in the germinationmetrics documentation (**C**) Kaplan–Meier “survival curve” plot shows proportion of germinated seeds for each group plotted against time. The test compares mean germination time calculated for all seeds present in the group. Results of the test are summarized in the file germination.kaplan-meier_test.tsv.

j. For troubleshooting, please refer to this table.

k. Please submit reports about errors and suggestions for optimizations as a GitHub issue.

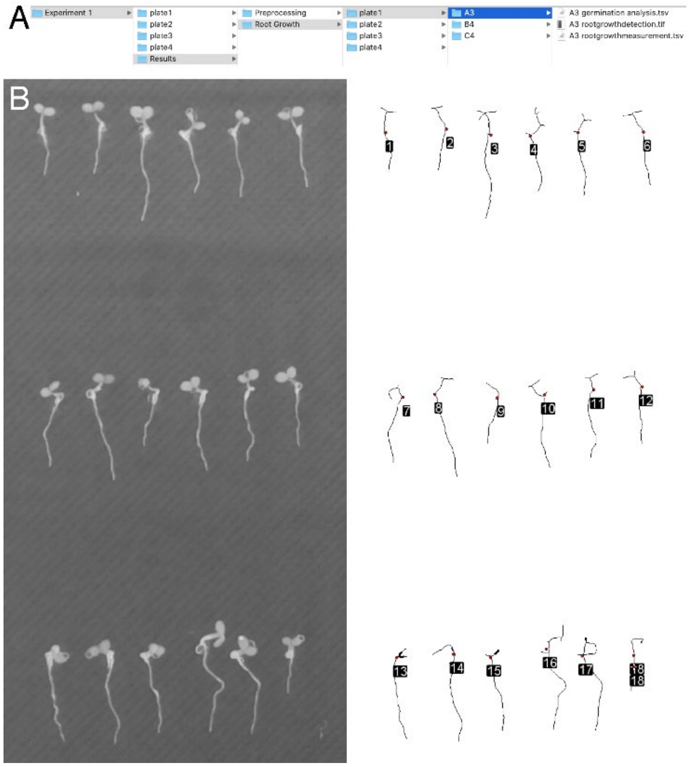


**Figure 9. Output of the root growth macro.**
(**A**) The results of the root growth macro will be saved in the subfolder Results/Root Growth/platen/group_name. The file group_name rootgrowthdetection.tif containing graphical output of root growth estimation, group_name rootgrowthmeasurement.tsv, contains quantitative results of root growth estimation, and group_name germination analysis.tsv, contains quantitative data about size and perimeter changes for each seed over time. (**B**) An example of a single frame from a group_name rootgrowthdetection.tif stack file, which show side by side raw image used for the analysis (left side) and results of this image segmentation (right side). Detected seedlings are presented as skeletonized lines, red dots denote position of the detected root start, the numbers indicate the number of a seedling that can be tracked to the the data in the .tsv file.

q. For troubleshooting, please refer to this table.

r. Please submit reports about errors and suggestions for optimizations as a GitHub issue.

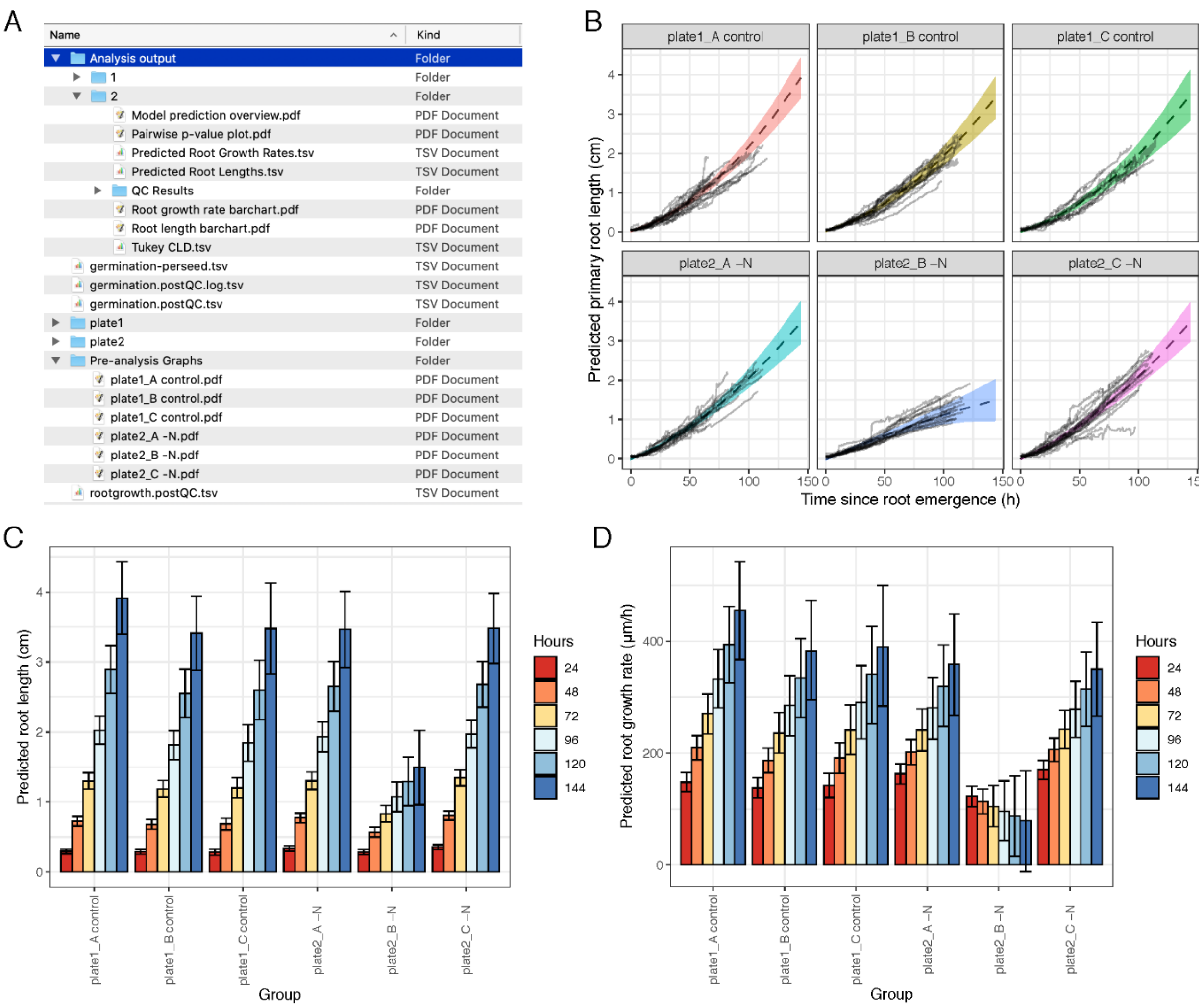


**Figure 10. Output from the root growth R scripts.**
**(A)** Folder structure after running all R scripts, showing locations of the pre-analysis graphs and the analysis output folders. **(B)** Model prediction overview indicating the model fit with standard error for each group. **(C)** Predicted root lengths at 24 h intervals for each group. **(D)** Predicted root growth rates at 24 h intervals for each group.

### 4.7 SPIRO Assay Customizer

SPIRO Assay Customizer is a companion utility for merging and editing experimental data after quality control but prior to statistical analysis. The tool can:

- merge data from several experiments
- reassign seedlings to existing or new groups
- exclude seedlings from analysis

The two main modes, *Merge Assays* and *Customize Assay*, are described below.

#### *4.7.1 Merge assays*

In the Merge Assays mode, you can combine data from several independently analyzed experiments. Experiments are joined after performing data quality control and normalization, i.e. either cleanup_germination_data.R (for germination assay), or after running consolidate_rootgrowth_data.R (root growth assay). After choosing the Merge Assays mode, you are presented with a simple interface (**Fig. 11**).

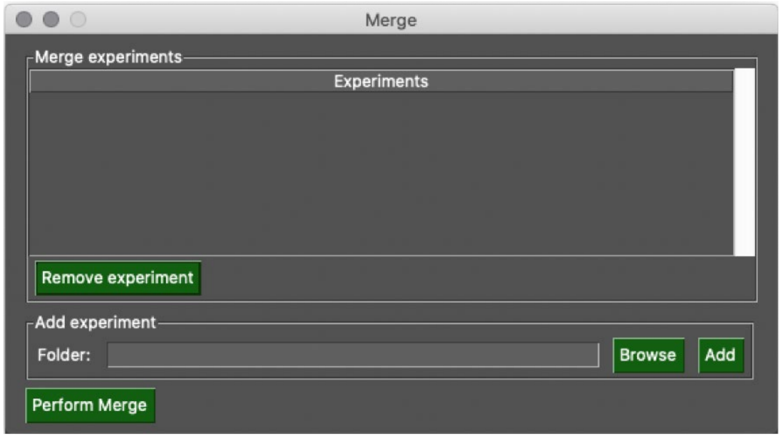


**Figure 11. SPIRO Assay Customizer *Merge Assays* mode.**
For each experiment you want to include in the merged output, click on *Browse*, and locate **either the *Root Growth* or *Germination* subfolder** of the *Results* subfolder of the main Experiment folder. After locating the folder, click *Add*. When all experiments are present in the *Merge Experiments* pane, click *Perform Merge*. The program will ask you to choose a folder. Create a new empty folder and select it. The merged data will be saved to this folder, and you may now run the appropriate processing script on the merged folder (or further customize the assay, see below).

#### 4.7.2 Customize assay

In *Customize Assay* mode, you will be presented with the following window after opening a rootgrowth.postQC.tsv or germination.postQC.tsv file (**Fig. 12**). In the pane to the left, all included seedlings are displayed together with their current group assignment. In the right pane, the groups available for assignment are listed.

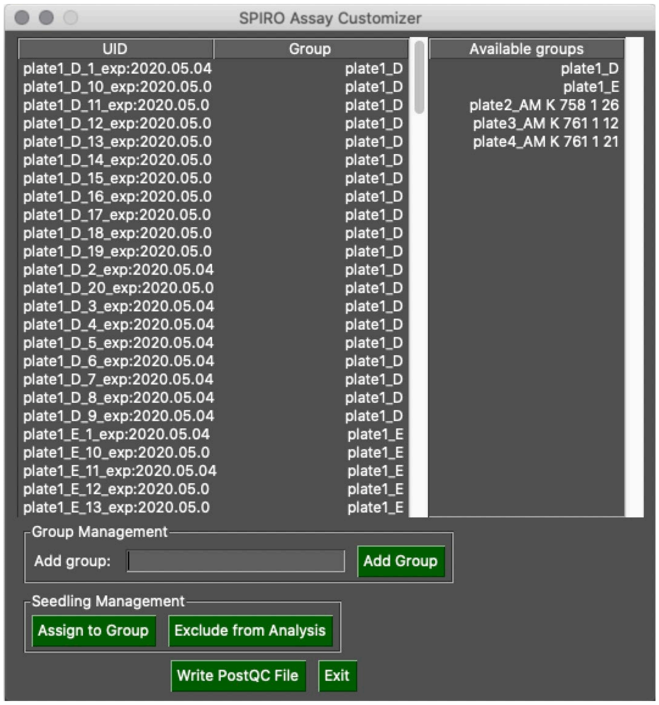


**Figure 12. SPIRO Assay Customizer *Customize Assay* mode.**

- **Adding a new group:** To make a new group available for assignment, type the name of the new group into the *Add group* textbox, then click the *Add Group* button. This makes the new group appear in the right pane.
- **Excluding a seedling from analysis:** To exclude a seedling, select it in the left pane and click *Exclude from Analysis*. Its group assignment will change to *NA*. Please note that several seedlings can be selected if either Shift or Control are held down during selection.
- **Reassigning seedlings to new groups:** With one or more seedlings selected in the left pane, and the desired group selected in the right pane, click *Assign to Group*. The assigned group will change.
- **Saving your changes:** Click on *Write PostQC File* to save your changes.
