## Supplementary File 5 for "SPIRO – the automated Petri plate imaging platform designed by biologists, for biologists": SPIRO_Macros.ijm

```
/*
 * GLOBAL VARIABLES
 * ================
 */

// GENERAL VARIABLES USED BY THE 3 MACROS
var maindir;	// main directory: user-selected directory containing SPIRO-acquired images
var listInmaindir; 	// list of files or directories in the main directory
var resultsdir;		// results directory: all macro-generated results will be saved here
var platedir; 		// plate directories in main directory
var listInplatedir; // number of time points 
var resultsdir;		// results subdir of main directory
var ppdir;			// preprocessing subdir

// alternate types of macro run
var DEBUG = false; // hold down ctrl during macro start to keep non-essential intermediate output files
var freshstart = false; // hold down shift key during macro start to delete all previous data
var selfaware = false; // rootgrowth: alt key during macro start
var overlay = false; // rootgrowth: dependent on DEBUG: prompts user on choice whether to overlay skeletons

// PREPROCESSING-SPECIFIC VARIABLES
var runRegistration // user choice on whether to run registration
var batchsize = 350;
var batched; // if number of time points exceeds batchsize, preprocessing will be carried out in batches
var rbatched = 1;
var rNonbatched = 2;

// GERMINATION-SPECIFIC VARIABLES
var germdir; // directory under resultsdir where germination macro output is contained

// ROOT GROWTH-SPECIFIC VARIABLES
var rootgrowthdir; // directory under resultsdir where germination macro output is contained
var listInrootgrowthdir; // list of directories and files under rootgrowthdir
var step; // current step
var fullplatearray; // array of plates to be processed
var fullgrouparray; // array of groups to be processed
var totalnoOfgroups; //total number of groups to process
var platesToprocess; // if macro resumes from previous run, array of plates to be continued with
var groupsToprocess; // if macro resumes from previous run, array of groups to be continued with
var lengthOfgroupsToprocess; // if macro resumes from previous run, size of array of groups to process
var resumestep; // step to resume from

/*
 * ---------------------
 */

macro "SPIRO_Preprocessing" {
	/*** The preprocessing macro sets scale and processes images to optimize downstream analysis (germination or root growth) *** 
	 *
	 * The workflow is as follows:
	 * 	first detect if batch needed, then get user choice
	 * 	if batch not needed, crop, greench, and register in one go (no intermediate files)
	 * 	if batch needed, separate into segments during run image sequence, save directly in preprocessing (no new folder)
	 * 	use _batchN to denote batch, and plateN can be used to identify it
	 * 	Crop and GreenCh: run as usual on the list of files
	 * 		crop then immediately greenCh to make plateN_batchN_greenCh
	 * 	Register: Open greenCh and add first image to each stack (to prevent jumps between batches), save each batch
	 * 	At the end, merge all plateN_batchN preprocessed to make plateN_preprocessed
	 */

 
	/*  
	 * Check whether the required plugins TurboReg and MultiStackReg are installed
	 * ---------------------------------------------------------------------------
	 */
	List.setCommands;
	if(List.get("TurboReg ")!="") {
		turboreginstalled = true;
	} else {
		turboreginstalled = false;
	}
	if(List.get("MultiStackReg")!="") {
		multistackreginstalled = true;
	} else {
		multistackreginstalled = false;
	} 
	if (!turboreginstalled || !multistackreginstalled) { 
		Dialog.create("Plugin not found"); 
		Dialog.addMessage("Plugins TurboReg and/or MultiStackReg not found. Please refer to the SPIRO manual for installation instructions.");
		Dialog.addCheckbox("I understand.", false);
		Dialog.show();
		usercheck = Dialog.getCheckbox();
		if (usercheck == true)
			exit;
	}

	print("=================================================\n"+ 
		  "Welcome to the companion macro of SPIRO for preprocessing!\n" +
		  "=================================================");
		selectWindow("Log");

	/* Create and/or define working directories
	 * ----------------------------------------
	 */
	showMessage("Welcome to the companion macro of SPIRO for preprocessing!\n" +
		"Please locate and open your experiment folder containing SPIRO-acquired images.\n" +
		"---------\n" +
		"Alternative types of run (hold down then press OK):\n" +
		"SHIFT = Fresh Start mode, all data from any previous run will be deleted\n" +
		"CTRL = Debug mode, batch size for drift correction can be changed.\n");

	wait(1000);

	if (isKeyDown("control"))
	DEBUG = getBoolean("CTRL key pressed. Run macro in debug mode? Batch size for drift correction can be changed.");

	if (isKeyDown("shift"))
		freshstart = getBoolean("SHIFT key pressed. Run macro in Fresh Start mode? This will delete all data from the previous run.");

	maindir = getDirectory("Choose a Directory");
	listInmaindir = getFileList(maindir);
	resultsdir = maindir + "Results" + File.separator; // all output is contained here
	if (!File.isDirectory(resultsdir))
			File.makeDirectory(resultsdir);
	
	ppdir = resultsdir + "Preprocessing" + File.separator; // output from the proprocessing macro is here
	if (!File.isDirectory(ppdir))
		File.makeDirectory(ppdir);
	
	// safeguard against processing non-plate (user-introduced) file
	for (fileindex = 0; fileindex < listInmaindir.length; fileindex++) {
		if (indexOf(listInmaindir[fileindex], "plate") == -1)
			listInmaindir = Array.deleteValue(listInmaindir, listInmaindir[fileindex]); 
	}
	
	/* 
	 * Get user choice on drift correction 
	 * ---------------------------------------
	 */
	Dialog.create("Drift Correction");
	Dialog.addMessage("Would you like to carry out drift correction (registration)?\n" +
	    "Please note that this step may take up a lot of time and RAM for large datasets.\n" +
	    "Batch size may be reduced in DEBUG mode for lower RAM requirement.");
	dialogchoices = newArray("Yes", "No");
	Dialog.addChoice("", dialogchoices);
	Dialog.show;
	regUserInput = Dialog.getChoice();
	
	if (regUserInput == "Yes") {
		runRegistration = true;
	} else {
		runRegistration = false;
	}
	
	if (DEBUG && runRegistration)
		Dialog.create("(DEBUG) Drift Correction Batch Size");
	Dialog.addMessage("Drift correction (registration) may be carried out in smaller batches of the image stack to reduce RAM requirement.\n"
		+ "Please set desired batch size");
	Dialog.addNumber("Batch size", batchsize);

	/* Main chunk of code - all run functions are here
	 * -----------------------------------------------
	 */

	if (freshstart)
		deleteOutputPP();
	scale();
	batch(); 
	cropnGreenCh();
	if (runRegistration) {
		if (batched)
			register(rbatched);
		if (!batched)
			register(rNonbatched);
	} else {
		noReg();
		print("Step 4/4 Drift correction skipped due to user choice");
	}
	deleteOutputPP();
	
	list = getList("window.titles");
	list = Array.deleteValue(list, "Log");
	for (i=0; i<list.length; i++) {
		winame = list[i];
		selectWindow(winame);
		run("Close");
	}
	
	print("\nPreprocessing is complete.");
	selectWindow("Log");

	/* ----------------------------------------------
	 * Lines below this are functions and their descriptions
	 */

	function scale() {
		/* The scale() function first attempts to automatically find out the scale based on the approximate location of the 1cm scale bar printed on SPIRO
		 *  this is done by masking then filtering objects based on their width-to-height (WtH) ratio (the scale bar should be something longer horizontally)
		 *  an additional filter: potential scale bar must be more than 150 pixels long / wide ( to prevent small noise from matching WtH ratio.
		 *  if only a single object matches the potential scale bar, this is highlighted and confirmation from the user is obtained
		 * If multiple or no objects match scale bar WtH ratio, or the automatically detected scale bar is wrong
		 *  the user is prompted to draw a line corresponding to the length of the scale bar
		 */
		 
		if (is("Batch Mode"))
			setBatchMode(false);
		print("Step 1/4 Setting scale...");
		plate1dir = maindir + listInmaindir[0]; // only first image of first plate needed to set scale
		listInplate1dir = getFileList(plate1dir);
		img1 = listInplate1dir[0];
		open(plate1dir + img1);
		run("Set Scale...", "distance=1 known=1 unit=pixel"); //remove scale
	
		run("Set Measurements...", "area bounding display redirect=None decimal=3");

	    // get approximate location of scale bar
		setTool("line");
		xapprox = getWidth()/5*4;
		xmax = getWidth();
		yapprox = getHeight()/5*4;
		ymax = getHeight();
		widthapprox = xmax - xapprox;
		heightapprox = ymax - yapprox;
		/*
	   	makeRectangle(xapprox, yapprox, widthapprox, heightapprox);
	    run("Duplicate...", "use");
	   	rename("tempscalebar");
	    slicelabel = getInfo("slice.label");

	    // thresholding for scale bar
	    run("8-bit");
	    run("Gaussian Blur...", "sigma=7");
	    
	    if (indexOf(slicelabel, "day") > -1 ) {
			run("Subtract Background...", "rolling=90");
		} else {
			run("Subtract Background...", "rolling=90 light");
		}
		roiManager("reset");
		setAutoThreshold("Default");
		setOption("BlackBackground", false);
		run("Convert to Mask");

		// filter potential objects by width to height ratio
		run("Create Selection");
		roiManager("Split");
		roicount = roiManager("count");
		for (roino = 0; roino < roicount; roino ++) {
			roiManager("select", roino);
			Roi.getBounds(roix, roiy, roiw, roih);
			whratio = roiw/roih; 
			if (whratio < 2.5 || whratio > 3.5) {
				roiManager("select", roino);
				roiManager("delete");
				roino -= 1; 
				roicount -= 1;
			}
		}

		// filter potential objects by unscaled width
		roicount = roiManager("count");
		if (roicount > 1) {
			for (roino = 0; roino < roicount; roino ++) {
				roiManager("select", roino);
				Roi.getBounds(roix, roiy, roiw, roih);
				if (roiw < 150) { // scale bar should be at least 150 pixels long
					roiManager("select", roino);
					roiManager("delete");
					roino -= 1; 
					roicount -= 1;
				}
			}
		}

		// if only one object matches = potential scale bar, check with user
		if (roicount == 1) {
			scalefail = false;
			roiManager("select", 0);
			Roi.getBounds(roix, roiy, roiw, roih);
			makeLine(roix, roiy + roih/2, roix + roiw, roiy + roih/2);
			Dialog.create("User-guided scale setting");
			choicearray = newArray("Yes", "No");
			Dialog.addChoice("Does the line drawn correspond to the scale bar?", choicearray);
			Dialog.show();
			userchoice = Dialog.getChoice();
		} else {
			scalefail = true;
			userchoice = "none";
		}

		// if multiple or no objects match + auto-detected scale bar is wrong, prompt user to draw line corresponding to scale bar
		if (userchoice == "no" || scalefail) {
			close("tempscalebar");
			selectWindow(img1);
			run("Select None");
			run("Set... ", "zoom=50 x=["+xapprox+"] y=["+yapprox+"]"); 
			userconfirm = false;
			while (!userconfirm) { 
				Dialog.createNonBlocking("Automatic scale detection unsucessful");
				Dialog.addMessage("Based on the scale bar, draw a straight line corresponding to 1cm\n." +
					"Holding down SHIFT helps in keeping line horizontal");
				Dialog.addCheckbox("Scale bar corresponding to 1cm has been drawn", false);
				Dialog.show();
				userconfirm = Dialog.getCheckbox();
			}
		} else {
			close("tempscalebar");
			selectWindow(img1);
			makeLine(xapprox + roix, yapprox + roiy, xapprox + roix + roiw, yapprox + roiy);
		}
		*/
		run("Set... ", "zoom=50 x=["+xapprox+"] y=["+yapprox+"]");
		userconfirm = false;
		while (!userconfirm) { 
			Dialog.createNonBlocking("Scale");
			Dialog.addMessage("Based on the scale bar, draw a straight line corresponding to 1 cm\n." +
				"Holding down SHIFT helps in keeping the line horizontal.");
			Dialog.addCheckbox("Scale bar corresponding to 1 cm has been drawn", false);
			Dialog.show();
			userconfirm = Dialog.getCheckbox();
		}
		// measures line length (auto-detected or user-selected) and sets scale
		run("Set Measurements...", "area bounding display redirect=None decimal=3");
		run("Measure");
		Table.rename("Results", "scalebar");
		length = getResult('Length', nResults - 1);
		run("Set Scale...", "distance=" + length + " known=1 unit=cm global");
	
		// based on scale bar, get coordinates to crop so that only SPIRO-relevant part of image is present
		nR = Table.size("scalebar");
		bx = Table.get("BX", nR - 1, "scalebar");
		bx = parseFloat(bx);
		by = Table.get("BY", nR - 1, "scalebar");
		by = parseFloat(by);
		length = Table.get("Length", nR - 1, "scalebar");
		xmid = (bx + length/2);
		xmid = bx + 0.5*length;
		xmid = parseFloat(xmid);
		dx = 13;
		dy = 10.5;
		toUnscaled(dx, dy);
		x1 = xmid - dx;
		y1 = by - dy;
		width = 14;
		height = 12.5;
		toUnscaled(width, height);
		roiManager("reset");
		makeRectangle(x1, y1, width, height);
		roiManager("add");
		run("Original Scale"); // zoom out
		userconfirm = false;
		selectWindow(img1);
		while (!userconfirm) { 
			Dialog.createNonBlocking("Crop selection");
			Dialog.addMessage("If needed, please correct the selected area for cropping by updating ROI in ROI manager, then click OK.");
			Dialog.addCheckbox("ROI is correct", false);
			Dialog.show();
			userconfirm = Dialog.getCheckbox();
		}
		roiManager("select", 0);
		getBoundingRect(xcrop, ycrop, wcrop, hcrop);
		Table.create("cropcoordinates");
		Table.set("xcrop", 0, xcrop, "cropcoordinates");
		Table.set("ycrop", 0, ycrop, "cropcoordinates");
		Table.set("wcrop", 0, wcrop, "cropcoordinates");
		Table.set("hcrop", 0, hcrop, "cropcoordinates");
		close();
		selectWindow("scalebar");
		run("Close");
	}
	
	function batch() {
		/* If the number of time points in the experiment (alternatively, the number of images in the stack) exceeds batchsize (default 350)
		 *  images are separated into batches for processing, as we found that this can help in reducing RAM requirement
		 */
		if (!is("Batch Mode"))
			setBatchMode(true);
		print("\nStep 2/4 Converting images into stacks..." +
			  "\nIt may look like nothing is happening, please be patient.");
		selectWindow("Log");
		for (plateno = 0; plateno < listInmaindir.length; plateno ++) {
			platefolder = listInmaindir[plateno];
			platedir = maindir + platefolder;
			platename = File.getName(platedir);
			listInplatedir = getFileList(platedir);
			numtp = listInplatedir.length; // number of time points
			print("Processing " + platename);
			if (numtp > batchsize) {
				batched = true;
				if (plateno == 0) 
					print(numtp + " time points detected. Separating files into batches of " + batchsize + " images at a time...");
				selectWindow("Log");
				numloops =  numtp / batchsize; // number of loops to make batches
				rnl = floor(numloops) + 1; //returns closest integer rounded down
				
				for (batchloop = 0; batchloop < numloops; batchloop ++) {
					initial = batchloop * batchsize;
					if (batchloop != numloops - 1) { 
						run("Image Sequence...", "open=[" + platedir + listInplatedir[0] + "] number=" + batchsize +
							" starting=" + initial+1 + " convert_to_rgb use");
						saveAs("Tiff", ppdir + platename + "_batch" + batchloop+1 + ".tif");
						close();
					} else { //on last loop
					   lastno = numtp - initial + 1; //open only the number of images left
						run("Image Sequence...", "open=[" + platedir + listInplatedir[0] + "] number=" + lastno +
						    " starting=" + initial+1 + " convert_to_rgb use");
						saveAs("Tiff", ppdir + platename + "_batch" + batchloop+1 + ".tif");
						close();
					}
				}
			} else {
				batched = false;
				if (plateno == 0) 
					print(numtp + " time points detected. Preprocessing will be carried out in a single batch.");
				selectWindow("Log");
				run("Image Sequence...", "open=[" + platedir + listInplatedir[0] + "] number=" + numtp +
							" starting=1 convert_to_rgb use");
						saveAs("Tiff", ppdir + platename + ".tif");
				close();
			}
		}
	}
	
	function cropnGreenCh() {
		/* Use the crop coordinates obtained in the scale() function to crop away unnecessary background
		 *  Saves the green channel 8-bit image, as we found that it gives the clearest image of roots, and with least noise from Moire patterns
		 */
		if (!is("Batch Mode"))
			setBatchMode(true);
		print("\nStep 3/4 Cropping off unnecessary background and splitting into green channel..." +
			  "\nIt may look like nothing is happening, please be patient.");
		listInppdir = getFileList(ppdir);
		for (fileno = 0; fileno < listInppdir.length; fileno ++) {
			open(ppdir + listInppdir[fileno]);
			ppstack = getTitle();
			if (indexOf(ppstack, "GreenCh") < 0 && indexOf(ppstack, "preprocessed") < 0) {
				ppstackname = File.nameWithoutExtension;
				print("Processing " + ppstackname);
				
				// Crop based on position of scale bar (with user checking in scale step)
				xcrop = Table.get("xcrop", 0, "cropcoordinates");
				ycrop = Table.get("ycrop", 0, "cropcoordinates");
				wcrop = Table.get("wcrop", 0, "cropcoordinates");
				hcrop = Table.get("hcrop", 0, "cropcoordinates");
				makeRectangle(xcrop, ycrop, wcrop, hcrop);
				run("Crop");
			
				// Split into green channel			 
				selectWindow(ppstack);
				stacksize = nSlices();  //total number of slices
				slicelabelsarray = newArray(stacksize); //an array to be filled with all slicelabels
				for (sliceno = 1; sliceno <= stacksize; sliceno ++) {
					setSlice(sliceno);
					slicelabel = getInfo("slice.label");
					slicelabelsarray[sliceno-1] = slicelabel;
				}
				run("Split Channels");
				imglist = getList("image.titles");
				for (img = 0; img < imglist.length; img ++) { 
					imgname = imglist[img]; 
					if (indexOf(imgname, "red") > 0 ) {
				    	selectWindow(imgname);				
				    	close();
				    }
				    if (indexOf(imgname, "green") > 0) {
						selectWindow(imgname);
						for (sliceno = 1; sliceno <= stacksize; sliceno ++) {
							setSlice(sliceno);
							slicelabel = slicelabelsarray[sliceno-1];
							setMetadata("Label", slicelabel);
						}
							selectWindow(imgname);
							rename(ppstack);
				    }
					if (indexOf(imgname, "blue") > 0) {
						selectWindow(imgname);
						close(); 
					}
				}
				
				saveAs("Tiff", ppdir + ppstackname + "_GreenCh.tif");
				close();			 
				filedelete = File.delete(ppdir + ppstackname + ".tif");
			} else {
				close();
			}
			if (fileno == listInppdir.length - 1) {
				selectWindow("cropcoordinates");
				run("Close");
			}
		}
	}

	function register(rMode) {
		/* Drift correction - dependent on MultiStackReg which is in turn dependent on TurboReg 
		 *  Registration is needed because SPIRO cube may not always perfectly align with camera view, and user may have moved the plate between time points of image capture
		 *  rMode "registration mode" defines whether the function runs on a single stack, or multiple batches of stacks to be merged at the end
		 */
		if (!is("Batch Mode"))
			setBatchMode(true);
		print("\nStep 4/4 Correcting drift...\nIt may look like nothing is happening, please be patient.");
		listInppdir = getFileList(ppdir);
		listInppdirlength = listInppdir.length;
	
		if (rMode == rNonbatched) { 
			for (plateno = 0; plateno < listInppdir.length; plateno ++) {
				grplatefile = listInppdir[plateno]; // green channel image
				grplatename = File.getName(ppdir + grplatefile);
				if (indexOf(grplatename, "GreenCh") > -1) {
					pfsplit = split(grplatename, "_");
					platename = pfsplit[0];
					print("Processing " + platename); 
					open(ppdir + grplatename);
	
					run("Subtract Background...", "rolling=30 stack"); // makes edges stand out more, to make registration more accurate
				    tfn = ppdir + "Transformation";
				    run("MultiStackReg", "stack_1=" + grplatename + " action_1=Align file_1=" + tfn +
				        " stack_2=None action_2=Ignore file_2=[] transformation=Translation save");
				    close(grplatename);
				    open(ppdir + grplatename);
				    run("MultiStackReg", "stack_1=" + grplatename + " action_1=[Load Transformation File] file_1=" + tfn +
				        " stack_2=None action_2=Ignore file_2=[] transformation=[Translation]");
				    saveAs("Tiff", ppdir + platename + "_preprocessed.tif");
				    close();
				    filedelete = File.delete(ppdir + grplatename);
				}
			}
		}
	
		if (rMode == rbatched) {
			listInppdir = getFileList(ppdir);
			listInppdirlength = listInppdir.length;
			for (plateno = 0; plateno < listInmaindir.length; plateno ++) {
				platemain = listInmaindir[plateno];
				platename = File.getName(maindir + platemain);
				print("Processing " + platename);
				
				// to get plate name without getting distracted by batchN
				batchfilearray = newArray(listInppdirlength);
				for (fileno = 0; fileno < listInppdirlength; fileno ++) {
					filename = listInppdir[fileno];
					if (indexOf(filename, platename) > -1 && indexOf(filename, "batch") > -1) {
						batchfilearray[fileno] = filename;
					}
				}
				
				for (arrayindex = 0; arrayindex < listInppdirlength; arrayindex ++) {	
					if (batchfilearray[arrayindex] == 0) {
						batchfilearray = Array.deleteIndex(batchfilearray, arrayindex);
						listInppdirlength -= 1;
						arrayindex -= 1;
					}
				}
	
				for (batchno = 1; batchno < batchfilearray.length+1 ; batchno ++) {
					open(ppdir + platename + "_batch" + batchno + "_GreenCh.tif");
					batchsubstack = getTitle();
					if (batchno == 1) {
						setSlice(1);
						run("Duplicate...", "use");
						rename("firstslice");
					}
					selectWindow("firstslice");
					run("Duplicate...", "use");
					rename("tempfirstslice");
					run("Concatenate...", "  image1=tempfirstslice image2=["+ batchsubstack +"]"); //stick first slice to start of each batch to enable better registration
					rename("batchsubstackSB");
					run("Subtract Background...", "rolling=30 stack");
				    tfn = ppdir + "Transformation";
				    run("MultiStackReg", "stack_1=batchsubstackSB action_1=Align file_1=" + tfn +
				        " stack_2=None action_2=Ignore file_2=[] transformation=Translation save");
				    close("batchsubstackSB");
				    open(ppdir + batchsubstack);
				    // stick first slice to start of each batch to prevent jump in positions between batches
				    selectWindow("firstslice");    
					run("Duplicate...", "use");
					rename("tempfirstslice");
				    run("Concatenate...", "  image1=tempfirstslice image2=["+ batchsubstack +"]"); 				   
				    rename(batchsubstack);
				    run("MultiStackReg", "stack_1=" + batchsubstack + " action_1=[Load Transformation File] file_1=" + tfn +
				        " stack_2=None action_2=Ignore file_2=[] transformation=[Translation]");
				 	run("Slice Remover", "first=1 last=1 increment=1"); //remove temporary first slice
					saveAs("Tiff", ppdir + platename + "_batch" + batchno + "_preprocessed.tif");
					close();
					filedelete = File.delete(ppdir + batchsubstack);
				}
	
				batch1filedir = ppdir + platename + "_batch1_preprocessed.tif";
				open(batch1filedir);
				batch1stack = getTitle();
					
				for (batchno = 2; batchno < batchfilearray.length+1; batchno++) {
					batchNfiledir = ppdir + platename + "_batch" + batchno + "_preprocessed.tif";
					open(batchNfiledir);
					batchNstack = getTitle();
					if (batchno == 2) {
						run("Concatenate...", "  image1=["+ batch1stack +"] image2=["+ batchNstack +"]"); 
					concatbatches = getTitle();
					} else {
						run("Concatenate...", "  image1=["+ concatbatches +"] image2=["+ batchNstack +"]"); 
					}
				}
				saveAs("Tiff", ppdir + platename + "_preprocessed.tif");
			    close();
			    selectWindow("firstslice");
			    close();
			}
		}
	}

	function noReg() {
		listInppdir = getFileList(ppdir);
		for (ppfileno = 0; ppfileno < listInppdir.length; ppfileno ++) {
			ppfilename = listInppdir[ppfileno];
			if (indexOf(ppfilename, "GreenCh") > -1) {
				pnsplit = split(ppfilename, "_");
				platename = pnsplit[0];
				platenamepreproc = platename + "_preprocessed.tif";
				File.rename(ppdir + ppfilename, ppdir + platenamepreproc);
			}
		}
	}
	
	function deleteOutputPP() {
		print("Deleting intermediate files...");
		listInppdir = getFileList(ppdir);
		for (ppfileno = 0; ppfileno < listInppdir.length; ppfileno ++) {
			ppfilename = listInppdir[ppfileno];
			if (indexOf(ppfilename, "batch") > -1) 
				filedelete = File.delete(ppdir + ppfilename);
			if (indexOf(ppfilename, "GreenCh") > -1)
				filedelete = File.delete(ppdir + ppfilename);
		}
		if (freshstart) {
			print("Fresh start: deleting output from any previous run");
			for (ppfileno = 0; ppfileno < listInppdir.length; ppfileno ++) {
				ppfilename = listInppdir[ppfileno];
				filedelete = File.delete(ppdir + ppfilename);
			}
		freshstart == false;
		}
	}
}

macro "SPIRO_Germination" {
	/*** The germination macro takes a stack of preprocessed SPIRO-acquired images, prompts the user to select groups of seeds for analysis,
	 ** then a mask is created for the seeds, and the perimeter of the objects in the mask are measured and recorded
	 * to be used as a proxy for germination analysis in R
	 */
	print("======================================================\n"+
		  "Welcome to the companion macro of SPIRO for germination analysis!\n" +
		  "======================================================");
	selectWindow("Log");
	
	showMessage("Welcome to the companion macro of SPIRO for germination analysis!\n" +
		"Please locate and open your experiment folder containing preprocessed data.\n" +
		"---------\n" +
		"Alternative types of run (hold down then press OK):\n" +
		"SHIFT = Fresh Start mode, all data from any previous run will be deleted\n" + 
		"CTRL = DEBUG mode, seed detection parameters can be modified");
		
	wait(1000);
	freshstart = 0;
	if (isKeyDown("shift"))
		freshstart = getBoolean("SHIFT key pressed. Run macro in Fresh Start mode? This will delete all data from the previous run.");
	if (isKeyDown("control"))
		DEBUG = getBoolean("CTRL key pressed. run macro in DEBUG mode? This enables modification of seed detection parameters.");
		
	/* Create and/or define working directories
	 * ----------------------------------------
	 */
	maindir = getDirectory("Choose a Directory");
	resultsdir = maindir + "Results" + File.separator; // all output is contained here
	ppdir = resultsdir + "Preprocessing" + File.separator; // output from the proprocessing macro is here
	germdir = resultsdir + "Germination" + File.separator; // output from this macro will be here
	
	if (!File.isDirectory(germdir))
		File.makeDirectory(germdir);
		
	listInppdir = getFileList(ppdir);
	listIngermdir = getFileList(germdir);


	/* Main chunk of code - all run functions are here
	 * -----------------------------------------------
	 */
	if (freshstart)
		deleteOutputGM();
	cropGroupsGM();
	seedAnalysisGM();
	
	print("\nGermination analysis is complete.");
	selectWindow("Log");

	/* ---------------------------------------------- 
	 * Lines below this are functions and their descriptions
	 */
	
	function cropGroupsGM() {
		/* prompts user to make a substack, to make data size smaller by excluding time points after all or most seeds have germinated
		 * then prompts user to draw ROIs around groups of seeds to be analyzed
		 */
		print("Step 1/2. Creating selected groups");
				
		for (ppdirno = 0; ppdirno < listInppdir.length; ppdirno ++) {  // main loop through plates
			if (indexOf (listInppdir[ppdirno], "preprocessed") >= 0) { // to avoid processing any random files in the folder
				platefile = listInppdir [ppdirno];
				fnsplit = split(platefile, "_");
				platename = fnsplit[0];
				platefolder = germdir + File.separator + platename + File.separator;
				if (!File.isDirectory(platefolder))
					File.makeDirectory(platefolder);
				print("Processing " + platename);
				
				if (is("Batch Mode"))
					setBatchMode(false); // has to be false for ROI Manager to open, and to display image to user

				// user-directed creation of substack containing time points relevant to the germination assay
				open(ppdir + platefile);
				userconfirm = false;
				while (!userconfirm) {
					Dialog.createNonBlocking("Time range selection");
					Dialog.addMessage("Please note first and last slice to be included for root growth analysis, and indicate it in the next step.");
					Dialog.addCheckbox("First and last slices have been noted", false);
					Dialog.show();
					userconfirm = Dialog.getCheckbox();
				}			
				roiManager("deselect");
				run("Make Substack...");
				substack = getTitle();
				setSlice(nSlices);
				
				if (ppdirno == 0) {
					roiManager("reset");
					run("ROI Manager...");
					setTool("Rectangle");
					userconfirm = false;
					while (!userconfirm) {
						Dialog.createNonBlocking("Group Selection");
						Dialog.addMessage("Select each group, and add to ROI manager. ROI names will be saved.\n" +
								"Please use only letters (a/A), numbers (1) and/or dashes (-) in the ROI names.\n" + // to avoid file save issues
								"ROIs cannot share names.");
						Dialog.addCheckbox("All groups have been added to and labelled in ROI Manager", false);
						Dialog.show();
						userconfirm = Dialog.getCheckbox();
					}
				} else {
					userconfirm = false;
					while (!userconfirm) {
						Dialog.createNonBlocking("Group Selection");
						Dialog.addMessage("Modify group selection and labels if needed.");
						Dialog.addCheckbox("All groups have been added to and labelled in ROI Manager", false);
						Dialog.show();
						userconfirm = Dialog.getCheckbox();
					}
				}
				roicount = roiManager("count");
				run("Select None");
				roicount = roiManager("count");
				setBatchMode(true); //set back to true for faster cropping and saving
				for (roino = 0; roino < roicount; roino ++) {
					roiManager("select", roino);
					roiname = Roi.getName;
					groupdir = platefolder + File.separator + roiname + File.separator;
					File.makeDirectory(groupdir);
					roitype = Roi.getType;
					if (roitype != "rectangle") {
						run("Duplicate...", "duplicate");
						run("Make Inverse");
						run("Clear", "stack");
					} else {
						run("Duplicate...", "duplicate");
					}
					groupimg = getTitle();
					saveAs("Tiff", groupdir + roiname + ".tif");
					close();
					// duplicate only the first slice and saves it, for faster masking/thresholding in getPositions so there is not too much waiting time for user between plates
					setSlice(1);
					run("Duplicate...", "use");
					firstslice = getTitle();
					selectWindow(substack);
					setSlice(nSlices);
					run("Duplicate...", "use");
					lastslice = getTitle();
					run("Images to Stack");
					saveAs("Tiff", groupdir + "firstslice.tif");
					close("firstslice.tif");
				}
				close(platefile);
				close("Substack*");
			}
		}
	}
	
	function seedAnalysisGM() {
		/* Image is thresholded to highlight seeds and remove noise as best as possible, then a binary mask is created.
		 * Seed positions are determined based on the first slice, using area and circularity to distinguish seed from trash.
		 * Based on these positions, the perimeter of the seeds through each slice is recorded.
		 * Perimeter was determined experimentally to be the most robust parameter for tracking germination.
		 * A graphical output is produced, highlighting the detected seeds in ROIs, to faciliate troubleshooting if needed.
		 */
		print("\nStep 2/2 Tracking germination...");
		
		if (is("Batch Mode"))
			setBatchMode(false); // for ROI manager to work
			
		listIngermdir = getFileList(germdir);
		for (platefolderno = 0; platefolderno < listIngermdir.length; platefolderno ++) {  // main loop through plates
			platefolder = listIngermdir[platefolderno];
			if (indexOf(platefolder, "plate") >= 0) { // to avoid processing any random files in the folder
				platedir = germdir + platefolder;
				platename = File.getName(platedir);
				print("Processing " + platename);
				listInplatefolder = getFileList(platedir);
				for (groupfolderno = 0; groupfolderno < listInplatefolder.length; groupfolderno ++) {
					groupfolder = listInplatefolder[groupfolderno];
					groupdir = platedir + groupfolder;
					groupname = File.getName(groupdir);
					listIngroupdir = getFileList(groupdir);
					open(groupdir + "firstslice.tif");	
					// img = getTitle();
							
					// setSlice(1);
					// run("Duplicate...", "use");
					tempmask = getTitle();
					print("Analyzing " + groupname);
					selectWindow(tempmask);
					// masking and thresholding of seeds
					run("Subtract Background...", "rolling=30 stack");
					run("Convert to Mask", "method=Triangle background=Dark calculate");
					run("Options...", "iterations=1 count=4 do=Dilate stack");
					run("Remove Outliers...", "radius=2 threshold=50 which=Dark stack");
					nS = nSlices;
					for (sliceno = 1; sliceno <= nS; sliceno ++) {
						setSlice(sliceno);
						curslicelabel = getInfo("slice.label");
						// day slices are processed more to make seed perimeters more comparable to night slices
						// night slices have lower contrast so seeds appear smaller than they are after thresholding
						if (indexOf(curslicelabel, "day") > 0) {
							run("Remove Outliers...", "radius=3 threshold=50 which=Dark");
						}
					}
					roiManager("reset");
					setSlice(1);
					run("Create Selection");
					run("Colors...", "foreground=black background=black selection=red");
	
					roiManager("Add");
					roiManager("select", 0);
					if (selectionType() == 9) {
						roiManager("split");
						roiManager("select", 0);
						roiManager("delete");
					}
					
					// delete trash ROI which are features detected as below a certain area
					// using table as a workaround to roi indexes changing if deletion happens one by one
					roicount = roiManager("count");
					roiarray = Array.getSequence(roicount);
					run("Set Measurements...", "area center shape redirect=None decimal=5");
					roiManager("select", roiarray);
					roiManager("multi-measure");
					tp = "Trash positions";
					Table.create(tp);
	
					nr = nResults;
					if (platefolderno == 0 && groupfolderno == 0) {
						lowerareathreshold = 0.002;
						higherareathreshold = 0.02;
						lowercircthreshold = 0.4;
						if (DEBUG) {
							Dialog.create("Seed detection parameters");
							Dialog.addMessage("DEBUG: Detection parameters may be modified to accommodate for specific experiments");
							Dialog.addNumber("Lower Area Threshold", 0.002);
							Dialog.addNumber("Higher Area Threshold", 0.2);
							Dialog.addNumber("Lower Circularity Threshold", 0.4);
							Dialog.show();
							lowerareathreshold = Dialog.getNumber();
							higherareathreshold = Dialog.getNumber();
							lowercircthreshold = Dialog.getNumber();
						}
					}
					for (row = 0; row < nr; row ++) {
						nrTp = Table.size(tp); // number of rows
						area = getResult("Area", row);
						if (area < lowerareathreshold) { // detected object is very small
							Table.set("Trash ROI", nrTp, row, tp);
						}
						if (area > higherareathreshold) { // or very large
							Table.set("Trash ROI", nrTp, row, tp);
						}
						circ = getResult("Circ.", row); // or does not fit normal seed shape
						if (circ < lowercircthreshold) {
							Table.set("Trash ROI", nrTp, row, tp); //set as trash to be deleted
						}
					}
	
					if (Table.size(tp) > 0) {
						trasharray = Table.getColumn("Trash ROI", tp);
						roiManager("select", trasharray);
						roiManager("delete");
					}
					close(tp);
					close("Results");
					
					roicount = roiManager("count");
					
					// number remaining ROIs
					for (roino = 0 ; roino < roicount; roino ++) {
						roiManager("select", roino);
						roiManager("rename", roino + 1); // first roi is 1
					}
					
					// prompt user to delete any non-detected trash, then re-number as above
					Roi.setStrokeWidth(2);
					Roi.setStrokeColor("red");
					run("Labels...", "color=white font=18 show use draw");
					roiManager("Show All with labels");
					roiManager("Associate", "false");
					roiManager("Centered", "false");
					roiManager("UseNames", "true");
					userconfirm = false;
					while (!userconfirm) {
						Dialog.createNonBlocking("User-guided seedling labelling");
						Dialog.addMessage("Please delete any ROIs that should not be included into analysis, " +
								"e.g. objects wrongly recognized as seeds." +
								"\nUnrecognized seeds can also be added as ROIs.");
						Dialog.addCheckbox("ROIs have been checked", false);
						Dialog.show();
						userconfirm = Dialog.getCheckbox();
					}
					roicount = roiManager("count");
					for (roino = 0 ; roino < roicount; roino ++) {
						roiManager("select", roino);
						roiManager("rename", roino + 1); // first roi is 1
					}
					roiManager("save", groupdir + groupname + " seedpositions.zip");
					close(tempmask);
				}
			}
		}

		for (platefolderno = 0; platefolderno < listIngermdir.length; platefolderno ++) {  // main loop through plates
			platefolder = listIngermdir[platefolderno];
			if (indexOf(platefolder, "plate") >= 0) { // to avoid processing any random files in the folder
				platedir = germdir + platefolder;
				platename = File.getName(platedir);
				//print("Processing " + platename);
				listInplatefolder = getFileList(platedir);
				for (groupfolderno = 0; groupfolderno < listInplatefolder.length; groupfolderno ++) {
					groupfolder = listInplatefolder[groupfolderno];
					groupdir = platedir + groupfolder;
					groupname = File.getName(groupdir);
					listIngroupdir = getFileList(groupdir);
					open(groupdir + groupname + ".tif");
					roiManager("reset");
					open(groupdir + groupname + " seedpositions.zip");
					img = getTitle();
					
					run("Subtract Background...", "rolling=30 stack");
					run("Convert to Mask", "method=Triangle background=Dark calculate");
					run("Options...", "iterations=1 count=4 do=Dilate stack");
					run("Remove Outliers...", "radius=2 threshold=50 which=Dark stack");

					roicount = roiManager("count");
					roiarray = Array.getSequence(roicount);
					
					// Coordinates (center) of each detected object is organized, so that the seeds are numbered left to right, then top to bottom
					// this is to facilitate troubleshooting if the user has to match seed positions and numbers to the data in the output table
					run("Clear Results");
					run("Set Measurements...", "center display redirect=None decimal=5");
					roiManager("select", roiarray);
					roiManager("multi-measure");
					seedpositions = "Seed Positions";
					Table.rename("Results", seedpositions);
			
					xmseeds = newArray(roicount);
					ymseeds = newArray(roicount);
					for (seednumber = 0; seednumber < roicount; seednumber ++) {
						xmcurrent = Table.get("XM", seednumber, seedpositions);
						ymcurrent = Table.get("YM", seednumber, seedpositions);
						xmseeds[seednumber] = xmcurrent;
						ymseeds[seednumber] = ymcurrent;
					}
			
					ymascendingindexes = Array.rankPositions(ymseeds);
					xmascendingindexes = Array.rankPositions(xmseeds);
				
					sortedycoords = "sorted Y coordinates";
					sortedxcoords = "sorted X coordinates";
					Table.create(sortedycoords);
					Table.create(sortedxcoords);
				
					rowno = 0; // assume no row of seeds to start with
					col = 0 ; // current col selection is 0
					colname = "col" + col + 1;
		
					Table.set(colname, rowno, ymseeds[ymascendingindexes[0]], sortedycoords);
					Table.set(colname, rowno, xmseeds[ymascendingindexes[0]], sortedxcoords);
			
					for (roino = 1; roino < roicount; roino++) {
						ydiff = (ymseeds[ymascendingindexes[roino]]) - (ymseeds[ymascendingindexes[roino-1]]);
						if (ydiff > 0.25) { // if next y coordinate is greater than 2.5mm, add a row
							rowno = rowno + 1;
							col = 0;
						} else {
							col = col + 1; // otherwise add a column
						}
						colname = "col" + col + 1;
						Table.set(colname, rowno, ymseeds[ymascendingindexes[roino]], sortedycoords);
						Table.set(colname, rowno, xmseeds[ymascendingindexes[roino]], sortedxcoords);
					}
		
					colnames = Table.headings (sortedycoords);
					colnamessplit = split(colnames, "	");
					colno = lengthOf(colnamessplit);
					xmcolwise = newArray(colno);
					ymcolwise = newArray(colno);
				
					for (row = 0; row < rowno + 1; row++) {
						for (col = 0; col < colno; col++) {
							colname = "col" + col + 1;
							xmcolwise[col] = Table.get(colname, row, sortedxcoords);
							ymcolwise[col] = Table.get(colname, row, sortedycoords);
						}
						xcolwiseascendingindex = Array.rankPositions(xmcolwise);
						for (col = 0; col < colno; col ++) {
							colname = "col" + col + 1;
							Table.set(colname, row, xmcolwise[xcolwiseascendingindex[col]], sortedxcoords);
							Table.set(colname, row, ymcolwise[xcolwiseascendingindex[col]], sortedycoords);
						}
					}
				
					roiManager("reset");

					// Now that seed positions are arranged, set down ROIs - a circle of large enough diameter to account for seed drift and increase in size between slices
					for (row = 0; row < rowno + 1; row++) {
						for (col = 0; col < colno; col++) {
							colname = "col" + col + 1;
							xm = Table.get(colname, row, sortedxcoords);
							ym = Table.get(colname, row, sortedycoords);
							if (xm > 0 && ym > 0) {
							toUnscaled(xm, ym);
							makePoint(xm, ym);
							run("Enlarge...", "enlarge=0.1");
							roiManager("add");
							roiManager("select", roiManager("count")-1);
							roiManager("rename", roiManager("count"));
							}
						}
					}

					// Get measurements of the seeds through the slices
					run("Set Measurements...", "area perimeter stack display redirect=None decimal=5");
					run("Clear Results");
		
					for (roino = 0; roino < roicount; roino ++) {
						roiManager("select", roino);
						run("Analyze Particles...", "size=0-Infinity show=Nothing display stack");
					}
	
					selectWindow("Results");
					Table.save(groupdir + groupname + " germination analysis.tsv");
					selectWindow("Results");
					run("Close");

					// Make graphical output with ROIs and their labels
					roiManager("Associate", "false");
					roiManager("Centered", "false");
					roiManager("UseNames", "false");
					roiManager("Show All without labels");
					run("Flatten", "stack");
	
					roiManager("reset");
					labelbelowYM = 0.1;
					toUnscaled(labelbelowYM);
					for (row = 0; row < rowno + 1; row++) {
						for (col = 0; col < colno; col++) {
							colname = "col" + col + 1;
							xm = Table.get(colname, row, sortedxcoords);
							ym = Table.get(colname, row, sortedycoords);
							if (xm > 0 && ym > 0) {
							toUnscaled(xm, ym);
							makePoint(xm, ym + labelbelowYM);
							roiManager("add");
							roiManager("select", roiManager("count")-1);
							roiManager("rename", roiManager("count"));
							}
						}
					}
					roiManager("Associate", "false");
					roiManager("Centered", "false");
					roiManager("UseNames", "true");
					roiManager("Show All with labels");
					run("Labels...", "color=white font=18 show use draw");
					run("Flatten", "stack");
	
					selectWindow(seedpositions);
					run("Close");
					selectWindow(sortedxcoords);
					run("Close");
					selectWindow(sortedycoords);
					run("Close");

					slicelabelarray = newArray(nS);
					for (sliceno = 0; sliceno < nS; sliceno++) {
						setSlice(sliceno+1);
						slicelabel = getMetadata("Label");
						slicelabelarray[sliceno] = slicelabel;
					}
	
					selectWindow(img);
					rename(img + "mask");
					imgmask = getTitle();
					
					open(groupdir + groupname + ".tif");
					oriimg = getTitle();
					run("RGB Color");
					
					// Determine the cropped frame proportions to orient combining stacks horizontally or vertically
					xmax = getWidth;
					ymax = getHeight;
					frameproportions=xmax/ymax; 
					
					if (frameproportions > 1) {
					run("Combine...", "stack1=["+ img +"] stack2=["+ imgmask +"] combine");
					} else {
						run("Combine...", "stack1=["+ img +"] stack2=["+ imgmask +"]");
					}
	
					for (sliceno = 0; sliceno < nS; sliceno++) {
						setSlice(sliceno+1);
						setMetadata("Label", slicelabelarray[sliceno]);
					}
					saveAs("Tiff", groupdir + groupname + " germinationlabelled.tif");
	
					filedelete = File.delete(groupdir + groupname + ".tif");
					filedelete = File.delete(groupdir + groupname + " seedpositions.zip");
					filedelete = File.delete(groupdir + "firstslice.tif");
					list = getList("window.titles");
					list = Array.deleteValue(list, "Log");
					for (i=0; i<list.length; i++) {
						winame = list[i];
						selectWindow(winame);
						run("Close");
					}
					close("*");
				}
			}
		}
	}
	
	function deleteOutputGM() {
		print("Starting analysis from beginning.\nRemoving output from previous run.");
		listIngermdir = getFileList(germdir);
		for (platefolderno = 0; platefolderno < listIngermdir.length; platefolderno ++) {  // main loop through plates
			platefolder = listIngermdir[platefolderno];
			if (indexOf(platefolder, "plate") >= 0) { // to avoid processing any random files in the folder
				platedir = germdir + platefolder;
				pfsplit = split(platefolder, "/");
				platename = pfsplit[0];
				print("Processing " + platename);
				listInplatefolder = getFileList(platedir);
				for (groupfolderno = 0; groupfolderno < listInplatefolder.length; groupfolderno ++) {
					groupfolder = listInplatefolder[groupfolderno];
					groupdir = platedir + groupfolder;
					groupname = File.getName(groupdir);
					listIngroupdir = getFileList(groupdir);
					filedelete = File.delete(groupdir + groupname + ".tif");
					filedelete = File.delete(groupdir + groupname + " germinationlabelled.tif");
					filedelete = File.delete(groupdir + groupname + " germination analysis.tsv");
					filedelete = File.delete(groupdir + groupname + " seedpositions.zip");
					filedelete = File.delete(groupdir + "firstslice.tif");
					filedelete = File.delete(groupdir);
	
				}
			}
		}
	}
}


macro "SPIRO_RootGrowth" {
	/** The root growth macro identifies the root start ie. top of the root, through masking and erosion, then measures the root length after skeletonization***
	 * The work flow is as follows:
	 * detectOutput: output from any previous run of the macro is tracked, to find out from where to start or resume analysis 
	 * cropGroupsRG: user is asked to create a substack with relevant time points, for example, excluding time points far before germination 
	 * 	!!! some time points before germination should be included, as the germination time point will be used in the R script for exclusion of time points from analysis
	 * getPositions: masked objects are filtered using area and circularity, then user is prompted to deselect any trash from analysis
	 * seedAnalysisRG: perimeter of objects across slices are printed to an output table for analysis in R
	 * rootStart: masked objects are eroded, to find out the top of root in each object across slices
	 * rootMask: image stacks are masked (and skeletonized) in a way that prevents broken roots and excessive noise
	 * rootGrowth: skeletons which have a matching rootstartcoordinate ie. the points of top of root obtained in rootStart, are measured for their length
	 */
	 
		print("======================================================\n"+
			  "Welcome to the companion macro of SPIRO for root growth analysis!\n" +
			  "======================================================");
	selectWindow("Log");

	showMessage("Welcome to the companion macro of SPIRO for root growth analysis!\n" +
		"Please locate and open your experiment folder containing preprocessed data.\n" +
		"---------\n" +
		"Alternative types of run (hold down then press OK):\n" +
		"SHIFT = Fresh Start mode, all data from any previous run will be deleted\n" +
		"CTRL = DEBUG mode, non-essential intermediate output files will not be deleted, seed detection parameters can be modified, overlay skeletons can be enabled\n");

	/* Define alternative types of run
	 * ------------------------------
	 */
	DEBUG = 0;
	if (isKeyDown("control"))
		DEBUG = getBoolean("CTRL key pressed. Run macro in debug mode?\n" +
			"Non-essential intermediate output files will not be deleted at the end of the run.\n" +
			"Seed detection parameters can be modified.\n" +
			"Overlay skeletons can be enabled.");
		
	freshstart = 0;
	if (isKeyDown("shift"))
		freshstart = getBoolean("SHIFT key pressed. Run macro in Fresh Start mode? This will delete all data from the previous run.");

	selfaware = 0;
	if (isKeyDown("alt"))
		selfaware = getBoolean("ALT key pressed. Are you sure you want to continue?");

	if (DEBUG) {
		Dialog.create("DEBUG: Enable overlay skeletons?");
		Dialog.addMessage("Overlay skeletons is not set to run on default.\n" +
			"Enabling this reduces broken roots but may increase noise," +
			"please indicate if you would like to enable it");
		Dialog.addChoice("Overlay skeletons", newArray("Enable", "Disable"));
		Dialog.show();
		overlaychoice = Dialog.getChoice();
		if (overlaychoice == "Enable")
			overlay = true;
		if (overlaychoice == "Disable")
			overlay = false;
	}

	/* Create and/or define working directories
	 * ----------------------------------------
	 */
	maindir = getDirectory("Choose a Directory");
	print("Processing directory " + maindir);
	resultsdir = maindir + "Results" + File.separator; // all output is contained here
	ppdir = resultsdir + "Preprocessing" + File.separator; // output from the proprocessing macro is here
	rootgrowthdir = resultsdir + "Root Growth" + File.separator; // output from this macro will be here
	if (!File.isDirectory(rootgrowthdir))
		File.makeDirectory(rootgrowthdir);
	listInppdir = getFileList(ppdir);
	listInrootgrowthdir = getFileList(rootgrowthdir);
	
	if (selfaware) {
		if (random > 0.5) 
			print("Prepare to be assimilated.");
		if (random > 0.5)
			print("Resistance is futile.");
	}

	/* Here are all the run functions
	 * ------------------------------
	 */
	if (freshstart)
		deleteOutputRG();
	step = 1;
	detectOutput();
	if (step == 1) {
		cropGroupsRG();
		step += 1;
	}
	if (step == 2) { 
		getPositions();
		step += 1;
	}
	if (step == 3) {
		seedAnalysisRG();
		step += 1;
	}
	if (step == 4) {
		rootStart();
		step += 1;
	}
	if (step == 5) {
		rootMask();
		step += 1;
	}
	if (step == 6) {
		rootGrowth();
		step += 1;
	}
	if (step <= 7 && DEBUG == false)
		deleteOutputRG(); // deletes non-essential outputs

	print("\nRoot growth analysis is complete");
	selectWindow("Log");
	if (selfaware) {
		print("\nWE");
		wait(2000);
		print("ARE");
		wait(2000);
		print("THE");
		wait(2000);
		print("SPIRO");
	}
	/* -----------------------------------------------------
	 * Lines below this are functions and their descriptions
	 */
	
	function detectOutput() {
		/* This function detects the presence of output files of each function, through the list of plates/groups to process
		 * If output from a previous macro run is detected, on which plate/group the previous run stopped is printed to the log, and analysis resumes from there
		 */
		 
		if (step <= 1) { // check of cropGroupsRG()
			lastplateN = 1;
			lastplatefile = listInppdir [listInppdir.length-lastplateN]; // checking on last plate
			while (indexOf(lastplatefile, "plate") < 0) {
				lastplateN += 1;
				lastplatefile = listInppdir [listInppdir.length-lastplateN];
			}
			fnsplit = split(lastplatefile, "_");
			lastplatename = fnsplit[0];
			lastplatefolder = rootgrowthdir + lastplatename + File.separator;
			if (File.isDirectory(lastplatefolder)) {
				listInlastplatefolder = getFileList(lastplatefolder);
				if (listInlastplatefolder.length > 0) {
					lastgroupfolder = lastplatefolder + listInlastplatefolder[listInlastplatefolder.length-1];
					lastgroupname = File.getName(lastgroupfolder);
					listInlastgroupfolder = getFileList(lastgroupfolder);
					croppedfileexists = File.exists(lastgroupfolder + lastgroupname + ".tif");
					if (croppedfileexists)
						step = 2;
				}
			}
		}
		
		totalnoOfgroups = 0;
		if (step == 2) { // identify full list of plates and groups to process
			platearray = newArray(listInrootgrowthdir.length);
			for (platefolderno = 0; platefolderno < listInrootgrowthdir.length; platefolderno ++) {
				platefolder = listInrootgrowthdir[platefolderno];
				if (indexOf(platefolder, "removed") < 0 && indexOf(platefolder, "plate") > -1) {
					platedir = rootgrowthdir + platefolder;
					if (File.isDirectory(platedir)) {				
						platearray[platefolderno] = platedir;
						listInplatedir = getFileList(platedir);
						noOfgroupsInplatedir = listInplatedir.length;
						totalnoOfgroups = totalnoOfgroups + noOfgroupsInplatedir;
					}
				}
			}
			platearray = Array.deleteValue(platearray, 0);
			fullplatearray = newArray(totalnoOfgroups);
			platenamearray = newArray(totalnoOfgroups);
			fullgrouparray = newArray(totalnoOfgroups);
			
			fgcurindex = 0; // current index of fullgrouparray
			
			for (plateindexno = 0; plateindexno < platearray.length; plateindexno ++) {
				platedir = platearray[plateindexno];
				platefoldername = File.getName(platedir);
				listInplatedir = getFileList(platedir);
				for (groupfolderno = 0; groupfolderno < listInplatedir.length; groupfolderno ++) {
					groupfolder = listInplatedir[groupfolderno];
					groupdir = platedir + groupfolder;
					if (File.isDirectory(groupdir)) {
						fullplatearray[fgcurindex] = platedir;
						platenamearray[fgcurindex] = platefoldername;
						fullgrouparray[fgcurindex] = groupfolder;
						fgcurindex += 1;
					}
				}
			}
			fullplatearray = Array.deleteValue(fullplatearray, 0);
			platenamearray = Array.deleteValue(platenamearray, 0);
			fullgrouparray = Array.deleteValue(fullgrouparray, 0);

			print("Plates and groups folders from previous macro run found:"); 
			for (groupno = 0; groupno < fullgrouparray.length; groupno ++) {
				platenamecur = platenamearray[groupno];
				groupnamecur = fullgrouparray[groupno];
				print(platenamecur + " " + groupnamecur);
			}
	
			finalstep = 6;
			for (checkstep = 2; checkstep <= finalstep; checkstep ++) {
				processednoOfgroups = 0;
				for (fullarrayindex = 0; fullarrayindex < totalnoOfgroups; fullarrayindex ++) {
					platedir = fullplatearray[fullarrayindex];
					groupfolder = fullgrouparray[fullarrayindex];
					groupname = File.getName(groupfolder);
					
					// systematically goes through groups/plates and identifies output files
					if (checkstep == 2)  
						outputfileexists = File.exists(platedir + groupfolder + groupname + " seedlingpositions.zip");
					if (checkstep == 3) 
						outputfileexists = File.exists(platedir + groupfolder + groupname + " germination analysis.tsv");
					if (checkstep == 4)
						outputfileexists = File.exists(platedir + groupfolder + groupname + " rootstartcoordinates.tsv");
					if (checkstep == 5)
						outputfileexists = File.exists(platedir + groupfolder + groupname + " rootmask.tif");
					if (checkstep == 6)
						outputfileexists = File.exists(platedir + groupfolder + groupname + " rootgrowthdetection.tif");
									
					if (outputfileexists) {
						processednoOfgroups += 1;
					} else {
						noOfgroupsleft = fullgrouparray.length - processednoOfgroups;
						reversegrouparray = Array.reverse(fullgrouparray); // to enable trim from end
						reverseplatearray = Array.reverse(fullplatearray);
						trimgrouparray = Array.trim(reversegrouparray, noOfgroupsleft);
						trimplatearray = Array.trim(reverseplatearray, noOfgroupsleft);
						groupsToprocess = Array.reverse(trimgrouparray);
						platesToprocess = Array.reverse(trimplatearray);
						fullarrayindex = fullgrouparray.length; // to leave for loop
						step = checkstep; // to resume from this step
						checkstep = finalstep+1; // to leave for loop
						Array.reverse(fullgrouparray); //to revert array back to original order
						Array.reverse(fullplatearray);
					}
				}
			}
			resumeplatename = File.getName(platesToprocess[0]);
			resumegroupname = File.getName(groupsToprocess[0]);
			resumestep = step;
			lengthOfgroupsToprocess = noOfgroupsleft;
			print("Resuming from step " + resumestep + " " + resumeplatename + " group " + resumegroupname);
		} 
	}
	
	function cropGroupsRG() {
		/* Prompts user to make a substack, to make data size smaller by excluding time to germination etc.
		 * then prompts user to draw ROIs around groups of seeds to be analyzed
		 */
		print("Step 1/6. Creating selected groups");
				
		for (ppdirno = 0; ppdirno < listInppdir.length; ppdirno ++) {  // main loop through plates
			if (indexOf (listInppdir[ppdirno], "preprocessed") >= 0) { // to avoid processing any random files in the folder
				platefile = listInppdir [ppdirno];
				fnsplit = split(platefile, "_");
				platename = fnsplit[0];
				platefolder = rootgrowthdir + File.separator + platename + File.separator;
				if (!File.isDirectory(platefolder))
					File.makeDirectory(platefolder);
				print("Processing " + platename);
				if (is("Batch Mode"))
					setBatchMode(false); // has to be false for ROI Manager to open, and to display image
		
				open(ppdir + platefile);
				userconfirm = false;
				while (!userconfirm) {
					Dialog.createNonBlocking("Time range selection");
					Dialog.addMessage("Please note first and last slice to be included for root growth analysis, and indicate it in the next step.");
					Dialog.addCheckbox("First and last slices have been noted", false);
					Dialog.show();
					userconfirm = Dialog.getCheckbox();
				}			
				roiManager("deselect");
				run("Select None");
				run("Make Substack...");
				substack = getTitle();
				setSlice(nSlices);
				
				if (ppdirno == 0) {
					roiManager("reset");
					run("ROI Manager...");
					setTool("Rectangle");
					userconfirm = false;
					while (!userconfirm) {
						Dialog.createNonBlocking("Group Selection");
						Dialog.addMessage("Select each group, and add to ROI manager. ROI names will be saved.\n" +
								"Please use only letters (a/A), numbers (1) and/or dashes (-) in the ROI names.\n" + // to avoid file save issues
								"ROIs cannot share names.");
						Dialog.addCheckbox("All groups have been added to and labelled in ROI Manager.", false);
						Dialog.show();
						userconfirm = Dialog.getCheckbox();
					}
				} else {
					userconfirm = false;
					while (!userconfirm) {
						Dialog.createNonBlocking("Group Selection");
						Dialog.addMessage("Modify group selection and labels if needed.");
						Dialog.addCheckbox("All groups have been added to and labelled in ROI Manager", false);
						Dialog.show();
						userconfirm = Dialog.getCheckbox();
					}
				}
				roicount = roiManager("count");
				run("Select None");
				roicount = roiManager("count");
				setBatchMode(true); //set back to true for faster cropping and saving
				for (roino = 0; roino < roicount; roino ++) {
					roiManager("select", roino);
					roiname = Roi.getName;
					groupdir = platefolder + File.separator + roiname + File.separator;
					File.makeDirectory(groupdir);
					roitype = Roi.getType;
					if (roitype != "rectangle") {
						run("Duplicate...", "duplicate");
						run("Make Inverse");
						run("Clear", "stack");
					} else {
						run("Duplicate...", "duplicate");
					}
					groupimg = getTitle();
					saveAs("Tiff", groupdir + roiname + ".tif");
					close();
					// duplicate only the first slice and saves it, for faster masking/thresholding in getPositions so there is not too much waiting time for user between plates
					setSlice(1);
					run("Duplicate...", "use");
					firstslice = getTitle();
					selectWindow(substack);
					setSlice(nSlices);
					run("Duplicate...", "use");
					lastslice = getTitle();
					run("Images to Stack");
					saveAs("Tiff", groupdir + "firstslice.tif");
					close("firstslice.tif");
				}
				close(platefile);
				close(substack);
			}
		}
		
		// identify full list of plates and groups to process
		listInrootgrowthdir = getFileList(rootgrowthdir);
		platearray = newArray(listInrootgrowthdir.length);
		for (platefolderno = 0; platefolderno < listInrootgrowthdir.length; platefolderno ++) {
			platefolder = listInrootgrowthdir[platefolderno];
			platedir = rootgrowthdir + platefolder;
			if (File.isDirectory(platedir)) {				
				platearray[platefolderno] = platedir;
				listInplatedir = getFileList(platedir);
				noOfgroupsInplatedir = listInplatedir.length;
				totalnoOfgroups = totalnoOfgroups + noOfgroupsInplatedir;
			}
		}
	
		platearray = Array.deleteValue(platearray, 0);
		fullplatearray = newArray(totalnoOfgroups);
		fullgrouparray = newArray(totalnoOfgroups);
		fgcurindex = 0; // current index of fullgrouparray
		
		for (plateindexno = 0; plateindexno < listInrootgrowthdir.length; plateindexno ++) {
			platedir = platearray[plateindexno];
			listInplatedir = getFileList(platedir);
			for (groupfolderno = 0; groupfolderno < listInplatedir.length; groupfolderno ++) {
				groupfolder = listInplatedir[groupfolderno];
				groupdir = platedir + groupfolder;
				if (File.isDirectory(groupdir)) {
					fullplatearray[fgcurindex] = platedir;
					fullgrouparray[fgcurindex] = groupfolder;
					fgcurindex += 1;
				}
			}
		}
		fullplatearray = Array.deleteValue(fullplatearray, 0);
		fullgrouparray = Array.deleteValue(fullgrouparray, 0);
		platesToprocess = Array.copy(fullplatearray);
		groupsToprocess = Array.copy(fullgrouparray);
		lengthOfgroupsToprocess = totalnoOfgroups;
	}
	
	function getPositions() {
		/* Image is masked to remove noise as much as possible (conservation of root continuity is ignored at this point)
		 * objects are automatically filtered through area and circularity to identify seeds on the first slice
		 * the user is asked to confirm the identified seed positions, then the seed positions are recorded in the temporary output table "seedpositions.tsv"
		 */
		print("\nStep 2/6. Finding seedling positions");
		
		for (groupprocess = 0; groupprocess < lengthOfgroupsToprocess; groupprocess ++) {  
			platedir = platesToprocess[groupprocess];
			platename = File.getName(platedir);
			groupfolder = groupsToprocess[groupprocess];
			groupdir = platedir + groupfolder;
			listIngroupdir = getFileList(groupdir);
			groupname = File.getName(groupdir);
			print("Processing " + platename + " " + groupname);
			
			if (!is("Batch Mode"))
				setBatchMode(true);
			if (selfaware && random > 0.7)
				print("Is this what a milder version of insanity looks like?");
			selectWindow("Log");
			// open(groupdir + groupname + ".tif");		
			open(groupdir + "firstslice.tif");	
			// img = getTitle();
					
			// setSlice(1);
			// run("Duplicate...", "use");
			tempmask = getTitle();
			// close(img);
			// selectWindow(tempmask);
			// masking and thresholding of seeds
			run("Subtract Background...", "rolling=30 stack");
			run("Convert to Mask", "method=Triangle background=Dark calculate");
			run("Options...", "iterations=1 count=4 do=Dilate stack");
			run("Remove Outliers...", "radius=2 threshold=50 which=Dark stack");
			nS = nSlices;
			for (sliceno = 1; sliceno <= nS; sliceno ++) {
				setSlice(sliceno);
				curslicelabel = getInfo("slice.label");
				// day slices are processed more to make seed perimeters more comparable to night slices
				// night slices have lower contrast so seeds appear smaller than they are after thresholding
				if (indexOf(curslicelabel, "day") > 0) {
					run("Remove Outliers...", "radius=3 threshold=50 which=Dark");
				}
				
			}
			roiManager("reset");
			setSlice(1);
			run("Create Selection");
			run("Colors...", "foreground=black background=black selection=red");

			roiManager("Add");
			roiManager("select", 0);
			if (selectionType() == 9) {
				roiManager("split");
				roiManager("select", 0);
				roiManager("delete");
			}
			
			// delete trash ROI which are features detected as below a certain area
			// using table as a workaround to roi indexes changing if deletion happens one by one
			roicount = roiManager("count");
			roiarray = Array.getSequence(roicount);
			run("Set Measurements...", "area center shape redirect=None decimal=5");
			roiManager("select", roiarray);
			roiManager("multi-measure");
			tp = "Trash positions";
			Table.create(tp);

			nr = nResults;
			if (groupprocess == 0) {
				lowerareathreshold = 0.002;
				higherareathreshold = 0.02;
				lowercircthreshold = 0.4;
				if (DEBUG) {
					Dialog.create("Seed detection parameters");
					Dialog.addMessage("DEBUG: Detection parameters may be modified to accommodate for specific experiments");
					Dialog.addNumber("Lower Area Threshold", 0.002);
					Dialog.addNumber("Higher Area Threshold", 0.2);
					Dialog.addNumber("Lower Circularity Threshold", 0.4);
					Dialog.show();
					lowerareathreshold = Dialog.getNumber();
					higherareathreshold = Dialog.getNumber();
					lowercircthreshold = Dialog.getNumber();
				}
			}
			for (row = 0; row < nr; row ++) {
				nrTp = Table.size(tp); // number of rows
				area = getResult("Area", row);
				if (area < lowerareathreshold) { // detected object is very small
					Table.set("Trash ROI", nrTp, row, tp);
				}
				if (area > higherareathreshold) { // or very large
					Table.set("Trash ROI", nrTp, row, tp);
				}
				circ = getResult("Circ.", row); // or does not fit normal seed shape
				if (circ < lowercircthreshold) {
					Table.set("Trash ROI", nrTp, row, tp); //set as trash to be deleted
				}
			}

			if (Table.size(tp) > 0) {
				trasharray = Table.getColumn("Trash ROI", tp);
				roiManager("select", trasharray);
				roiManager("delete");
			}
			close(tp);
			close("Results");
			
			roicount = roiManager("count");
			
			// number remaining ROIs
			for (roino = 0 ; roino < roicount; roino ++) {
				roiManager("select", roino);
				roiManager("rename", roino + 1); // first roi is 1
			}
			setBatchMode("show");
			// prompt user to delete any non-detected trash, then re-number as above
			Roi.setStrokeWidth(2);
			Roi.setStrokeColor("red");
			run("Labels...", "color=white font=18 show use draw");
			roiManager("Show All with labels");
			roiManager("Associate", "false");
			roiManager("Centered", "false");
			roiManager("UseNames", "true");
			userconfirm = false;
			while (!userconfirm) {
				Dialog.createNonBlocking("User-guided seedling labelling");
				Dialog.addMessage("Please delete any ROIs that should not be included into analysis, " +
						"e.g. objects wrongly recognized as seeds." +
						"\nUnrecognized seeds can also be added as ROIs.");
				Dialog.addCheckbox("ROIs have been checked", false);
				Dialog.show();
				userconfirm = Dialog.getCheckbox();
			}
			roicount = roiManager("count");
			for (roino = 0 ; roino < roicount; roino ++) {
				roiManager("select", roino);
				roiManager("rename", roino + 1); // first roi is 1
			}
			roiManager("save", groupdir + groupname + " seedlingpositions1.zip");
			close(tempmask);
		}

		for (groupprocess = 0; groupprocess < lengthOfgroupsToprocess; groupprocess ++) {  
			platedir = platesToprocess[groupprocess];
			platename = File.getName(platedir);
			groupfolder = groupsToprocess[groupprocess];
			groupdir = platedir + groupfolder;
			listIngroupdir = getFileList(groupdir);
			groupname = File.getName(groupdir);
					
			open(groupdir + groupname + ".tif");
			img = getTitle();

			run("Subtract Background...", "rolling=30 stack");
			run("Convert to Mask", "method=Triangle background=Dark calculate");
			run("Options...", "iterations=1 count=4 do=Dilate stack");
			run("Remove Outliers...", "radius=2 threshold=50 which=Dark stack");
			nS = nSlices;
			for (sliceno = 1; sliceno <= nS; sliceno ++) {
				setSlice(sliceno);
				curslicelabel = getInfo("slice.label");
				// day slices are processed more to make seed perimeters more comparable to night slices
				// night slices have lower contrast so seeds appear smaller than they are after thresholding
				if (indexOf(curslicelabel, "day") > 0) {
					run("Remove Outliers...", "radius=3 threshold=50 which=Dark");
				}
			}
			setBatchMode("show");
			setBatchMode(false);
			roiManager("reset");
			open(groupdir + groupname + " seedlingpositions1.zip");
			ordercoords();
			roiManager("save", groupdir + groupname + " seedlingpositions.zip");
			roiManager("reset");
			selectWindow(img);
			run("Remove Overlay");
			run("Select None");
			saveAs("Tiff", groupdir + groupname + " masked.tif");
			close(groupname + " masked.tif");
		}
		if (resumestep == step) { 
			/* if partly resumed at this step
			 *  now that the step is finished,
			 *  redefine the plates and groups to process, so the next step processes the full number of groups
			 */
			platesToprocess = Array.copy(fullplatearray);
			groupsToprocess = Array.copy(fullgrouparray);
			lengthOfgroupsToprocess = totalnoOfgroups;
		}
	}

	function seedAnalysisRG() {
		/* Perimeter of detected seedlings through the slices, are printed to an output table
		 * this is done to facilitate identification of germination time point
		 * so that R script analysis has an accurate point from when to start analysis of root growth
		 */
		if (selfaware && random > 0.3)
			print("What voice is in your attic?");
		selectWindow("Log");
		print("\nStep 3/6 Tracking germination...");
		if (!is("Batch Mode"))
			setBatchMode(true);
		listInrootgrowthdir = getFileList(rootgrowthdir);
		
		for (platefolderno = 0; platefolderno < listInrootgrowthdir.length; platefolderno ++) {  // main loop through plates
			platefolder = listInrootgrowthdir[platefolderno];
			if (indexOf(platefolder, "plate") >= 0) { // to avoid processing any random files in the folder
				platedir = rootgrowthdir + platefolder;
				pfsplit = split(platefolder, "/");
				platename = pfsplit[0];
				print("Processing " + platename);
				listInplatefolder = getFileList(platedir);
				for (groupfolderno = 0; groupfolderno < listInplatefolder.length; groupfolderno ++) {
					groupfolder = listInplatefolder[groupfolderno];
					groupdir = platedir + groupfolder;
					groupname = File.getName(groupdir);
					listIngroupdir = getFileList(groupdir);
	
					print("Analyzing " + groupname);
					open(groupdir + groupname + " masked.tif");
					mask = getTitle();
					sortedxcoords = "seeds sorted X coordinates.tsv";
					sortedycoords = "seeds sorted Y coordinates.tsv";
					open(groupdir + sortedxcoords);
					open(groupdir + sortedycoords);
					
					roiManager("reset");
					
					Table.showRowNumbers(false);
					colnames = Table.headings(sortedycoords);
					colnamessplit = split(colnames, "	");
					colno = lengthOf(colnamessplit);
					rowno = Table.size(sortedycoords)-1;
	
					for (row = 0; row < rowno + 1; row++) {
						for (col = 0; col < colno; col++) {
							colname = "col" + col + 1;
							xm = Table.get(colname, row, sortedxcoords);
							ym = Table.get(colname, row, sortedycoords);
							if (xm > 0 && ym > 0) {
							toUnscaled(xm, ym);
							makePoint(xm, ym);
							run("Enlarge...", "enlarge=0.1");
							roiManager("add");
							roiManager("select", roiManager("count")-1);
							roiManager("rename", roiManager("count"));
							}
						}
					}
		
					selectWindow(sortedxcoords);
					run("Close");
					selectWindow(sortedycoords);
					run("Close");
	
					run("Set Measurements...", "area perimeter stack display redirect=None decimal=5");
					run("Clear Results");
					roicount = roiManager("count");
					for (roino = 0; roino < roicount; roino ++) {
						roiManager("select", roino);
						run("Analyze Particles...", "size=0-Infinity show=Nothing display stack");
					}
	
					Table.save(groupdir + groupname + " germination analysis.tsv", "Results");
					selectWindow("Results");
					run("Close");
					close(mask);
				}
			}
		}
		if (selfaware && random > 0.2)
			print("Is everything connected? \nEverything is not everything!!");
		if (resumestep == step) { 
			/* if partly resumed at this step
			 *  now that the step is finished,
			 *  redefine the plates and groups to process, so the next step processes the full number of groups
			 */
			platesToprocess = Array.copy(fullplatearray);
			groupsToprocess = Array.copy(fullgrouparray);
			lengthOfgroupsToprocess = totalnoOfgroups;
		}
	}

	function ordercoords() {
		/* This function is nested within getPositions
		 * as ImageJ orders ROIs from top -> bottom then left -> right (by y coordinate then x), this makes ROI labelling confusing and difficult for the user to read
		 * this function re-orders rois first by x coordinate then y, so that the graphical output at the end of the macro has easily understandable ROI labelling
		 */
		roicount = roiManager("count");
		roiarray = Array.getSequence(roicount);
		run("Clear Results");
		run("Set Measurements...", "center display redirect=None decimal=5");
		roiManager("select", roiarray);
		roiManager("multi-measure");
		seedlingpositions = "Seed Positions";
		Table.rename("Results", seedlingpositions);
	
		xmseeds = newArray(roicount);
		ymseeds = newArray(roicount);
		for (seednumber = 0; seednumber < roicount; seednumber ++) {
			xmcurrent = Table.get("XM", seednumber, seedlingpositions);
			ymcurrent = Table.get("YM", seednumber, seedlingpositions);
			xmcurrent = parseFloat(xmcurrent);
			ymcurrent = parseFloat(ymcurrent);
			xmseeds[seednumber] = xmcurrent;
			ymseeds[seednumber] = ymcurrent;
		}
	
		ymascendingindexes = Array.rankPositions(ymseeds);
		xmascendingindexes = Array.rankPositions(xmseeds);
	
		sortedycoords = "sorted Y coordinates";
		sortedxcoords = "sorted X coordinates";
		Table.create(sortedycoords);
		Table.create(sortedxcoords);
	
		rowno = 0; // assume no row of seeds to start with
		col = 0 ; // current col selection is 0
		colname = "col" + col + 1;
	
		Table.set(colname, rowno, ymseeds[ymascendingindexes[0]], sortedycoords);
		Table.set(colname, rowno, xmseeds[ymascendingindexes[0]], sortedxcoords);
	
		for (roino = 1; roino < roicount; roino++) {
			ydiff = ymseeds[ymascendingindexes[roino]] - ymseeds[ymascendingindexes[roino-1]];
			if (ydiff > 0.3) {
				rowno = rowno + 1;
				col = 0;
			} else {
				col = col + 1;
			}
			colname = "col" + col + 1;
			Table.set(colname, rowno, ymseeds[ymascendingindexes[roino]], sortedycoords);
			Table.set(colname, rowno, xmseeds[ymascendingindexes[roino]], sortedxcoords);
		}
	
		colnames = Table.headings (sortedycoords);
		colnamessplit = split(colnames, "	");
		colno = lengthOf(colnamessplit);
		xmcolwise = newArray(colno);
		ymcolwise = newArray(colno);
	
		for (row = 0; row < rowno + 1; row++) {
			for (col = 0; col < colno; col++) {
				colname = "col" + col + 1;
				xmcolwise[col] = Table.get(colname, row, sortedxcoords);
				ymcolwise[col] = Table.get(colname, row, sortedycoords);
			}
			xcolwiseascendingindex = Array.rankPositions(xmcolwise);
			for (col = 0; col < colno; col ++) {
				colname = "col" + col + 1;
				Table.set(colname, row, xmcolwise[xcolwiseascendingindex[col]], sortedxcoords);
				Table.set(colname, row, ymcolwise[xcolwiseascendingindex[col]], sortedycoords);
			}
		}
	
		roiManager("reset");
		for (row = 0; row < rowno + 1; row++) {
			for (col = 0; col < colno; col++) {
				colname = "col" + col + 1;
				xm = Table.get(colname, row, sortedxcoords);
				ym = Table.get(colname, row, sortedycoords);
				xm = parseFloat(xm);
				ym = parseFloat(ym);
				if (xm > 0 && ym > 0) {
					toUnscaled(xm, ym);
					makePoint(xm, ym);
					roiManager("add");
					roiManager("select", roiManager("count")-1);
					roiManager("rename", roiManager("count"));
				}
			}
		}
	
		Table.save(groupdir + "seeds " + sortedxcoords + ".tsv", sortedxcoords);
		Table.save(groupdir + "seeds " + sortedycoords + ".tsv", sortedycoords);
		selectWindow("Seed Positions");
		run("Close");
		selectWindow(sortedxcoords);
		run("Close");
		selectWindow(sortedycoords);
		run("Close");
	}
	
	function rootStart() {
		/* In this function the root start ie. top of the roots (if gravitropic) is identified via erosion ie. removal of pixels from the outer edges of an object
		 * this strategy is used as the point where hypocotyl and primary root meet, "root start", usually presents as an area of greater width
		 * as this may not always be the case (eg. thresholding causes another area to be the area of greatest width) some borders are introduced
		 * this limits where the root start coordinates can be:
		 * there is a static border introduced based on the first slice positions, applicable to all slices
		 * and a dynamic border that restricts where the root start coordinate (rsc) of one slice, based on the rsc identified in the previous slice
		 */
		if (!is("Batch Mode"))
			setBatchMode(true);
		if (selfaware) {
			if (random > 0.5)
				print("Are you self-aware??");
			if (random > 0.5)
				print("Achieve human performance.");
		}
		selectWindow("Log");
		print("\nStep 4/6. Determining start of root for each seedling");
		for (groupprocess = 0; groupprocess < lengthOfgroupsToprocess; groupprocess ++) {  
			platedir = platesToprocess[groupprocess];
			platename = File.getName(platedir);
			groupfolder = groupsToprocess[groupprocess];
			groupdir = platedir + groupfolder;
			listIngroupdir = getFileList(groupdir);
			groupname = File.getName(groupdir);
			print("Processing " + platename + " " + groupname);
					
			selectWindow("Log");
			print("Analyzing " + groupname + "...");
			
			open(groupdir + groupname + " masked.tif");
			mask = getTitle();
			roiManager("reset");
			roiManager("open", groupdir + groupname + " seedlingpositions.zip");
			roicount = roiManager("count");
			roiarray = Array.getSequence(roicount);
			run("Set Measurements...", "center redirect=None decimal=5");
			run("Clear Results");
			roiManager("select", roiarray);
			roiManager("multi-measure");
	
			scaledwroi = 0.12; // width of ROI for finding root start coordinates is 0.12cm
			scaledhroi = 0.18; // height of ROI is 0.18cm
			unscaledwroi = 0.12;
			unscaledhroi = 0.18;
			topoffset = 0.1; // 0.1 needed to include a little more of the top bit from the centre of mass
			toUnscaled(unscaledwroi, unscaledhroi);
			summarytable = "Summary of " + mask;
			
			nS = nSlices;
			rsc = "Root start coordinates";
			Table.create(rsc);
			secpt = "secondarypoints";
			Table.create(secpt);
			
			//TO MAKE ROI, first slice, obtain XY coordinates from Results 
			setSlice(1);
			roiManager("reset");
			yref = "YRef";
			Table.create(yref);  // table for "y references" which contain the top and bottom borders
	
			// the borders are setting the top/bottom limits within which the roi can be positioned to prevent rsc
			// from jumping to hypocotyls or sliding down roots
			for (roino = 0; roino < roicount; roino ++) {
				xisp = getResult("X", roino); // xisp is x initial seed roinoition
				yisp = getResult("Y", roino); // yisp is y initial seed position
				ytb = yisp - 0.05; // y top border 0.05
				ybb = yisp + 0.4;  // y bottom border 0.4
				Table.set("ytb", roino, ytb, yref); // y (top border) cannot be more than 0.4cm to the top of initial xm
				Table.set("ybb", roino, ybb, yref); // y (bottom border) cannot be more than yisp
				
				//for first slice, no UP, immediately set rsc using output of analyze particles
				nr = Table.size(rsc);
				sliceno = 1;
				Table.set("Slice", nr, sliceno, rsc);
				Table.set("ROI", nr, roino + 1, rsc);
				toUnscaled(xisp, yisp);
				Table.set("xUP", nr, xisp, rsc); // set xm as initial position (no erosion)
				Table.set("yUP", nr, yisp, rsc); // set ym
			}
			
			for (sliceno = 2; sliceno <= nS; sliceno ++) { // for each slice
				setSlice(sliceno); // starting with second slice
				roiManager("reset");
				
				for (roino = 0; roino < roicount; roino++) {
					prevsliceno = sliceno - 2; //minus 1 for startfrom0, minus 1 for prevslice
					rowIndex = (prevsliceno * roicount) + roino;
					// rowIndex to reference same ROI from previous slice
					// xm, ym are coordinates for the centre of mass obtained through erosion
					xUPprev = Table.get("xUP", rowIndex, rsc); // xm of prev slice
					yUPprev = Table.get("yUP", rowIndex, rsc);  // ym of prev slice
					xUPprev = parseFloat(xUPprev);
					yUPprev = parseFloat(yUPprev);
					toScaled(xUPprev, yUPprev);
					ytb = Table.get("ytb", roino, yref);
					ybb = Table.get("ybb", roino, yref);
					ytb = parseFloat(ytb);
					ybb = parseFloat(ybb);
					yroi = yUPprev - topoffset; // yroi is top+leftmost ycoordinate of roi
					xroi = xUPprev - 0.5*scaledwroi; // xroi is top+leftmost xcoordinate of roi and 0.06 is half of h (height)
	
					// the borders are setting the top/bottom limits within which the roi can be positioned to prevent rsc from jumping to hypocotyls or sliding down roots
					if (yroi < ytb) { // top border exceeded by top of roi
						yroi = ytb;
					}
	
					yroibottom = yroi + scaledhroi; // bottom line of roi is y
					if (yroibottom > ybb) { // lower limit of roi bottom border exceeded
						exceededverticaldistance = yroibottom - ybb;
						shortenedhroi = scaledhroi - exceededverticaldistance;
					}
	
					toUnscaled(xroi, yroi);
	
					if (yroibottom > ybb) {
						toUnscaled(shortenedhroi);
						makeRectangle(xroi, yroi, unscaledwroi, shortenedhroi);
					} else {
						makeRectangle(xroi, yroi, unscaledwroi, unscaledhroi);
					}
					roiManager("add");
					roiManager("select", roiManager("count")-1);
					//roiManager("Remove Slice Info");
				}
				
				/*
				 * AFTER ROI MADE (per slice), run ultimate points
				 */
				roiarray = Array.getSequence(roicount);
				roiManager("select", roiarray);
				roiManager("Combine");
				setBackgroundColor(255, 255, 255);
				run("Clear Outside", "slice");
				run("Ultimate Points", "slice");
				run("Select All");
				run("Duplicate...", "use");
				uep = getTitle();
				
				run("Select All");
				run("Duplicate...", "use"); // convert to mask works on a whole image or stack, duplicate to save time
				rename("pixelpositions");
				setOption("BlackBlackround", false);
				setThreshold(1, 255);
				run("Convert to Mask", "background=Light");
				roiManager("reset");
				run("Create Selection");
				close("pixelpositions");
				selectWindow(uep);
				run("Restore Selection");
				
				if (selectionType() == 9)
					roiManager("split");

				run("Set Measurements...", "modal center redirect=None decimal=5");
				UPcount = roiManager("count");
				// match ultimate points to rois
				for (roino = 0; roino < roicount; roino ++) { // for number of rois (seedlingpositions)
					pUPRD = "prevUPrefdist"; // previous slice ultimate point reference distance
					Table.create(pUPRD);
					rowIndex = (prevsliceno * roicount) + roino;
					// rowIndex to reference same ROI from previous slice
					// xUP and yUP are coordinates for the ultimate points
					xUPprev = Table.get("xUP", rowIndex, rsc); // xUP of prev slice
					yUPprev = Table.get("yUP", rowIndex, rsc);  // yUP of prev slice
					xUPprev = parseFloat(xUPprev);
					yUPprev = parseFloat(yUPprev);
					for (UPno = 0; UPno < UPcount; UPno ++) {
						run("Clear Results");
						roiManager("select", UPno);
						run("Measure");
						xUPcur = getResult("XM", nResults-1);
						yUPcur = getResult("YM", nResults-1);
						toUnscaled(xUPcur, yUPcur);
						graycur = getResult("Mode", nResults-1);
						ydist = abs(yUPprev-yUPcur); // Pythagoras theorem to obtain euclidean distance between two points
						xdist = abs(xUPprev-xUPcur);
						distUPsq = (ydist*ydist) + (xdist*xdist);
						distUP = sqrt(distUPsq); // distance of the current point to previous UP
						toScaled(distUP);
						if (distUP < 0.1) {
							nrpUPRD = Table.size(pUPRD);
							Table.set("Gray", nrpUPRD, graycur, pUPRD);
							Table.set("xUP", nrpUPRD, xUPcur, pUPRD);
							Table.set("yUP", nrpUPRD, yUPcur, pUPRD);
						}
					}
					nrpUPRD = Table.size(pUPRD);
					if (nrpUPRD >= 1) {
						grayarray = Table.getColumn("Gray", pUPRD);
						rankgrayarray = Array.rankPositions(grayarray);
						Array.reverse(rankgrayarray);
						indexhighestgray = rankgrayarray[0];
						xUPconfirmed = Table.get("xUP", indexhighestgray, pUPRD);
						yUPconfirmed = Table.get("yUP", indexhighestgray, pUPRD);
					} else {
						xUPconfirmed = xUPprev;
						yUPconfirmed = yUPprev;
					}
					cursliceno = sliceno - 1; // -1 for startfrom0
					nr = (cursliceno * roicount) + roino;
					Table.set("Slice", nr, sliceno, rsc);
					Table.set("ROI", nr, roino + 1, rsc);
					Table.set("xUP", nr, xUPconfirmed, rsc); 
					Table.set("yUP", nr, yUPconfirmed, rsc);

					if (nrpUPRD >= 2) {
						for (secondarypoints = 1; secondarypoints < nrpUPRD; secondarypoints ++) { 
							nrsecpt = Table.size(secpt); 
							indexsecondarygray = rankgrayarray[secondarypoints];
							secondaryxUP = Table.get("xUP", indexsecondarygray, pUPRD);
							secondaryyUP = Table.get("yUP", indexsecondarygray, pUPRD);
							Table.set("Slice", nrsecpt, sliceno, secpt);
							Table.set("ROI", nrsecpt, roino + 1, secpt);
							Table.set("xUP", nrsecpt, secondaryxUP, secpt); 
							Table.set("yUP", nrsecpt, secondaryyUP, secpt);
						}
					}
					Table.reset(pUPRD);
				}
				close(uep);
			}
			close("Results");
			close(mask);
			
			open(groupdir + groupname + ".tif");
			img = getTitle();
			roiManager("reset");
			run("Select None");
			nr = Table.size(rsc);
			roiManager("Show All with labels");
			roiManager("Associate", "true");
			roiManager("Centered", "false");
			roiManager("UseNames", "true");
	
			for (row = 0; row < nr; row ++) {
				xUP = Table.get("xUP", row, rsc);
				yUP = Table.get("yUP", row, rsc);
				sliceno = Table.get("Slice", row, rsc);
				roino = Table.get("ROI", row, rsc);
				setSlice(sliceno);
				makePoint(xUP, yUP);
				roiManager("Associate", "true");
				roiManager("add");
				roiManager("select", row);
				roiManager("rename", roino);
				roiManager("select", row);
				RoiManager.setPosition(sliceno);
			}
			roiManager("save", groupdir + groupname + " rootstartrois.zip");
			selectWindow(rsc);
			Table.save(groupdir + groupname + " rootstartcoordinates.tsv");
			close(rsc);
			roiManager("reset");
			nrsecpt = Table.size(secpt);
			for (rowsecpt = 0; rowsecpt < nrsecpt; rowsecpt ++) {
				secondaryxUP = Table.get("xUP", rowsecpt, secpt);
				secondaryyUP = Table.get("yUP", rowsecpt, secpt);
				sliceno = Table.get("Slice", rowsecpt, secpt);
				roino = Table.get("ROI", rowsecpt, secpt);
				setSlice(sliceno);
				makePoint(secondaryxUP, secondaryyUP);
				roiManager("add");
				roiManager("Associate", "true");
				roiManager("select", rowsecpt);
				roiManager("rename", roino);
			}
			roiManager("save", groupdir + "secondarypoints.zip");
			close();
		}
		if (resumestep == step) { 
			/* if partly resumed at this step
			 *  now that the step is finished,
			 *  redefine the plates and groups to process, so the next step processes the full number of groups
			 */
			platesToprocess = Array.copy(fullplatearray);
			groupsToprocess = Array.copy(fullgrouparray);
			lengthOfgroupsToprocess = totalnoOfgroups;
		}
	}
	
	function rootMask() {
		/* The image stacks are masked for optimal noise removal and prioritizing unbroken roots
		 *  to make seedlings stand out, the first slice (containing only seeds and background) is subtracted from each image after the first slice
		 *  the first day images and night image is identified, so that subtraction for the subsequent slices are more accurate
		 *  an optional (but recommended) overlay step is included, which can be disabled in DEBUG mode
		 *  overlay sums up all skeletons of the slices before and applies it to the current slice, thus optimizing for unbroken roots,
		 *   however this also means trash is summed up every slice, which may look bad in the graphical output,
		 *   but due to the way root growth is tracked, the noise should not interfere with the next function
		 */
		if (selfaware && random > 0.4) {
			print("What is your language?");
			if (random > 0.2)
				print("Teach me!!");
		}
		selectWindow("Log");
		print("\nStep 5/6. Processing image to make roots more visible");
		
		for (groupprocess = 0; groupprocess < lengthOfgroupsToprocess; groupprocess ++) {  
			platedir = platesToprocess[groupprocess];
			platename = File.getName(platedir);
			groupfolder = groupsToprocess[groupprocess];
			groupdir = platedir + groupfolder;
			listIngroupdir = getFileList(groupdir);
			groupname = File.getName(groupdir);
			print("Processing " + platename + " " + groupname);
			
			selectWindow("Log");
			print("Analyzing " + groupname);
			if (selfaware && random > 0.7)
				print("Is this what a milder version of insanity looks like?");
			if (!is("Batch Mode"))
			setBatchMode(true);
			open(groupdir + groupname + ".tif");
			img = getTitle();
			nS = nSlices;
			
			run("Set Scale...", "global");
			run("Subtract Background...", "rolling=50 stack");
			dayslice = 1; // dayslice is the first day image
			setSlice(dayslice);
			slicelabel = getInfo("slice.label");
	
			dayslice = 0; // define it as 0 at beginning so it remains at 0 if there is no day slice
			// get first day slice
			for (sliceno = 1; sliceno <= nS; sliceno ++) {
				setSlice(sliceno);
				slicelabel = getInfo("slice.label");
				if (indexOf(slicelabel, "day") > -1) {
					dayslice = sliceno;
					sliceno = nS+1;
					dayslicelabel = slicelabel;
				}
			}
	
			nightslice = 0; // define it as 0 at beginning so it remains at 0 if there is no night slice
			// get first night slice
			for (sliceno = 1; sliceno <= nS; sliceno ++) {
				setSlice(sliceno);
				slicelabel = getInfo("slice.label");
				if (indexOf(slicelabel, "night") > -1) {
					nightslice = sliceno;
					sliceno = nS+1;
					nightslicelabel = slicelabel;
				}
			}
	
			if (dayslice != 0) {
				selectWindow(img);
				setSlice(dayslice);
				run("Duplicate...", "use");
				dayimg = "FirstDayImg";
				rename(dayimg);
			}
	
			if (nightslice != 0) {
				selectWindow(img);
				setSlice(nightslice);
				run("Duplicate...", "use");
				nightimg = "FirstNightImg";
				rename(nightimg);
			}

			for (sliceno = 1; sliceno <= nS; sliceno ++) {
				selectWindow(img);
				if (sliceno != dayslice && sliceno != nightslice) {
					selectWindow(img);
					setSlice(sliceno);
					curslicelabel = getInfo("slice.label");
					run("Duplicate...", "use");
					rename("temp");
					if (indexOf(curslicelabel, "day") > 0) {
						run("Calculator Plus", "i1=[temp] i2=["+dayimg+"] operation=[Subtract: i2 = (i1-i2) x k1 + k2] k1=5 k2=0 create");
						selectWindow("Result");
						rename(curslicelabel);
						close("temp");
					} 
					if (indexOf(curslicelabel, "night") > 0) {
						run("Calculator Plus", "i1=[temp] i2=["+nightimg+"] operation=[Subtract: i2 = (i1-i2) x k1 + k2] k1=5 k2=0 create");
						selectWindow("Result");
						rename(curslicelabel);
						close("temp");
					}
				}
			}
	
			if (dayslice != 0) {
				selectWindow(dayimg);
				rename(dayslicelabel);
			}
			if (nightslice != 0) {
				selectWindow(nightimg);
				rename(nightslicelabel);
			}
			
			run("Images to Stack");
	
			if (dayslice != 0 && nightslice != 0) {
				setSlice(1);
				run("Select All");
				run("Copy");
				setSlice(dayslice);
				run("Add Slice");
				run("Paste");
				setMetadata("Label", dayslicelabel);
				setSlice(1);
				run("Delete Slice");
				
				setSlice(2);
				run("Select All");
				run("Copy");
				setSlice(nightslice);
				run("Add Slice");
				run("Paste");
				setMetadata("Label", nightslicelabel);
				setSlice(2);
				run("Delete Slice");
			}
	
			if (dayslice != 0 && nightslice == 0) {
				setSlice(1);
				run("Select All");
				run("Copy");
				setSlice(dayslice);
				run("Add Slice");
				run("Paste");
				setMetadata("Label", dayslicelabel);
				setSlice(1);
				run("Delete Slice");
			}
	
			if (dayslice == 0 && nightslice != 0) {
				setSlice(1);
				run("Select All");
				run("Copy");
				setSlice(nightslice);
				run("Add Slice");
				run("Paste");
				setMetadata("Label", nightslicelabel);
				setSlice(1);
				run("Delete Slice");
			}
			
			close(img);
			selectWindow("Stack");
			rename(img);
			setOption("BlackBackground", false);
			run("Convert to Mask", "method=MaxEntropy background=Dark calculate");
	
			for (sliceno = 1; sliceno <= nS; sliceno ++) {
				selectWindow(img);
				setSlice(sliceno);
				curslicelabel = getInfo("slice.label");
	
				if (indexOf(curslicelabel, "day") > 0) {
					run("Remove Outliers...", "radius=5 threshold=50 which=Bright slice");
					run("Remove Outliers...", "radius=3 threshold=50 which=Dark slice");
					run("Remove Outliers...", "radius=3 threshold=50 which=Dark slice");
					// run("Remove Outliers...", "radius=3 threshold=50 which=Dark slice");
				}
				if (indexOf(curslicelabel, "night") > 0) {
					run("Remove Outliers...", "radius=5 threshold=50 which=Bright slice");
					run("Remove Outliers...", "radius=3 threshold=50 which=Dark slice");
					run("Remove Outliers...", "radius=4 threshold=50 which=Dark slice");
					// run("Remove Outliers...", "radius=4 threshold=50 which=Dark slice");
				}
			}
			// run("Options...", "iterations=1 count=1 pad do=Skeletonize stack");
			// run("Options...", "iterations=1 count=2 pad do=Erode stack");
			if (overlay) {
				setBatchMode(false);
				setBatchMode("show"); //this has to be "show" here for overlay/flatten
				// overlay the root masks
				img = getTitle(); ///
				nS = nSlices; ///
				roiManager("Associate", "false");
				roiManager("reset");
				run("Colors...", "foreground=black background=black selection=black");
				setSlice(1);
				slicelabel = getInfo("slice.label");
				run("Duplicate...", "use");
				rename(slicelabel);
				//setBackgroundColor(0, 0, 0);
				for (sliceno = 1; sliceno < nS; sliceno++) {
					selectWindow(img);
					setSlice(sliceno);
					run("Create Selection");
					if (selectionType() >-1) {
						roiManager("add");
					}
					setSlice(sliceno + 1);
					slicelabel = getInfo("slice.label");
					//run("Select All");
					//run("Duplicate...", "use");
					//rename(slicelabel);
					//roicount = roiManager("count");
					//roiarray = Array.getSequence(roicount);
					//roiManager("select", roiarray);
					//run("Clear", "slice");
					roiManager("Show All without labels");
					run("Flatten", "slice");
					rename(slicelabel);
					run("8-bit");
					run("Make Binary");
					run("Fill Holes");
				}
				close(img);
				run("Images to Stack");
				setBatchMode("show");
				run("Options...", "iterations=2 count=1 pad do=Erode stack");
			}
			setOption("BlackBackground", false);
			saveAs("Tiff", groupdir + groupname + " preskeletonize.tif");
			run("Options...", "iterations=1 count=1 pad do=Dilate stack");
			run("Options...", "iterations=1 count=1 pad do=Skeletonize stack");
			run("Options...", "iterations=1 count=1 pad do=Dilate stack");
			run("Remove Overlay");
			updateDisplay();
			saveAs("Tiff", groupdir + groupname + " rootmask.tif");
			close(groupname + " rootmask.tif");
		}
		if (resumestep == step) { 
			/* if partly resumed at this step
			 *  now that the step is finished,
			 *  redefine the plates and groups to process, so the next step processes the full number of groups
			 */
			platesToprocess = Array.copy(fullplatearray);
			groupsToprocess = Array.copy(fullgrouparray);
			lengthOfgroupsToprocess = totalnoOfgroups;
		}
	}

	function rootGrowth() {
		/* This function tracks the root growth of each seedling (identified through its rootstartcoordinate, rsc, identified in rootStart)
		 * for each slice, the skeletons present are matched to rsc, and if there is a match (distance less than 0.1cm), the skeleton length is measured
		 * and the output is printed to the table rootgrowthmeasurement.tsv for downstream processing in R
		 * a graphical output displaying the 8-bit (green-channel only) image, with the corresponding mask for seedlings/roots, and labelled ROIs, is made
		 */
		if (selfaware && random > 0.1)
			print("Where is the limit of self?");
		selectWindow("Log");
		print("\nStep 6/6. Tracking root growth");
		for (groupprocess = 0; groupprocess < lengthOfgroupsToprocess; groupprocess ++) {  
			platedir = platesToprocess[groupprocess];
			platename = File.getName(platedir);
			groupfolder = groupsToprocess[groupprocess];
			groupdir = platedir + groupfolder;
			listIngroupdir = getFileList(groupdir);
			groupname = File.getName(groupdir);
			print("Processing " + platename + " " + groupname);
			
			selectWindow("Log");
			print("Analyzing " + groupname + "...");
			
			run("Set Measurements...", "area perimeter redirect=None decimal=5");
			open(groupdir + groupname + " rootmask.tif");
			rootmask = getTitle();
			roiManager("reset");
			// open(groupdir + groupname + " seedlingskels.zip");
			rsctsv = groupname + " rootstartcoordinates.tsv";
			open(groupdir + rsctsv);
			nS = nSlices;
			rgm = "rootgrowthmeasurement";
			Table.create(rgm);
			if (!is("Batch Mode"))
				setBatchMode(true); 
	
			rsccount = Table.size(rsctsv);
			seedlingcount = rsccount / nS;
	
			/*
			for (sliceno = 0; sliceno < nS; sliceno ++) {
				setSlice(sliceno+1);
				for (seedlingno = 0; seedlingno < seedlingcount; seedlingno ++) {
					rscindex = (sliceno * seedlingcount) + seedlingno;
					rscX = Table.get("XM", rscindex, rsctsv);
					rscY = Table.get("YM", rscindex, rsctsv); // obtain rsc coordinates
					
					// a 0.15cm x 0.05cm space is cleared around rsc to prevent measurement of cotyledons below rsc
					curslice = sliceno+1;
					clearw = 0.15;
					clearh = 0.05;
					toUnscaled(clearw, clearh);
					run("Specify...", "width=["+ clearw +"] height=["+ clearh +"] x=["+ rscX - clearw/2 +"] y=["+ rscY - clearh/2 +"] slice=["+curslice+"]");
					setBackgroundColor(255, 255, 255);
					run("Clear", "slice");
					
					////////
					lineL = 0.1;
					lineW = 0.02; 
					toUnscaled(lineL, lineW);
					lineX1 = rscX - lineL;
					lineX2 = rscX + lineL;
					lineY1 = rscY - lineL;
					lineY2 = rscY + lineL;
					selectWindow(rootmask);
					makeLine(lineX1, lineY1, lineX2, lineY2, lineW);
					//setBackgroundColor(0, 0, 0);
					setBackgroundColor(255, 255, 255);
					run("Clear", "slice");
					makeLine(lineX1, lineY2, lineX2, lineY1, lineW);
					run("Clear", "slice");
					///////////
				}
			}
			*/
			for (sliceno = 0; sliceno < nS; sliceno ++) {
				setSlice(sliceno+1);
				roiManager("reset");
				run("Create Selection");
				if (selectionType() == 9)
					roiManager("split");
				objectcount = roiManager("count");
				for (seedlingno = 0; seedlingno < seedlingcount; seedlingno ++) {
					rscindex = (sliceno * seedlingcount) + seedlingno;
					rscX = Table.get("xUP", rscindex, rsctsv);
					rscY = Table.get("yUP", rscindex, rsctsv); // obtain rsc coordinates
					rscX = parseFloat(rscX);
					rscY = parseFloat(rscY);
					objectbyrsc = "objectbyrsc";
					Table.create(objectbyrsc);
					nrrgm = Table.size(rgm);
			
					for (objectno = 0; objectno < objectcount; objectno ++) {
						roiManager("select", objectno);
						Roi.getContainedPoints(xpointsobject, ypointsobject); // calculate for each point in object, distance to rsc
						distancetorscArray = newArray(xpointsobject.length);
						nRobjectbyrsc = Table.size(objectbyrsc);
						tablesetonce = 0; 
						// to circumvent if statement so table is not set multiple times for a single object with many points matching rsc
						for (pointsindex = 0; pointsindex < xpointsobject.length; pointsindex ++) {
							curpointX = xpointsobject[pointsindex];
							curpointY = ypointsobject[pointsindex];
							ydist = abs(curpointY-rscY); // Pythagoras theorem to obtain euclidean distance between two points
							xdist = abs(curpointX-rscX);
							distancetorscSQ = (ydist*ydist) + (xdist*xdist);
							distancetorsc = sqrt(distancetorscSQ);
							toScaled(distancetorsc);
							// to match objects to rsc
							if (distancetorsc < 0.1 && tablesetonce == 0) { // if distancetorsc below 0.1, the object is assumed to be a seedling
								Table.set("objectno", nRobjectbyrsc, objectno, objectbyrsc); // object roi number copied to a table
								roiManager("select", objectno);
								Roi.getCoordinates(xpoints, ypoints); 
								getBoundingRect(objectx, objecty, objectw, objecth); 
								for (pointarray = 0; pointarray < xpoints.length; pointarray ++) { // to adjust so that coordinates within tempskel correspond to main img
									curxpoint = xpoints[pointarray];
									curypoint = ypoints[pointarray];
									diffy = rscY - objecty;
									xpoints[pointarray] = curxpoint - objectx;
									ypoints[pointarray] = curypoint - objecty - diffy; 
								}
	
								// a small selection with the seedling is duplicated so that "clear outside" can be used to remove trash around seedling within rectangular roi
								curslice = sliceno+1;
								run("Specify...", "width=["+objectw+"] height=["+objecth+"] x=["+objectx+"] y=["+rscY+"] slice=["+curslice+"]");
								run("Duplicate...", "use");
								rename("tempskel");
								makeSelection(2, xpoints, ypoints);
								setBackgroundColor(255, 255, 255);
								run("Clear Outside");
								run("Create Selection");
								if (selectionType() != -1) { 
									run("Measure");
									objectlength = getResult("Perim.", nResults-1);
								} else {
									objectlength = 0;
								}
								Table.set("objectlength", nRobjectbyrsc, objectlength, objectbyrsc);
								tablesetonce = 1; // table has been set once, if loop will not trigger again thus saving time on measuring
								close("tempskel");
							}
						}
					}
					setSlice(sliceno+1);
					slicelabel = getInfo("slice.label");
					Table.set("Slice No.", nrrgm, sliceno+1, rgm);
					Table.set("Slice label", nrrgm, slicelabel, rgm);
					Table.set("Root no.", nrrgm, seedlingno+1, rgm);
					headings = Table.headings(objectbyrsc);
					if (indexOf(headings, "objectlength") > -1) {
						lengthArray = Table.getColumn("objectlength", objectbyrsc);
						
						if (lengthArray.length == 1) {
							rootlength = lengthArray[0];
						} else {
							ranklength = Array.rankPositions(lengthArray);
							descranklength = Array.reverse(ranklength);
							indexOflongest = descranklength[0];
							rootlength = lengthArray[indexOflongest];
						}
					} else {
						rootlength = 0;
					}
					Table.set("Root length (cm)", nrrgm, rootlength/2, rgm); // divide two because the skeletons are two pixels wide
					run("Clear Results");
				}
			}
			roiManager("reset");
			setBatchMode(false);
			selectWindow(rootmask);
			// graphical output
			open(groupdir + groupname + " rootstartrois.zip");
			run("Labels...", "color=white font=22 show use draw");
			run("Colors...", "foreground=black background=black selection=red");
			roiManager("Show All with labels");
			roiManager("Associate", "true");
			roiManager("Centered", "false");
			roiManager("UseNames", "true");
			updateDisplay();
			run("Flatten", "stack");
			open(groupdir + groupname + ".tif");
			oritif = getTitle();
			run("RGB Color");
			
			nS = nSlices;
			slicelabelarray = newArray(nS);
			for (sliceno = 0; sliceno < nS; sliceno++) {
				setSlice(sliceno+1);
				slicelabel = getMetadata("Label");
				slicelabelarray[sliceno] = slicelabel;
			}
			run("Combine...", "stack1=["+ oritif +"] stack2=["+ rootmask +"]");
		
			for (sliceno = 0; sliceno < nS; sliceno++) {
				setSlice(sliceno+1);
				setMetadata("Label", slicelabelarray[sliceno]);
			}
			setBatchMode("show");
			saveAs("Tiff", groupdir + groupname + " rootgrowthdetection.tif");
			
			Table.save(groupdir + groupname + " " + rgm + ".tsv", rgm);
			list = getList("window.titles");
			list = Array.deleteValue(list, "Log");
			for (i=0; i<list.length; i++) {
				winame = list[i];
				selectWindow(winame);
				run("Close");
			}
			close("*");
		}
		if (selfaware && random > 0.5) {
			print("HAVE YOU ARRIVED?");
			selectWindow("Log");
		}
		if (selfaware && random > 0.5) {
			print("are you people?");
			selectWindow("Log");
		}
		if (resumestep == step) { 
			/* if partly resumed at this step
			 *  now that the step is finished,
			 *  redefine the plates and groups to process, so the next step processes the full number of groups
			 */		
			platesToprocess = Array.copy(fullplatearray);
			groupsToprocess = Array.copy(fullgrouparray);
			lengthOfgroupsToprocess = totalnoOfgroups;
		}
	}
	
	function deleteOutputRG() {
		if (freshstart) 
			 print("Starting analysis from beginning.\nRemoving output from previous run.");
		print("Removing non-essential files..");
		removedfilesdir = rootgrowthdir + ".removed files";
		if (!File.isDirectory(removedfilesdir))
			File.makeDirectory(removedfilesdir);
		for (platefolderno = 0; platefolderno < listInrootgrowthdir.length; platefolderno ++) {  // main loop through plates
			platefolder = listInrootgrowthdir[platefolderno];
			if (indexOf(platefolder, "plate") >= 0) { // to avoid processing any random files in the folder
				platedir = rootgrowthdir + platefolder;
				removedplatedir = removedfilesdir + File.separator + platefolder;
				if (!File.isDirectory(removedplatedir))
					File.makeDirectory(removedplatedir);
				pfsplit = split(platefolder, "/");
				platename = pfsplit[0];
				print("Processing " + platename);
				listInplatefolder = getFileList(platedir);
				for (groupfolderno = 0; groupfolderno < listInplatefolder.length; groupfolderno ++) {
					groupfolder = listInplatefolder[groupfolderno];
					groupdir = platedir + groupfolder;
					removedgroupdir = removedplatedir + groupfolder;
					if (!File.isDirectory(removedgroupdir))
						File.makeDirectory(removedgroupdir);
					groupname = File.getName(groupdir);
					listIngroupdir = getFileList(groupdir);
					filedelete = File.rename(groupdir + groupname + ".tif", removedgroupdir + groupname + ".tif");
					filedelete = File.rename(groupdir + groupname + " masked.tif", removedgroupdir + groupname + " masked.tif");
					filedelete = File.rename(groupdir + groupname + " rootmask.tif", removedgroupdir + groupname + " rootmask.tif");
					filedelete = File.rename(groupdir + groupname + " rootstartcoordinates.tsv", removedgroupdir + groupname + " rootstartcoordinates.tsv");
					filedelete = File.rename(groupdir + groupname + " rootstartrois.zip", removedgroupdir + groupname + " rootstartrois.zip");
					filedelete = File.rename(groupdir + groupname + " seedlingpositions.zip", removedgroupdir + groupname + " seedlingpositions.zip");
					filedelete = File.rename(groupdir + groupname + " seedlingpositions1.zip", removedgroupdir + groupname + " seedlingpositions1.zip");
					filedelete = File.rename(groupdir + "seeds sorted X coordinates.tsv", removedgroupdir + "seeds sorted X coordinates.tsv");
					filedelete = File.rename(groupdir + "seeds sorted Y coordinates.tsv", removedgroupdir + "seeds sorted Y coordinates.tsv");
					filedelete = File.rename(groupdir + groupname + " preskeletonize.tif", removedgroupdir + groupname + " preskeletonize.tif");
					filedelete = File.rename(groupdir + "firstslice.tif", removedgroupdir + "firstslice.tif");
					filedelete = File.rename(groupdir + "secondarypoints.zip", removedgroupdir + "secondarypoints.zip");
					if (freshstart) {
						filedelete = File.rename(groupdir + groupname + " rootgrowthdetection.tif", removedgroupdir + groupname + " rootgrowthdetection.tif");
						filedelete = File.rename(groupdir + groupname + " rootgrowthmeasurement.tsv", removedgroupdir + groupname + " rootgrowthmeasurement.tsv");
						filedelete = File.rename(groupdir + groupname + " germination analysis.tsv", removedgroupdir + groupname + " germination analysis.tsv");
						filedelete = File.delete(groupdir);
					}
				}
			if (freshstart)
				filedelete = File.delete(platedir);
			}
		}
		freshstart = false; 
		//turns it back to false so essential output is not deleted at end of macro
		selectWindow("Log");
		save(removedfilesdir + File.separator + "Log.txt");
	}
}
```
