## Supplementary File 6 for "SPIRO – the automated Petri plate imaging platform designed by biologists, for biologists": release.yml

```
name: Make release

on:
  push:
    tags:
      - 'v*.*'

jobs:
  
  MacOS:
    runs-on: macos-latest
    
    steps:
    - uses: actions/checkout@v2
    - name: Get tag
      id: tag
      run: |
        echo ::set-output name=tag::${GITHUB_REF#refs/tags/}
    - name: Set up Python
      uses: actions/setup-python@v1
      with:
        python-version: '3.8'
    - name: Install requirements
      run: |
        python -m pip install -r requirements.txt
        python -m pip install pyinstaller
    - name: Create MacOS app
      run: |
        pyinstaller --add-binary='/System/Library/Frameworks/Tk.framework/Tk':'tk' --add-binary='/System/Library/Frameworks/Tcl.framework/Tcl':'tcl' --osx-bundle-identifier slu.spiro.customizer --windowed --noconfirm -n 'SPIRO Assay Customizer' -i resources/spiro-assay-customizer.icns src/main.py
    - name: Build Zip
      run: |
        cd dist
        zip -r SPIRO_Assay_Customizer_Mac_${{ steps.tag.outputs.tag }}.zip "SPIRO Assay Customizer.app"
    - name: Upload release
      uses: actions/upload-artifact@v2
      with:
        name: SPIRO Assay Customizer Mac ${{ steps.tag.outputs.tag }}.zip
        path: dist/*.zip

  Windows:
    runs-on: windows-latest
    
    steps:
    - uses: actions/checkout@v2
    - name: Get tag
      id: tag
      shell: bash
      run: |
        echo ::set-output name=tag::${GITHUB_REF#refs/tags/}
    - name: Set up Python
      uses: actions/setup-python@v1
      with:
        python-version: '3.8'
    - name: Install requirements
      run: |
        python -m pip install -r requirements.txt
        python -m pip install pyinstaller
    - name: Create Windows exe
      run: |
        pyinstaller --windowed --noconfirm -n "SPIRO Assay Customizer" -i resources/spiro-assay-customizer.ico -F src/main.py
    - name: Build Zip
      run: |
        cd dist
        7z a SPIRO_Assay_Customizer_Windows_${{ steps.tag.outputs.tag }}.zip "SPIRO Assay Customizer.exe"
    - name: Upload release
      uses: actions/upload-artifact@v2
      with:
        name: SPIRO Assay Customizer Windows ${{ steps.tag.outputs.tag }}.zip
        path: dist/*.zip

  Linux:
    runs-on: ubuntu-16.04
    
    steps:
    - uses: actions/checkout@v2
    - name: Get tag
      id: tag
      run: |
        echo ::set-output name=tag::${GITHUB_REF#refs/tags/}
    - name: Set up Python
      uses: actions/setup-python@v1
      with:
        python-version: '3.8'
    - name: Install requirements
      run: |
        python -m pip install -r requirements.txt
        python -m pip install pyinstaller
    - name: Create Linux executable
      run: |
        pyinstaller --noconfirm -n 'SPIRO Assay Customizer' --onefile src/main.py
    - name: Run GZip
      run: |
        gzip dist/'SPIRO Assay Customizer'
    - name: Upload release
      uses: actions/upload-artifact@v2
      with:
        name: SPIRO Assay Customizer Linux ${{ steps.tag.outputs.tag }}.gz
        path: dist/*.gz

  Release:
    needs: [MacOS, Linux, Windows]
    runs-on: ubuntu-latest
    
    steps:
    - name: Check out
      uses: actions/checkout@v2
    - name: Get tag
      id: tag
      run: |
        echo ::set-output name=tag::${GITHUB_REF#refs/tags/}
    - name: Download MacOS release
      uses: actions/download-artifact@v2
      with:
        name: SPIRO Assay Customizer Mac ${{ steps.tag.outputs.tag }}.zip
        path: releases/
    - name: Download Windows release
      uses: actions/download-artifact@v2
      with:
        name: SPIRO Assay Customizer Windows ${{ steps.tag.outputs.tag }}.zip
        path: releases/
    - name: Download Linux release
      uses: actions/download-artifact@v2
      with:
        name: SPIRO Assay Customizer Linux ${{ steps.tag.outputs.tag }}.gz
        path: releases/
    - name: Create release
      id: create_release
      uses: actions/create-release@v1
      env:
        GITHUB_TOKEN: ${{ secrets.GITHUB_TOKEN }}
      with:
        tag_name: ${{ steps.tag.outputs.tag }}
        release_name: ${{ steps.tag.outputs.tag }}
        draft: false
        prerelease: false
    - name: Upload MacOS release asset
      uses: actions/upload-release-asset@v1
      env:
        GITHUB_TOKEN: ${{ secrets.GITHUB_TOKEN }}
      with:
        upload_url: ${{ steps.create_release.outputs.upload_url }}
        asset_path: releases/SPIRO Assay Customizer Mac ${{ steps.tag.outputs.tag }}.zip
        asset_name: SPIRO Assay Customizer Mac ${{ steps.tag.outputs.tag }}.zip
        asset_content_type: application/zip
    - name: Upload Windows release asset
      uses: actions/upload-release-asset@v1
      env:
        GITHUB_TOKEN: ${{ secrets.GITHUB_TOKEN }}
      with:
        upload_url: ${{ steps.create_release.outputs.upload_url }}
        asset_path: releases/SPIRO Assay Customizer Windows ${{ steps.tag.outputs.tag }}.zip
        asset_name: SPIRO Assay Customizer Windows ${{ steps.tag.outputs.tag }}.zip
        asset_content_type: application/zip
    - name: Upload Linux release asset
      uses: actions/upload-release-asset@v1
      env:
        GITHUB_TOKEN: ${{ secrets.GITHUB_TOKEN }}
      with:
        upload_url: ${{ steps.create_release.outputs.upload_url }}
        asset_path: releases/SPIRO Assay Customizer.gz
        asset_name: SPIRO Assay Customizer.gz
        asset_content_type: application/gzip
```
