## Supplementary File 6 for "SPIRO – the automated Petri plate imaging platform designed by biologists, for biologists": README.md

# SPIRO Assay Customizer
SPIRO Assay Customizer is a simple GUI for managing output from [SPIRO Assays](https://github.com/jiaxuanleong/SPIRO.Assays). The GUI allows adding new groups, rearranging seeds/seedlings between groups, and removing seeds/seedlings from analysis. It can handle output files from both the Germination and Root Growth assays.

This application is cross-platform, and should run on MacOS, Windows, common Unixes, and Linux.

## Customize Assay

The Customize Assay mode allows you to reassign seedlings to groups, and to exclude seedlings or groups of seedlings from analysis.

<img width="602" alt="image" src="https://user-images.githubusercontent.com/6480370/86357062-ce289280-bc5c-11ea-816f-a656977b224c.png">

## Merge Assays

In the Merge Assays mode, analysis results from multiple runs may be joined together in order to be jointly analyzed.

<img width="683" alt="image" src="https://user-images.githubusercontent.com/6480370/80723195-9d16cf00-8b00-11ea-9d25-f47389d92d13.png">
