## Supplementary File 6 for "SPIRO – the automated Petri plate imaging platform designed by biologists, for biologists": assay_editor.py

```
#!/usr/bin/env python
import sys
import PySimpleGUI as sg
import pandas as pd
import numpy as np
from icon import icon


def file_picker():
    """shows a file picker for selecting a postQC.tsv file. Returns None on Cancel."""
    chooser = sg.Window('Choose file', [
        [sg.Text('Filename')],
        [sg.Input(), sg.FileBrowse(key='-FILE-', file_types=(('PostQC TSV files', '*.postQC.tsv'),))],
        [sg.OK(), sg.Cancel()]], icon=icon)

    event, values = chooser.read()
    if event in (None, 'Cancel'):
        chooser.close()
        return None
    elif event == 'OK':
        chooser.close()
        return values['-FILE-']


def postqc_window(uid_groups, avail_groups):
    """main interface. uid_groups is a list of [UID, Group] combinations.
       avail_groups is a list of the available groups. returns the main window object."""
    table_height = min(25, len(uid_groups))
    mgmt_layout = [[sg.B('Add New Group', key='Add')],
                   [sg.B('Assign Seedling to Group', key='Change'),
                    sg.B('Exclude Seedling from Analysis', key='Exclude')]]
    layout = [
        [sg.Table(values=uid_groups, headings=['UID', 'Group'], display_row_numbers=False,
                  auto_size_columns=True, num_rows=table_height, key="-COMBOS-"),
            sg.Table(values=avail_groups, headings=['Available groups', ], display_row_numbers=False,
                     auto_size_columns=True, num_rows=table_height, key="-GROUPS-",
                     select_mode=sg.TABLE_SELECT_MODE_BROWSE)],
        [sg.Frame('Seedling and Group Management', layout=mgmt_layout)],
        [sg.Sizer(h_pixels=120), sg.B('Write PostQC File', key='Write'), sg.B('Exit')]]
    return sg.Window('SPIRO Assay Customizer', layout, grab_anywhere=False, icon=icon)


def get_uid_groups(df):
    """gets the unique uids and groups in the specified dataframe. returns a
       tuple of uid/group combos (list) and the unique groups (list)."""
    uids = pd.unique(df['UID'])

    # XXX: there is a better wya of doing this
    uid_groups = list()
    groupset = set()
    for uid in uids:
        group = str(df[df['UID'] == uid].iloc[0, 1])
        if group == 'nan':
            # sg tables act really weird if you feed them nans
            group = 'NA'
        uid_groups.append([uid, group])
        groupset.add(group)

    # needs to be list of lists
    groups = sorted(([x, ] for x in groupset), key=lambda x: x[0])
    
    return uid_groups, groups


def start_editor(file=None):
    if file is None:
        file = file_picker()

    if file is None:
        # no file was provided in file picker
        sys.exit()
    elif not file.endswith('.postQC.tsv'):
        sg.Popup('The file must be a postQC.tsv file.', icon=icon)
        sys.exit()
    else:
        try:
            df = pd.read_csv(file, sep='\t', engine='python')
        except OSError as e:
            sg.Popup("Couldn't read file: " + e.strerror, icon=icon)
            sys.exit()

    # get uid/group combos, and save a working copy in a dataframe
    # groups can just be a list
    uid_groups, groups = get_uid_groups(df)
    uid_groups_df = pd.DataFrame(uid_groups)

    # display main window
    window = postqc_window(uid_groups, groups)

    # main loop
    while True:
        event, values = window.read()
        if event in (None, 'Exit'):
            break
        elif event == 'Change' and len(values['-GROUPS-']) > 0 and len(values['-COMBOS-']) > 0:
            uid_groups_df.loc[values['-COMBOS-'], 1] = groups[values['-GROUPS-'][0]]
            uid_groups = uid_groups_df.values.tolist()
            window['-COMBOS-'].Update(values=uid_groups)
        elif event == 'Write':
            for (uid, group) in uid_groups:
                df.loc[df['UID'] == uid, 'Group'] = group
            try:
                df.to_csv(file, sep='\t', index=False)
            except OSError as e:
                sg.Popup('Unable to write file ' + e.filename + ': ' + e.strerror, icon=icon)
        elif event == 'Add':
            window.Hide()
            newgroup = sg.popup_get_text('Name of new group', 'SPIRO Assay Customizer', icon=icon)
            if isinstance(newgroup, str):
                groups.append([newgroup, ])
                groups.sort(key=lambda x: x[0])
                window['-GROUPS-'].Update(values=groups)
            window.UnHide()
        elif event == 'Exclude':
            uid_groups_df.loc[values['-COMBOS-'], 1] = 'NA'
            uid_groups = uid_groups_df.values.tolist()
            window['-COMBOS-'].Update(values=uid_groups)

    window.close()
```
