## Supplementary File 6 for "SPIRO – the automated Petri plate imaging platform designed by biologists, for biologists": main.py

```
#!/usr/bin/env python
import sys
import PySimpleGUI as sg
import merge
import assay_editor
from icon import icon

sg.set_options(auto_size_buttons=True, font='Any 12')
sg.ChangeLookAndFeel('Dark')

if len(sys.argv) == 2:
    assay_editor.start_editor(file=sys.argv[1])
elif len(sys.argv) > 2:
    sg.Popup('Assay editor can only work on one file at a time. Please merge files first.',
             title='SPIRO Assay Customizer', icon=icon)
    sys.exit()
else:
    layout = [
        [sg.T('What do you want to do?')],
        [sg.B('Customize Assay', key='Customize'), sg.B('Merge Assays', key='Merge')]]
    window = sg.Window('SPIRO Assay Customizer', layout=layout, icon=icon)

    event, values = window.read()

    if event == 'Customize':
        window.close()
        assay_editor.start_editor()
    elif event == 'Merge':
        window.close()
        merge.start_merge()
```
