## Supplementary File 6 for "SPIRO – the automated Petri plate imaging platform designed by biologists, for biologists": merge.py

```
#!/usr/bin/env python
import sys
import os
import PySimpleGUI as sg
import pandas as pd
from icon import icon


def merge_window():
    """window for merging experiments. will deal with either root growth or germination data,
       but not both at the same time."""
    # the table of experiments
    explist = [
        [sg.Table(headings=['Experiments', ], display_row_numbers=False,
                  auto_size_columns=False, values=list([]), def_col_width=65,
                  num_rows=8, key='-EXPERIMENTS-', select_mode=sg.TABLE_SELECT_MODE_BROWSE)],
        [sg.B('Remove experiment', key='Remove')]]
    # rest of ui
    addexp = [
        [sg.T('Folder:'), sg.I(key='exp', size=(45, 1)), sg.FolderBrowse(), sg.B('Add')]]
    layout = [
        [sg.Frame('Merge experiments', layout=explist)],
        [sg.Frame('Add experiment', layout=addexp)],
        [sg.B('Perform Merge', key='Merge'), sg.B('Exit')]]
    return sg.Window('Merge', layout=layout, grab_anywhere=False, icon=icon)


def save_germination(g_df, log_df):
    """save merged germination data to a new folder"""
    savelayout = [
        [sg.Text('Choose location for merged results')],
        [sg.T('Folder:'), sg.I(key='dir', size=(45, 1)), sg.FolderBrowse()],
        [sg.OK(), sg.Cancel()]]
    save_window = sg.Window('Save merged data', layout=savelayout, icon=icon)
    event, values = save_window.read()
    
    if event in (None, 'Cancel'):
        save_window.close()
        sg.Popup('Merged data not saved.', title='SPIRO Assay Customizer', icon=icon)
        return
    elif event == 'OK':
        save_window.close()
        dir = os.path.join(values['dir'], 'Results', 'Germination')
        os.makedirs(dir, exist_ok=True)
        g_file = os.path.join(dir, 'germination.postQC.tsv')
        log_file = os.path.join(dir, 'germination.postQC.log.tsv')
        if os.path.exists(g_file):
            sg.Popup('Selected directory already contains germination data. Aborting.',
                     title='SPIRO Assay Customizer', icon=icon)
            return
        try:
            g_df.to_csv(g_file, index=False, na_rep='NA', sep='\t')
            log_df.to_csv(log_file, index=False, na_rep='NA', sep='\t')
        except OSError as e:
            sg.Popup('Unable to write file ' + e.filename + ': ' + e.strerror, icon=icon)


def save_rootgrowth(r_df, ps_df):
    """save merged root growth data to a new folder"""
    savelayout = [
        [sg.Text('Choose location for merged results')],
        [sg.T('Folder:'), sg.I(key='dir', size=(45,1)), sg.FolderBrowse()],
        [sg.OK(), sg.Cancel()] ]
    save_window = sg.Window('Save merged data', layout=savelayout, icon=icon)
    event, values = save_window.read()
    
    if event in (None, 'Cancel'):
        save_window.close()
        sg.Popup('Merged data not saved.')
        return
    elif event == 'OK':
        save_window.close()
        dir = os.path.join(values['dir'], 'Results', 'Root Growth')
        os.makedirs(dir, exist_ok=True)
        r_file = os.path.join(dir, 'rootgrowth.postQC.tsv')
        ps_file = os.path.join(dir, 'germination-perseed.tsv')
        for file in r_file, ps_file:
            if os.path.exists(file):
                sg.Popup('Selected directory already contains root growth data: ' + os.path.basename(file) +
                         '. Aborting.', title='SPIRO Assay Customizer', icon=icon)
                return

        try:
            r_df.to_csv(r_file, index=False, na_rep='NA', sep='\t')
            ps_df.to_csv(ps_file, index=False, na_rep='NA', sep='\t')
        except OSError as e:
            sg.Popup('Unable to write file ' + e.filename + ': ' + e.strerror,
                     title='SPIRO Assay Customizer', icon=icon)
        sg.Popup('Merged data saved.', title='SPIRO Assay Customizer', icon=icon)


def unlist(l):
    """returns a list of values from a list of lists of values"""
    return [x[0] for x in l]


def merge_experiments(exps):
    """takes a list of a lists of experiment directories and returns merged_germination,
       merged_germination_log, merged_rootgrowth, merged_perseed"""
    rtsv = lambda x: pd.read_csv(x, sep='\t')
    bail = lambda x: sg.Popup("File " + x + "doesn't exist, but is needed for merge. Aborting.", icon=icon)
    fail = None, None, None, None
    
    exps = unlist(exps)
    mode = os.path.basename(exps[0])
    if not all(mode == os.path.basename(x) for x in exps):
        sg.Popup('Cannot mix Germination and Root Growth assays, not merging.',
                 title='SPIRO Assay Customizer', icon=icon)
        return fail
    
    r_df = pd.DataFrame()
    g_df = pd.DataFrame()
    log_df = pd.DataFrame()
    ps_df = pd.DataFrame()

    if mode == 'Root Growth':
        for exp in exps:
            r_file = os.path.join(exp, 'rootgrowth.postQC.tsv')
            ps_file = os.path.join(exp, 'germination-perseed.tsv')
            for file in r_file, ps_file:
                if not os.path.exists(file):
                    bail(file)
                    return fail
            try:
                r_tsv = rtsv(r_file)
                ps_tsv = rtsv(ps_file)
            except OSError as e:
                sg.Popup("Couldn't open file " + e.filename + ": " + e.strerror, icon=icon)
                return fail
            r_df = pd.concat([r_df, r_tsv])
            ps_df = pd.concat([ps_df, ps_tsv])
        return None, None, r_df, ps_df
    elif mode == 'Germination':
        for exp in exps:
            g_file = os.path.join(exp, 'germination.postQC.tsv')
            log_file = os.path.join(exp, 'germination.postQC.log.tsv')
            for file in g_file, log_file:
                if not os.path.exists(file):
                    bail(file)
                    return fail
            try:
                g_tsv = rtsv(g_file)
                log_tsv = rtsv(log_file)
            except OSError as e:
                sg.Popup("Couldn't open file " + e.filename + ": " + e.strerror, icon=icon)
                return fail
            g_df = pd.concat([g_df, g_tsv])
            log_df = pd.concat([log_df, log_tsv])
        return g_df, log_df, None, None


def start_merge():
    window = merge_window()
    exps = list()

    while True:
        event, values = window.read()
        if event in (None, 'Exit'):
            break
        elif event == 'Add' and values['exp'] != '':
            if not values['exp'].endswith(('Germination', 'Root Growth')):
                sg.Popup('Selected directory must be named either "Germination" or "Root Growth"',
                         title='SPIRO Assay Customizer', icon=icon)
            else:
                exps.append([values['exp']])
                window['-EXPERIMENTS-'].Update(values=exps)
        elif event == 'Remove':
            del exps[values['-EXPERIMENTS-'][0]]
            window['-EXPERIMENTS-'].Update(values=exps)
        elif event == 'Merge':
            # commence merging
            g_df, log_df, r_df, ps_df = merge_experiments(exps)
            if g_df is not None:
                save_germination(g_df, log_df)
                sg.Popup('Merged germination data saved.', title='SPIRO Assay Customizer', icon=icon)
            elif r_df is not None:
                save_rootgrowth(r_df, ps_df)
                sg.Popup('Merged root growth data saved.', title='SPIRO Assay Customizer', icon=icon)
    window.close()
```
