## Supplementary Table 1 for "SPIRO – the automated Petri plate imaging platform designed by biologists, for biologists"

| | # | The components | Distributor | Catalog number | Quantity required for one SPIRO system |
|---|---|---|---|---|---|
| | | **Electronics** | | | |
| ☐ | 1 | Raspberry Pi 3B+ | Mouser | 358-RPI3-MODBP-BULK | 1 |
| ☐ | 2 | Power supply 5V 2.5A, micro USB | Mouser | 358-1101001000045 | 1 |
| ☐ | 3 | Adafruit TB6612 h-bridge dc-stepper motor driver breakout board | Mouser | 485-2448 | 1 |
| ☐ | 4 | Unipolar stepper motor NEMA-17, 12V, 0.35A | Mouser | 485-324 | 1 |
| ☐ | 5 | Power adapter 12V 3A wih with 2.1 power plug | Amazon | | 1 |
| ☐ | 6 | Female DC connector 5.5 x2.1mm | Amazon | | 1 |
| ☐ | 7 | SparkFun MOSFET Power Control Kit | Mouser | 474-COM-12959 | 1 |
| ☐ | 8 | Green LED strip 12V (note: must be possible to cut into 5-cm segments!) | Mouser | 828-OVQ12S30G7 | 1 meter |
| ☐ | 9 | 90 degree connectors for 8 mm LED strip + terminal LED strip connector | Amazon | | 4 corners + 1 terminal connector |
| ☐ | 10 | 2-pin 3.5 mm pitch PCB Mount Screw Terminal Connector | Mouser | 710-691214110002S | 1 |
| ☐ | 11 | Jumper cables 10 cm, female to female (Dupont wires) | Amazon | | 20 |
| ☐ | 12 | SD Memory Card Class 10 or higher, 32 GB or more | Mouser | 467-SDSDQAE-032G | 1 |
| ☐ | 13 | Heat shrink tubes, diameter 1 mm and 2 mm | Amazon | | 5 cm of each |
| ☐ | 14 | Wire end ferrules, 0.5 mm2, not insulated | Amazon | | 20 |
| ☐ | 15 | AWG 28 wires, longer than 40 cm | Amazon | | ca 3 m |
| ☐ | 16 | Spiral wire cover (Coiled cable protection) | Amazon | | ca 30 cm |
| | | **Camera** | | | |
| ☐ | 17 | Adafruit Flex Cable for Raspberry Pi Camera | Amazon | | 25-30 cm |
| | 18a | **SPIRO configuration with manual focus:** | | | |
| ☐ | | Raspberry Pi Camera v2 - 8 MP | Mouser | 485-3099 | 1 |
| | 18b | **SPIRO configuration with motorized focus:** | | | |
| ☐ | | Raspberry Pi Camera v2 - 8 MP | Mouser | 485-3099 | 1 |
| ☐ | | Arducam IMX219 8MP Auto Focus Camera Module, drop-in replacement for Raspberry Pi V2 Camera | Uctronics | SKU: B0182 | 1 |
| | | **Positional sensor** | | | |
| ☐ | 19 | Mini microswitch | Reichelt elektronik | MAR 1050.5202 | 1 |
| | | **Structural parts** | | | |
| ☐ | 20 | Aluminium profile T2020, **black** | Amazon | | 1x 30cm for the vertical rail, 1x ca 34 cm for the horizontal rail, 2x ca 8cm for each leg, 2x 14 cm for the LED frame mounting |
| ☐ | 21 | 2020 Aluminum Profile Slot Cover, black | Banggood | | 1 m |
| ☐ | 22 | 635zz - R-1950ZZ - 5972K263 Miniature Ball Bearings | Amazon | | 3 |
| ☐ | 23 | Rigid shaft coupling M5 to M5 | Amazon | | 1 |
| | | **Screws** | | | |
| ☐ | 24 | M2.5 x 12mm, button head | | | 2 |
| ☐ | 25 | M2.5 x 12mm, flat head countersunk, black | | | 2 |
| ☐ | 26 | M2.5 x 8mm, flat head countersunk | | | 8 |
| ☐ | 27 | M2.5 hex nuts | | | 12 |
| ☐ | 28 | M3 x 12 mm, button head, hex socket, black | | | 8 |
| ☐ | 29 | M3 x 20mm, flat head countersunk | | | 4 |
| ☐ | 30 | M3 hex nuts | | | 8 |
| ☐ | 31 | M5 x 8 mm, flat head countersunk, black | | | 1 |
| ☐ | 32 | M5 x 12 mm, flat head countersunk | | | 8 |
| ☐ | 33 | M5 x 8 mm screw, button head, hex socket, black | | | 21 |
| ☐ | 34 | M5 x 16 mm screw, button head, hex socket, black | | | 3 |
| ☐ | 35 | M5 x 16mm bolt, hex head | | | 5 |
| ☐ | 36 | M5 x 20mm bolt, hex head | | | 1 |
| ☐ | 37 | M5 T-nuts for T2020 aluminium profile | | | 35 |
| ☐ | 38 | M5 hex nuts | | | 3 |
| ☐ | 39 | M5 washer, OD 10 mm, flat | | | 6 |
| ☐ | 40 | M5 slide-in bracket for T2020 aluminium profile | | | 3 |
| ☐ | 41 | M5 x 11 mm , button head, hex socket, black | | | 6 |
| | | **Plastic for the 3D printer** | | | |
| ☐ | 42 | PLA filament, matte black | 3Dprinthings | Matteforge Pro Filament PLA 1.75 mm, black | ca 1.5 kg |
| ☐ | 43 | PLA filament, milky white | Amazon | Sunlu, PLA+, 1.75 mm, milky white / semi-transparent | ca 200 g |
| | | **Tools** | | | |
| ☐ | 44 | Wire stripper suitable for AWG 28 wires | Amazon | | |
| ☐ | 45 | Ferrule crimper suitable for 0.5 mm2 ferrules | Amazon | | |
| ☐ | 46 | Soldering iron & solder | Amazon | | |
| ☐ | 47 | Screw drivers for M2, M2.5 and M3 (cross and flat head) | | | |
| ☐ | 48 | Allen keys for M5 & M3 | | | |
| ☐ | 49 | A saw for aluminium profiles | | | |
| ☐ | 50 | Superglue | any would do | | |
| | | **Optional** | | | |

| | # | The components | Distributor | Catalog number | Quantity required for one SPIRO system |
|---|---|---|---|---|---|
| ☐ | 51 | Ethernet cable | Amazon | | 1 cable, the length will depend on the distance between the robot and ethernet port. we strongly recommend using flat ethernet cables to enable closing incubators door |
