## Supplementary Table 2 for "SPIRO – the automated Petri plate imaging platform designed by biologists, for biologists"

### Structural parts

| | .stl, .3mf, .f3d file names* | Rotation | Supports | Brim | Infill | | Printing | Components list |
|---|---|---|---|---|---|---|---|---|
| ☐ | Set 01. Alyona Minina. 2019. UHEI | X: 90° | Build plate | No | 20% | ● | Matte black filament, 135 g<br>Printing time: 13 h | 1. Motor hub<br>2. 2x bearing guides<br>3. 1x bearing guide with the mount for the mini microswitch<br>4. Mini microswitch housing<br>5. Mini microswitch pin (tiny dick)<br>6. Holder for the vertical rail<br>7. Holder for the LED frame rail<br>8. 2x bottom for a thumb screw<br>9. 2x cap for a thumb screw |
| ☐ | Set 02. Alyona Minina. 2019. UHEI | – | Build plate | No | 20% | ● | Matte black filament, 210 g<br>Printing time: 18.5 h | 1. Borg's nest<br>2. Lid for the Borg's nest<br>3. Lid for the DC connector<br>4. Holder for the SparkFun MOSFET power control kit |
| ☐ | Set 03a. Alyona Minina. 2019. UHEI | X: 90° | Build plate | No | 20% | ● | Matte black, 50 g<br>Printing time: 5 h | 1. Camera housing for Arducam<br>2. Lid for the camera housing<br>3. 2x bottom for a thumb screw<br>4. 2x cap for a thumb screw<br>5. 2x washer |
| ☐ | Set 03b. Alyona Minina. 2019. UHEI | X: 90° | Build plate | No | 20% | ● | Matte black, 25 g<br>Printing time: 3 h | 1. Camera housing for RPi camera<br>2. Lid for the camera housing<br>3. 2x bottom for a thumb screw<br>4. 2x cap for a thumb screw<br>5. Stabilizer for the camera<br>6. Focusing tool |
| ☐ | Set 04. Alyona Minina. 2019. UHEI | X: 270° | Build plate | No | 20% | ● | Matte black, 380 g<br>Printing time: 28.5 h | 1. Cube-shaped stage with holders for Petri plates |
| ☐ | Set 05. Alyona Minina. 2019. UHEI | Z: 345° | Everywhere | Yes | 50% | ● | Matte black, 260 g<br>Printing time: 23 h<br>Notes: Please ignore messages about toolpath being detected outside the perimeter. | 1. The right half of the light screen |
| ☐ | Set 06. Alyona Minina. 2019. UHEI | Z: 355° | Everywhere | Yes | 50% | ● | Matte black, 230 g<br>Printing time: 20.5 h<br>Notes: Please ignore messages about toolpath being detected outside the perimeter. | 1. The left half of the light screen |
| ☐ | Set 07. Alyona Minina. 2019. UHEI | Z: 50° | Everywhere | Yes | 20% | ● | Matte black, 140 g<br>Printing time: 14 h<br>Notes: Please ignore messages about toolpath being detected outside the perimeter. | 1. The right half of the LED frame |
| ☐ | Set 08. Alyona Minina. 2019. UHEI | Z: 50° | Everywhere | Yes | 20% | ● | Matte black, 130 g<br>Printing time: 12.5 h<br>Notes: Please ignore messages about toolpath being detected outside the perimeter. | 1. The left half of the LED frame |
| ☐ | Set 09. Alyona Minina. 2019. UHEI | Z: 50° | – | Yes | 30% | ○ | Transparent/milky white, 35 g<br>Printing time: 2.5 h | 1. Left diffuser<br>2. Bottom left diffuser<br>3. Bottom right diffuser |
| ☐ | Set 10. Alyona Minina. 2019. UHEI | Z: 40° | – | Yes | 30% | ○ | Transparent/milky white, 20 g<br>Printing time: 1.5 h | 1. Top diffuser |
| ☐ | Set 11. Alyona Minina. 2019. UHEI | Z: 50° | – | Yes | 30% | ○ | Transparent/milky white, 20 g<br>Printing time: 1.4 h | 1. Right diffuser |

**Notes**

As of today, printing structural parts has been tested using PLA filaments only. Required total amount of black matt PLA = 1685 g

Required total amount of milky white PLA = 75 g

Total estimated time of printing = 159.5 h

### Accessories

| | .stl, .3mf, .f3d file names* | Rotation | Supports | Brim | Infill | | Filament | Additional info |
|---|---|---|---|---|---|---|---|---|
| | Anti-reflection lids for square 12 cm plates | Y: 180° | - | No | 20% | ◐ | PETG black and white, 41 g<br>Printing time: 6 h | Printing instructions: For the sake of durability, we strongly recommend using PETG plastic.<br>The lid should be printed in two colors: the inner pane of the lid should be black and ironed to provide evenly contrasting background for seeds and seedlings, while the rims of the lid should be white to reduce shading and reflect the light back into the plate. |

| | | | | | | | | |
|---|---|---|---|---|---|---|---|---|
| | Anti-reflection lids for round 9 cm plates | Y: 180° | - | No | 20% | ◐ | PETG black and white, 18 g<br>Printing time: 3.5 h | Ironing during printing might result in an unwanted shiny finish of the surface, this can be avoided by manually lowering down the nozzle temperature during printing the ironed level. In our experience, using 190–200 C nozzle temperature results in a matte finish while using PETG filament.<br>The .3mf file contains information required for printing using a Prusa i3 MK3/S printer and can be modified for any other printer. |
| | Seed plating guide for square 12 cm plates | - | - | No | 15% | | Any filament, 59 g<br>Printing time: 14.5 h | |
| | Seed plating guide for round 9 cm plates | Y: 270° | - | No | 15% | | Any filament, 51 g<br>Printing time: 9.5 h | |
| | Seed plating guides and insets for multi-well Petri plates | - | - | No | - | | - | For more information please refer to Thingiverse: https://www.thingiverse.com/thing:4196441 |
| | * Supplementary file 1.zip | | | | | | | |
